# A Latent Variable Model for Evaluation of Disparate Ratings of Stem Cell Colonies by Two Experts

**DOI:** 10.1101/746057

**Authors:** Michael Halter, Steven Lund, Adele Peskin, Ya-Shian Li-Baboud, Peter Bajcsy, Oleg Aulov, Daniel J. Hoeppner, Joshua G. Chenoweth, Suel-Kee Kim, Ronald D. McKay, Anne L. Plant

## Abstract

The visual inspection of pluripotent stem cell colonies by microscopy is widely used as a primary method to assess the quality of the preparations and degree of pluripotency. The lack of ground truth and the possible inconsistency of evaluations from multiple experts within and between stem cell laboratories are sources of uncertainty about the state of the cells, the reproducibility of preparations, and the efficiency of expansion protocols. To examine how to evaluate the level of confidence one has in disparate rating from experts, we explored a statistical method for assessing the differences in ratings of pluripotent stem cells by two different experts. Two experts rated phase contrast microscope images of human embryonic stem cell (hESC) colonies on a scale of 1 (poor) to 5 (maximum pluripotency character) but agreed with one another only 48% of the time. To assess whether experts used similar criteria to rate colonies, we developed custom image feature algorithms based on the stated visual criteria provided by the experts for selection of colonies. These features, plus others, were then used to develop pluripotency scoring algorithms trained to reflect ratings of both experts. We treated expert ratings as inexact indicators of a continuous pluripotency score and considered the inconsistency between expert ratings in developing our models. The model suggests that the two experts use somewhat different scales for discriminating between colony quality. Covariance analysis indicated that both experts use features that are not included in the model. Two image features, colony perimeter and a feature based on texture, were the most important for both experts for predicting the ratings. Interestingly, colony perimeter was not one of the expert-provided criteria for rating colonies, showing that this modeling approach allowed identification of features that the experts were not aware they were using. A linear model based on both experts identified each expert’s top-rated colonies as well as, or better than, the ratings of the other expert, as indicated by receiver operator characteristic curve analysis. By providing an understanding of the differences and similarities in disparate sets of expert ratings, this analysis helps to establish confidence in the ratings and the criteria for ratings, even when the experts disagree.

## INTRODUCTION

The commercial significance of stem cell-based therapies and diagnostics has led to increased interest in minimizing variability in culturing and characterizing these cells in research as well as manufacturing settings. Unambiguous metrics for evaluation of stem cell colonies do not existconfidence in assessing colonies. There is uncertainty about the adequacy of commonly used pluripotency markers for the identification of colonies with immunofluorescence [1](and references therein), in part because of heterogeneity in the expression of markers in cells and insufficient understanding in the field of how to interpret the observed heterogeneity to classify individual cells. Morphology assessment is often preferred for evaluating stem cell colonies, and it is often thought that visual assessment provides indications of desirable properties of pluripotency, viability, and other characteristics that are putatively recognized by an expert observer using phase contrast microscopy. Morphology assessment by experts is, of course, also ambiguous because different experts often have different opinions and may even infer characteristics that are not explicit in the images of cells being examined, such as how do the cells look at this point in time in comparison to some other time. The lack of ground truth and the ambiguity in expert opinion makes it difficult to be confident about any individual’s assessment. In this study we show how to use a combination of expert opinions to add to the confidence and understanding of the stem cell colony ratings even when those expert opinions differ.

We have used descriptive explanations of visual cues from stem cell experts to provide insight into the colony characteristics they use to identify desirable colonies. We have designed image feature algorithms that aim to numerically describe those characteristics and scoring algorithms that attempt to represent the assessments of the stem cell experts as an explicit function of those image features.

We provide here a set of 14 computational features inspired by the experts’ descriptions as potential components of signatures of pluripotent stem cell colonies. Many examples of analyzing cell and tissue images by translating an expert’s assessment of biological response into mathematical expressions have been reported [2–4], and we have considered other common features to our colony image data in addition to expert-inspired features. A number of machine learning approaches have been used to develop automated methods for classifying stem cell colonies, including unsupervised clustering approaches [5], and supervised methods using neural networks [6, 7], pattern recognition [7], and morphology-based feature classification [8], as well as recent work using deep learning [9]. Here, we want to know what features of colonies experts are using, but instead of a traditional classification strategy, we used a probabilistic approach to convert expert-dependent ratings to provide a basis for a variable pluripotency scoring range. Unlike typical classification schemes, this approach uses the information provided by the ordering of the experts’ rating when fitting and enables the estimation of what proportion of evaluation patterns shared by both experts are captured by a given model. To our knowledge, this work represents the first application of latent variable modeling [10] (LVM) to the automated evaluation of stem cell pluripotency.

Colony ratings by the two stem cell experts were not always consistent. This lack of ‘ground truth’ is a common problem with assessing uncertainty in analysis of biological data in general, and care must be given when interpreting subjective expert assessments. In this application, we abide by the following tenets for expert ratings when developing and assessing the performance of the image feature-based pluripotency scoring algorithms: 1) expert ratings do not correspond to categorical states of nature (i.e., ratings are not a literal ground truth); 2) expert ratings are not exactly precise in that each rating may correspond to a range of perceived relative pluripotency (e.g., colony A is preferred over colony B, but both received a rating of 4); 3) the meaning of a given rating value can differ between experts (e.g., experts could agree on the relative pluripotency of a given colony compared to other colonies, but assign it different rating values); 4) experts can disagree in their perceived pluripotency of a given colony (e.g., experts disagree whether colony A appears more desirable than colony B).

Collectively, these considerations dictate that the expert ratings should be recognized as ordinal and expert-specific, rather than categorical or continuous, and consequently latent variable modeling, as opposed to either classification or ordinary regression provide the appropriate modeling framework for these data. We address these perspectives by modeling each rating value from each expert as corresponding to a range, rather than an exact value, on a latent, continuous pluripotency scoring scale. The predicted score for each colony is given by a function of the image feature values for the colony (e.g., weighted combination or random forest) and evaluated through comparison with the scoring range(s) corresponding to the rating value(s) provided by the expert(s).

## MATERIALS AND METHODS

### Human pluripotent stem cell culture

The hESC H9 line was routinely maintained in culture with replacement by frozen stocks every 30–40 passages. Cells are expanded on mouse embryonic fibroblast (MEF) feeder cells and routinely tested for pluripotency characteristics (by expression of OCT4, SOX2, NANOG and SSEA-3/4) and their potential to differentiate into multiple germ layers (by expression of eomesodermin and nuclear export signal), as indicated by specific antibody staining. The MEF feeder cells were from CF-1 mice at embryonic day 13.5. The MEFs were passaged at least three times before use with hESCs to minimize residual non-fibroblast cells and were used as feeders between passage 3 and 4. The feeder cells were plated onto gelatinized tissue culture plates at the density of 44,000 cells/cm2 and irradiated with gamma irradiation. The hESCs were cultured on the irradiated MEF cells in DMEM/F12 medium (Thermo Fisher) supplemented with 20% Knockout SR (Thermo Fisher), 0.1mM nonessential amino acids (Thermo Fisher), 0.1 mM β-mercaptoethanol (Sigma-Aldrich), and 5 ng/ml human basic fibroblast growth factor (hbFGF, R&D Systems). Cultures of hESCs were routinely passaged at approximately 4 - 5 day intervals when the colonies reached an average size of 400-500 cells by visual inspection. For passaging of uniform-sized colonies, cells were treated with 1 mg/ml type IV collagenase (Thermo Fisher) for 5 min and the colonies were cut by StemPro EZPassage Tool (Thermo Fisher). The clumps of 50-100 cells were selectively collected by using 100 μm Cell Strainer (Corning) and washed to remove residual collagenase and then cultured on the fresh feeders.

### Image data collection

Three 100mm dishes containing stem cell colonies were used for the study. The cultures were fixed using 4% paraformaldehyde for 10 min prior to imaging. To test whether the fixation perturbed the appearance of the colonies, images were acquired from colonies before and after fixation and no observable changes were noted. The phase contrast image data was acquired on a Zeiss 135 TV microscope (Carl Zeiss USA, Thornwood, NY) with a Zeiss 10X/0.3NA Ph1 objective. The microscope was equipped with a CoolSNAP HQ2 CCD camera (Photometrics, Tucson, Arizona) and a motorized stage. Stage, filters and shutters were controlled by the Zeiss Axiovision software. The stage was programmed to move from field to field, 22 fields horizontally and 24 fields vertically with an overlap of adjacent fields of 10%. The total imaged area was approximately 2.3 cm x 2.1 cm. The fields were stitched using the Zeiss Axiovision software. A reference material for alignment of phase rings was developed and used to facilitate reproducible imaging conditions and assure the quality of the images. The material is composed of an array of micron-size features of polydimethylsiloxane (PDMS). An image of this patterned material is shown in Supplemental Information 1, Supplemental Figure 1 with a description of the procedure for collecting phase contrast images in which the material is used.

### Image processing

Prior to colony rating by experts, image objects that corresponded to colonies were identified and segmented using an automated macro routine in ImageJ (see Supplemental Information 1, section II) to visually separate the colonies from the background and from the surrounding MEF feeder cells. The macro blurs the images, applies the rolling ball algorithm to increase the intensity difference between larger colonies and the MEF layer, and then applies the ImageJ AutoThreshold function to identify colony objects. Objects smaller than 50,000 pixels (52,000 µm^2^) were not presented to the experts.

A separate algorithm was implemented for identifying the colony margin. A binary image from pixels that have both a high local mean intensity (I_mean_local_), and a high local standard deviation (SD_local_), computed over a 20×20 pixel neighborhood (21.2 µm x 21.2µm) was created. The mean of I_mean_local_ was computed over the entire image as I_mean_image_ and the mean and standard deviation of SD_local_ were computed over the entire image as SD_mean_image_ and SD_SD_image_, respectively. Pixels with I_mean_local_ > (I_mean_image_ + 0.5 *SD_image_) and SDlocal > (SD_mean_image_ + SD_SD_image_) were used to create a binary mask image. A morphological hole-filling operation was performed on the resulting binary image to remove internal holes from the contiguous objects. Colony objects were separated using the Fogbank algorithm [11], in which, once cell boundaries are located, each internal pixel is assigned to a cell based on its geodesic distance from the cell boundaries, i.e. the path from pixel to boundary must be internal to the pixel cluster. The details of the segmentation steps are presented in Peskin et al. [12], and the segmentation algorithm is available at [https://gitlab.nist.gov/gitlab/peskin/stem_cell_segmentation].

### Rating of colonies by experts

Colony images were examined and rated by two experts independently. Colonies were given subjective ratings by each expert on a scale from 5 (for the best pluripotent colonies), to 1 (those with the least pluripotent character). Rules for ratings were discussed and agreed on by the two experts, and these rules are described in Table 1. Not all colonies were provided rating by both experts. Experts did not rate a colony if, for example, the expert was uncertain about how to rate the colony. Expert 1 rated 450/480 colonies, expert 2 rated 476/480 colonies and 449/480 colonies were rated by both experts. The dataset of 480 colony images shown to raters is displayed in Supplemental Information 2 and the modeling was based on 477 colonies that were rated by at least one expert.

**Table 1.** Characteristics that experts agreed on for rating colonies

| **Rating** | **Description** |
| --- | --- |
| 5 | (a)distinct margins, homogeneous phase-bright internal cells, small internuclear distance, epithelial characteristics (elongated cells throughout). (b)distinct margins, homogeneous phase-bright internal cells, small internuclear distance, epithelial characteristics at margin only. |
| 4 | Intermediate |
| 3 | Distinct margins, heterogeneous phase bright/phase dark cells, variable inter-nuclear distance. |
| 2 | Intermediate |
| 1 | Fully differentiated, indistinct margins, heterogeneous internal cell types, large internuclear distance. |

### Image analysis features

Custom algorithms were developed to mathematically represent visual cues that experts used and are listed in Table 2. These 11 features were supplemented with the features perimeter, circularity and area, and the 14 image analysis features are further described in Supplemental Information 1, section III and in reference [12]. Because of the apparent importance of the colony margin, the features were developed based on three regions: the center region of the colony, the region immediately inside of the margin of the colony, and the region just outside the colony margin (as shown Supplemental Figure 2). Additional Haralick and wavelet features were applied in some models (see Supplemental Information1, section 3 for more details). A number of features, particularly wavelet features, are correlated, and an analysis of feature correlations is provided in Supplemental Figure 3.

**Table 2.** Colony attributes, interpretations for custom algorithm development, and feature names

| **Colony Attribute** | **Interpretations for algorithm development** | **Feature name** |
| --- | --- | --- |
| distinct margins | Differences between margin and outside of colony; homogeneity of margin | Edge quality, Edge contrast, Fgbg |
| homogeneous phase-bright internal cells | Contrast in pixel intensities; homogeneity in center; texture in center | Local std, Entropy, Hg1, Hg2, AreaRatio |
| small internuclear distance | Dark pixels surrounded by bright pixels relative to total colony pixels | Holes/Area |
| epithelial characteristics | Homogeneity of pixel contrast immediately inside and immediately outside the colony margin | E1, E2 |
| colony size and shape | The number of pixels in each segmentation mask; the number of pixels at the margin of each colony; $4.0 \cdot \pi \cdot \text{Area} / \text{Perimeter}^2$ | Area, Perimeter, Circularity |

### Model development and evaluation

We are seeking deterministic functions of image feature values that when applied to each of the colonies in our dataset mimic the ordering of preference indicated by the expert ratings. We refer to a candidate function as a scoring algorithm (i.e., a scoring function produces a predicted pluripotency score for each colony as a function of the image feature values).

Several scoring algorithm approaches were used. Primary emphasis was on linear models, although random forests were used to see if a non-linear scoring algorithm improved performance. Algorithms were initially developed using the 14 expert-inspired image features and later revisited using an expanded feature set that included 42 additional Haralick and wavelet image features.

Our modeling approach uses a bivariate normal distribution to represent deviations of each expert from the pluripotency scores predicted by a scoring algorithm. The extent to which a given scoring algorithm is discordant with the ratings provided by a given expert is represented by a variance parameter for that expert. The extent to which the experts agree in their disagreements with the scoring algorithm is represented by a covariance parameter. Collectively, the two variance parameters and the covariance parameter provide the elements of the variance-covariance matrix for the bivariate normal distribution.

In this analysis, the experts’ ratings are treated as ordinal responses. The provided numbers indicate the relative preference of the expert, but not the magnitude of difference; e.g., 1,2,3,4,5 could be respectively translated as worst < worse < intermediate < better < best. Because the ratings provided by the experts are clearly discrete (and therefore do not resemble a sample from a normal distribution), we treat the ratings of the experts as representing a range of possible pluripotency scores (i.e., ratings are treated as censored scores). Initial fitting assumed that each rating corresponded to the range of scores for which that rating was the nearest available value (i.e., a rating of 1 indicates the score was less than 1.5; a rating of 2 indicates the score was between 1.5 and 2.5; …; a rating of 5 indicates the score was greater than 4.5). Subsequent modeling efforts allowed the spacing to vary for each rating and expert. For example, a rating of 2 by expert 1 may correspond to a score between 1.05 and 2.3, but the exact value is otherwise unknown. A rating of 2 by expert 2 may correspond to a score between 1.55 and 2.62. The intervals for each expert are identified by 4 thresholds, which are model parameters optimized during model fitting.

A brief technical description of our implementation of LVM is as follows: let *y*_*ij*_ represent the rating provided by expert *j* when evaluating colony *i*. We assume that although the experts provide a categorical observation, there exists an unstated (i.e., latent) perception, say 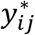, for each evaluation that could be considered continuous scores. We further suppose that an expert’s perceived score determines the what rating will be provided according to the following relationship:

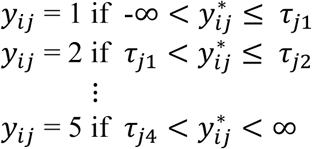

where τ_*j1*_ through τ_*j4*_ are the four rating thresholds. Our modeling approach can be represented as 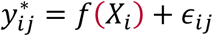, where 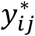 represents the expert’s perceived score, the function, *f*, represents the scoring algorithm, *X*_*i*_ denotes the vector of image feature values for colony *i*, *f*(*X*_*i*_) denotes the predicted pluripotency score for colony *i*, and ϵ_*ij*_ represents the difference between the predicted pluripotency score for colony *i* and the latent score corresponding to expert *j*’s perception of colony *i*.

Our modeling approach uses a bivariate normal distribution to represent deviations (i.e., ϵ_*ij*_) of each expert from the pluripotency scores predicted by a scoring algorithm. Because the continuous scores 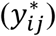 are not actually observed, the errors (ϵ_*ij*_) cannot be directly computed. Rather, the extent to which a given scoring algorithm is discordant with the scoring intervals corresponding to ratings provided by expert *j* is represented by the variance parameter for that expert (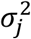, *j* = 1,2). The extent to which the experts agree in their disagreements with the scoring algorithm is represented by a covariance parameter 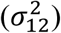. Collectively, the two variance parameters and the covariance parameter provide the elements of the variance-covariance matrix for the bivariate normal distribution.

Because we treat the ratings of the experts as censored observations, linear models were fit using the numerical optimization function, optim, in R [13] to maximize the likelihood of the observed data, rather than by conventional least squares. Random forests were fit using the R package random forest [14] while treating the ratings as exact values to facilitate computation. The predicted values from the fitted random forest were used as the scoring algorithm and incorporated into the evaluation of the thresholds and bivariate normal distribution best representing the discordance between the random forest predictions and the ratings provided by the experts. Additional modeling details are provided in Supplemental Information 1, sections V and VI.

During evaluation, we monitor the values of the variance-covariance parameters. Specifically, we compare the expert variances (representing their respective discordances with the predicted scores) to the variance of the difference between the experts (representing the discordance between the experts themselves). When a scoring algorithm produces an expert variance lower than the variance of differences between the experts, we say the scoring algorithm is more consistent with that expert than the two experts are with each other. Ideally, a scoring algorithm would be more consistent with each expert than the experts are with each other. In addition, the value of the covariance parameter is helpful in assessing the completeness of a scoring algorithm. An ideal scoring algorithm would produce a covariance estimate close to zero; large positive values indicate the model is overlooking evaluation trends exhibited by both experts, while negative covariance values are suggestive of over-fitting.

Additionally, we construct receiver operating characteristic (ROC) curves to assess how well a scoring algorithm identifies the highest rated colonies for each expert. This evaluation criterion is intended to reflect the envisioned usage of a pluripotency scoring algorithm to select the most promising colonies.

### Statistical Analysis

Statistical analyses were conducted in R and are displayed in Figures 3 and 4 and Supplemental Figures 4, 5, and 6. The predicted colony scores from each model in this study were computed using cross-validation, whereby a colony was never included in the training data for the model used to assign its predicted score (see Supplemental Information 1, section VI).

## RESULTS

### Image data set, expert rating, and LVM modelling approach

A total of 449 pluripotent stem cell colonies were rated by two experts. Colonies were given subjective ratings by experts on a scale from 5 (for the best pluripotent colonies), to 1 (those with the least pluripotent character). Characteristic criteria for rating were discussed and agreed on by the two experts, and these characteristics are described in Table 1. Examples of colonies are shown in Figure 1 and all colonies are shown in Supplemental Information 2. The colonies shown in Figure 1 were rated identically by the two experts, but often the experts provided different ratings for the same colonies.

**Figure 1.**
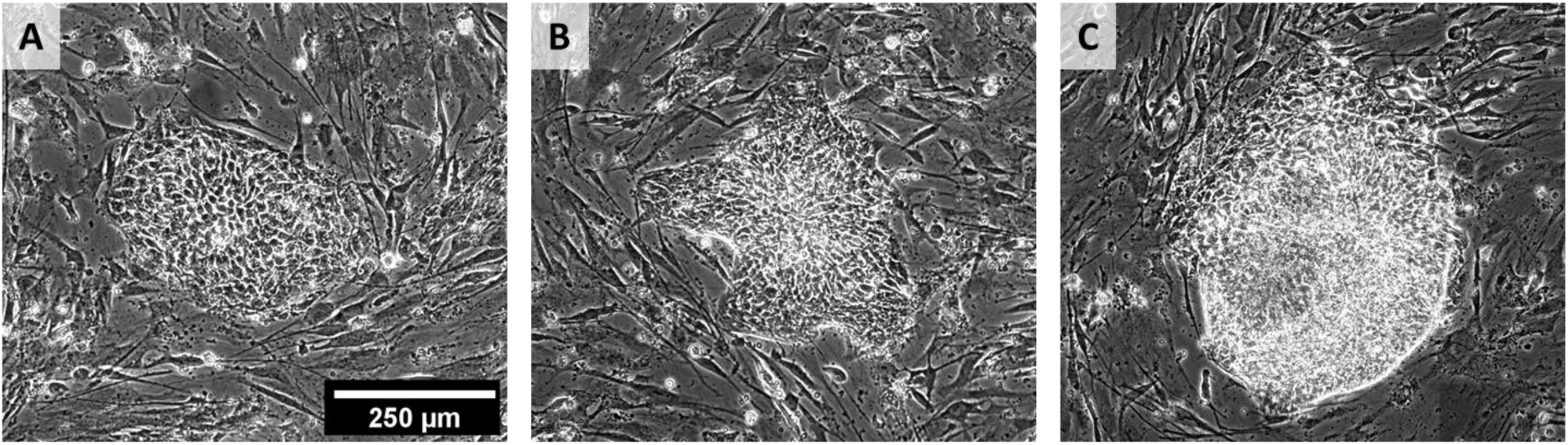
Example colony images illustrating the range of colony appearance over the 5 ratings. (A) A colony rated 5 by both experts (well 1, colony 10). (B) A colony rated 4 by both experts (well 2, colony 22). (C) A colony assigned rated 2 by both experts (well 1, colony 132). All colonies and their ratings are shown in Supplemental Information 2.

In the dataset of 449 images rated by two experts, only 48% of the colonies were given the same rating by both experts. Table 3 is a confusion matrix showing how the two experts varied from one another in their ratings.

**Table 3.** Confusion matrix of expert-assigned ratings illustrating correspondences and differences.

|  |  | **Expert 2 Ratings** |  |  |  |  |
| --- | --- | --- | --- | --- | --- | --- |
| **Expert 1**<br>**Ratings** |  | **1** | **2** | **3** | **4** | **5** |
|  | **1** | 7 | 5 | 0 | 0 | 0 |
|  | **2** | 2 | 31 | 19 | 11 | 4 |
|  | **3** | 1 | 21 | 52 | 49 | 11 |
|  | **4** | 0 | 4 | 35 | 105 | 43 |
|  | **5** | 0 | 0 | 2 | 25 | 22 |

Our goal was to develop models that allowed us to compare the basis on which experts rated stem cell colonies for pluripotency. This involved assessing whether experts used the same features in their evaluation, and the relative importance of the features they used. Our general approach to developing the linear models is described schematically in Figure 2A (additional details regarding implementation are provided in the Supplemental Information 1, sections V and VI). When constructing a linear model for a given set of features, three types of parameters were fit simultaneously to maximize the likelihood of the observed ratings: the weights assigned to the image features, which define the scoring algorithm; the score range associated with each unique rating value for each expert, defined by a vector of eight thresholds; and the variance covariance matrix of the bivariate normal distribution consisting of the variance between expert 1 and the model, the variance between expert 2 and the model and the covariance between the experts’ differences from the model.

**Figure 2.**
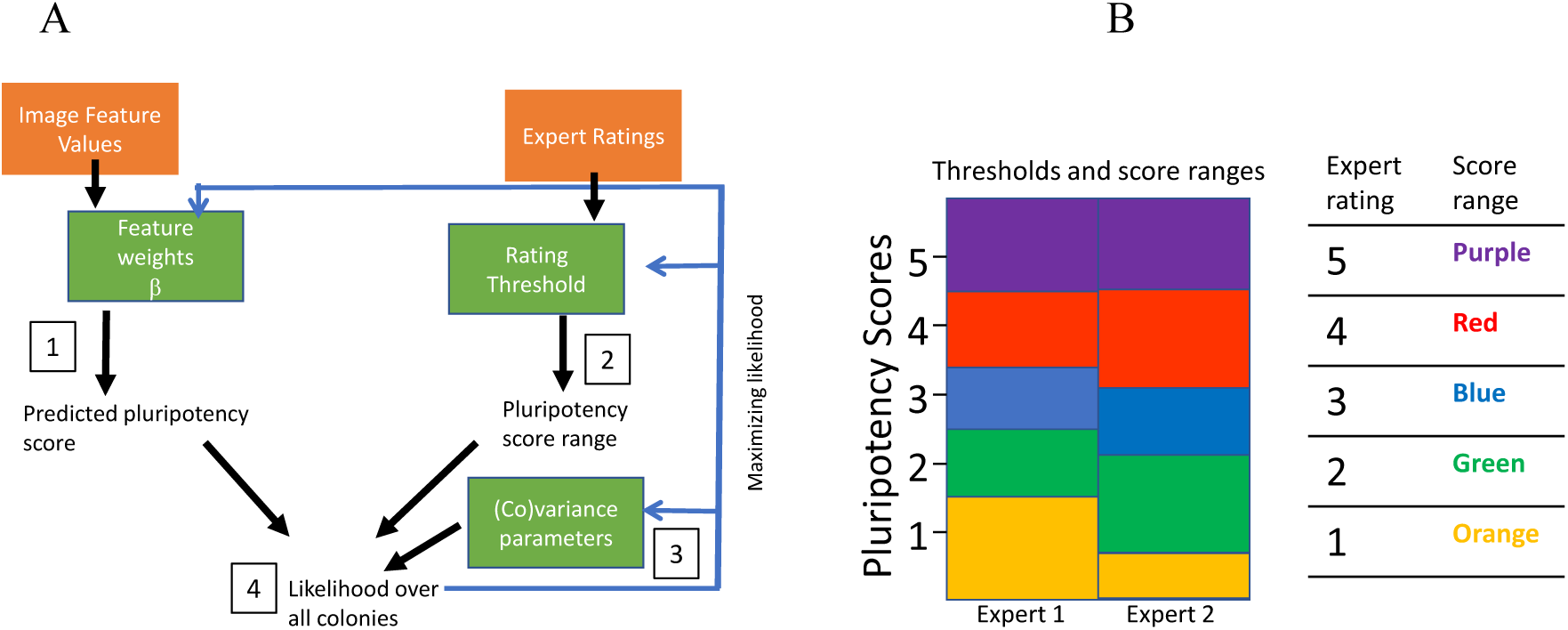
A. Schematic showing the relationship between the steps of the modeling process. The orange boxes represent input values and the green boxes represent variables that are to be optimized. Step 1: The predicted scores are computed as a linear combination of image feature values, using feature weights that are fixed across all colonies to a value chosen during model fitting. Step 2: Each expert has rated each colony on a scale of 1,2,3,4 or 5. Because of ambiguity in how an expert perceives quality, a set of four thresholds for each expert identifies the scores that divide an expert’s ratings as depicted in 2B. Each rating from each expert is translated to a range of possible predicted scores (according to the corresponding thresholds). Step 3: The discordance between the predicted scores for all colonies and the experts’ ranges of scores and the covariance between the experts’ discordance with the model are represented via a bivariate normal distribution. Step 4: Maximum likelihood is achieved when the concordance between predicted scores and corresponding pluripotency score ranges over all colonies is optimized. In a final step, the average expert-dependent score ranges are applied over all models. The differences in score ranges calculated for different models was not large and applying an average score range for each expert simplified data comparison (see Supplemental Figures 4 and 5). B. The relationship between expert ratings (1-5) and latent expert scores is not identical for the two experts. The rating thresholds optimized by the model define the expert score ranges. Expert ratings were constrained at a score of 1.5 and 2.5. For more details on the model, refer to Supplemental Information 1.

Figure 2B indicates how the ratings provided by each expert are interpreted by the model as scoring ranges, which are expert-specific. For each expert rating, the model assumes there is one precise but unknown value that represents the expert’s exact perception of pluripotency and falls somewhere within the corresponding expert score interval. We refer to this putative value as a latent score.

The linear models were constructed using a forward selection routine. The modeling process begins by giving all features a weight of 0, which causes all colonies to be given the same predicted score. At each iteration, a new model is created by assigning a non-zero weight to an additional feature that can best improve the performance of the model. The weight for each feature in a model is allowed to freely vary to whatever real number maximizes the likelihood, while features that are not included in the models retain weights of exactly 0. This process resulted in a collection of linear models. We report the models that include zero to 10 features, after which there was little improvement seen with any additional features.

After an initial round of fitting, we discovered that a common set of thresholds (defining the score ranges associated with each experts’ ratings) could be imposed across all 11 models without significantly reducing the model performance. Using a fixed set of thresholds across the models puts their scores and variances onto a common scale, which greatly aids comparison across models. Further details are provided in the Supplemental information 1, section VII.

### Evaluation of similarities and differences between expert ratings with LVM

The performance of the models developed within the 14 expert-inspired features and trained on both experts’ scores is shown in Figure 3. (The 14 features are described in Supplemental Information, section III and the algorithms are at https://github.com/usnistgov/stem_cell_segmentation. Receiver operating characteristic (ROC) curves were constructed using predicted scores obtained via 50-fold cross validation. The method and rationale for the cross-validation strategy is described in the Supplemental Information 1, section VI. The ROC curves in Figure 3A indicate that the model is quite effective at predicting expert 1 scores for colonies with a rating of 5 or a rating of either 4 or 5 (i.e., colonies that expert 1 rated as either 4 or 5 received high values from the scoring algorithm). In fact, the values produced by the scoring algorithm were more effective than the ratings from expert 2 at identifying expert 1’s highest rated colonies. On the other hand, colonies that expert 2 rated as either 4 or 5 had a greater spread in values from the scoring algorithm, and the scoring algorithm identifies colonies ranked highly by expert 2 only about as well as the ratings from expert 1 do. Adding more than two features had a small effect on ROC performance. Using a random forest, evaluated using the complete set of 14 image features, did not improve ROC performance over the linear models.

**Figure 3.**
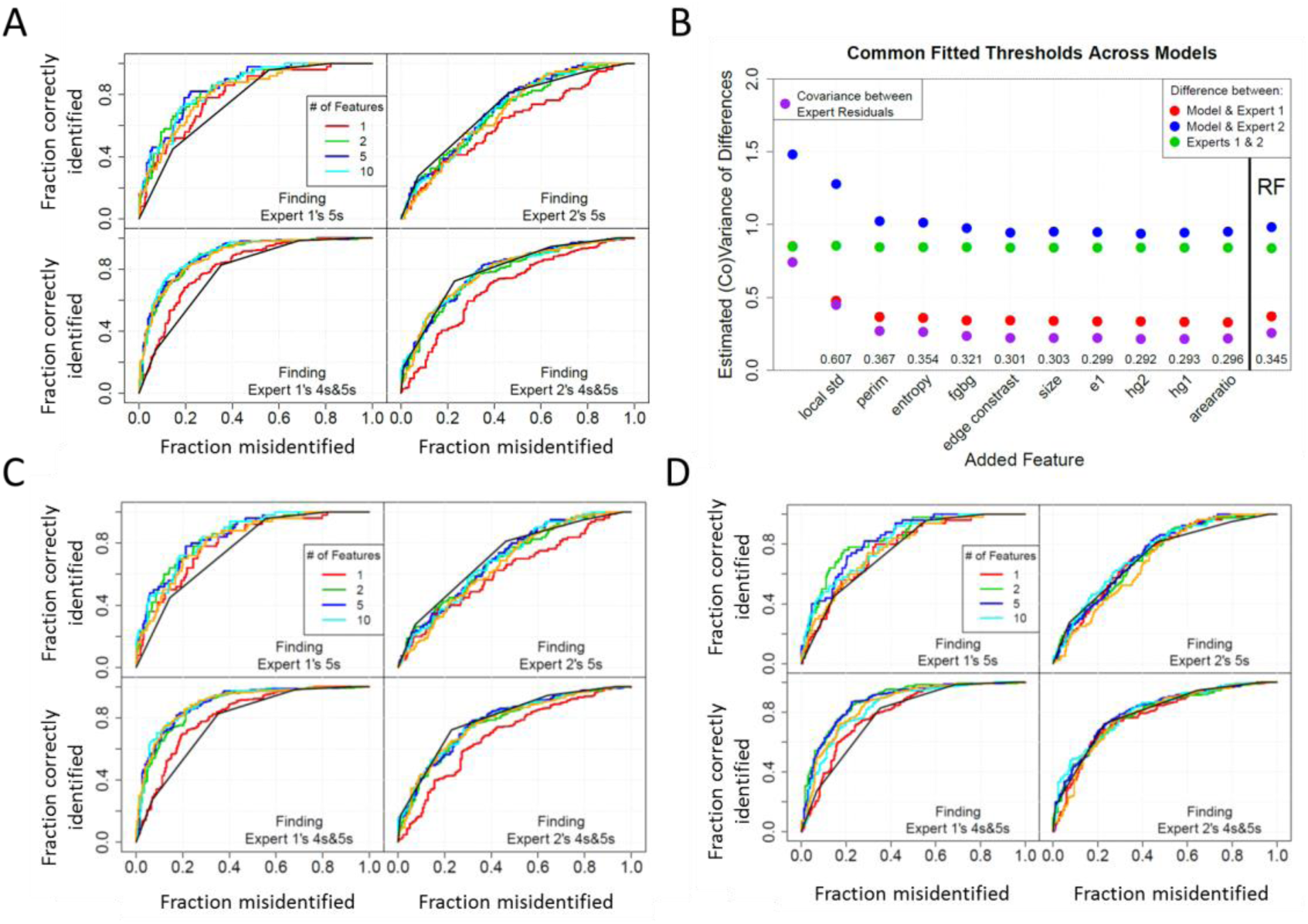
A and B. Fitting results from linear models with different numbers of features trained on scores from both experts and accessing 14 expert-inspired features. A. ROC curves for the model predictions of colonies given a score of 5 or 4-5 and compared to scores from expert 1 and from expert 2. The orange curve is the result of RF fit to 14 features. The black ROC curve is computed from the confusion matrix of expert 1 and expert 2 ratings. B. Estimated variances and covariances in the differences between expert scores and model predictions. Shown are results of 10 models, where each new model includes all features from the previous model and is created by the addition of the indicated feature. This plot represents the variance in the differences between the predicted scores from the models and the scores from expert 1 (red) and expert 2 (blue). The purple markers indicate covariance in the differences calculated between the experts scores and the model predictions. The numerical values of these covariances appear above the designation of the corresponding model on the x-axis. The green markers indicate the variance between the experts that is calculated according to equation 1.C and D. Training models on each expert individually. C. Models trained on expert 1 ratings. D. Models trained on expert 2 ratings.

The order in which features were added to the model is shown in Figure 3B, together with the variance in the differences between the predicted colony scores and the latent scores of each expert, evaluated with the addition of each feature. The red and blue markers represent how far, on average, the model’s predicted scores are from the putative (latent) scores of the respective expert over all colonies. Because each expert’s score is represented as a range of scores (and not a single value), the variance parameter is not equated to any explicit sample variance but is a characteristic of the bivariate normal distribution used to represent the disagreement between the scoring algorithm and the expert ratings.

A measure of the direct discordance between the experts’ scoring ranges over all colonies (shown with green markers) is attainable from the elements of the variance-covariance matrix of the bivariate normal distribution:

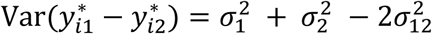

This quantity provides a benchmark against which to evaluate scoring algorithm performance. When 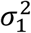 (red markers) and/or 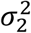 (blue markers) are less than 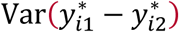 (green markers), the variance-covariance estimates indicate that the disparity between the scoring algorithm and the latent scores of the expert(s) would be less than the disparity between the latent scores of the two experts. The scoring algorithms do a much better job at predicting expert 1 than expert 2; the discordance between expert 2 and the predicted scores is similar to the discordance between experts as was also apparent from the ROC curve evaluations. Also, as seen in the ROC curves, a model with two features captures most of the predictive power. The addition of more features to the model has a minimal effect on reducing the variance of the differences.

The covariance in the differences between each expert and the model predictions is shown with the purple markers. The covariance serves as a measure of how much information is missing in the model; for an ideal model that captured all the patterns with features of interest that are common to both experts, the expected covariance in residuals between experts would be 0. The magnitude of the covariance indicates that ~70% of the systematic response to image characteristics, shared by these experts, is captured by the model. When the scoring algorithm under-values or over-values a given colony with respect to the rating of one expert, it tends to do the same with respect to the other expert as well. This tendency indicates that the model fails to capture all of the experts’ information. The amount of variation associated with expert 1 that is not also reflected by expert 2’s ratings is small (as indicated by the small difference between the red and purple markers). The average magnitude of the residuals between the scoring algorithm and expert 2, however is much larger, indicating the presence of either greater random variations in the responses from expert 2 or the presence of rating tendencies not shared by expert 1 (e.g., consideration of additional subjective image features).

The plot of covariance in the differences between each expert and the model predictions indicates that both experts exhibit a common rating tendency, or shared perception, that is not captured in the model. The plot indicates that adding two features to the model captures all but 36% of their shared perception. By sequentially adding features to the model, the covariance in residuals is slightly reduced; for the 10-feature model, the shared perception still missing from the model is reduced to about 27%. There are several possibilities for how the experts’ shared perception still missing in the model could be captured. It is possible that the segmentation routine we used does not reflect what the experts identify, and, hence, even the right image feature algorithms may produce non-ideal values because they are presented with non-ideal pixel sets. Another possibility is that there are additional feature algorithms that would improve our ability to mimic expert assessments. A third possibility is that we have the right image features and values but have not found the optimal function that translates the features into a pluripotency score.

To test if experts used more features than provided so far, the model was trained on both experts using 56 features consisting of the 14 used to generate the data in Figure 3 plus 42 Haralick and wavelet features (as listed in Supplemental Information 1, section III). The addition of these features has no significant effect on performance as shown in Supplemental Figure 6.

To assess whether a more flexible class of scoring algorithms would provide different outcome, we also trained a Random Forest model with the 14 or 56 feature sets. The results are shown in Figure 3A as the orange curve, in 3B under the label RF, and in Supplemental Figure 6. The more flexible Random Forest model performed approximately the same as the linear approach.

Although the models were trained on both experts, the differences in the effectiveness of the models to predict the ratings of the different experts are large. While the experts agreed on the rating criteria, the lack of concordance that is apparent in the confusion matrix suggests that the experts could be basing their ratings on different features or could be applying very different levels of importance to different features. We repeated the forward-selection development of linear models, and the Random Forest fits, training to each expert separately, again providing 14 features. Figure 3C and D show that training the model on the individual experts has a nominal effect compared to training the model on both experts, for predicting either expert 1 or 2. Thus both experts contribute to, and neither expert dominates, the selection of model parameters for predicting colony scores.

### Evaluation of image features useful for predicting expert ratings

Surprisingly, Figure 3 shows that for the model trained on two experts, just two image features, “local std”, a custom feature that is associated with high phase contrast in local regions, and “perimeter” account for nearly all the predictive power. Adding more features to the model only improves the model performance slightly. We plotted the weights for features used in the models trained on both experts, and the models trained in either expert 1 or expert 2. The results are shown in Figure 4, and confirm that for both experts, the predominant features are “local std” and “perimeter”.

**Figure 4.**
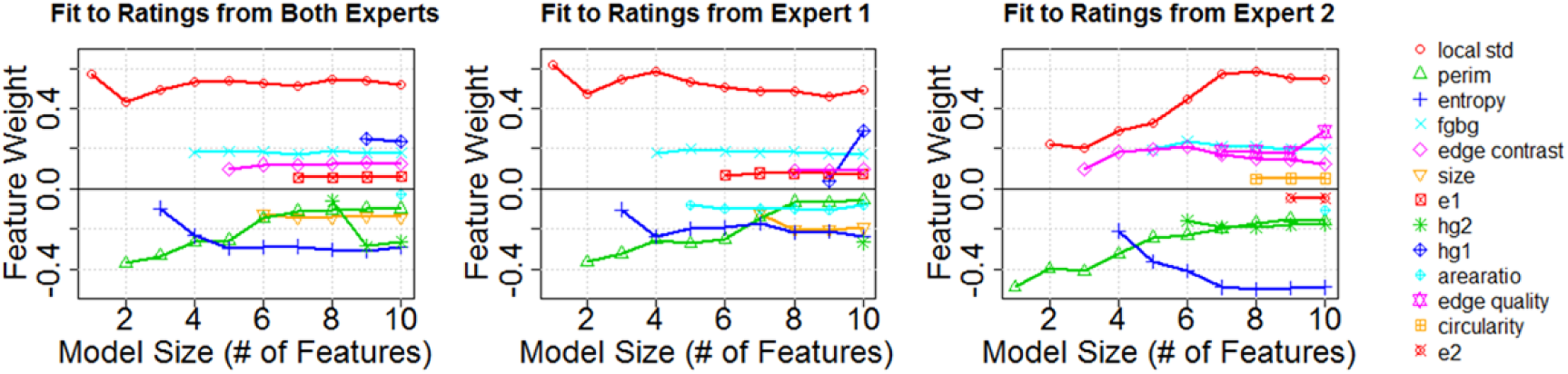
Features and weights for models using 14 features and trained on both experts (A), expert 1 (B) and expert 2 (C). The 10 most important features in the models are shown. The additional 4 features provided minimal improvement to the models. Under all cases, the most prominent features that appear in the models are “local std” and “perimeter”.

## DISCUSSION

For this study, we used an approach based on identified image features and algorithms developed to capture the characteristics indicated by experts as important for evaluating pluripotent stem cell colony qualities. Many of the features that our experts identified are similar to features identified for similar studies [8, 15, 16]. The experts of Wakui et al [15] identified intercellular spacing, round, well defined margin, and smooth margin as important criteria. Maddah et al [8] identified biologically relevant features that included colony margin character, as did Marx et al [16], who also used a feature associated with heterogeneity within the colony (that is likely similar to our features of local std). We also employed other features, including some Haralick features, which are not based explicitly on human visual cues, but have been used successfully by other groups to detect intensity patterns in sub-cellular features present in optical microscopy images [17, 18]. In other work, wavelet features used without any additional features were found to correlate well with the apparent pluripotency of cultured stem cells [19]. In this work, the 14 expert-inspired features were sufficient to capture most of the experts’ perceptions, and the addition of Haralick and wavelet features did not improve the performance of the models for comparing expert scores. By developing algorithms that are intended to quantify what the experts look for, we are able to assess the criteria that they each apparently use for visual evaluation.Inconsistencies between expert ratings and molecular markers of pluripotency have been observed by others [1, 15, 20]. In this work, we acknowledge that neither expert represented ground truth and allowed each expert’s ratings to represent a range of values. This general approach has been discussed by Frenay [21]. We compared models trained on both experts with models trained on one expert at a time. All models produced colony scores at least as concordant with each expert’s scores as the experts were with one another. Models were always better at predicting the scores of expert 1 than expert 2 was at predicting expert 1.

Interestingly, two image features dominated all models, whether they were trained on ratings from one or both experts. These image features relate to intra-colony contrast (“local std”) and a quality of the colony margin (“perimeter”), suggesting that these are robust features for this application, and that both experts were similarly influenced by these features when rating colonies. The importance of the “perimeter” image feature probably reflects that experts are aware that a colony that is large is more likely to spontaneously differentiate than a smaller colony, even though this characteristic was not initially acknowledged as one of their rating criteria.

The consideration of multiple experts can thus provide evidence of robustness of stem cell classification that cannot be achieved if only one expert were used. The relatively poor performance of the model when predicting expert 2’s scores might indicate that expert 2 is responding to features that are not used by expert 1. But cross correlation analysis indicated that only a small amount of discordance that could be attributed to features was not shared between the experts. Since we also trained the model on expert 2 alone, we conclude that these would be features that are outside of the feature space we’ve considered and are not shared by expert 1. Alternatively, this observation could also simply be the result of relative inconsistency by this expert in scoring.

The correlations in the differences between expert scores and the models indicate that both experts share perceptions that were not accounted for in the models. It appears that the models we employ based on our feature set account for about 70% of the experts’ decisions, suggesting that the evaluation of stem cell pluripotency by these experts may be improved by acknowledgement of additional, as yet undiscovered, features..

The latent variable modeling approach can be extended to more than two experts. The only requirement is that one must estimate or specify the parameters for a larger variance-covariance matrix. The simplest case would be to assume the expert-specific tendencies are all independent of one another. Then there would be a variance parameter for each expert and one additional covariance parameter. Otherwise, each pair of experts could have a different covariance.

## CONCLUSIONS

This study demonstrates the use of non-concordant ratings from multiple experts to add confidence to decisions about colony ratings. Each expert’s ratings contribute to the model independently. This approach allows us to extract information about commonalities and differences in the ratings of different experts. The differences between these experts can be attributed to slight differences in perception of relative importance of features, and consistency in scoring. Although there were differences between these experts, models based on each of their ratings were highly concordant with respect to features used and the presence of shared, unidentified features. This concordance enhances confidence that the experts are evaluating similar content in the images.

## Availability of Data and Materials

The raw image data used in this study are available at https://isg.nist.gov/deepzoomweb/data.

Images and expert ratings are shown in Supplemental Information 2. The segmentation and analysis algorithms are available at https://github.com/usnistgov/stem_cell_segmentation.

## Author Contributions

MH, DJH, ALP, RM conceived and designed the study; DJH JGC, SKK and MH collected the data; SL, AP, YL, PB, OA performed the analysis; and MH, SL, AP, ALP wrote the paper.

