## Supplemental Information for "A Latent Variable Model for Evaluation of Disparate Ratings of Stem Cell Colonies by Two Experts"

### Supplemental Materials Sections

**I.** Protocol for collecting phase contrast images of stem cell colonies (pg. 2-3)
  a. Supplemental Figure 1. Phase contrast reference material.
**II.** ImageJ macro for identifying colonies to present to experts for evaluation (pg. 4-6)
**III.** Description of image analysis features (pg. 7-9)
  a. Supplemental Figure 2. Schematic of colony regions for image analysis.
**IV.** Similarities in Image Analysis Features: Correlation analysis of features (pg. 10)
  a. Supplemental Figure 3. Heat map of feature correlations.
**V.** Detailed Model Description (pg. 11-12)
**VI.** Additional Details about using Cross Validation (pg. 13)
**VII.** Additional Details about Fitting Thresholds (pg. 14-16)
  a. Supplemental Figure 4. Analysis of fitting expert specific thresholds.
  b. Supplemental Figure 5. Significance testing of expert specific thresholds.
**VIII.** Model evaluation with will full feature 56 feature set (pg. 17-19)
  a. Supplemental Figure 6. ROC and variance-covariance analysis.

**Disclaimer**: Commercial products are identified in this document in order to specify the experimental procedure adequately. Such identification is not intended to imply recommendation or endorsement by the National Institute of Standards and Technology, nor is it intended to imply that the products identified are necessarily the best available for the purpose.

### I. Protocol for collecting phase contrast images of stem cell colonies

1. Collect a 'dark count' image (an image in the absence of light)
2. Phase condenser must be setup correctly (Kohler illumination) and microscope warmed up (lamp on for 15 minutes).
3. Imaging the STAGE MICROMETER (to record the spatial dimensions of the imaging field): place the reticule on the microscope surface and focus on the ruler gratings.
4. Remove the STAGE MICROMETER and capture a BLANK image to record artifacts in the image due to dust and debris in the optical train.
5. IMAGING the PHASE CONTRAST QC BENCHMARK (to ensure appropriate alignment of phase contrast optics): place patterned standard onto surface, focus on edge of the 500 µm squares. Collect image. Also reference the lamp intensity to the features on the standard. There should no changing of any optics after this step, though the lamp intensity can be changed.
6. IMAGING STEM CELL COLONIES: Wet kim wipe with a small amount of 100% ethanol and wipe off bottom of well plate (may be an issue with ink marking on bottom of plate). Evaluate well bottom to make sure dust and finger prints are removed.
7. Place stem cell sample on stage, focus on feeder cell layer adjacent to colony. Center colony into field of view. Collect image.

<u>Protocol notes</u>

Use the same exposure time of throughout the protocol (approximately 200 ms). Collect images on a scientific monochromatic camera at 1x1 binning. Collect a dark count image to ensure the dark image is above 100. Make sure camera is set up properly. May set up a specification on the background of image for a particular exposure time and use lamp intensity for any additional adjustments. Assume CCD bias offset and exposure time are identical between runs.

**For the initial dataset, we will focus on colonies that are sufficiently far from the edge of the well that the phase contrast image is NOT distorted by edge effects. This is to assemble a set of images that are clean and provide clear information when used as a reference dataset online. Non-distorted images are also expected to facilitate the automated identification of good/bad colonies compared to distorted colonies. So, collect images only from colonies that are sufficiently far from the edge of the well that the phase contrast image is NOT distorted by edge effects.

1. image of dark current
2. image of MICROMETER
3. image of BLANK

4. image of (TEST) PHASE CONTRAST QC BENCHMARK
5. images of colony (good and bad)

*Analysis notes*

1. (TEST) PHASE CONTRAST QC BENCHMARK should have a uniform, low intensity inside the large squares (background) and the profile between two squares should transitions from low intensity to high intensity with a positive slope. No drops below background intensity should be observed. This suggests that the microscope phase rings are well aligned and are adequate for cell imaging.
2. Correction by dark count subtraction and division by no sample image (collected with identical phase optics as cells) corrects for phase artifacts due to the fixed optical system.

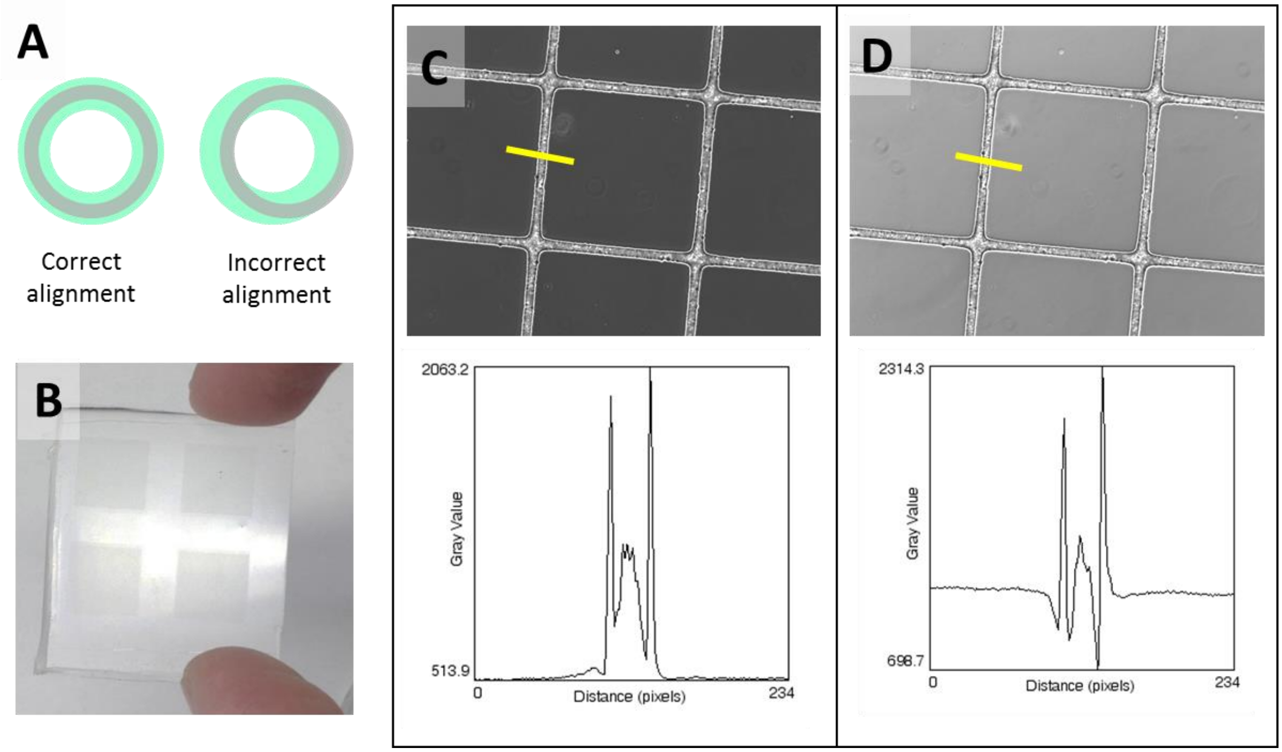


**Supplemental Figure 1. (**A) Schematic diagram of the illumination annulus (grey) and the phase plate (green) used in Zernike phase contrast. Rings on the left are correctly aligned and the rings on the right are incorrectly aligned. (B) Photograph of the PDMS benchmarking material that has been formed by casting PDMS prepolymer in a microfabricated mold. (C and D) Zernike phase contrast images of the benchmark material acquired with a 10x/0.3 NA Ph1 objective. The square features in the images are squares with sides approximately 500 µm in length with valleys between the squares that are approximately 50 µm across and 25 µm deep. The illumination annulus and the phase plate are correctly aligned in (C) and mis-aligned in (D). The line scan beneath the images, corresponding to the yellow line in each image, indicates the intensity response difference that can be observed between images acquired when the alignment is correct (C) or incorrect (D).

### II.. ImageJ macro for identifying colonies to present to experts for evaluation

```
//Designed to segment Human ES cell colonies captured at 10x phase
//Test input file captured on Zeiss 135 TV with 10x 0.3NA Phase 1 lens and condenser ring
//Camera is CoolSnapHQ2
//Test image 20100602_BG01_38_well2-0044.zvi and others from same series
//Performance is ok, but could be optimized.  MEFS are not confused with colonies, but the
//edges are sometimes ambiguous - probably reflects their true ambiguity.
//"big_circle.roi" is the user selected roi outlining the meniscus effected zone at the edges of the
dish.
//Requires ImageJ 1.45d or later

save_path=getDirectory("Choose a working directory to save the analysis results");

run("Colors...", "foreground=white background=black selection=yellow");

waitForUser("Select the image with colonies and select the roi for analysis...");
colonyID=getImageID();
run("Clear Outside");
saveAs("Selection", save_path+"big_circle.roi");
run("Select None");

run("Duplicate...", "title=full_colony_image");
saveAs("tif", save_path+"full_colony_image.tif");
selectImage(colonyID);
close();
selectWindow("full_colony_image.tif");
run("Find Edges");
run("Gaussian Blur...", "sigma=10");
run("Subtract Background...", "rolling=30 create");
run("Gaussian Blur...", "sigma=10");
setAutoThreshold("Default dark");
run("Analyze Particles...", "size=20824.66-Infinity circularity=0.00-1.00 show=Nothing display
clear summarize add");
close();

//this loop finds the max width and height of the colonies to save them in an image sequence
open(save_path+"full_colony_image.tif");
clnies= roiManager("count");
avg_size=0;
max_width=0;
max_height=0;
for (i=0; i<clnies; i++)		{
	roiManager("Select", i);
	getSelectionBounds(x, y, width, height);
	avg_size=avg_size+width+height;
```

```
	if (width>max_width) {
	max_width=width;
	}
	if (height>max_height) {
	max_height=height;
	}
}
avg_size=avg_size/(2*clnies);
pad_size=round(0.4*avg_size);
print("average colony x,y dimension is: "+avg_size+" pixels");
print("pad size is: "+pad_size+" pixels");

//loop writes an image sequence containing each of the colonies
for (i=0; i<clnies; i++)		{
	selectWindow("full_colony_image.tif");
	roiManager("Select", i);
		run("To Bounding Box");
		run("Enlarge...", "enlarge="+pad_size);
	run("Duplicate...", "title=temp_colony_segmented");
	roiManager("Select", i);
	run("Draw");
	run("Select All");
	run("Copy");
	newImage("colony", "16-bit Black", max_width, max_height, 1);
	run("Paste");
	saveAs("tif", save_path+"colony_"+IJ.pad(i+1, 3));
	close();
	selectWindow("temp_colony_segmented");
	close();
}
	run("Select None");
	File.delete(save_path+"full_colony_image.tif");
```

### III. Description of image analysis features

We used a number of different features to quantify how well-defined the margins of colonies appeared to be, a second set of features to calculate the extent of intra-colony phase brightness which provided an indication of size and shape of cells, and a number of other standard image features including measure of colonies size, shape, and texture

The features listed below are based on 3 different regions of the colonies, depicted in Figure 2: Region 1, the center region of the colony, Region 2, the region just inside the colony margin, and Region 3, the region just outside the colony margin. Features 1-5 and 9-14 (below) were developed based on experts' descriptions of criteria they used for evaluating pluripotency.

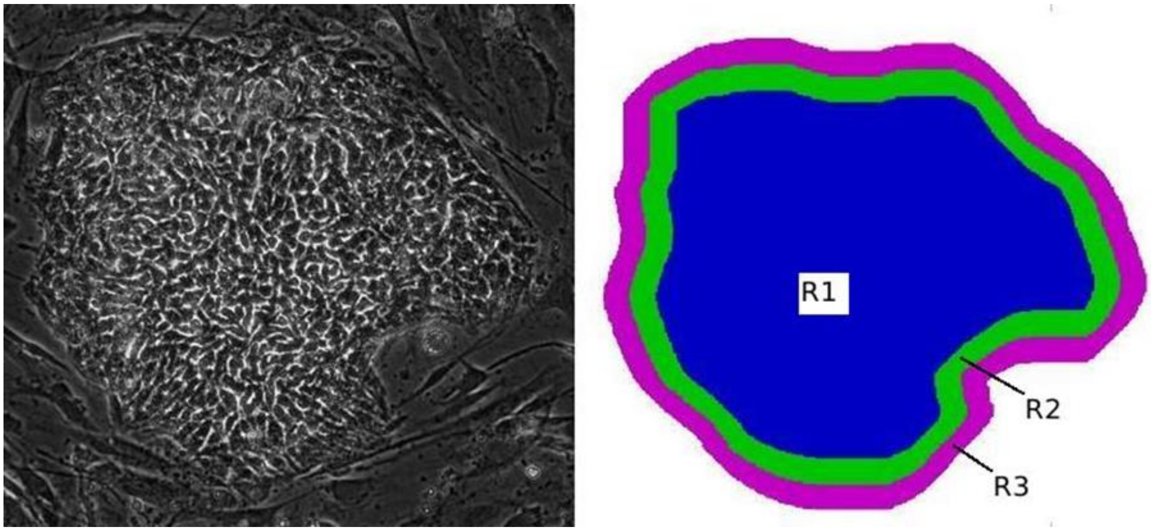


**Supplemental Figure 2.** Schematic illustrating colony regions of interest, including the exterior margins to the colony (R3), the interior margins of the colony (R2), and the center region of the colony (R1).

<u>Features describing colony margins:</u>

1 *Edge texture difference* (fgbg): A ratio of the entropy (see description of Entropy algorithm #10 below) of pixels in R2 divided by the entropy of pixels in R3 gave us a good measure of how distinct the textures inside and outside the margin were, where each region was 50 pixels thick. The choice of 50 pixels allows for some ambiguity in the segmentation while obtaining sufficient characteristics between the foreground and background.

2 *Edge intensity difference* (edge quality): For each pixel on the R2-R3 margin, we averaged the intensities of the 5 pixels just inside and the 5 pixels just outside each pixel at the colony margin. The ratio of these 2 averages is collected around the margin and the average of all colony margin ratios found. This gives a measure for how sharp the intensity differences are at the colony margin.

3 *Edge smoothness* (edge contrast): Otsu segmentation was performed on each colony image and the resultant threshold applied. Holes in the resulting mask are filled, leaving most of the unfilled pixels near the colony margin. A ratio of the Otsu segmented and filled area to the area of the colony mask is found; the closer to 1.0, the smoother the edge.

4-5 *Local standard deviation difference at edge* (e1) and *Local standard deviation extrema at edge* (e2): Both of these features quantify the contrast at the colony margin and are derived from local intensity standard deviation values in a 20 by 20 neighborhood of each pixel. E1 represents the fraction of pixels in R2 and R3 that are more than half a standard deviation (sd) above the mean value for the image. Levels are assigned based on the number of increments above the 0.5*sd mark the fraction is. Values range from 1 (0.5+0.05) sd to 20 (0.5+20*0.05) sd. E2 is assigned the lowest level for an individual pixel in the R2-R3 region.

6 *Area* (area): The number of pixels in each segmentation mask was counted.

7 *Perimeter* (perim): The number of pixels at the R2-R3 margin of each colony was counted.

8 *Circularity* (circularity): The quantity: 4.0*pi*Area/Perimeter^2, measures how evenly the colony grows in the outward direction.

Features to calculate the extent of intra-colony phase brightness:

9 *Local standard deviation magnitude within colony* (local std): To quantify the large local pixel intensity differences in the colonies, we found a local intensity standard deviation for each 20 by 20 neighborhood surrounding each pixel. We found the mean (lsm) and standard deviation (lssd) of these values over the image. This feature measures the fraction of pixels with very high local standard deviation values, at least 3*lssd above the image mean, lsm.

10 *Entropy of colony pixels* (entropy): From the histogram of pixel intensities within the segmentation mask, R1 and R2, the following sum is computed over all *i* bins in the histogram: *100.0- (maxlog – p(i)* log(p(i))/rangelog*, where *p(i)* is the probability of the $i^{th}$ bin, *maxlog* = -log(1.0/65536.0) and *rangelog* = *maxlog*/100.0.

11-12 *Local standard deviation dispersion 1* (hg1) and *Local standard deviation dispersion 2* (hg2): These 2 features measure how similar local standard deviation values are across the colony. Local standard deviation values are divided into 5 groups according to the number of 0.5 sd increments above the image mean they lie. These groups are ordered by how many pixels are in them. Hg1 is then computed as: (highest + 0.5*second highest – 0.25)/0.75; highest and second highest are in the range 1-5. Hg2 is the standard deviation of the number of pixels in each group.

13 *Otsu colony area ratio* (arearatio): Otsu segmentation was performed on each colony image and the resultant threshold applied. A ratio of the area of the resulting mask to the total colony area gives an estimate of the fraction of contrasting dark and light regions.

14 *Otsu colony threshold object count* (holes/area): Otsu segmentation was performed on each colony image and the resultant threshold applied. This feature measures the ratio of the number of holes in the resulting mask to the total area of the mask, and quantifies the size of individual cells within the colony.

These segmentation and analysis algorithms are at
*https://github.com/usnistgov/stem_cell_segmentation*

Other off-the shelf, general image analysis features applied to 56 feature models:

Wavelets: A set of 30 wavelet-based features, 10 levels using Debauchies 4, was used to quantify textural features of each colony. The set of 30 wavelet texture features was calculated using software from *http://murphylab.web.cmu.edu/services/SLF/*.

Haralick feature set: This is the set of 13 Haralick texture features, calculated using MATLAB software (Mathworks, Natick, MA). One of the computed Haralick features produced the same value for all colonies. Because it provided no discriminatory power, the feature was omitted so that 12 Haralick texture features remained.

### IV. Similarities in Image Analysis Features: Correlation analysis of features

Highly correlated image analysis features provide similar information to the model for predicting expert scores. A heatmap of correlation coefficients between all 56 features used in this study is shown in Supplementary Figure 3 and can be used to identify highly correlated features. The off-the-shelf features (Haralick and wavelet) were highly correlated amongst themselves.

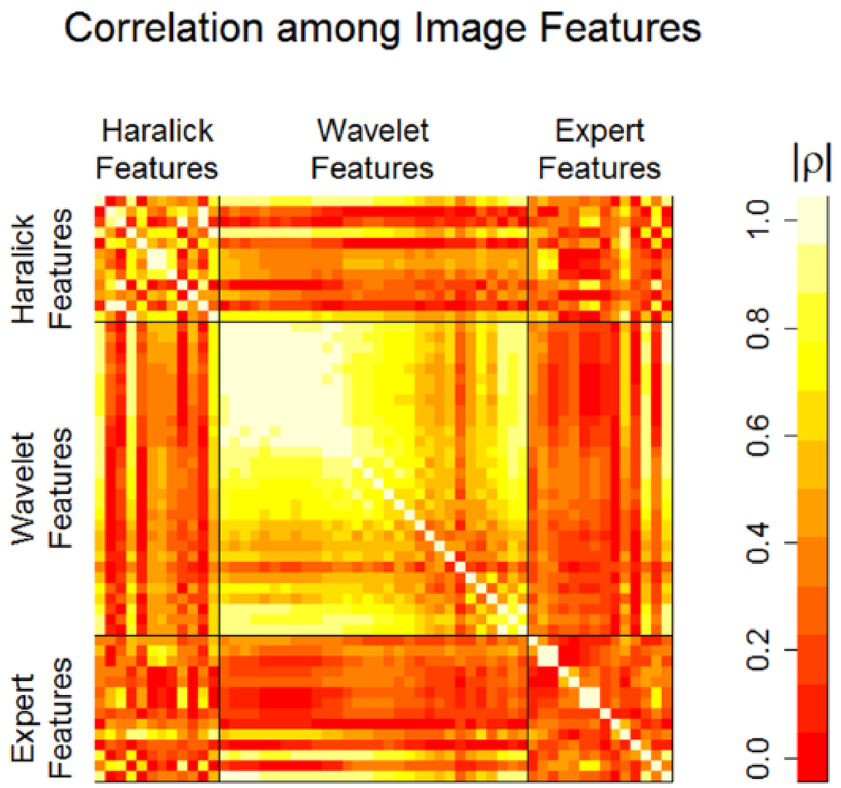


**Supplemental Figure 3.** Similarity between image features. The features are categorized as Haralick features, wavelet features, and expert features. The magnitude of the correlation coefficient (Pearson's) is indicated by both the color and scale.

### V. Detailed Model Description

Let $y_{ij}$ denote the rating for colony $i$ ($i = 1, \dots, 480$) reported by expert $j$ ($j = 1,2$). We model $Y_{ij}$ as being a censored (i.e., rounded) function of a latent perceived score $y_{ij}^*$ for colony $i$ from expert $j$. Specifically, for each expert, we allow four thresholds, $\tau_{jk}$ ($j = 1,2; k = 1, \dots, 4$) that identify the ranges of perceived scores that would lead to the expert providing a rating of 1, 2, 3, 4, or 5, respectively. The relationship between $y_{ij}$ and $y_{ij}^*$, given the thresholds, is assumed to be $y_{ij} = 1 + \sum_{k=1}^{4} I_{y_{ij}^* > \tau_{jk}}$, where $I_{condition}$ is a binary function equal to 1 if $condition$ is true and equal to 0 otherwise. Equivalently, if one assigns $\tau_{j0} = -\infty$ and $\tau_{j5} = \infty$, a rating $y_{ij}$ is interpreted as conveying that $y_{ij}^*$ falls somewhere within the interval given by $(\tau_{j(y_{ij}-1)}, \tau_{jy_{ij}})$. (This concept will be revisited when describing integration limits used to evaluate likelihoods.)

Let $f_j(I_i)$ represent the average latent score expert $j$ would perceive over many repeated evaluations of the image for colony $i$, $I_i$. At any one particular evaluation, the expert experiences a random perturbation around this average value, $y_{ij}^* = f_j(I_i) + \epsilon_{ij}$, and reports the corresponding rating $y_{ij}$ in accordance with the description previously provided. The evaluation tendencies for each expert can be partitioned in a component shared by both experts and a component specific to that expert. That is, $f_j(I) = g(I) + g_j(I)$, where function $g$ represents the component of evaluation tendencies that are common to both experts and the function $g_j$represents the component of evaluation tendencies present for expert $j$, but not the other expert. Among the infinitely many such decompositions, we seek those which maximize the proportion of variability attributed to common tendencies $g$and for which $\mathrm{cov}(g_1(X_i), g_2(X_i)) = 0$.

Our aim is to identify a scoring algorithm (i.e., a function of the image features) that best reflects the evaluation tendencies exhibited by both experts. That is, we are seeking a scoring algorithm $s(X_i)$ whose values across the set of colonies are as correlated with those of $g(I_i)$ as possible, where $X_i$ is a vector containing the values of the image features computed from the image for colony $i$ ($I_i$).

Our modeling approach can be represented as $y_{ij}^* = s(X_i) + e_{ij}$, where $s$ represents the scoring algorithm, $X_i$ denotes the vector of image feature values for colony $i$, $s(X_i)$ denotes the predicted pluripotency score for colony $i$, and $e_{ij}$ represents the difference between the predicted pluripotency score for colony $i$ and the latent score corresponding to expert $j$'s perception of colony $i$. That is,

$$e_{ij} = y_{ij}^* - s(X_i) = \left( g(I_i) - s(X_i) \right) + g_j(I_i) + \epsilon_{ij},$$

where $g(I_i) - s(X_i)$ represents the extent to which the scoring algorithm fails to capture the entire shared evaluation tendencies of the experts, $g_j(I_i)$ represents the contribution of evaluation tendencies specific to expert $j$, and $\epsilon_{ij}$ represents the random fluctuation around the average perceived score for colony $i$ corresponding to this particular instance of evaluation. Note that because each expert has rated each colony at most once, our data cannot distinguish random variability from rating tendencies exhibited by one expert and not the other. The failure of a scoring algorithm to capture the entire shared evaluation tendencies of the experts contributes equally to $e_{i1}$ and $e_{i2}$, thereby inducing a correlation that enables this component of error to be distinguished from the expert-specific contributions.

Our modeling approach uses a bivariate normal distribution to represent deviations of each expert from the pluripotency scores predicted by a scoring algorithm (i.e., $e_{ij}$). Because the continuous scores ($y^*_{ij}$) are not actually observed, the errors ($e_{ij}$) cannot be directly computed. Rather, the extent to which a given scoring algorithm is discordant with the scoring intervals corresponding to ratings provided by expert $j$ is represented by the variance parameter for that expert ($\sigma_j^2,\ j = 1,2$). The parameter $\sigma_j^2$ characterizes the average value of $e_{ij}^2$ across the colonies. When the three components of error are independent, this quantity can be interpreted as the sum of the respective variances for $g(I_i) - s(X_i)$, $g_j(I_i)$, and $\epsilon_{ij}$. The extent to which the experts agree in their disagreements with the scoring algorithm is represented by a covariance parameter ($\sigma_{12}^2$), which characterizes the average value of $\left(g(I_i) - s(X_i)\right)^2$ across the colonies. Collectively, the two variance parameters and the covariance parameter provide the elements of the variance-covariance matrix for the bivariate normal distribution.

For a given set of model parameters, including the scoring algorithm $s$ thresholds $\tau_{jk}$ ($j = 1,2; k = 1, \ldots, 4$, variances $\sigma_j^2$ ($j = 1,2$), and covariance $\sigma_{12}^2$, the likelihood for the reported scores is computed as

$$L(s, \boldsymbol{\tau}, \boldsymbol{\Sigma}) = \prod_{i=1}^{480} \int_{\tau_{2(y_{i2}-1)}}^{\tau_{2y_{i2}}} \int_{\tau_{1(y_{i1}-1)}}^{\tau_{1y_{i1}}} \phi_{\boldsymbol{\mu}_i,\boldsymbol{\Sigma}}(s_1, s_2) \partial S_1 \partial S_2,$$

where $\phi_{\boldsymbol{\mu}_i,\boldsymbol{\Sigma}}$ is the density function of a bivariate normal distribution with mean vector $\boldsymbol{\mu}_i = \begin{pmatrix} s(X_i) \\ s(X_i) \end{pmatrix}$ and variance-covariance matrix $\boldsymbol{\Sigma} = \begin{bmatrix} \sigma_1^2 & \sigma_{12}^2 \\ \sigma_{12}^2 & \sigma_2^2 \end{bmatrix}$. For clarity, the limits of integration for the outer most integral are the upper and lower bounds of the interval corresponding to the rating reported by expert 2 for colony $i$, determined from the corresponding thresholds as ($\tau_{2(y_{i2}-1)}$, $\tau_{2y_{i2}}$), where $\tau_{2,0}$ and $\tau_{2,5}$ are given fixed values of $-\infty$ and $\infty$, respectively. The limits of integration for the inner most integral are the upper and lower bounds of the interval corresponding to the rating reported by expert 1 for colony $i$, determined from the corresponding thresholds as ($\tau_{1(y_{i1}-1)}$, $\tau_{1y_{i1}}$), where $\tau_{1,0}$ and $\tau_{1,5}$ are given fixed values of $-\infty$ and $\infty$, respectively. If expert $j$ did not provide a rating for colony $i$, the corresponding upper and lower limits of integration are set to $-\infty$ and $\infty$, respectively.

### VI. Additional Details about using Cross Validation

To reduce the risk of over-fitting, we employed 10-fold cross-validation to produce predicted values from the linear scoring algorithms that were then used evaluate performance via receiver operating characteristic curves. The collection of colonies was randomly partitioned into 10 sets, with a roughly equal number of colonies in each set. To obtain predicted scores under cross validation, a given set was held out as a test set and the parameter values of the model were fit using maximum likelihood, applied to the training data from colonies within the nine other sets. The resulting feature weights and intercept were then used to obtain predicted scores for each of the colonies in the set that was held out during parameter fitting. This process was repeated so that each set of colonies was held out in turn, with each colony in the set receiving its predicted score during its turn as the test set.

For random forest models, the final predicted score for each colony is given by the average predicted scores among trees for which the colony was omitted by the random resampling process.

### VII. Additional Details about Fitting Thresholds

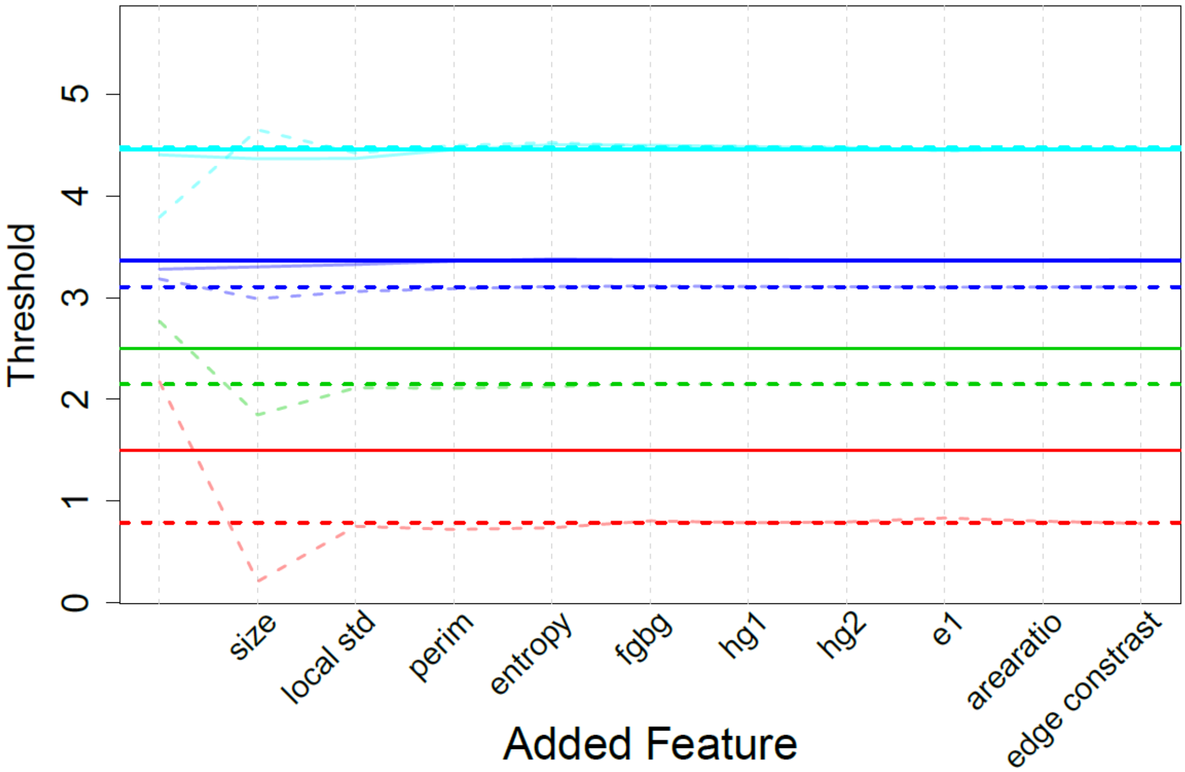


**Supplemental Figure 4.** Threshold values obtained via maximum likelihood during forward selection. The x-axis indicates the order in which features were added to the model. The faint lines in the plot above shows the value of $\tau_{jk}$ obtained by maximum likelihood for each of the linear models obtained by forward selection, beginning with no features (0) and culminating in a model with ten features. The median value of each threshold across the 11 models is depicted as a horizontal line in bold color.

To prevent over parameterization, we fixed the first two thresholds of the first expert to be $\tau_{1,1}$ =1.5 and $\tau_{1,2}$ =2.5 in all models and fit the intercept, image feature weights, variance, and other thresholds to maximize the likelihood around this constraint. If no parameters were fixed, the same likelihood could be obtained for any linear transformations of the scale for latent scores, and the model (including feature weights, thresholds, and components of the variance-covariance matrix) would be unidentifiable. As seen in **Supplementary Figure 4,** the fitted thresholds were each very stable among models with two or more features. We then refit the weights for each of the 11 models obtained during forward-selection after fixing each threshold to the median value across the 11 models.

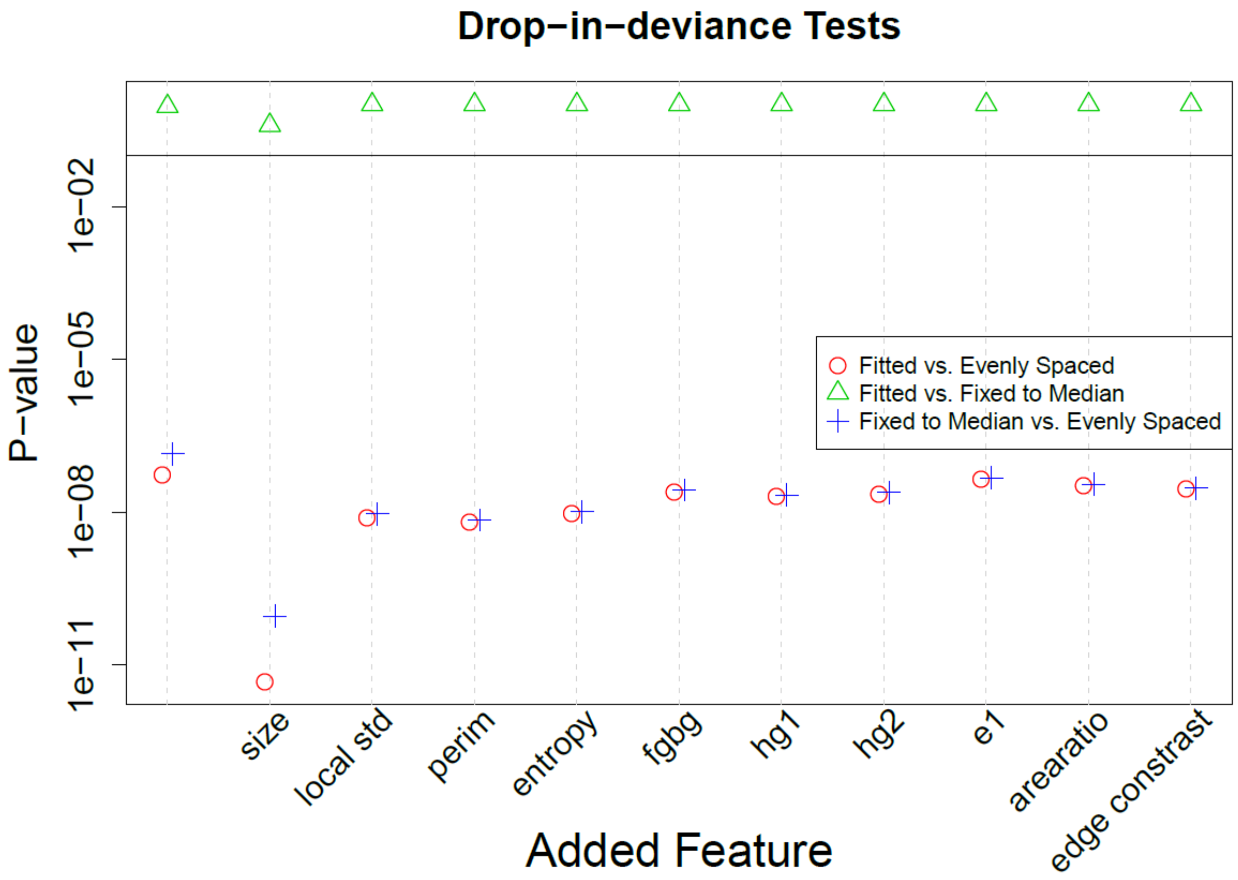


**Supplemental Figure 5.** P-values resulting from drop-in-deviance tests that compare models that include the same image features but place different constraints on the threshold values. "Fitted" denotes models in which the values of six threshold parameters may vary freely during likelihood maximization (the first two thresholds for expert 1 are fixed at 1.5 and 2.5, respectively, to make the model identifiable; see below). "Evenly Spaced" denotes models in which the thresholds were fixed at 1.5, 2.5, 3.5 and 4.5, respectively, for both experts. "Fixed to Median" denotes models in which each threshold parameter was assigned the median of the 11 values obtained for that parameter under the "Fitted" models. The x-axis indicates the order in which features were added to the model. The drop-in-deviance test statistic was computed as $|l_1 - l_2|$, where $l_m$ ($m$ =1,2) denotes the respective log likelihoods obtained for the two models being compared. P-values were computed as the right tail of a chi-squared distribution with six degrees of freedom.

As shown in **Supplemental Figure 5**, conducting drop-in-deviance tests comparing the original models to the corresponding model that contains the same image features but fixes the thresholds to their respective median values produced p-values ranging between 0.392 and 0.999. This assessment suggests that using a common set of thresholds does not substantially adversely affect the model fit.  Using a common set of thresholds helps place other model parameters such as image feature weights and the variance parameters onto a common scale across models with different numbers of image features, which greatly aids comparison between models of different sizes.

For each of the forward selection models, a set of evenly spaced thresholds (i.e., $\tau_{j1}$= 1.5, $\tau_{j2}$= 2.5, $\tau_{j3}$=3.5 and $\tau_{j4}$=4.5 for j=1,2) was rejected by the drop-in-deviance test when compared to either the corresponding model with fitted thresholds (as obtained in the first round of fitting) or the corresponding model with thresholds fixed to the median values.  (P-values ranging from $4.84 \times 10^{-12}$ to $1.41 \times 10^{-7}$; see **Supplementary Figure 5**). This analysis suggests that using a model that naively fixes evenly spaced thresholds results in a statistically significantly poorer fit to this data.

### VIII. Model evaluation with 56 feature set

The performance of the models developed with the 14 expert-inspired features plus the 42 off-the-shelf features (Haralick and wavelet) and trained on both experts' scores is shown in Supplementary Figure 6. The order in which features were added to the model is shown in Supplementary Figure 6A, together with the variance in the differences between the predicted colony scores and the latent scores of each expert, evaluated with the addition of each feature. The red and blue markers represent how far, on average, the model's predicted scores are from the putative (latent) scores of the respective expert over all colonies. The green markers are a measure of the direct discordance between the experts' scoring ranges over all colonies. This quantity provides a benchmark against which to evaluate scoring algorithm performance. When the red markers and/or the blue markers are less than green markers, the variance-covariance estimates indicate that the disparity between the scoring algorithm and the latent scores of the expert(s) would be less than the disparity between the latent scores of the two experts. The covariance in the differences between each expert and the model predictions is shown with the purple markers. The covariance serves as a measure of how much information is missing in the model; for an ideal model that captured all the patterns with features of interest that are common to both experts, the expected covariance in residuals between experts would be 0.

Compared with the prediction model based solely on the 14 expert-inspired features, the addition of the 42 off-the-shelf features (Haralick and wavelet) provides no significant improvement to the model performance for predicting either of the two experts (red and blue markers), nor does it reduce the covariance in residual between experts (purple markers). Additionally, because the Haralick and wavelet features are difficult to interpret, they don't add intuitive knowledge of colony characteristics.

Receiver Operator Characteristic (ROC) curves shown in Supplementary Figure 6B were constructed using predicted scores obtained via 10-fold cross validation. The prediction models accessed the full 56 feature set (14 expert-inspired features and 42 off-the-shelf features) and were trained on both experts' scores. The ROC curves assess how well a scoring algorithm identifies the highest rated colonies for each expert and is intended to reflect the envisioned usage of a pluripotency scoring algorithm to select the most promising colonies. The ROC curves in Supplementary Figure 6B indicate that addition of the 42 off-the-shelf features (Haralick and wavelet) provides no significant improvement to the model performance compared the expert-inspired features alone. Using a random forest modeling approach, evaluated using the complete set of 56 image features, did not improve ROC performance over the linear models.

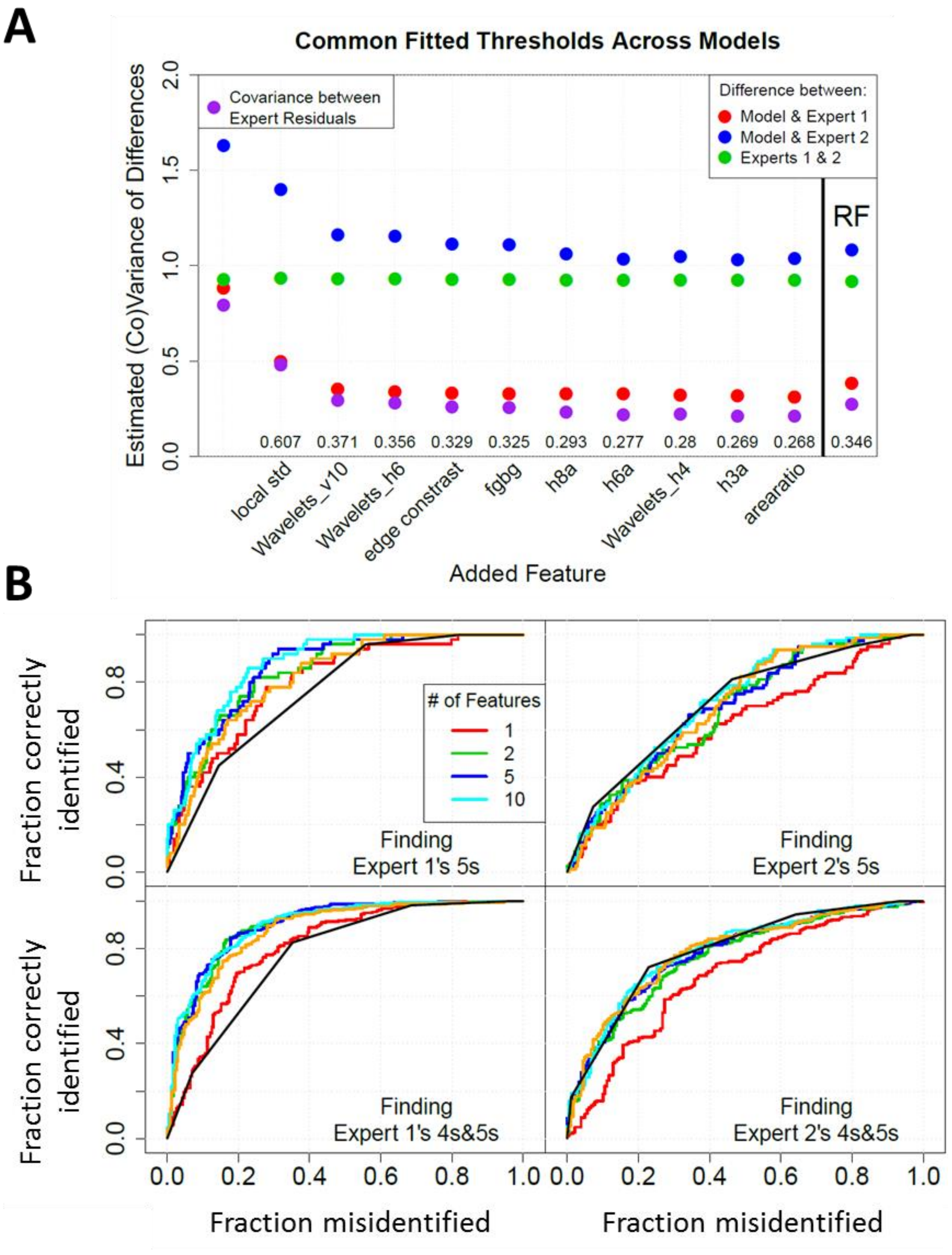


**Supplemental Figure 6.** Fitting results from linear models with different numbers of features trained on scores from both experts and accessing 56 features (14 expert-inspired features and 42 off-the-shelf Haralick and wavelet features). A. Estimated variances and covariances in the differences between expert scores and model predictions. Shown are results of 10 models, where each new model includes all features from the previous model and is created by the addition of the indicated feature. This plot represents the variance in the differences between the predicted scores from the models and the scores from expert 1 (red) and expert 2 (blue). The purple markers indicate covariance in the differences calculated between the experts scores and the model predictions. The green markers indicate the variance between the experts that is calculated according to Equation 1 in the Main Text. B. ROC curves for the model predictions of colonies given a score of 5 or 4-5 and compared to scores from expert 1 and from expert 2. The fraction correctly identified = (true positives/(true positives + false negatives)) and the fraction misidentified = (false positives/(false positives + true negatives)). The orange curve is the result of RF fit to 56 features. The black ROC curve is computed from the confusion matrix of expert 1 and expert 2 ratings.
