## Supplemental Information for "A Latent Variable Model for Evaluation of Disparate Ratings of Stem Cell Colonies by Two Experts"

This pdf file contains all 148 images that experts rated, and the ratings for each from each expert.

Well = 1, Colony = 1
Rater 1 = 2
Rater 2 = 1

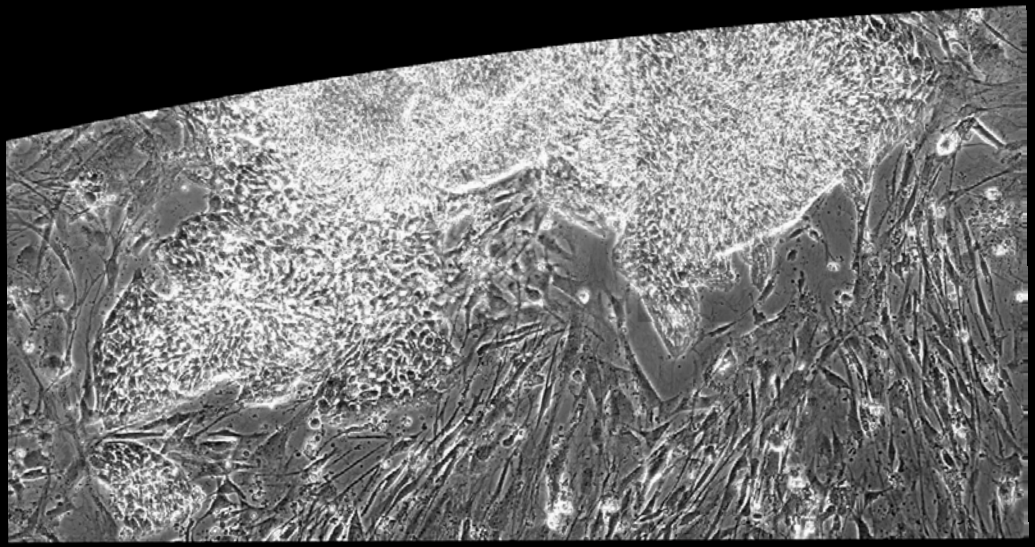

Well = 1, Colony = 2
Rater 1 = 3
Rater 2 = 3

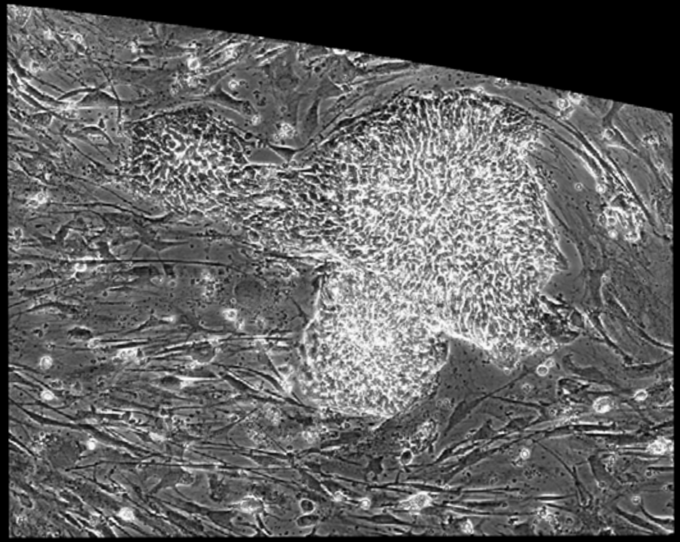

Well = 1, Colony = 3
Rater 1 = 3
Rater 2 = 4

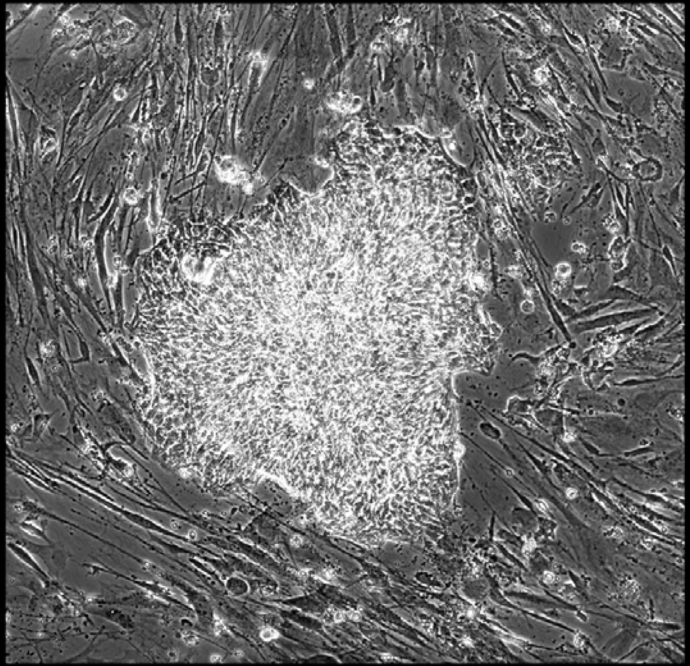

Well = 1, Colony = 4
Rater 1 = 4
Rater 2 = 3

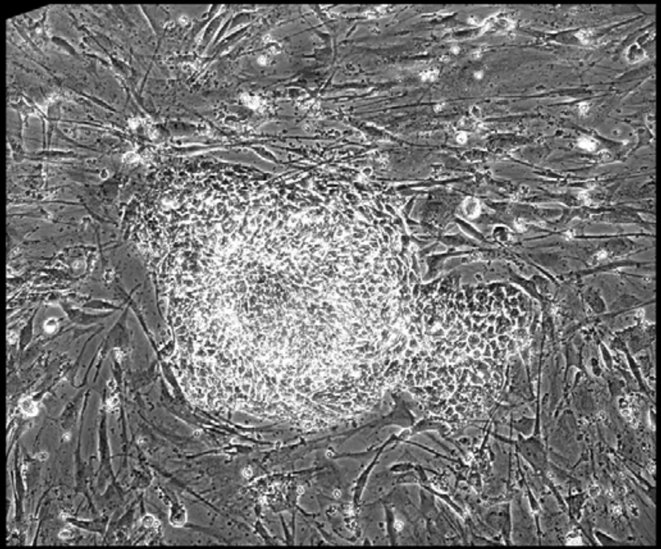

Well = 1, Colony = 5
Rater 1 = not scored
Rater 2 = not scored

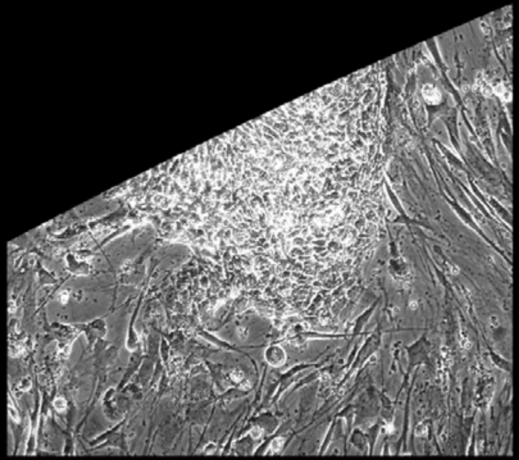

Well = 1, Colony = 6
Rater 1 = 3
Rater 2 = 1

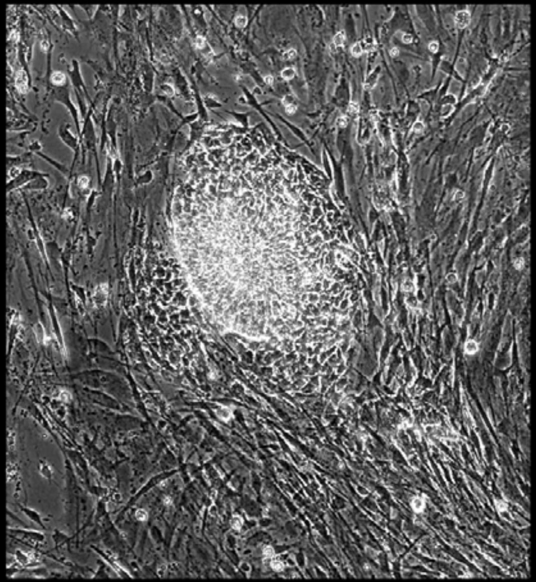

Well = 1, Colony = 7
Rater 1 = 4
Rater 2 = 3

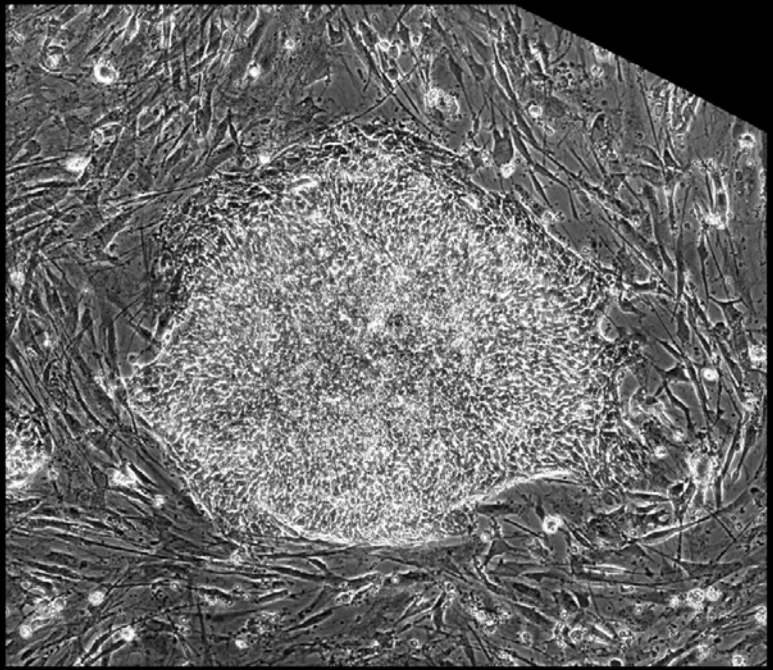

Well = 1, Colony = 8
Rater 1 = 4
Rater 2 = 3

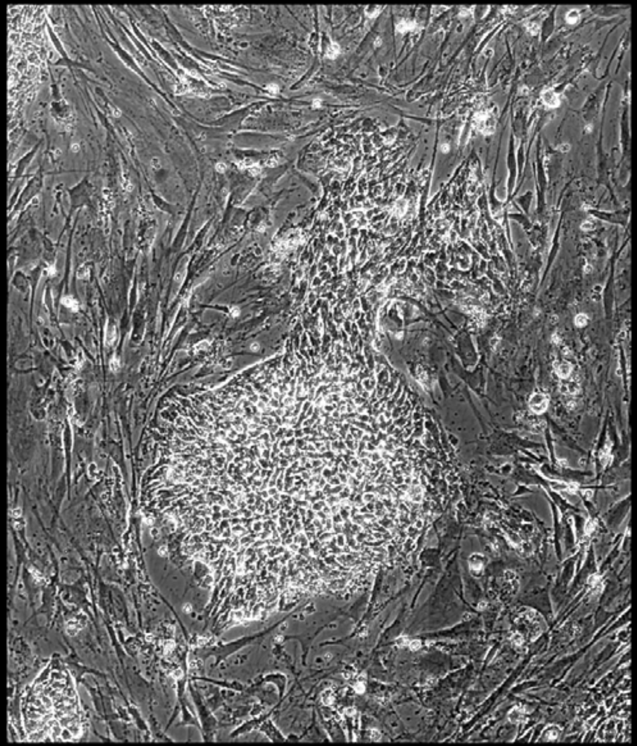

Well = 1, Colony = 9
Rater 1 = 5
Rater 2 = 5

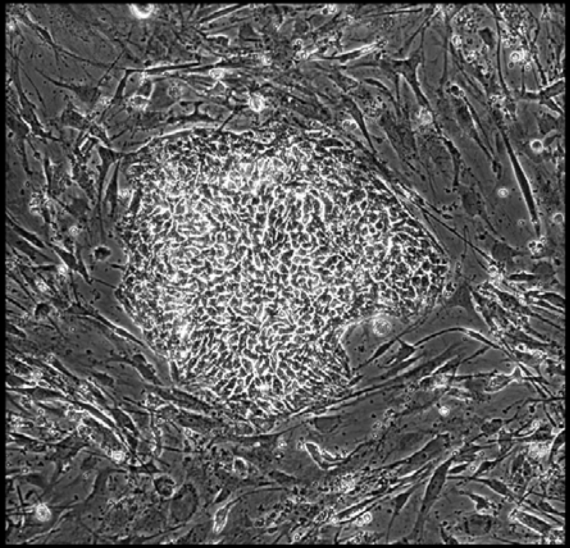

Well = 1, Colony = 10
Rater 1 = 5
Rater 2 = 5

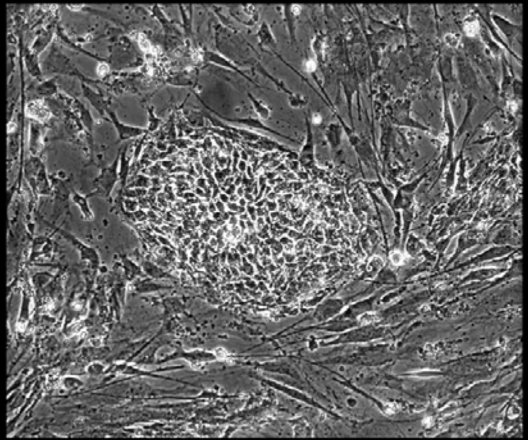

Well = 1, Colony = 11
Rater 1 = 5
Rater 2 = 5

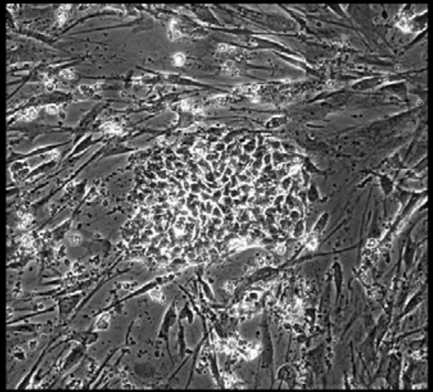

Well = 1, Colony = 12
Rater 1 = 4
Rater 2 = 4

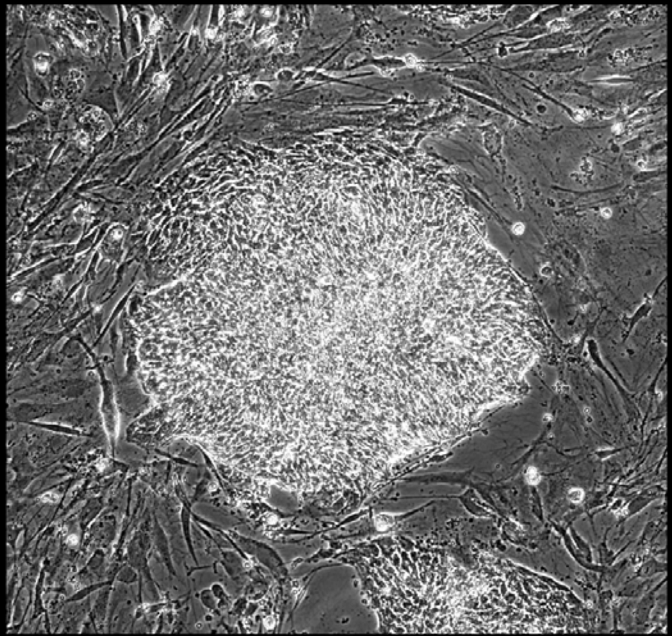

Well = 1, Colony = 13
Rater 1 = 5
Rater 2 = 4

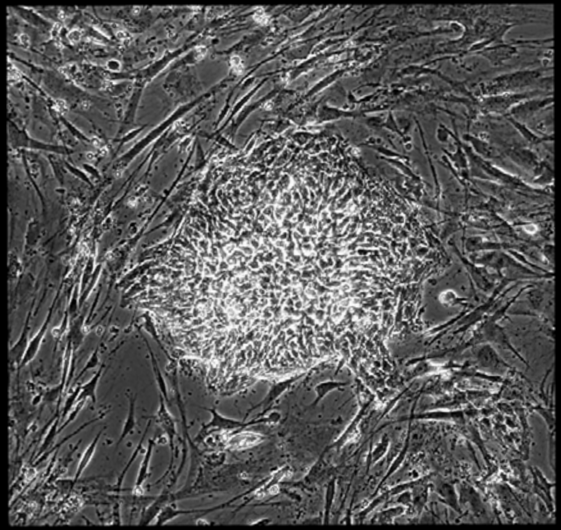

Well = 1, Colony = 14
Rater 1 = 4
Rater 2 = 2

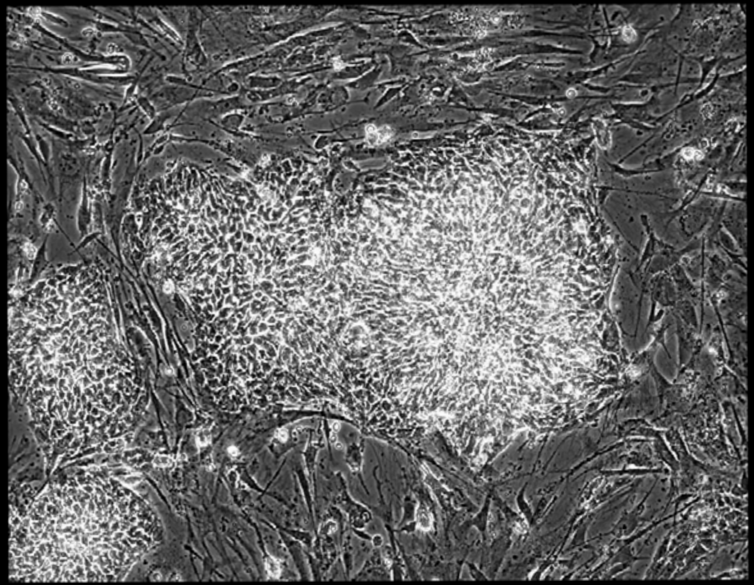

Well = 1, Colony = 15
Rater 1 = 4
Rater 2 = 5

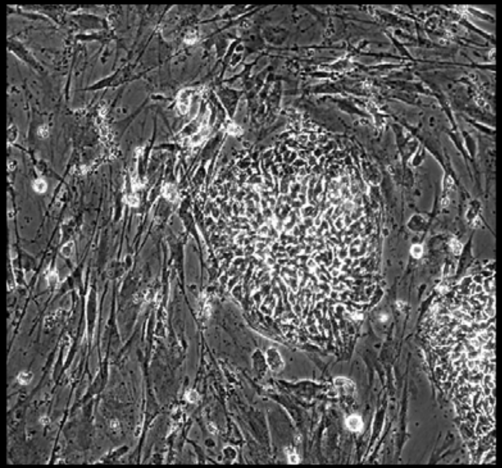

Well = 1, Colony = 16
Rater 1 = 4
Rater 2 = 4

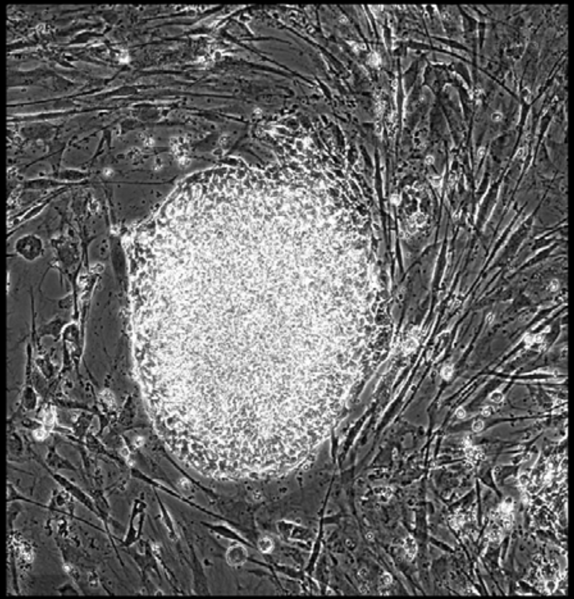

Well = 1, Colony = 17
Rater 1 = 5
Rater 2 = 5

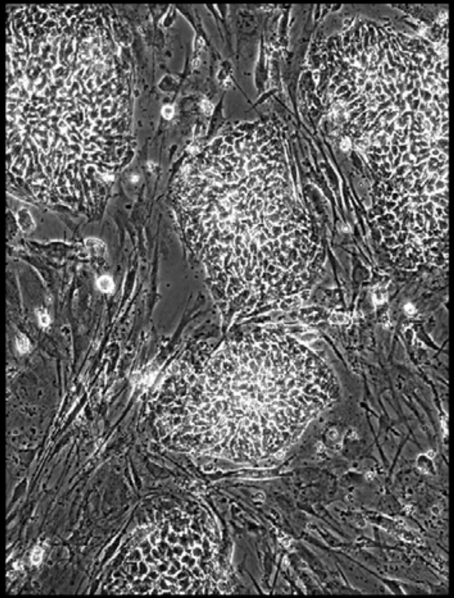

Well = 1, Colony = 18
Rater 1 = 5
Rater 2 = 4

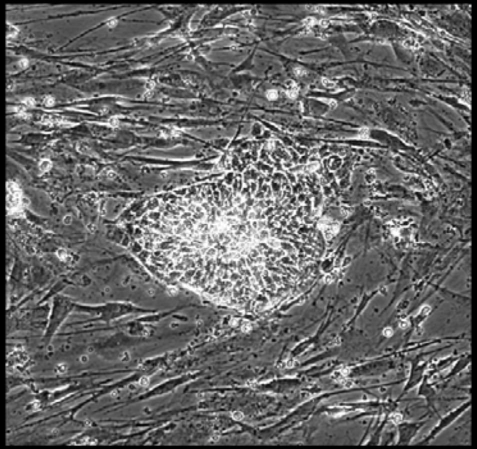

Well = 1, Colony = 19
Rater 1 = 2
Rater 2 = 2

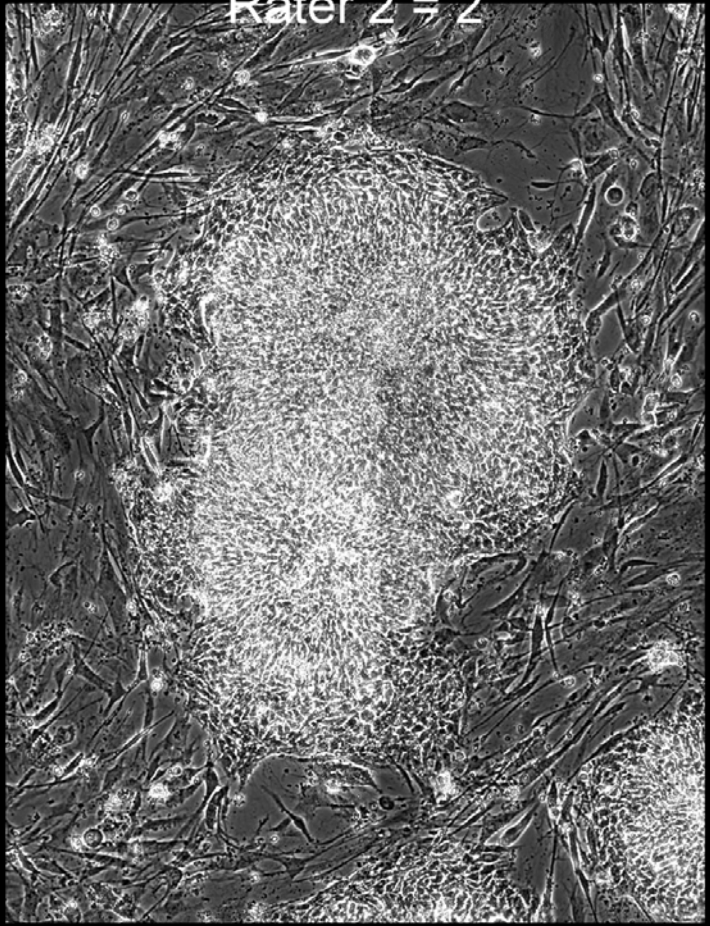

Well = 1, Colony = 20
Rater 1 = 3
Rater 2 = 4

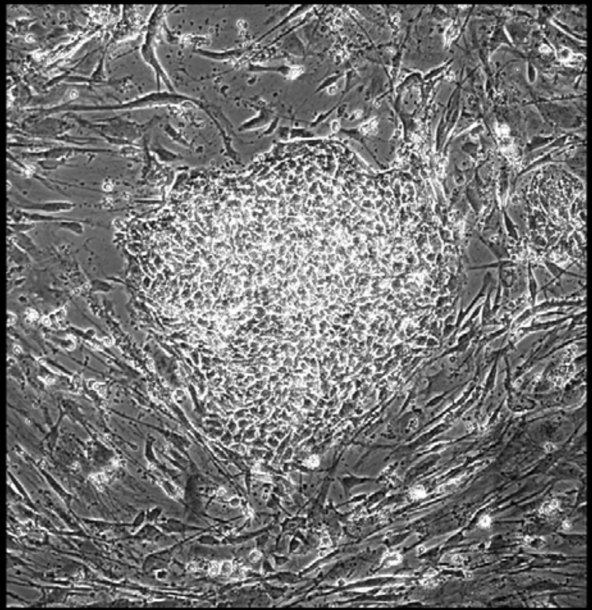

Well = 1, Colony = 21
Rater 1 = 3
Rater 2 = 3

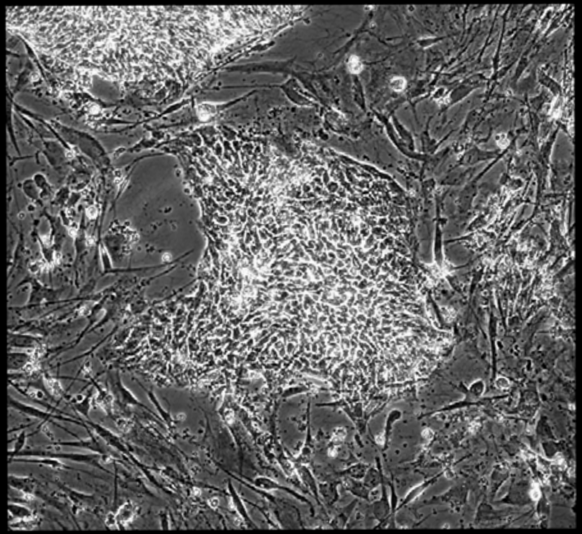

Well = 1, Colony = 22
Rater 1 = 5
Rater 2 = 4

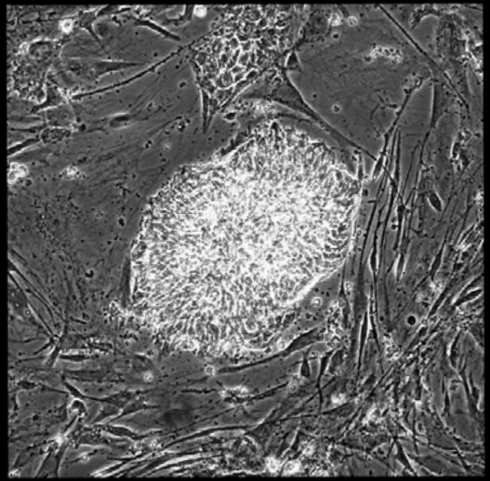

Well = 1, Colony = 23
Rater 1 = 3
Rater 2 = 2

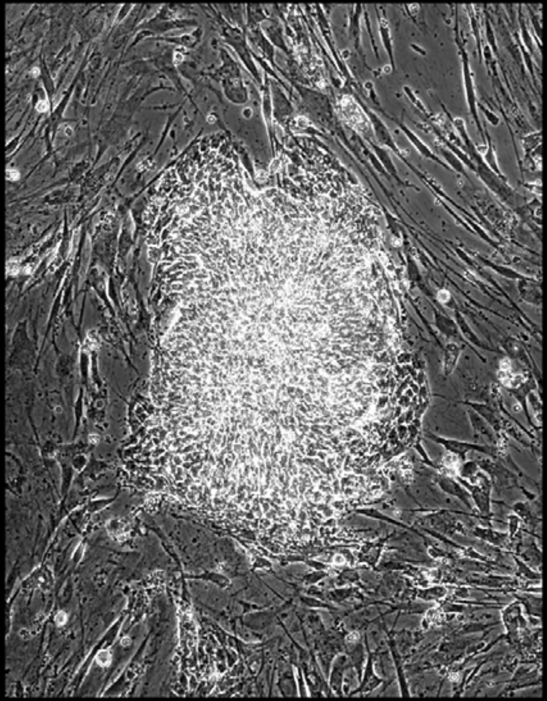

Well = 1, Colony = 24
Rater 1 = 3
Rater 2 = 3

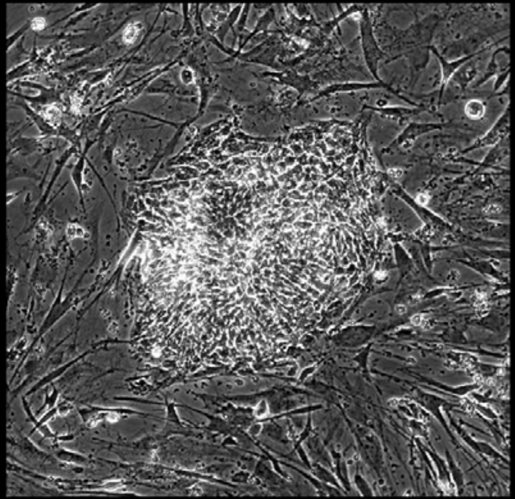

Well = 1, Colony = 25
Rater 1 = 3
Rater 2 = 2

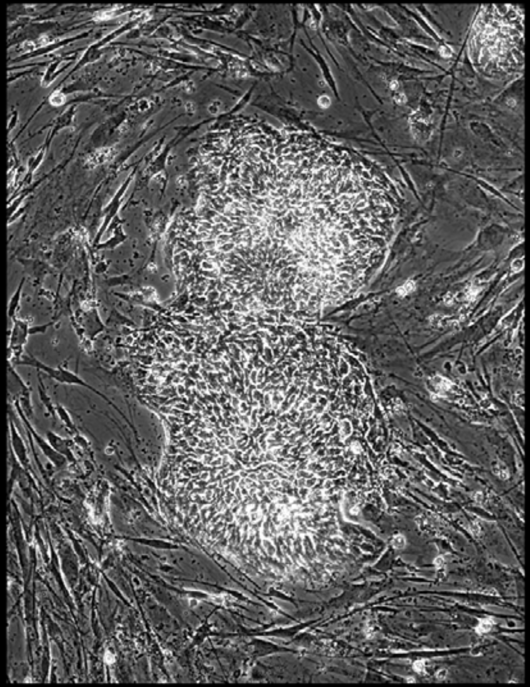

Well = 1, Colony = 26
Rater 1 = 2
Rater 2 = not scored

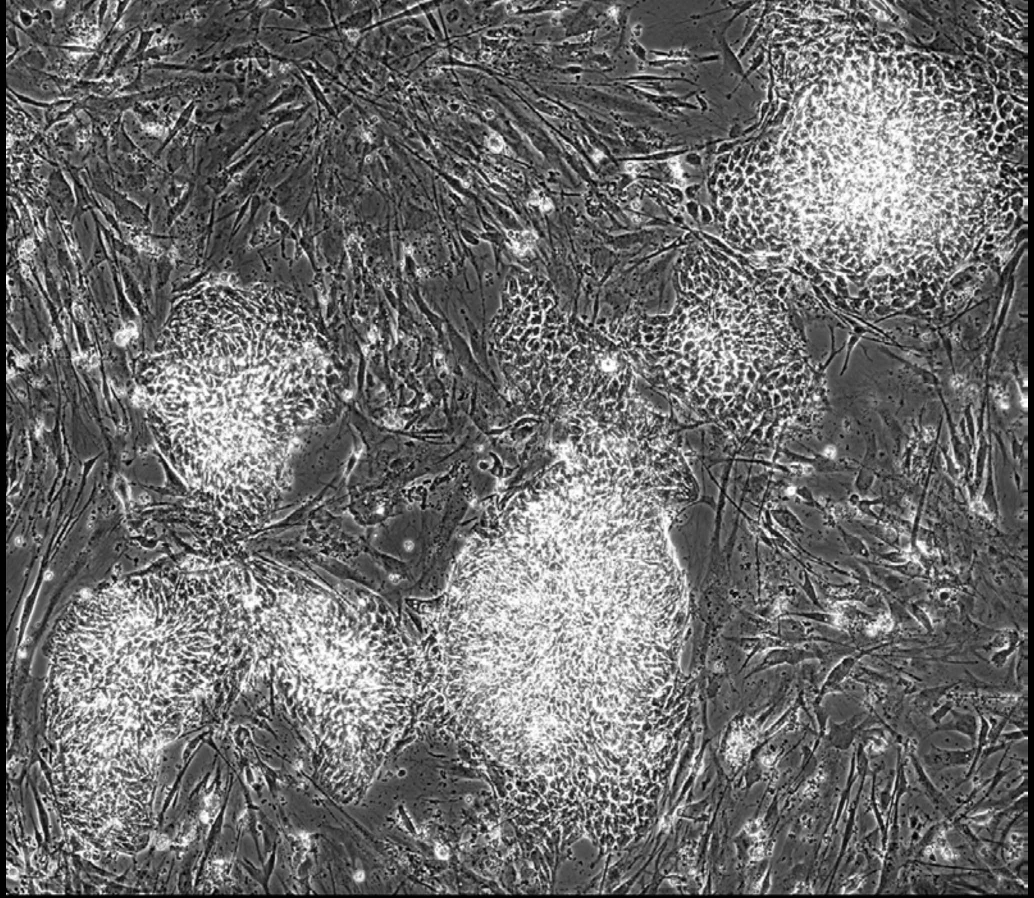

Well = 1, Colony = 27
Rater 1 = 5
Rater 2 = 3

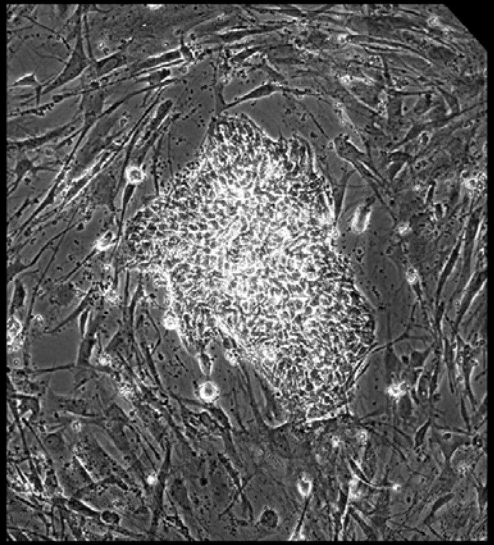

Well = 1, Colony = 28
Rater 1 = 4
Rater 2 = 3

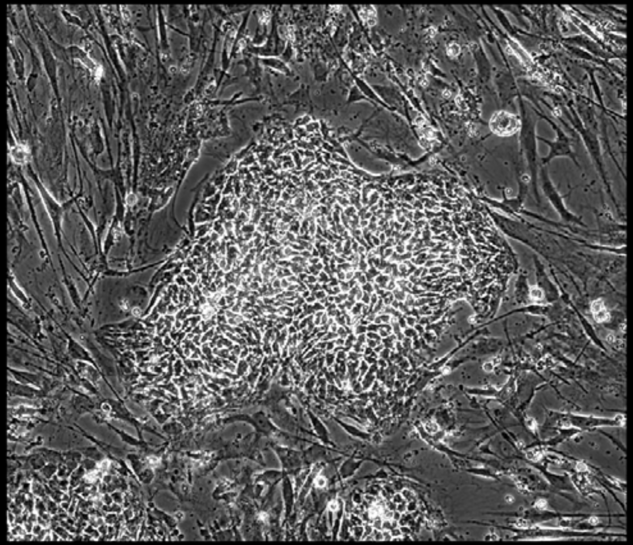

Well = 1, Colony = 29
Rater 1 = 3
Rater 2 = 3

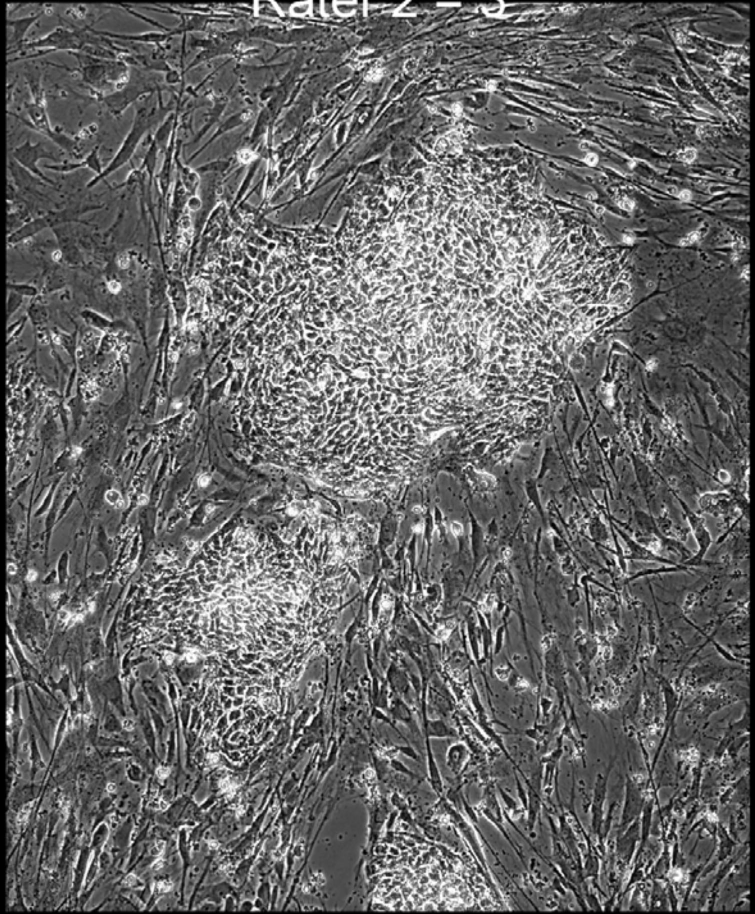

Well = 1, Colony = 30
Rater 1 = 5
Rater 2 = 4

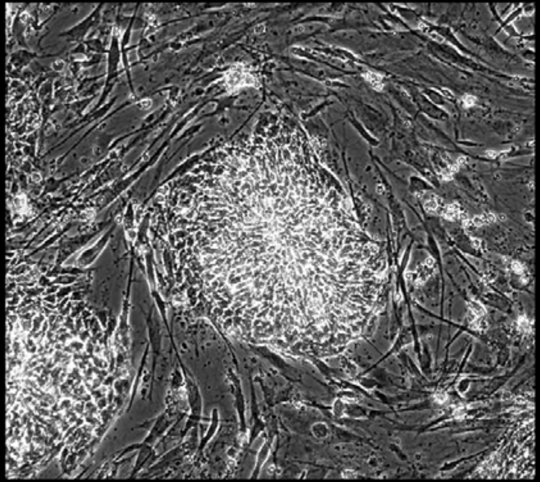

Well = 1, Colony = 31
Rater 1 = 4
Rater 2 = 5

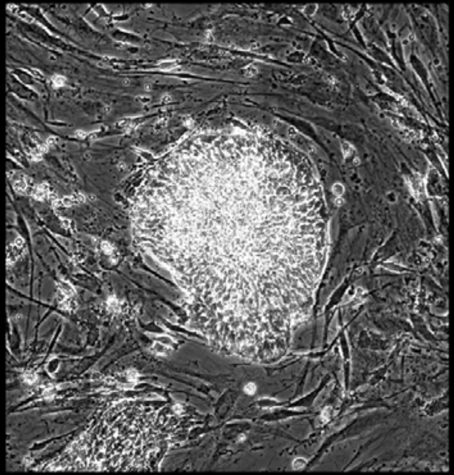

Well = 1, Colony = 32
Rater 1 = 3
Rater 2 = 3

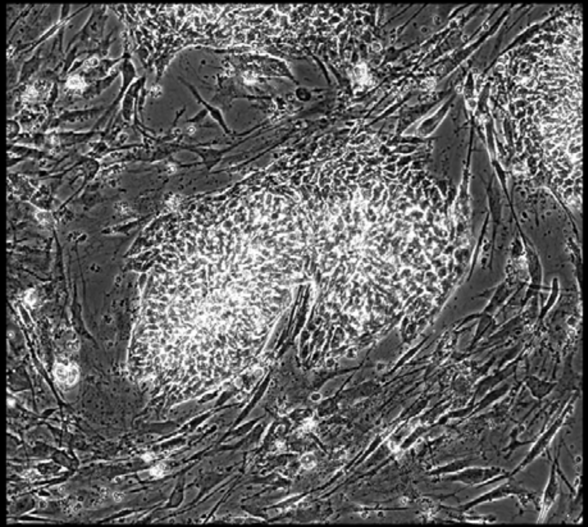

Well = 1, Colony = 33
Rater 1 = 5
Rater 2 = 3

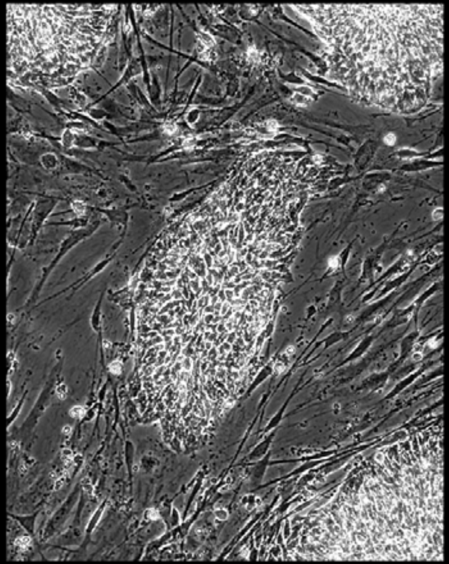

Well = 1, Colony = 34
Rater 1 = 4
Rater 2 = 2

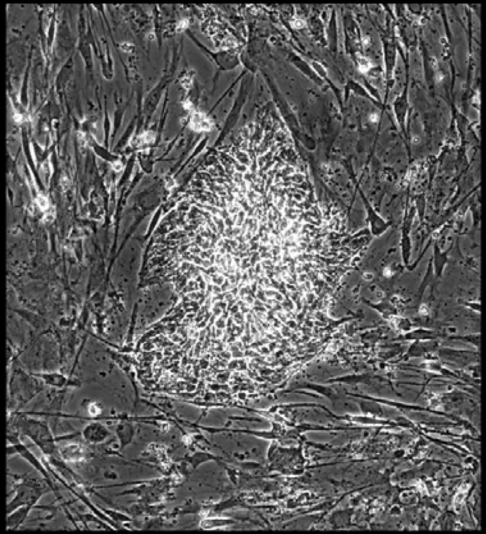

Well = 1, Colony = 35
Rater 1 = 3
Rater 2 = 2

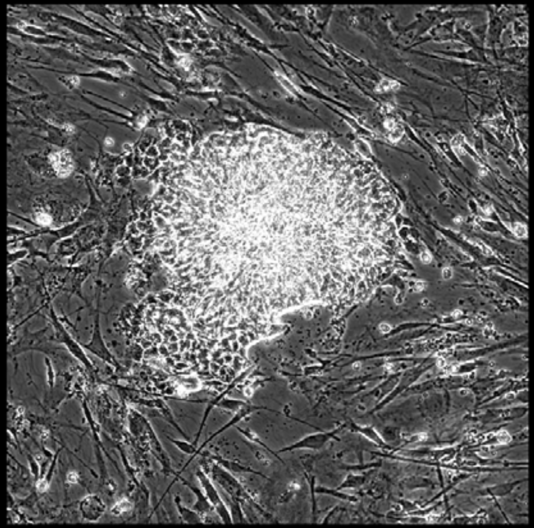

Well = 1, Colony = 36
Rater 1 = 1
Rater 2 = 1

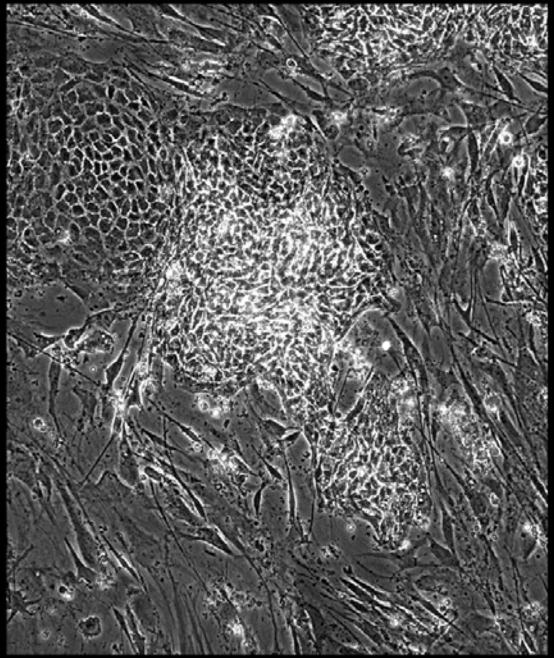

Well = 1, Colony = 37
Rater 1 = 3
Rater 2 = 3

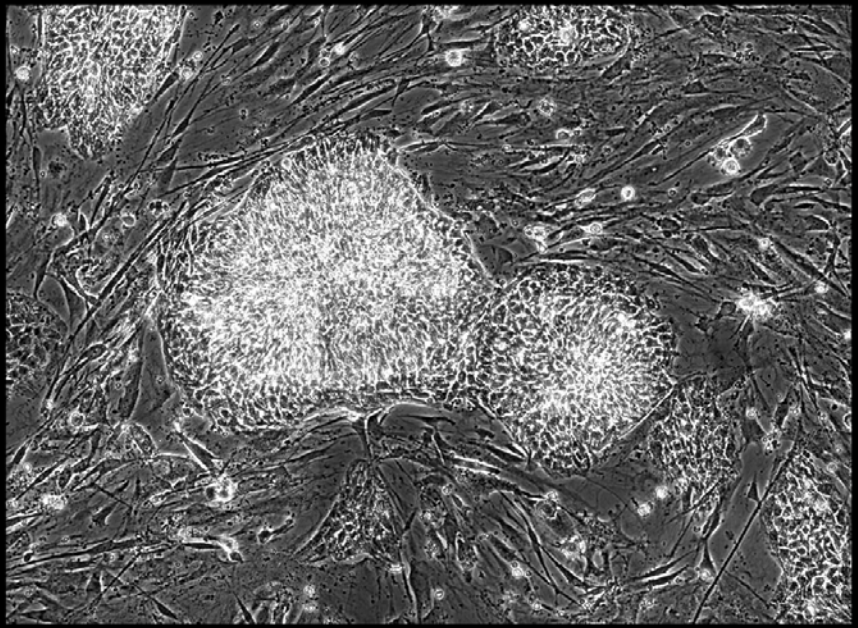

Well = 1, Colony = 38
Rater 1 = 4
Rater 2 = 4

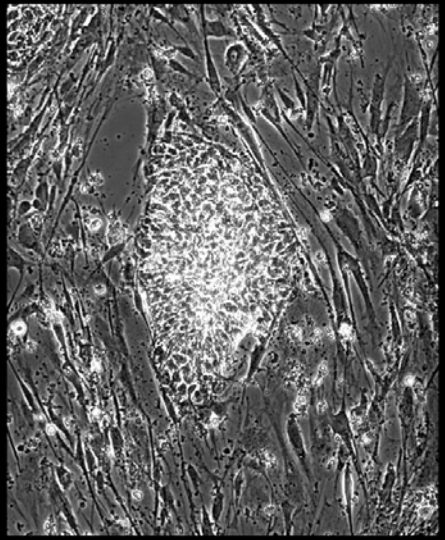

Well = 1, Colony = 39
Rater 1 = 4
Rater 2 = 3

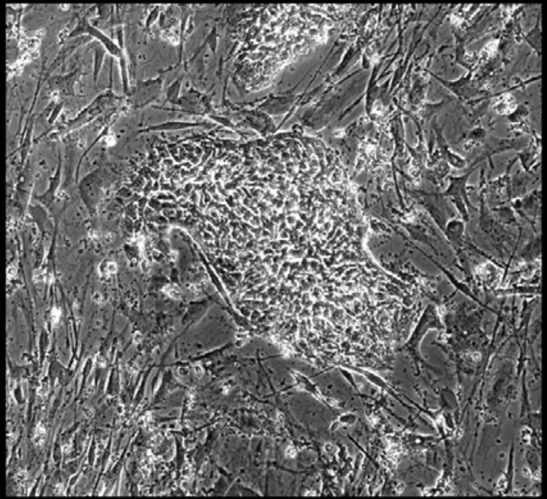

Well = 1, Colony = 40
Rater 1 = 4
Rater 2 = 5

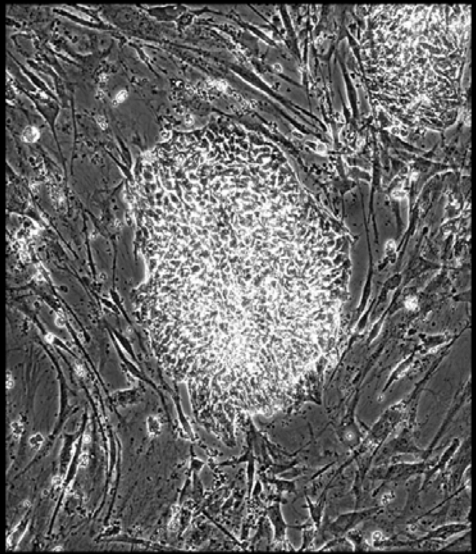

Well = 1, Colony = 41
Rater 1 = 4
Rater 2 = 4

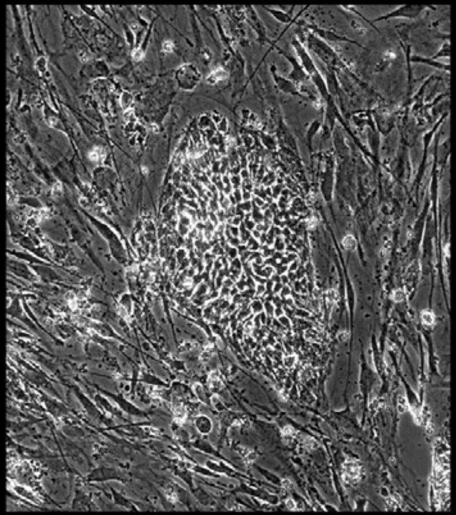

Well = 1, Colony = 42
Rater 1 = not scored
Rater 2 = not scored

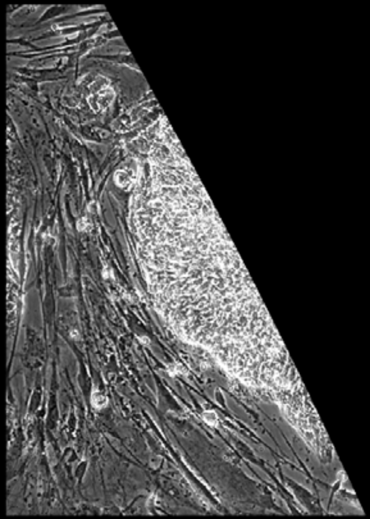

Well = 1, Colony = 43
Rater 1 = 3
Rater 2 = 3

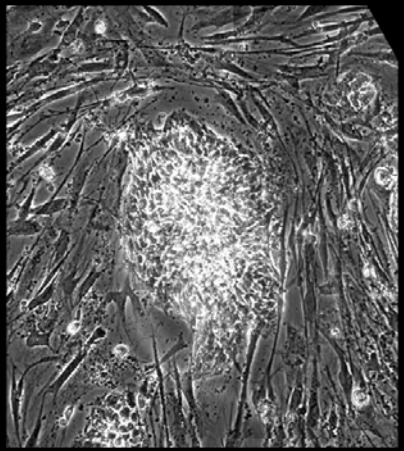

Well = 1, Colony = 44
Rater 1 = 4
Rater 2 = 4

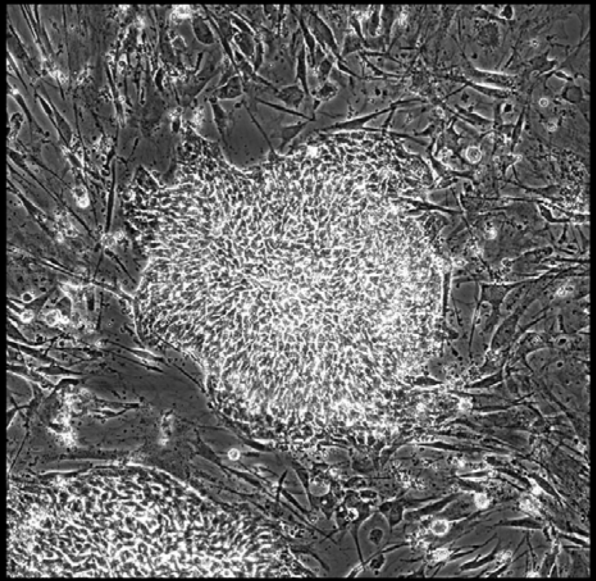

Well = 1, Colony = 45
Rater 1 = 1
Rater 2 = 1

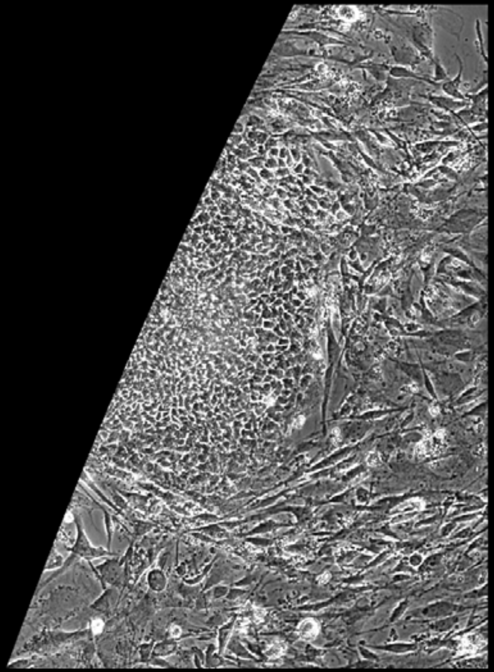

Well = 1, Colony = 46
Rater 1 = 4
Rater 2 = 4

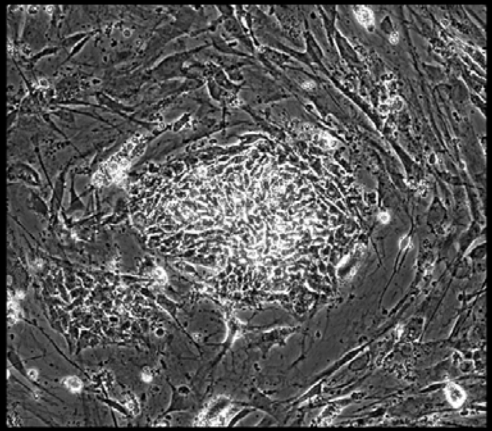

Well = 1, Colony = 47
Rater 1 = 3
Rater 2 = 4

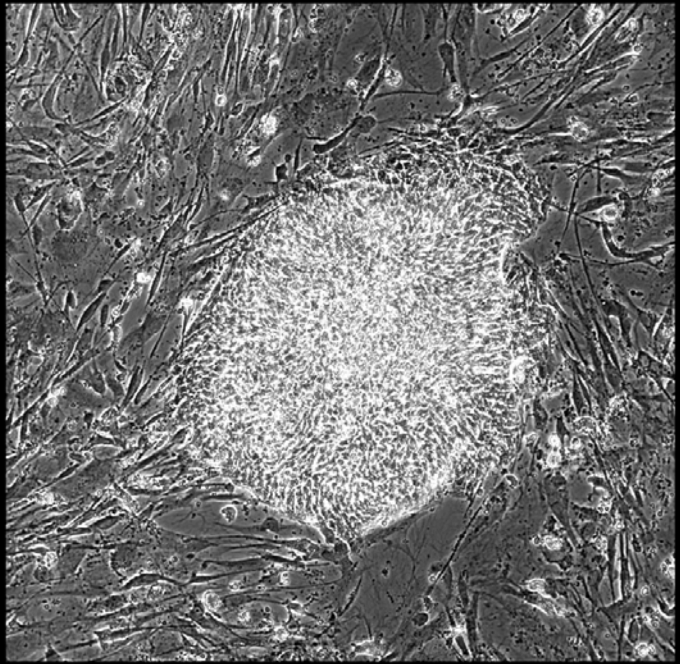

Well = 1, Colony = 48
Rater 1 = 4
Rater 2 = 2

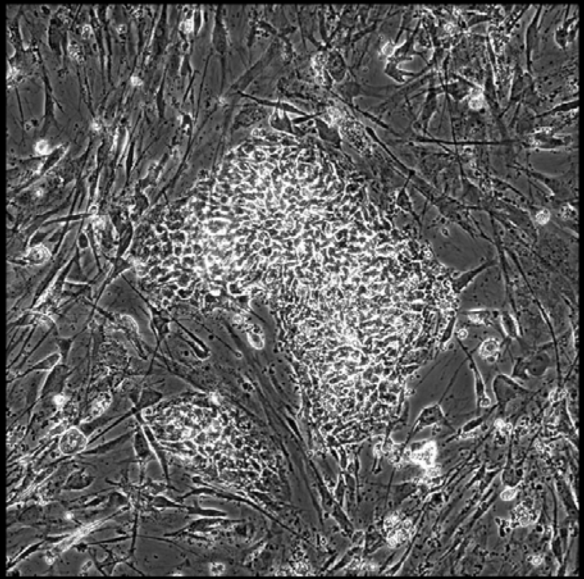

Well = 1, Colony = 49
Rater 1 = 3
Rater 2 = 3

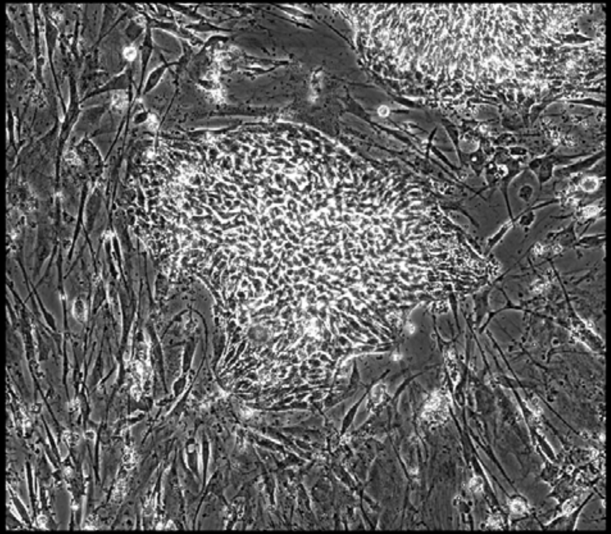

Well = 1, Colony = 50
Rater 1 = 3
Rater 2 = 3

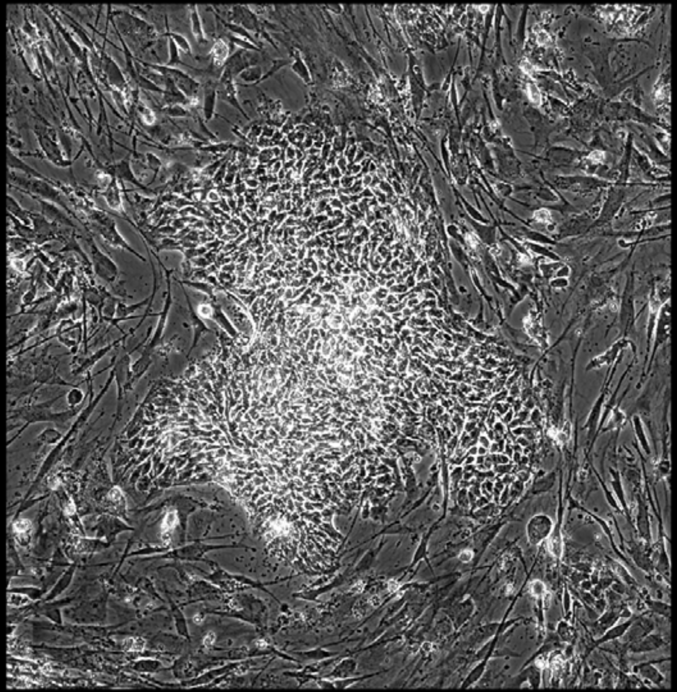

Well = 1, Colony = 51
Rater 1 = 4
Rater 2 = 4

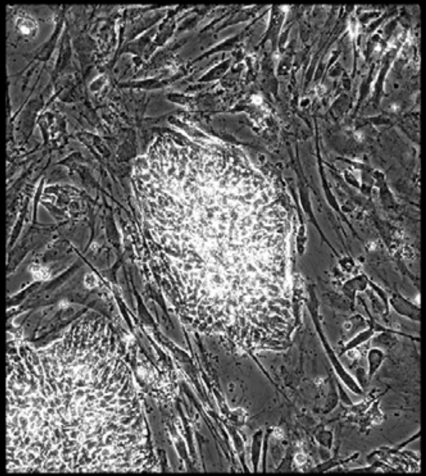

Well = 1, Colony = 52
Rater 1 = 3
Rater 2 = 2

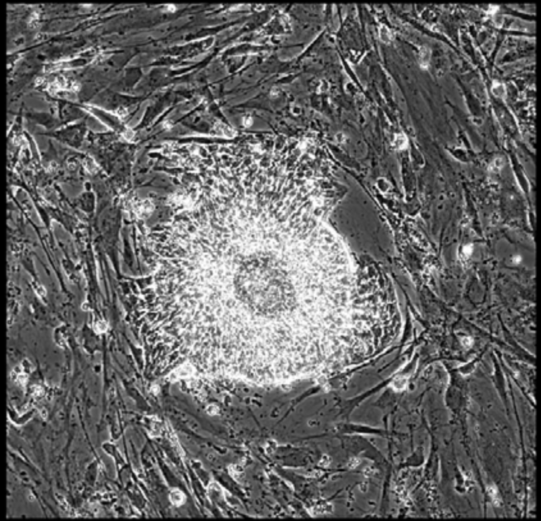

Well = 1, Colony = 53
Rater 1 = 3
Rater 2 = 3

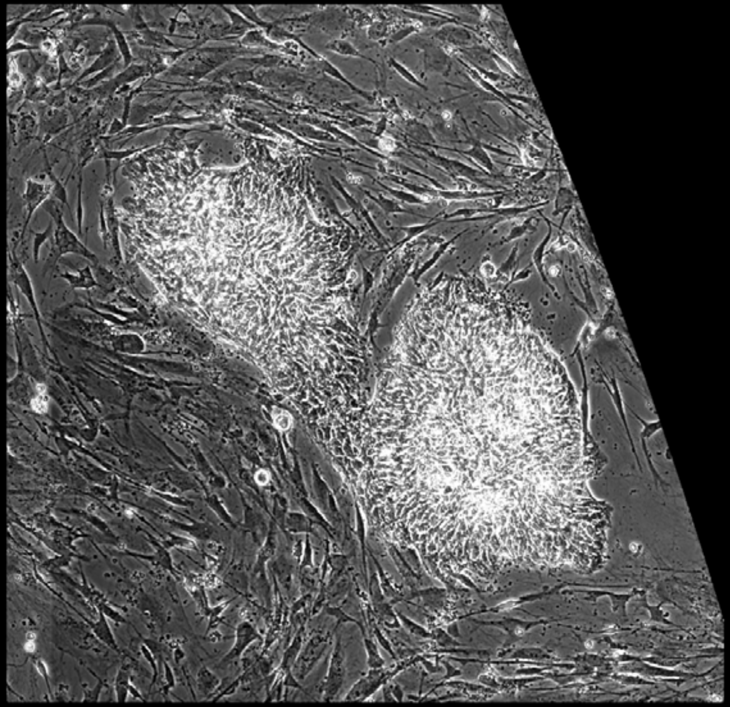

Well = 1, Colony = 54
Rater 1 = 4
Rater 2 = 4

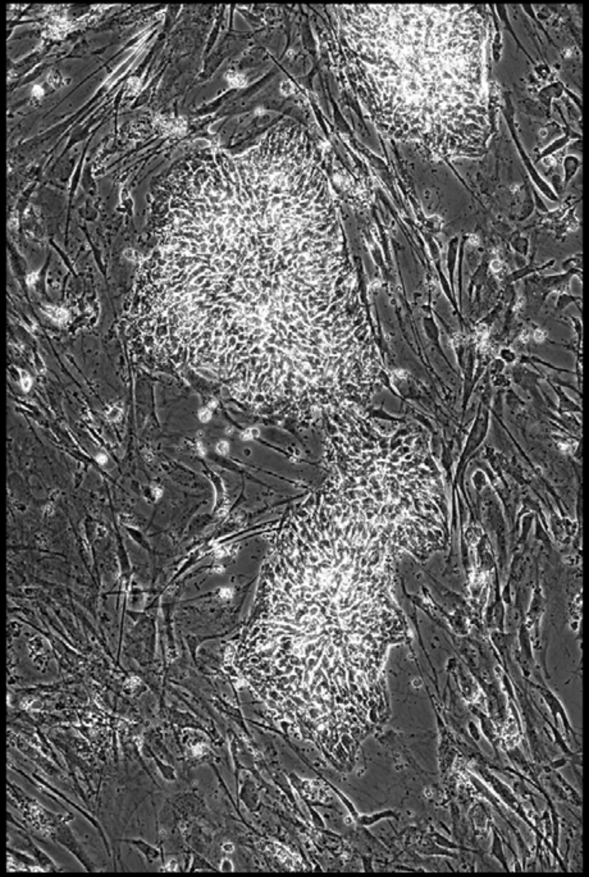

Well = 1, Colony = 55
Rater 1 = 2
Rater 2 = 4

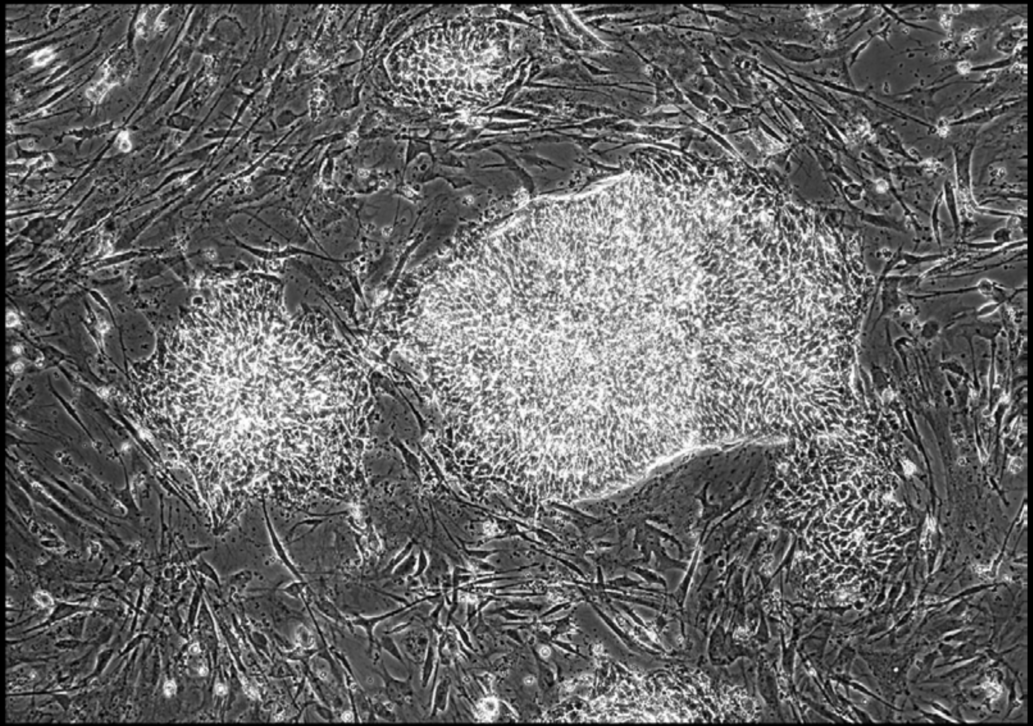

Well = 1, Colony = 56
Rater 1 = 3
Rater 2 = 2

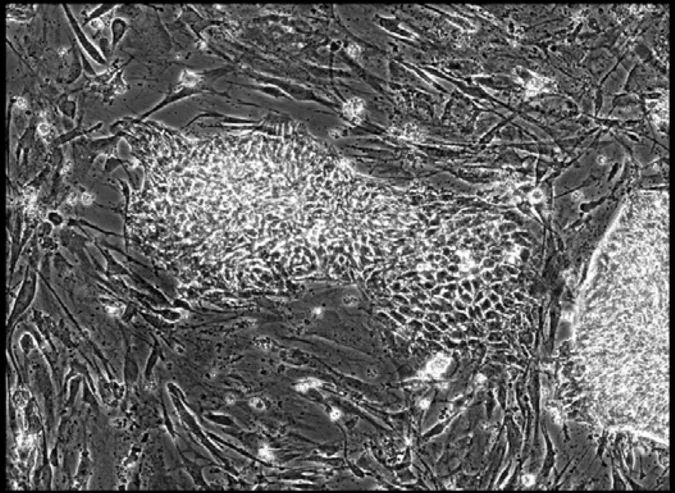

Well = 1, Colony = 57
Rater 1 = 2
Rater 2 = 2

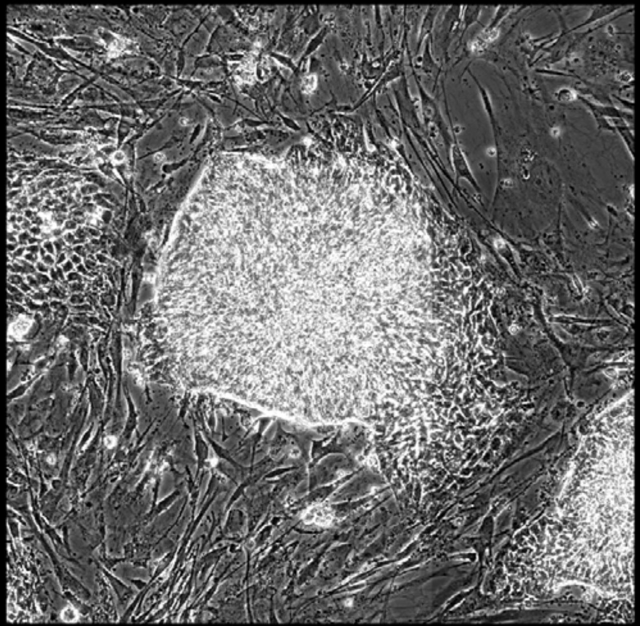

Well = 1, Colony = 58
Rater 1 = 5
Rater 2 = 4

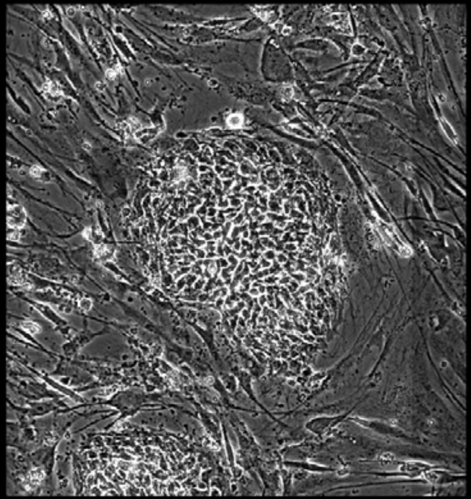

Well = 1, Colony = 59
Rater 1 = 5
Rater 2 = 4

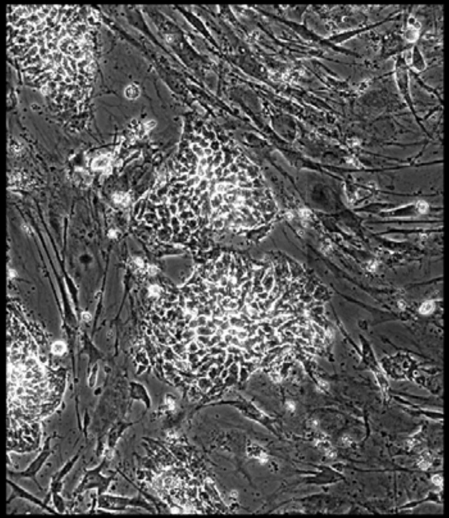

Well = 1, Colony = 60
Rater 1 = 3
Rater 2 = 2

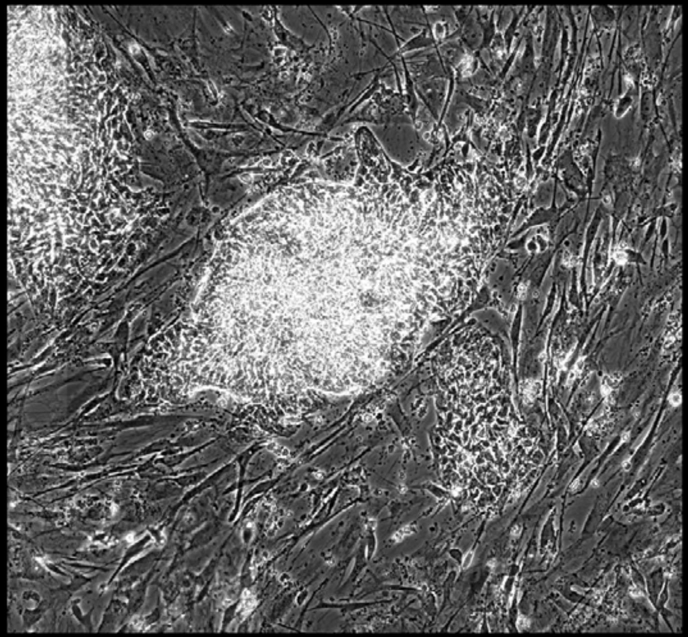

Well = 1, Colony = 61
Rater 1 = 5
Rater 2 = not scored

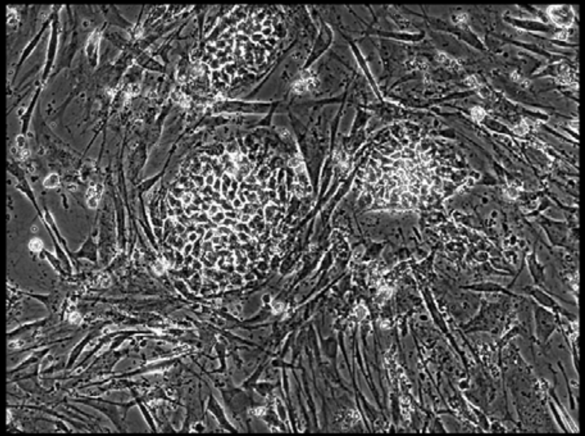

Well = 1, Colony = 62
Rater 1 = 5
Rater 2 = 4

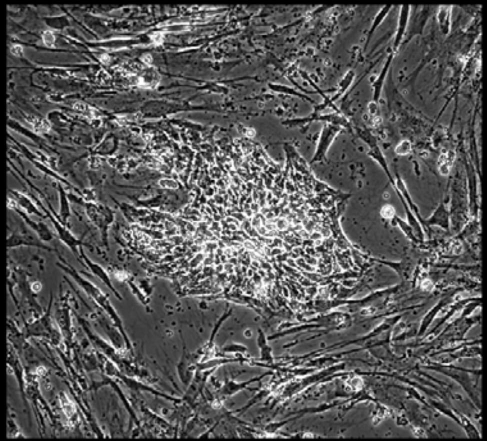

Well = 1, Colony = 63
Rater 1 = 5
Rater 2 = 4

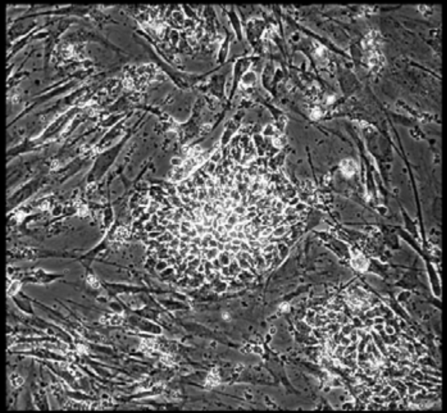

Well = 1, Colony = 64
Rater 1 = 3
Rater 2 = 3

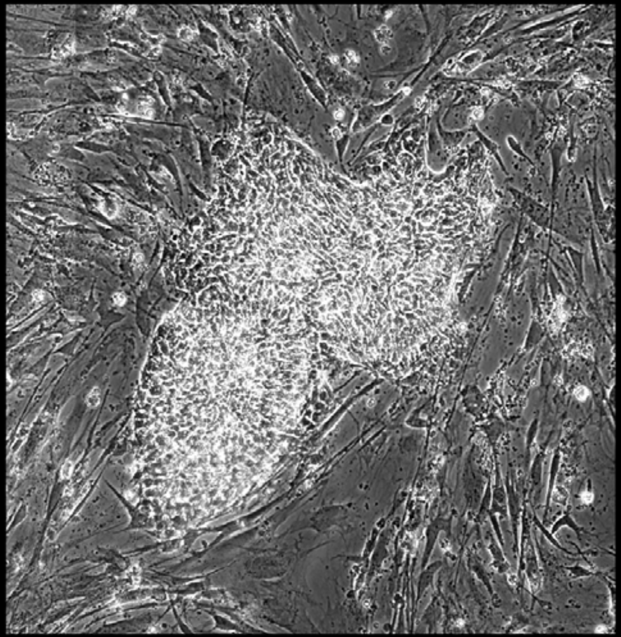

Well = 1, Colony = 65
Rater 1 = 2
Rater 2 = 4

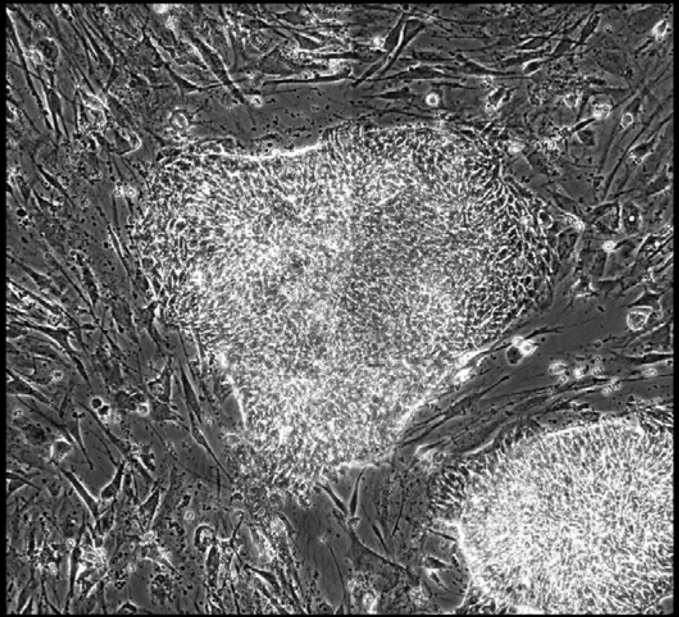

Well = 1, Colony = 66
Rater 1 = 4
Rater 2 = 5

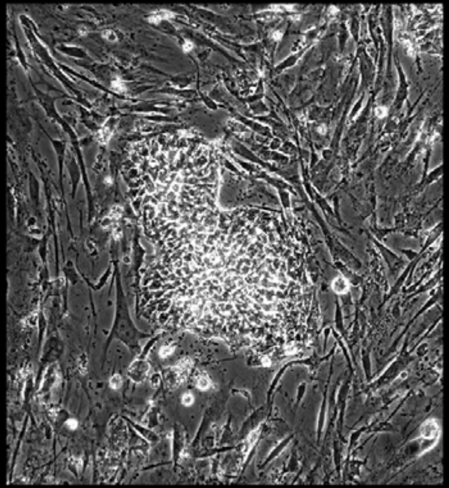

Well = 1, Colony = 67
Rater 1 = 5
Rater 2 = 5

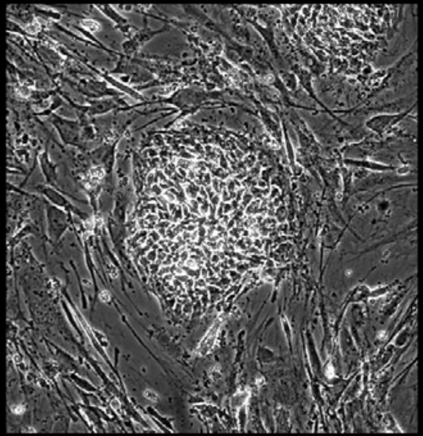

Well = 1, Colony = 68
Rater 1 = 3
Rater 2 = 4

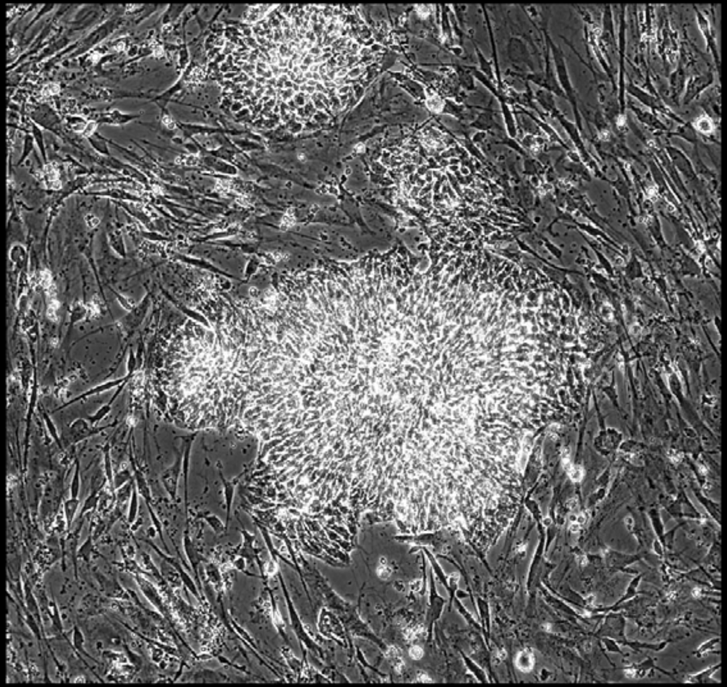

Well = 1, Colony = 69
Rater 1 = 3
Rater 2 = 3

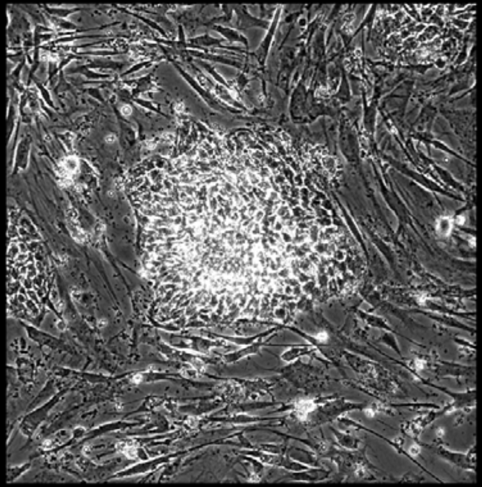

Well = 1, Colony = 70
Rater 1 = 3
Rater 2 = 3

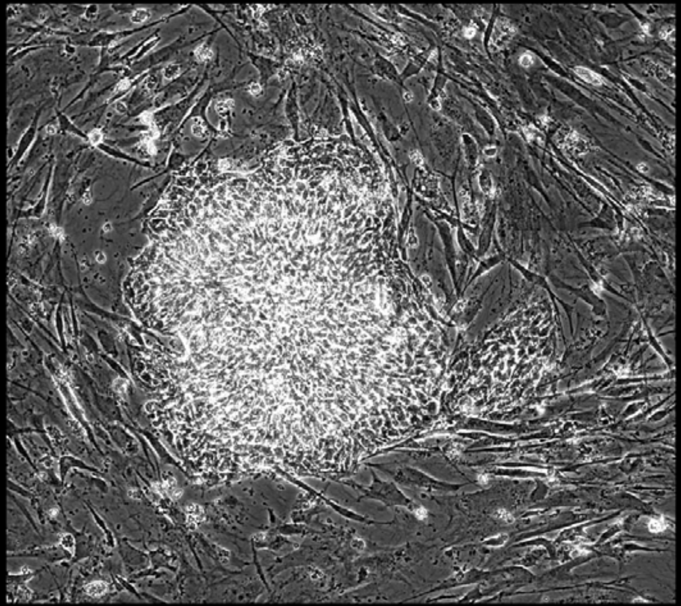

Well = 1, Colony = 71
Rater 1 = 3
Rater 2 = 4

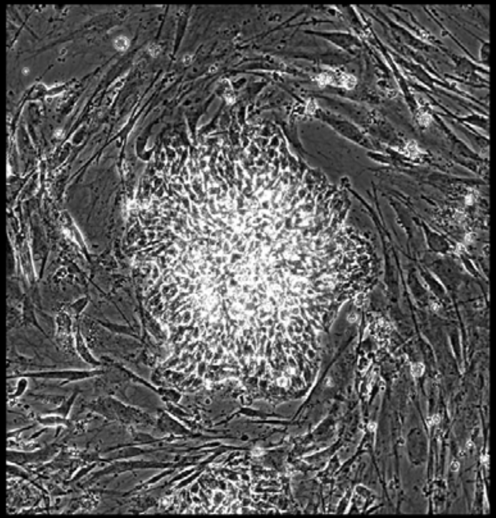

Well = 1, Colony = 72
Rater 1 = 2
Rater 2 = 3

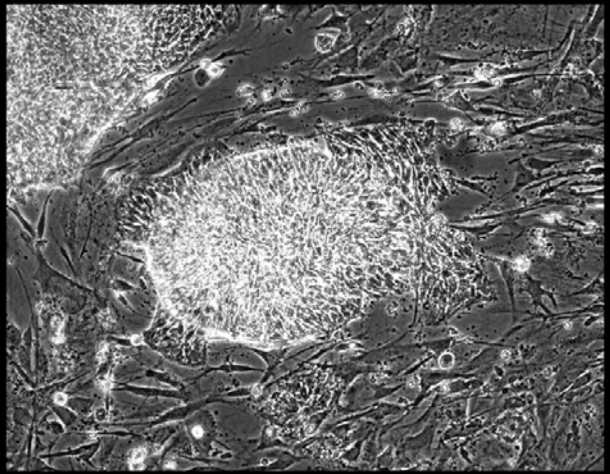

Well = 1, Colony = 73
Rater 1 = 4
Rater 2 = 5

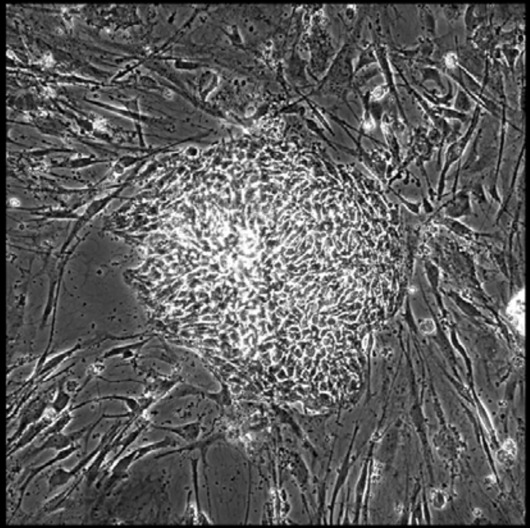

Well = 1, Colony = 74
Rater 1 = 4
Rater 2 = 4

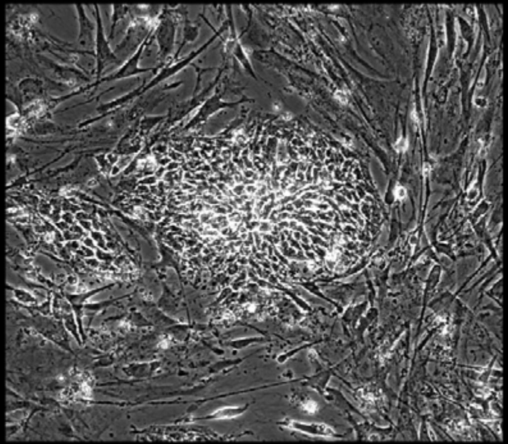

Well = 1, Colony = 75
Rater 1 = 4
Rater 2 = 4

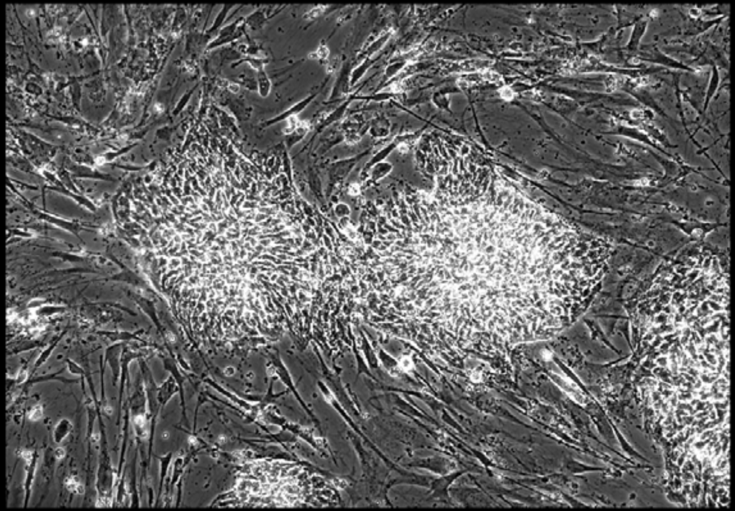

Well = 1, Colony = 76
Rater 1 = 1
Rater 2 = 1

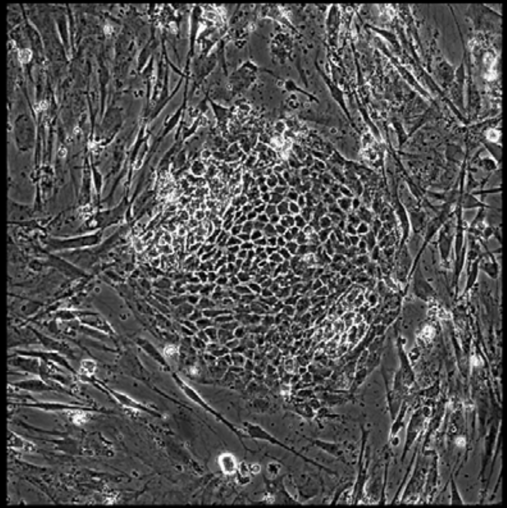

Well = 1, Colony = 77
Rater 1 = 3
Rater 2 = 3

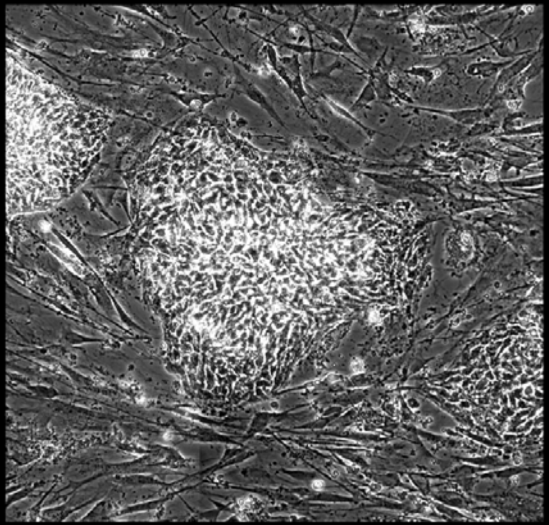

Well = 1, Colony = 78
Rater 1 = 4
Rater 2 = 4

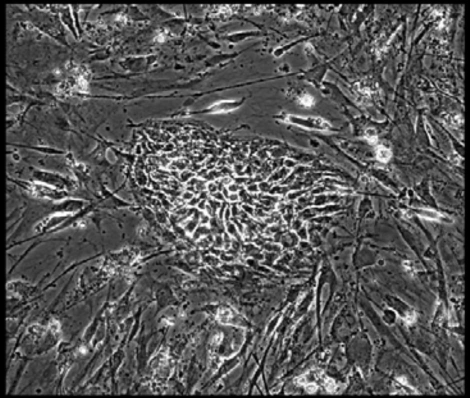

Well = 1, Colony = 79
Rater 1 = 4
Rater 2 = not scored

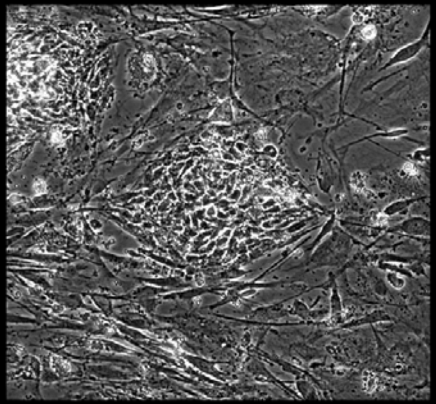

Well = 1, Colony = 80
Rater 1 = 4
Rater 2 = 4

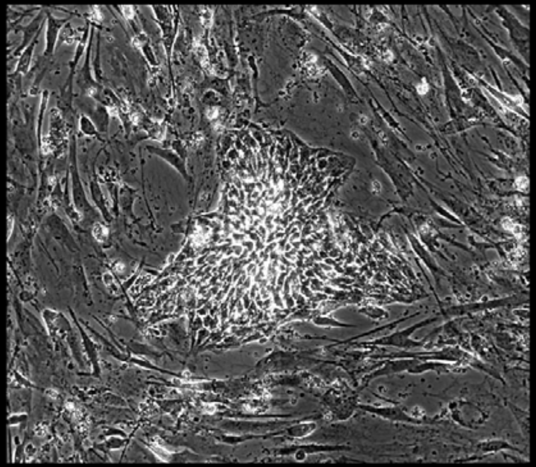

Well = 1, Colony = 81
Rater 1 = 4
Rater 2 = 5

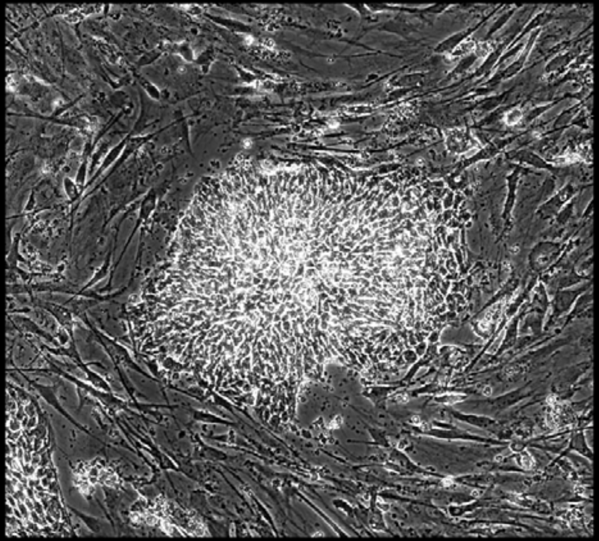

Well = 1, Colony = 82
Rater 1 = 4
Rater 2 = 4

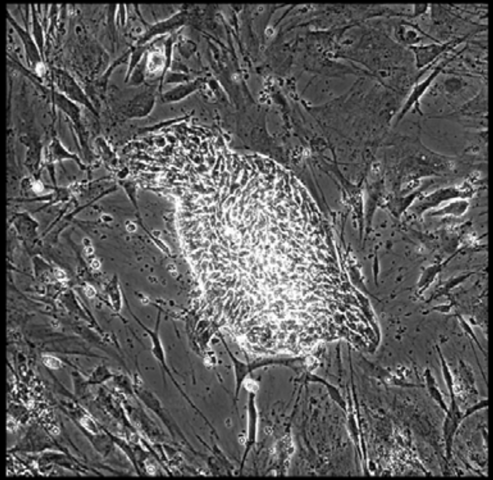

Well = 1, Colony = 83
Rater 1 = 4
Rater 2 = not scored

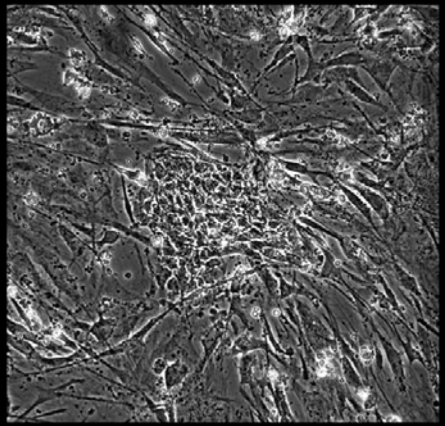

Well = 1, Colony = 84
Rater 1 = 3
Rater 2 = not scored

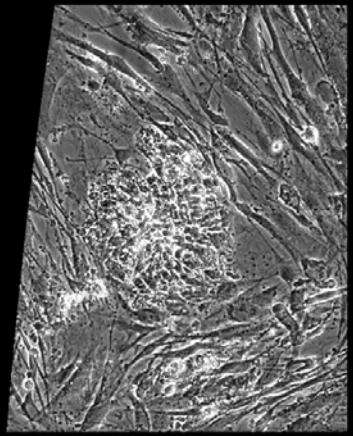

Well = 1, Colony = 85
Rater 1 = 4
Rater 2 = 4

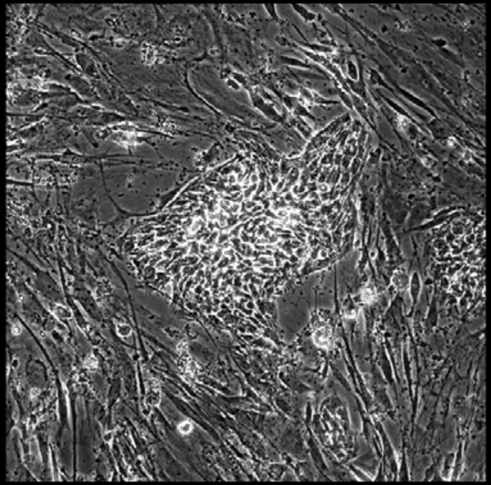

Well = 1, Colony = 86
Rater 1 = 4
Rater 2 = 3

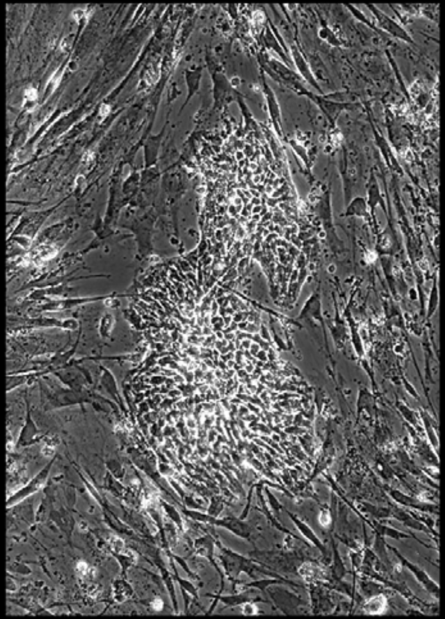

Well = 1, Colony = 87
Rater 1 = 4
Rater 2 = 4

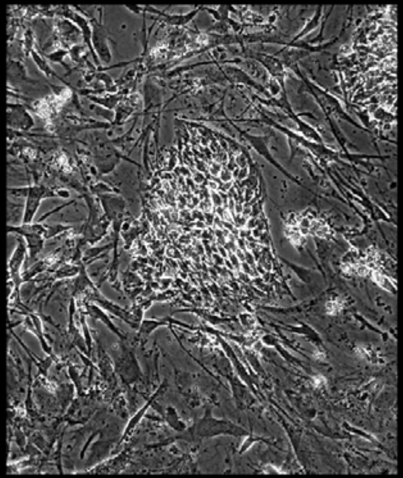

Well = 1, Colony = 88
Rater 1 = 3
Rater 2 = 3

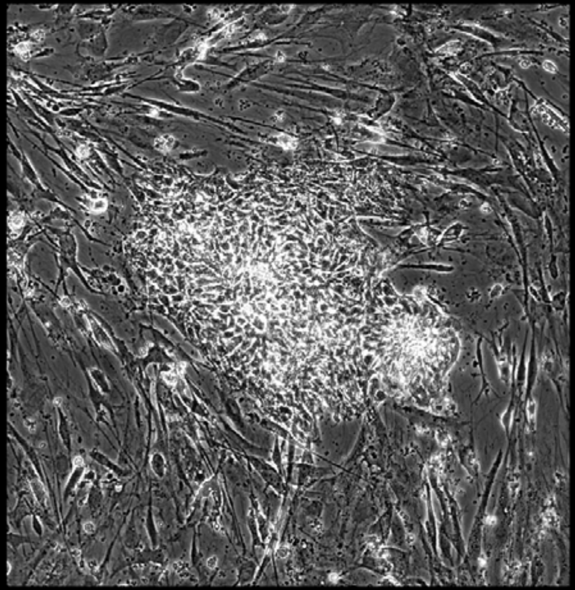

Well = 1, Colony = 89
Rater 1 = 4
Rater 2 = 4

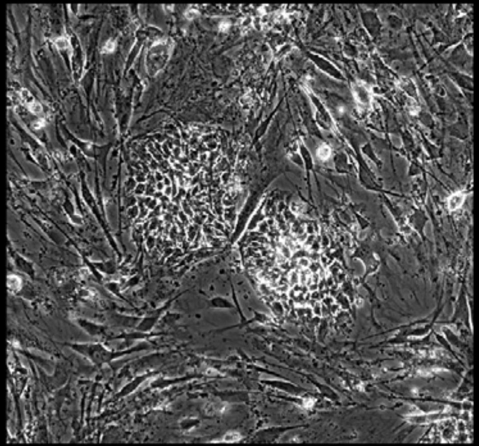

Well = 1, Colony = 90
Rater 1 = 4
Rater 2 = 4

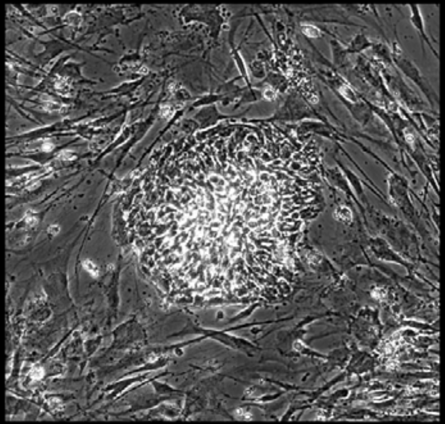

Well = 1, Colony = 91
Rater 1 = 5
Rater 2 = 4

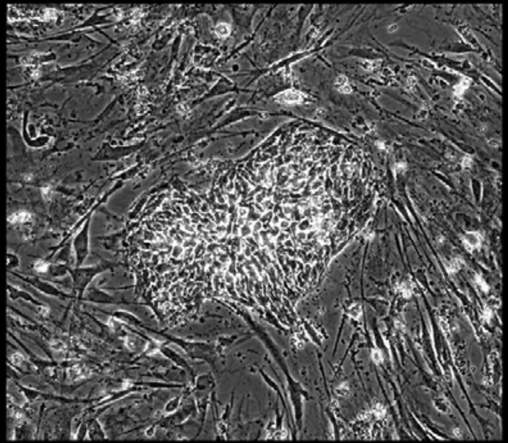

Well = 1, Colony = 92
Rater 1 = 4
Rater 2 = 3

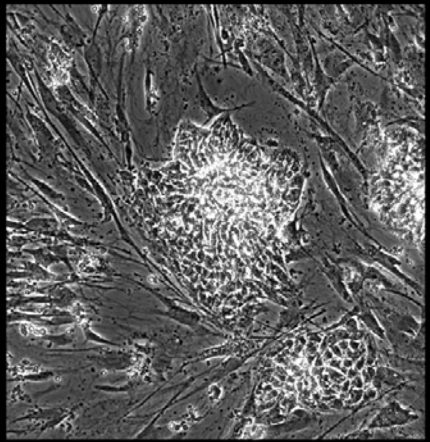

Well = 1, Colony = 93
Rater 1 = 2
Rater 2 = 3

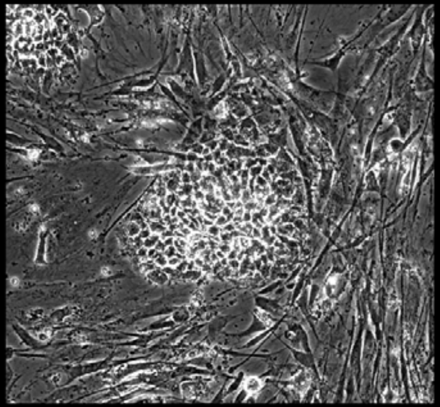

Well = 1, Colony = 94
Rater 1 = 5
Rater 2 = 4

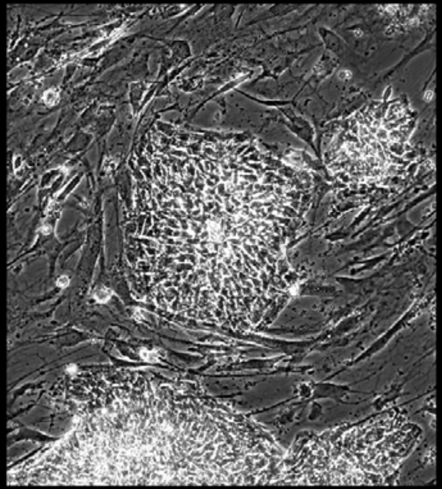

Well = 1, Colony = 95
Rater 1 = 4
Rater 2 = 4

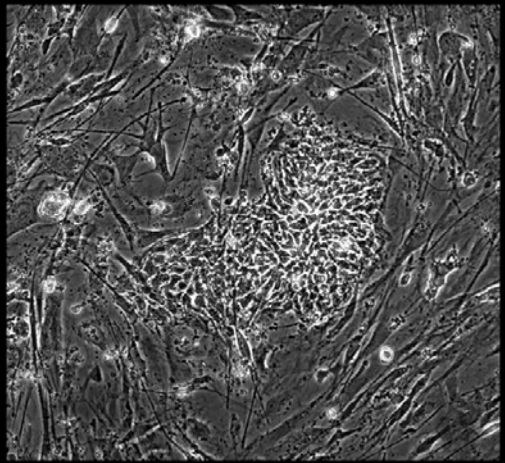

Well = 1, Colony = 96
Rater 1 = 5
Rater 2 = 5

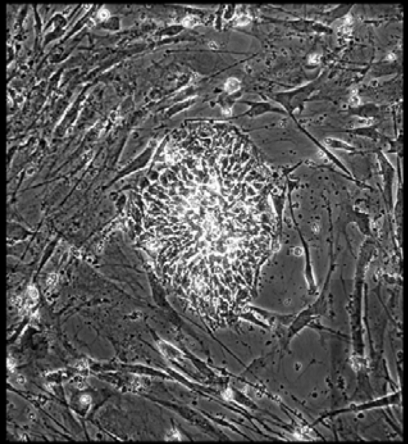

Well = 1, Colony = 97
Rater 1 = 4
Rater 2 = 4

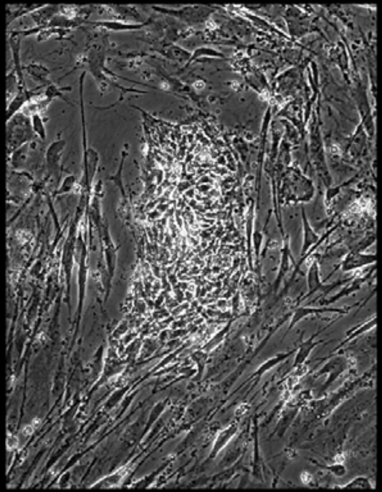

Well = 1, Colony = 98
Rater 1 = 2
Rater 2 = 2

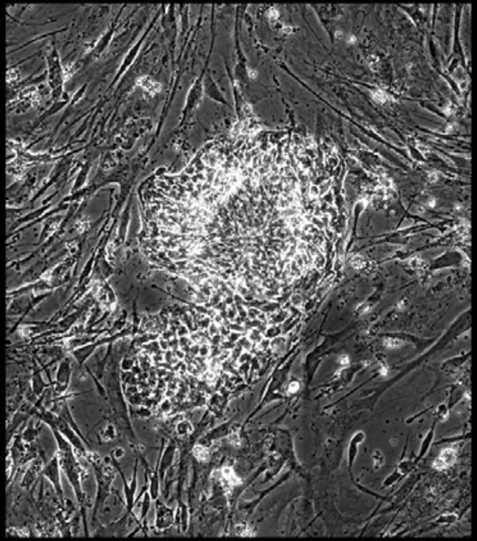

Well = 1, Colony = 99
Rater 1 = 2
Rater 2 = 2

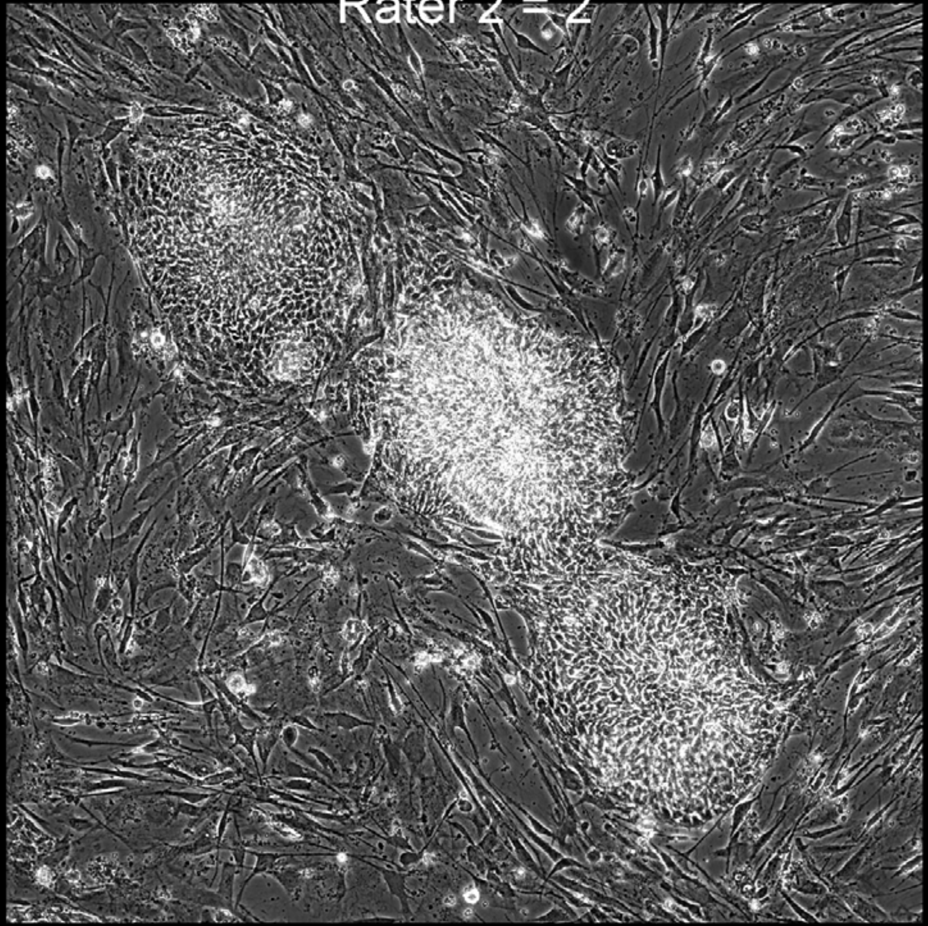

Well = 1, Colony = 100
Rater 1 = 3
Rater 2 = 4

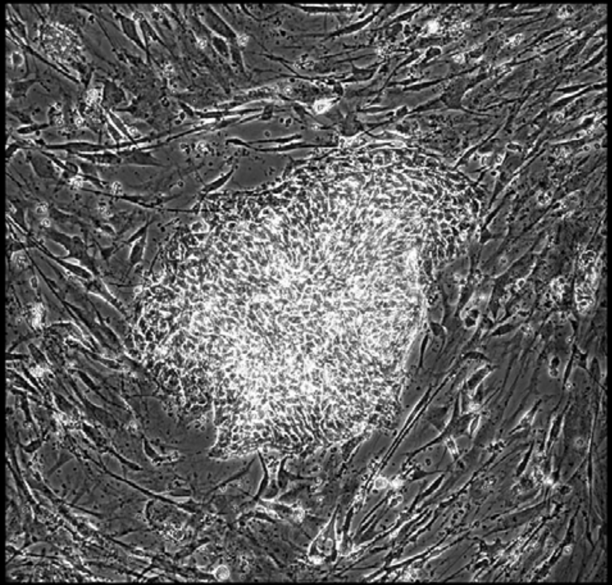

Well = 1, Colony = 101
Rater 1 = 2
Rater 2 = 4

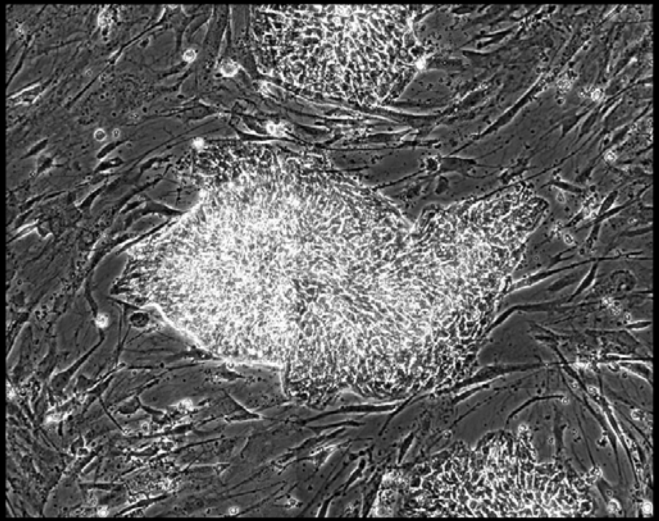

Well = 1, Colony = 102
Rater 1 = 3
Rater 2 = 4

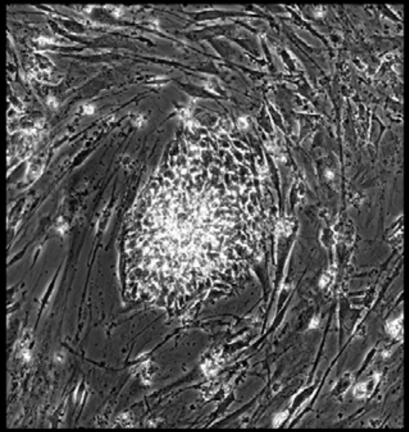

Well = 1, Colony = 103
Rater 1 = 3
Rater 2 = 4

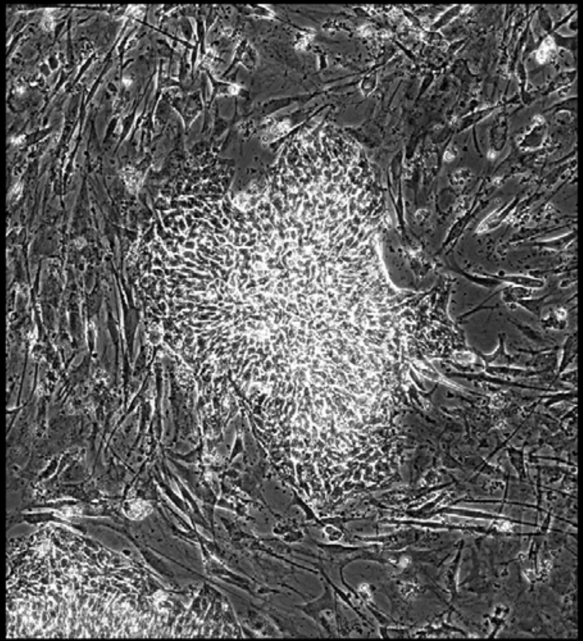

Well = 1, Colony = 104
Rater 1 = 4
Rater 2 = 4

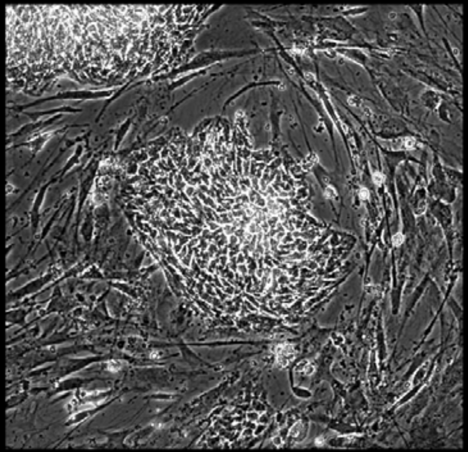

Well = 1, Colony = 105
Rater 1 = 4
Rater 2 = 5

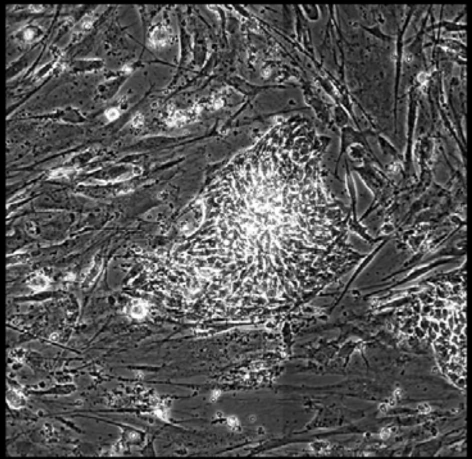

Well = 1, Colony = 106
Rater 1 = 3
Rater 2 = 4

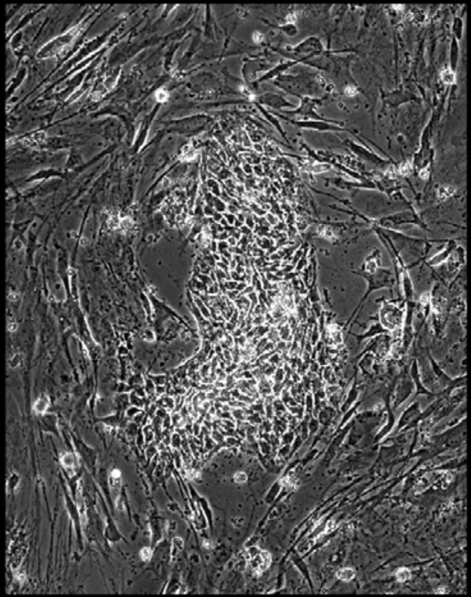

Well = 1, Colony = 107
Rater 1 = 4
Rater 2 = 4

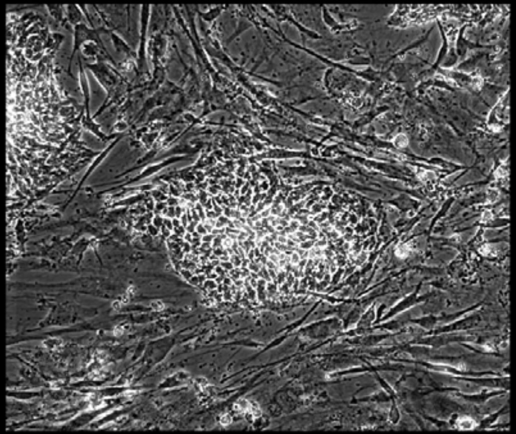

Well = 1, Colony = 108
Rater 1 = 2
Rater 2 = 2

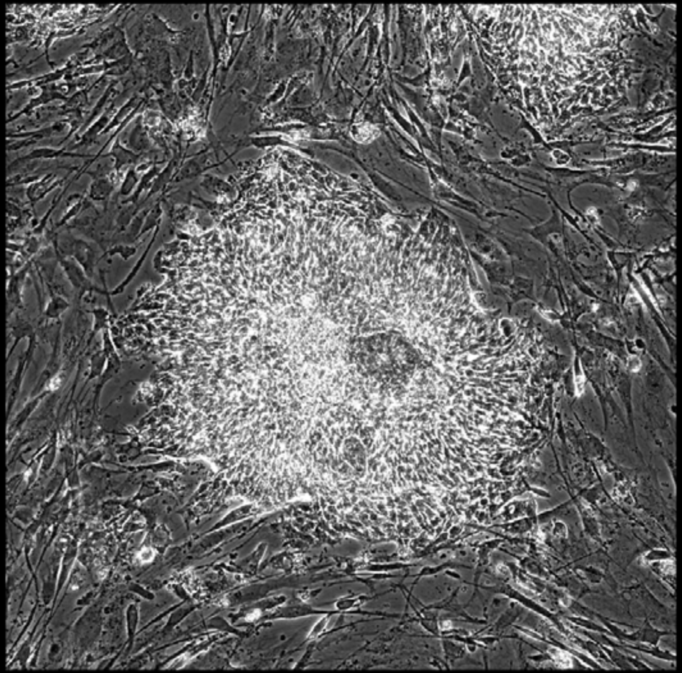

Well = 1, Colony = 109
Rater 1 = 4
Rater 2 = 5

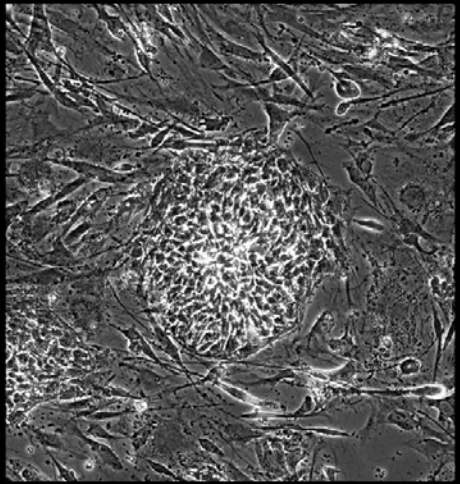

Well = 1, Colony = 110
Rater 1 = 2
Rater 2 = 4

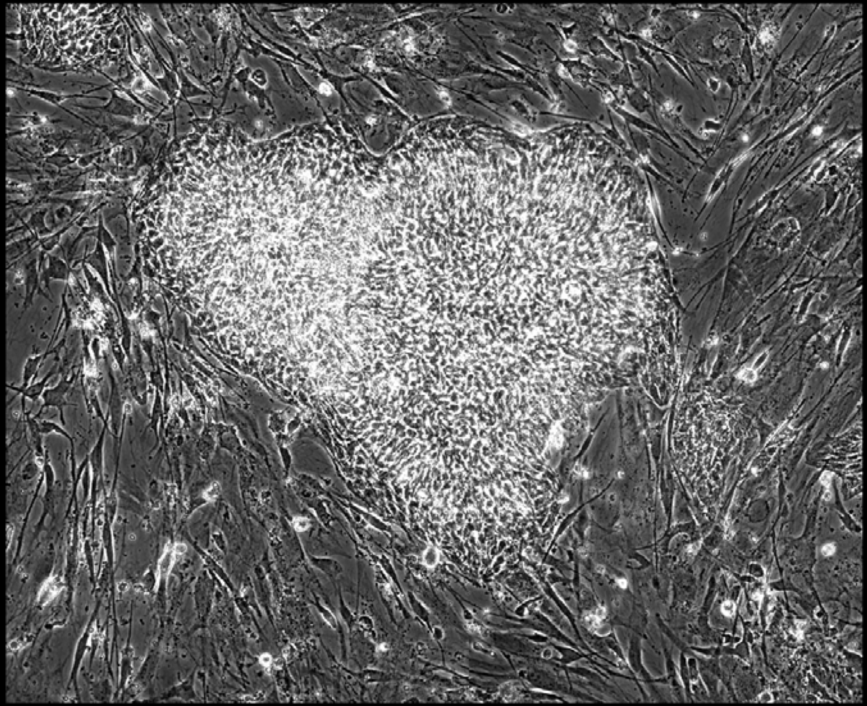

Well = 1, Colony = 111
Rater 1 = 4
Rater 2 = 5

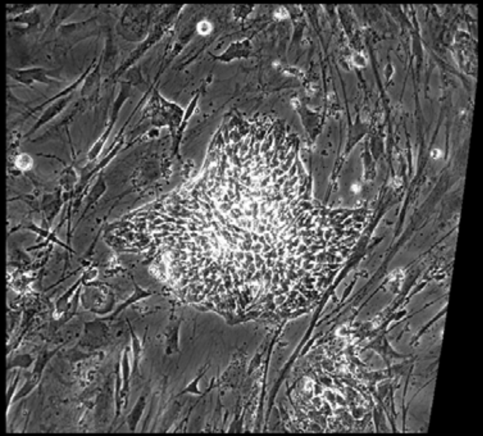

Well = 1, Colony = 112
Rater 1 = 2
Rater 2 = 2

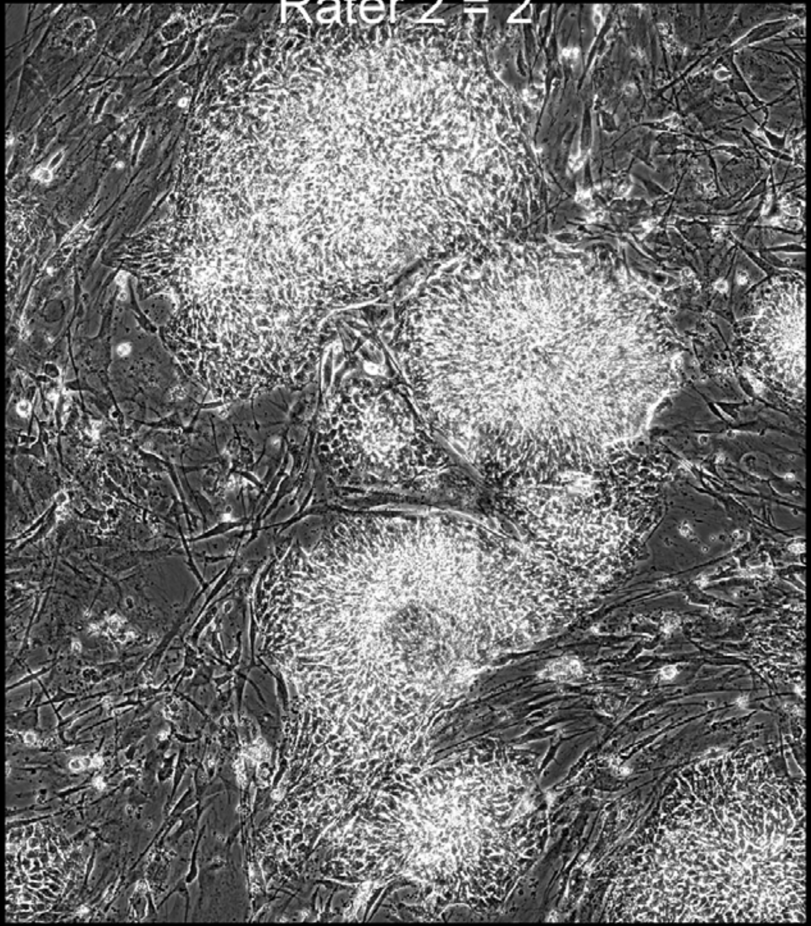

Well = 1, Colony = 113
Rater 1 = 2
Rater 2 = 5

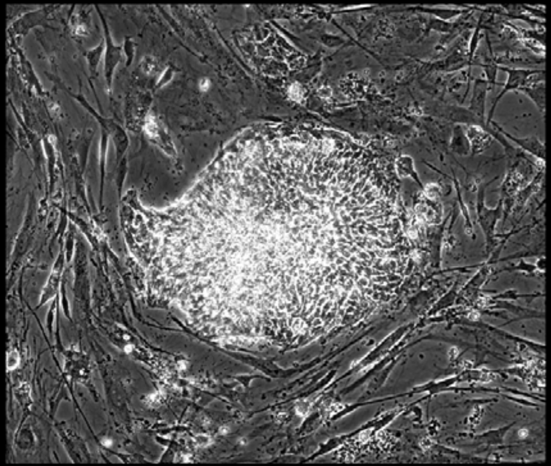

Well = 1, Colony = 114
Rater 1 = 5
Rater 2 = 5

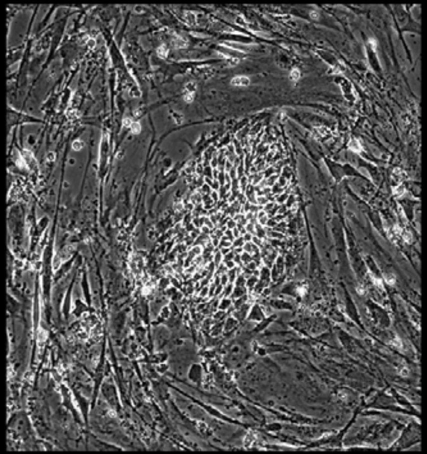

Well = 1, Colony = 115
Rater 1 = 2
Rater 2 = 2

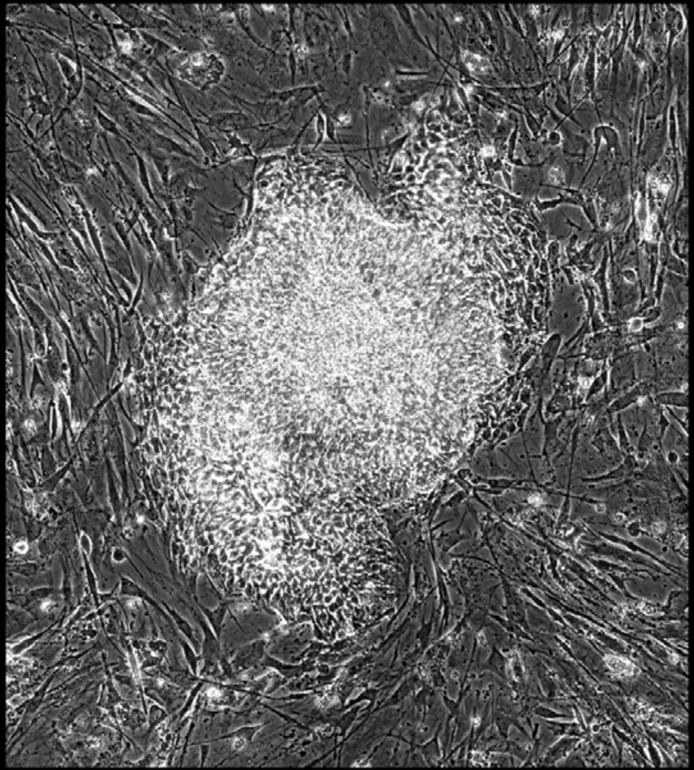

Well = 1, Colony = 116
Rater 1 = 2
Rater 2 = 2

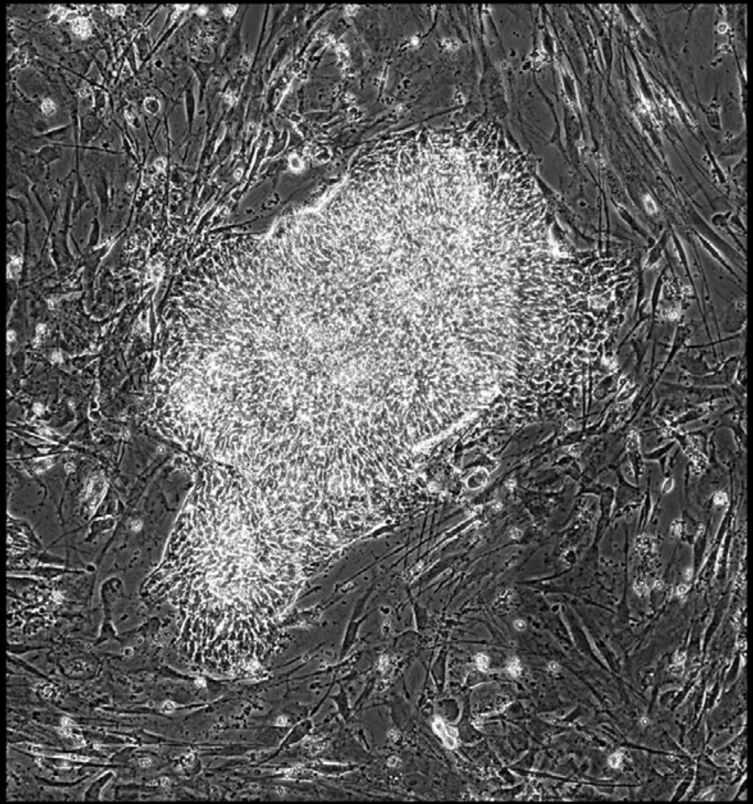

Well = 1, Colony = 117
Rater 1 = 3
Rater 2 = 3

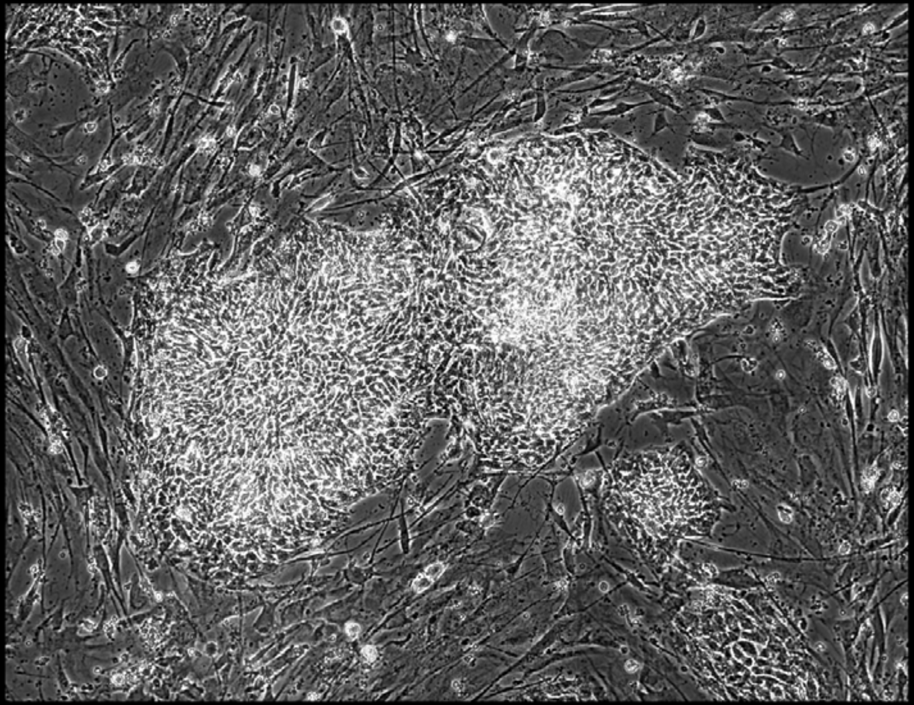

Well = 1, Colony = 118
Rater 1 = 2
Rater 2 = 3

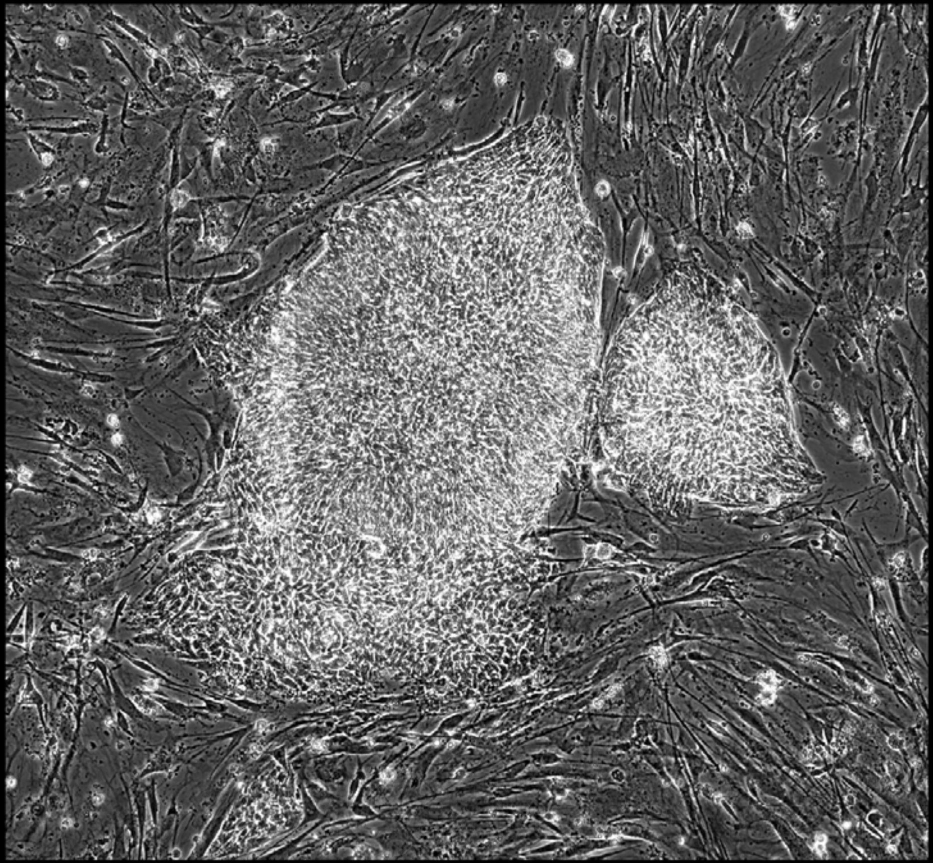

Well = 1, Colony = 119
Rater 1 = 4
Rater 2 = 4

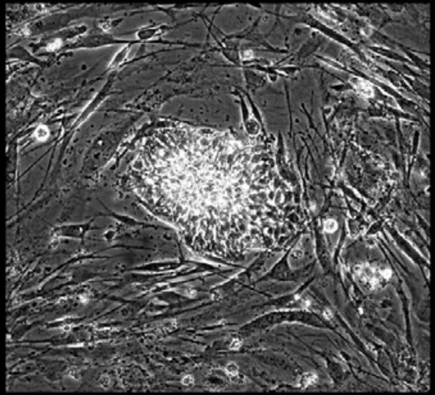

Well = 1, Colony = 120
Rater 1 = 4
Rater 2 = 4

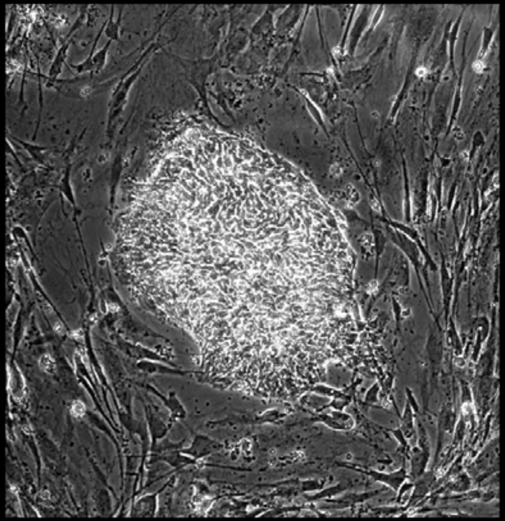

Well = 1, Colony = 121
Rater 1 = 5
Rater 2 = 4

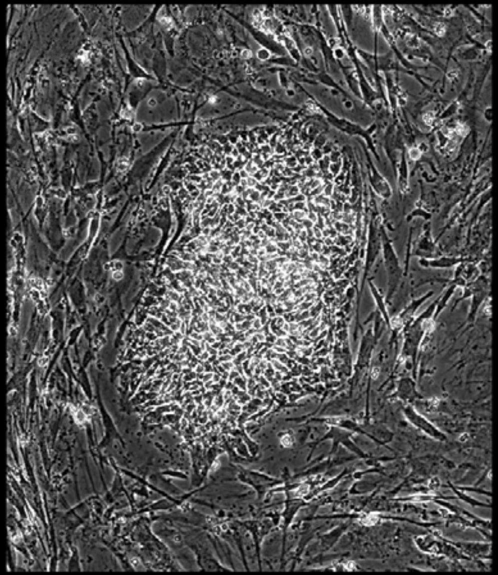

Well = 1, Colony = 122
Rater 1 = 3
Rater 2 = 3

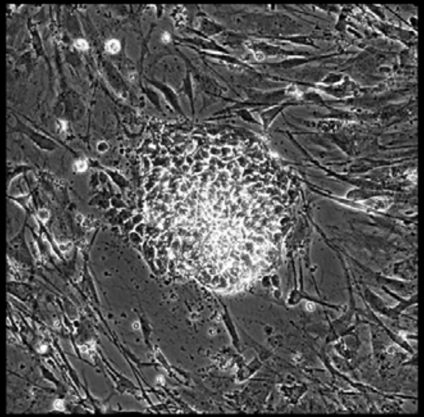

Well = 1, Colony = 123
Rater 1 = 4
Rater 2 = 3

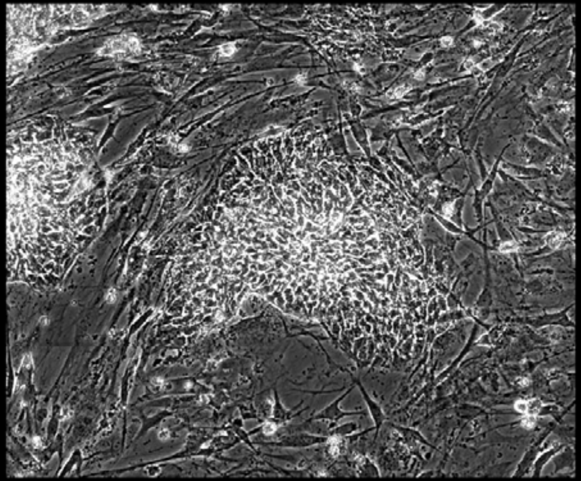

Well = 1, Colony = 124
Rater 1 = 4
Rater 2 = 4

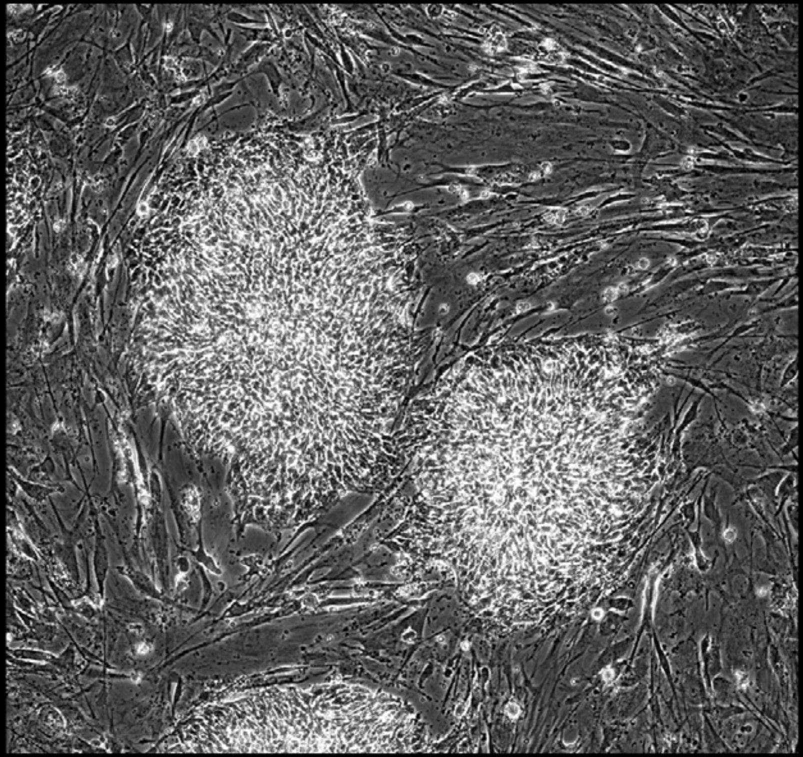

Well = 1, Colony = 125
Rater 1 = 4
Rater 2 = 4

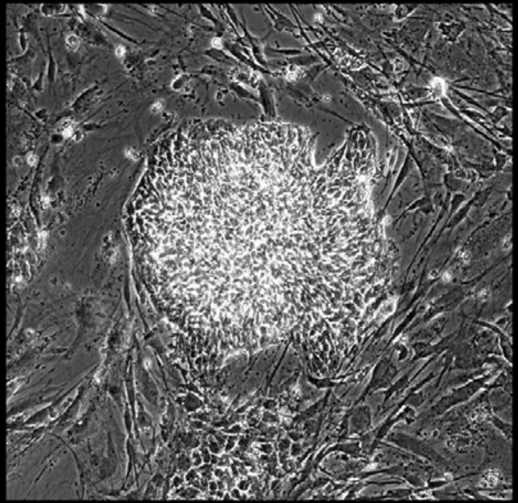

Well = 1, Colony = 126
Rater 1 = 2
Rater 2 = 2

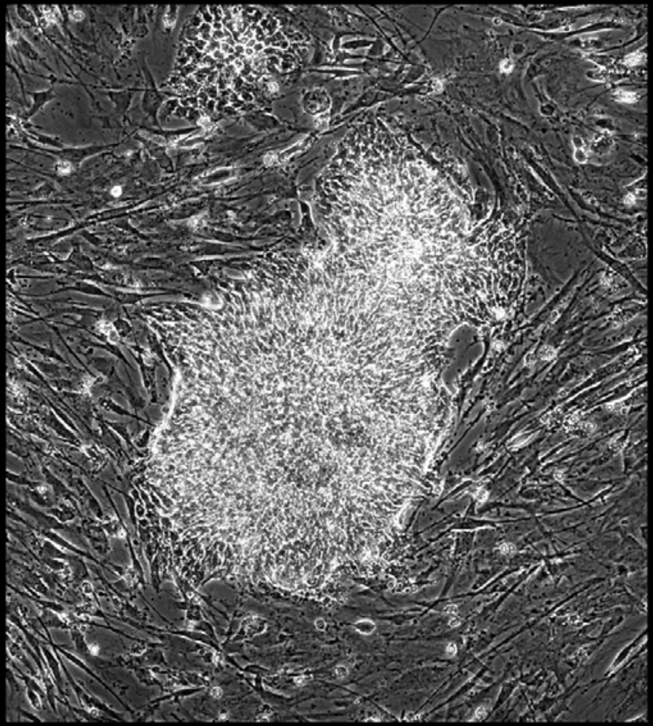

Well = 1, Colony = 127
Rater 1 = 3
Rater 2 = 4

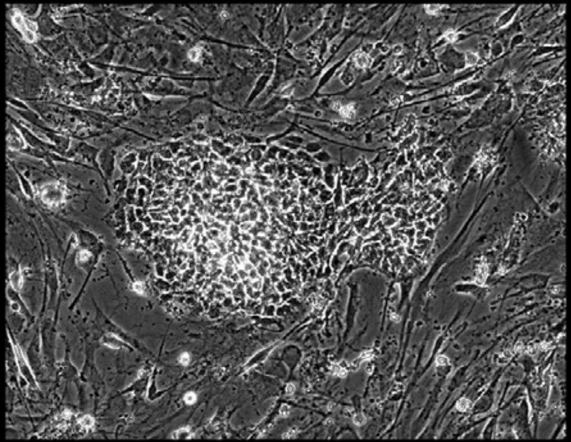

Well = 1, Colony = 128
Rater 1 = 2
Rater 2 = not scored

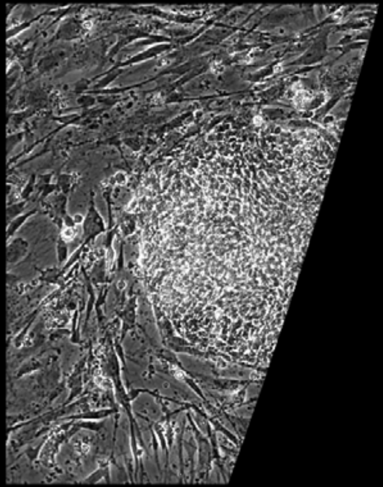

Well = 1, Colony = 129
Rater 1 = 3
Rater 2 = 4

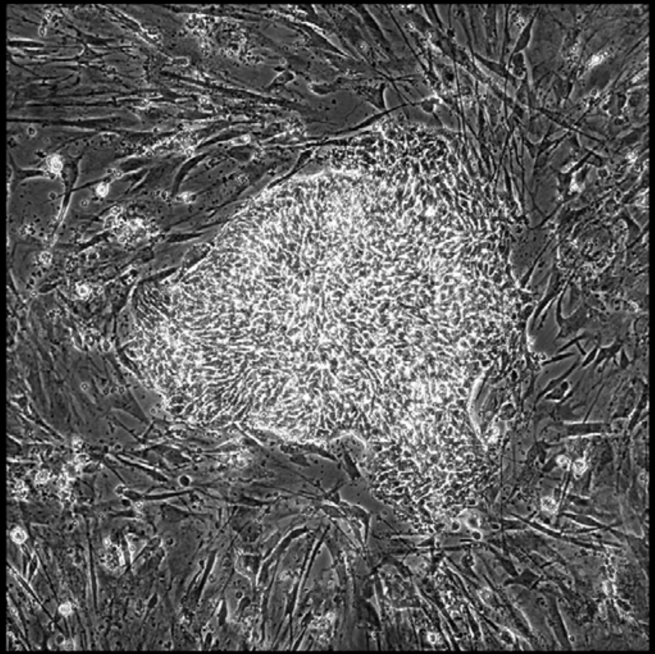

Well = 1, Colony = 130
Rater 1 = 1
Rater 2 = 2

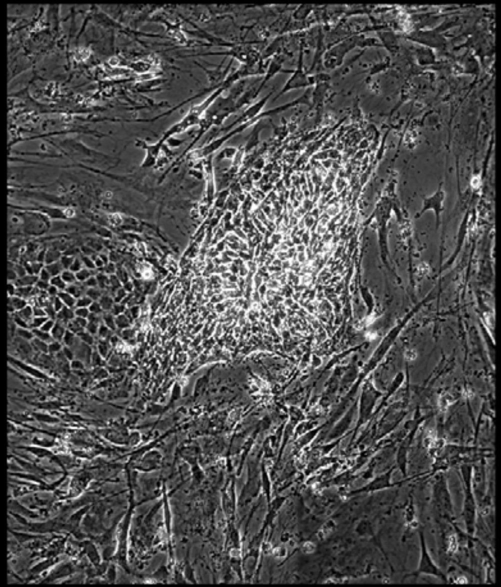

Well = 1, Colony = 131
Rater 1 = 1
Rater 2 = not scored

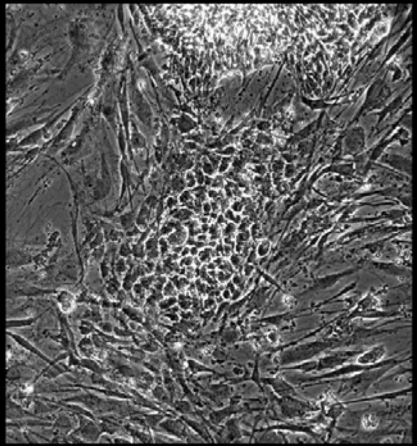

Well = 1, Colony = 132
Rater 1 = 2
Rater 2 = 2

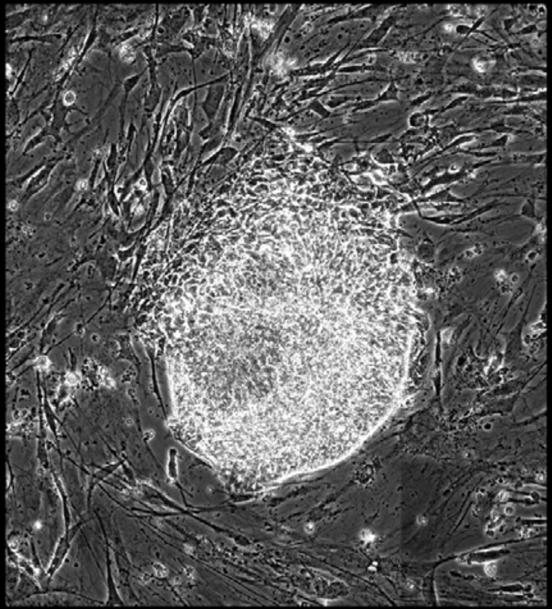

Well = 1, Colony = 133
Rater 1 = 2
Rater 2 = 2

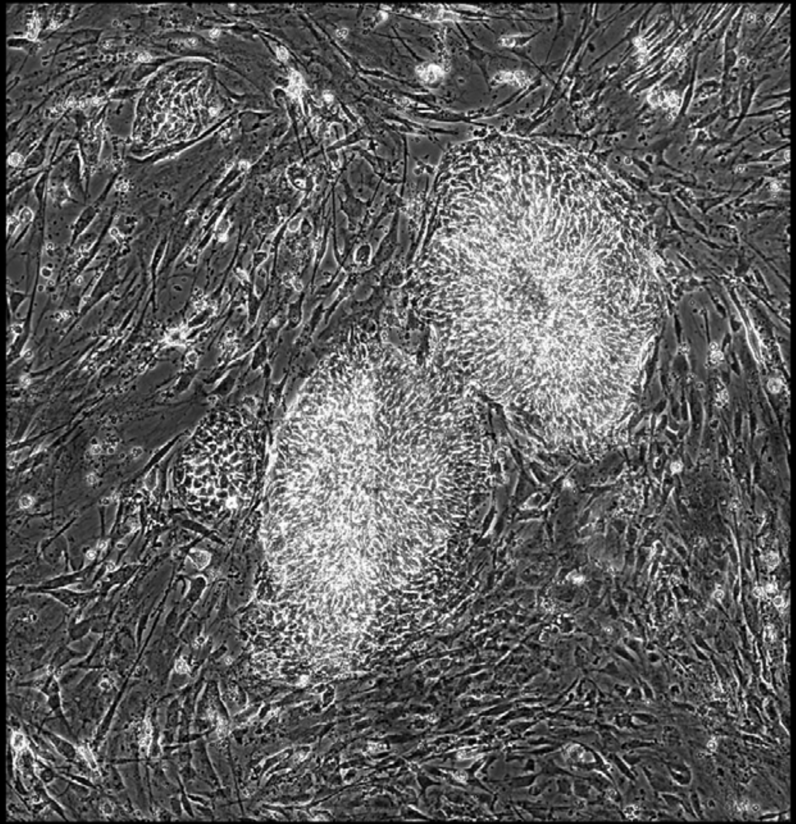

Well = 1, Colony = 134
Rater 1 = 5
Rater 2 = 5

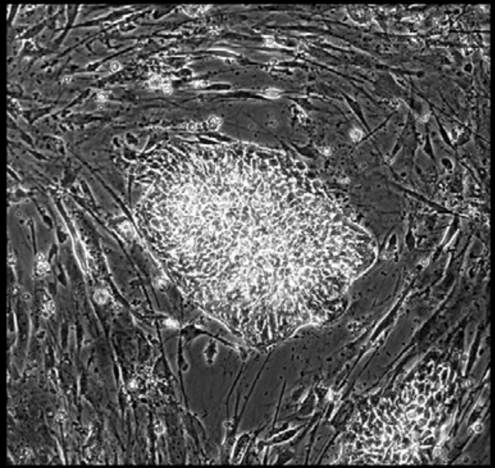

Well = 1, Colony = 135
Rater 1 = 4
Rater 2 = 4

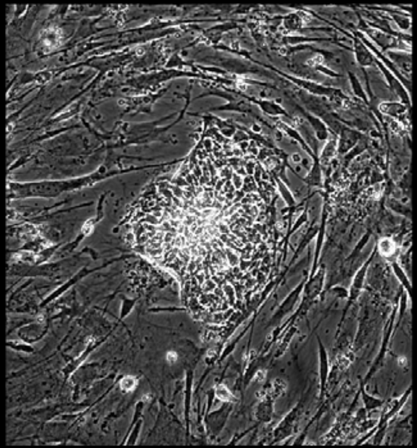

Well = 1, Colony = 136
Rater 1 = 5
Rater 2 = 4

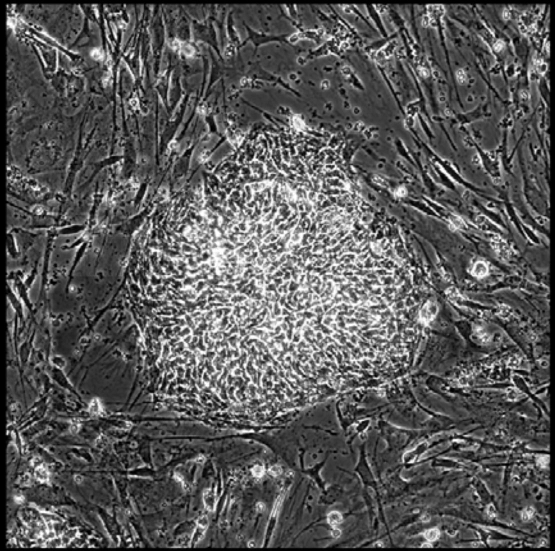

Well = 1, Colony = 137
Rater 1 = 4
Rater 2 = 4

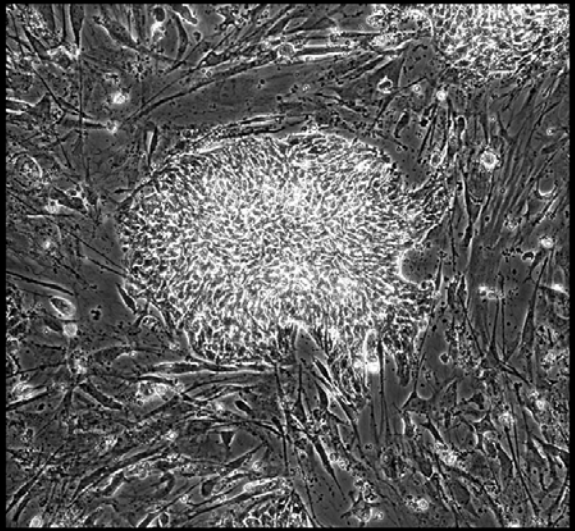

Well = 1, Colony = 138
Rater 1 = 4
Rater 2 = 4

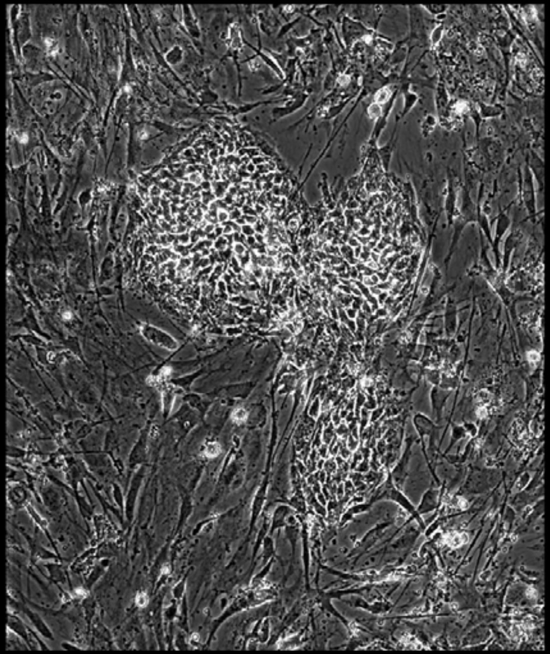

Well = 1, Colony = 139
Rater 1 = 4
Rater 2 = 4

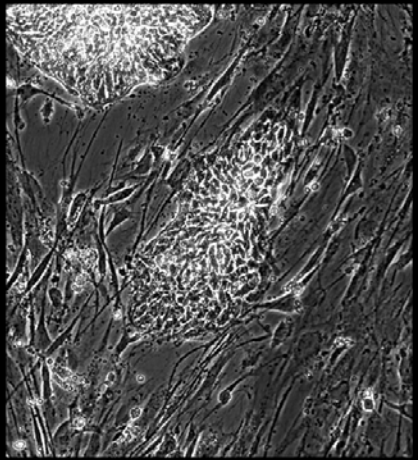

Well = 1, Colony = 140
Rater 1 = 4
Rater 2 = 4

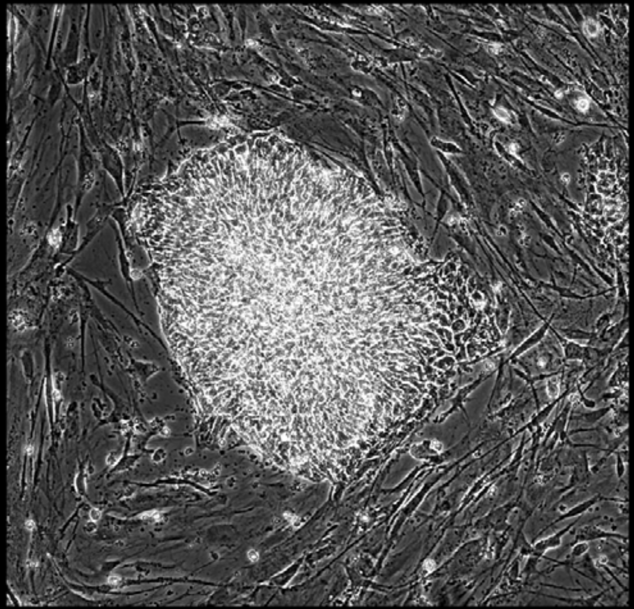

Well = 1, Colony = 141
Rater 1 = 4
Rater 2 = 4

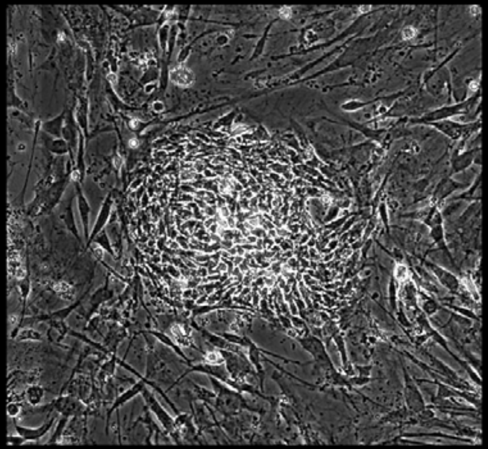

Well = 1, Colony = 142
Rater 1 = 4
Rater 2 = 4

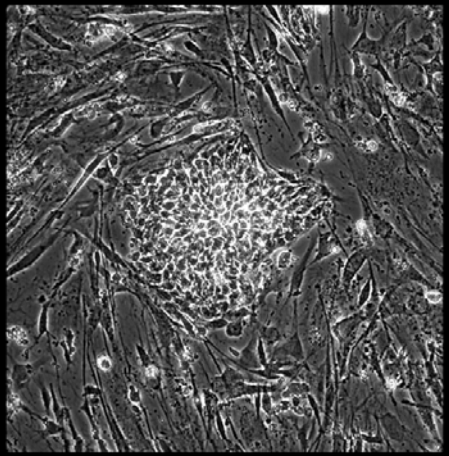

Well = 1, Colony = 143
Rater 1 = 5
Rater 2 = 4

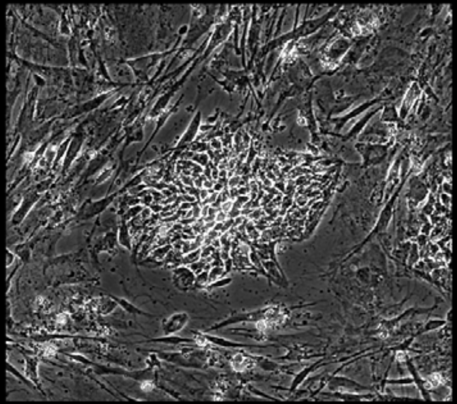

Well = 1, Colony = 144
Rater 1 = 2
Rater 2 = 2

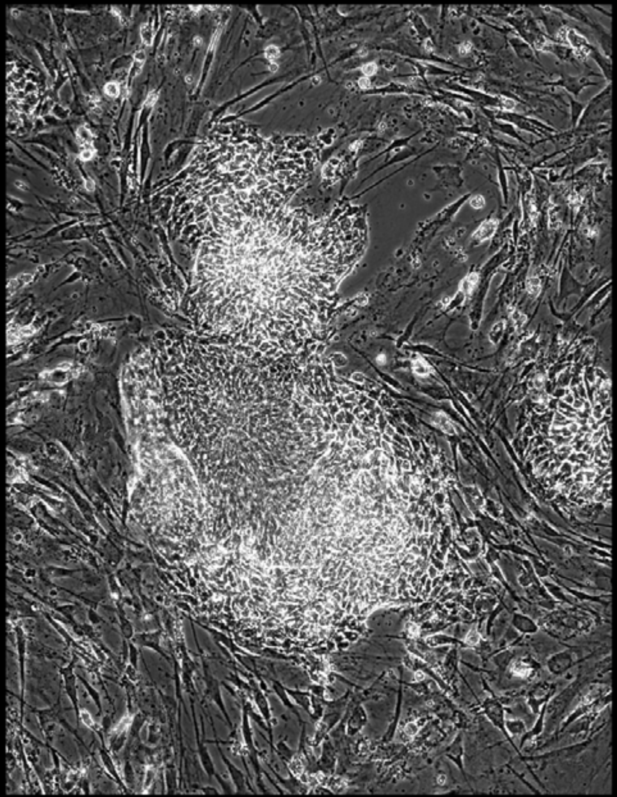

Well = 1, Colony = 145
Rater 1 = 3
Rater 2 = 4

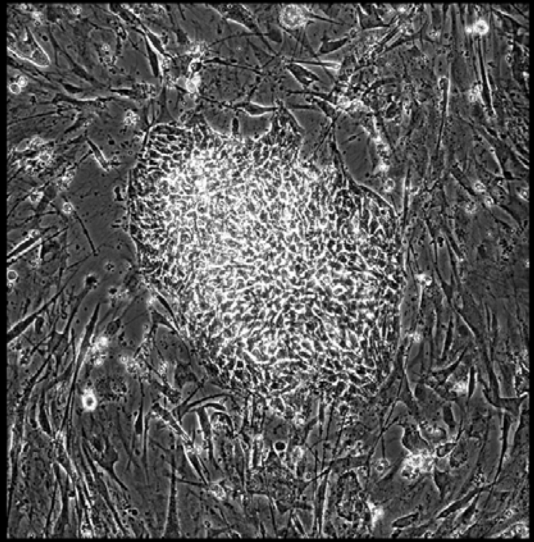

Well = 1, Colony = 146
Rater 1 = 2
Rater 2 = 3

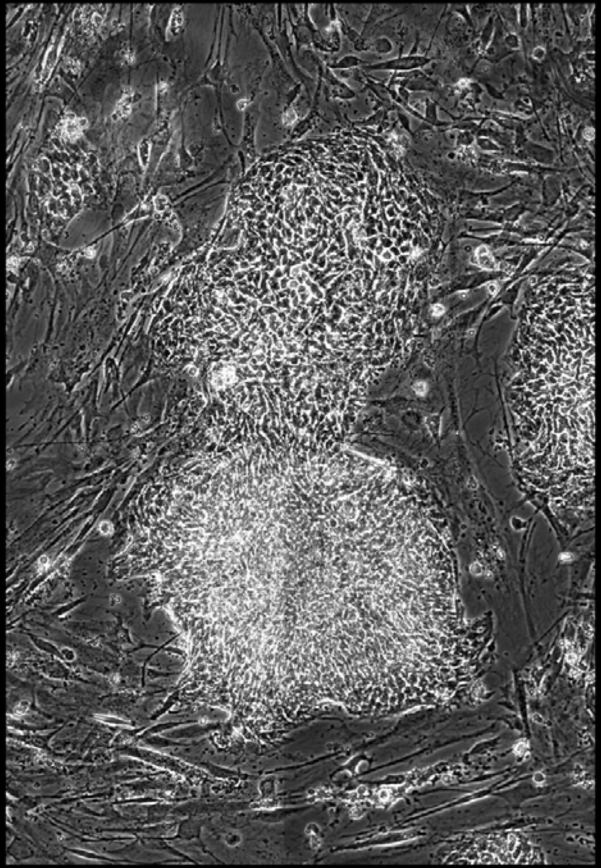

Well = 1, Colony = 147
Rater 1 = 3
Rater 2 = 4

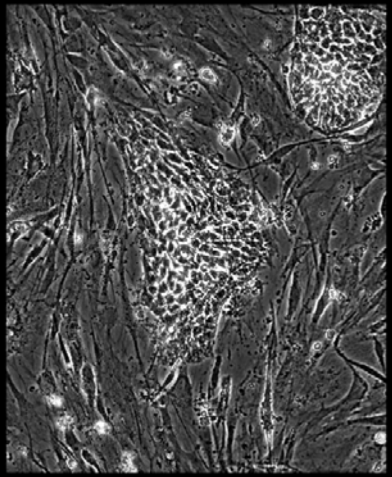

Well = 1, Colony = 148
Rater 1 = 3
Rater 2 = 4

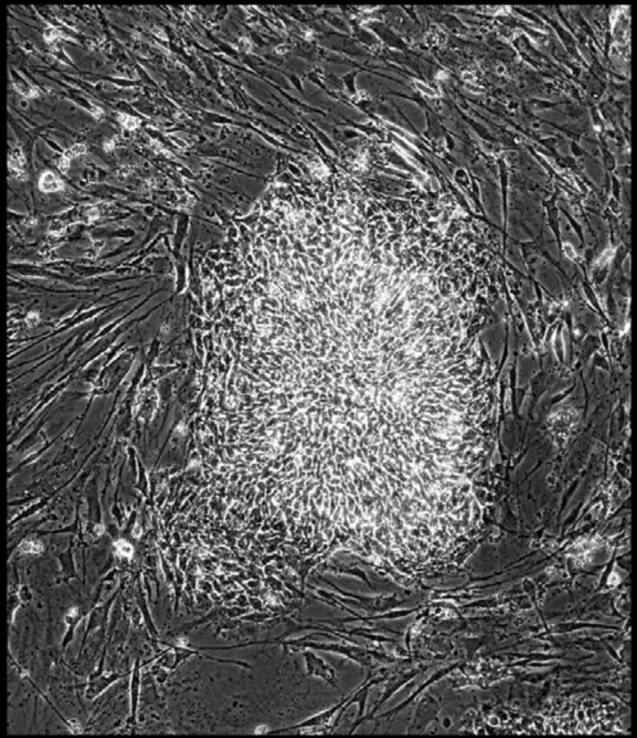

Well = 1, Colony = 149
Rater 1 = 4
Rater 2 = 5

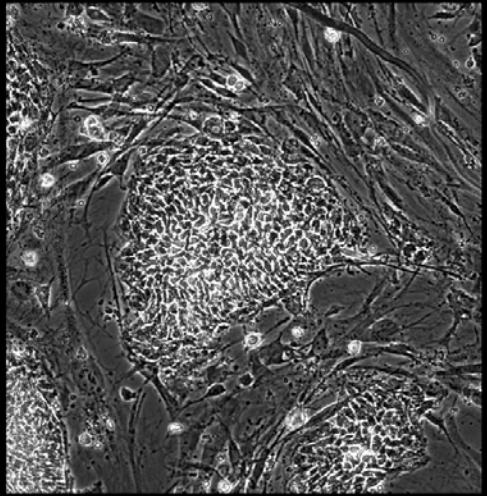

Well = 1, Colony = 150
Rater 1 = 4
Rater 2 = 5

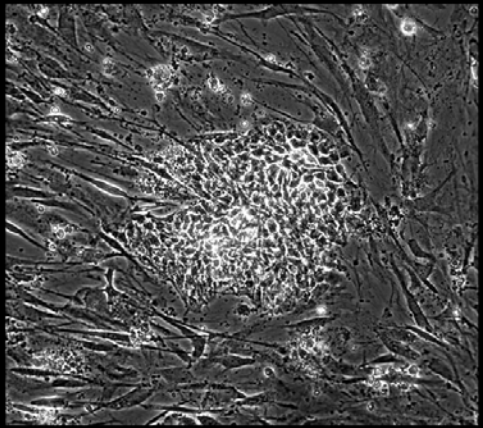

Well = 1, Colony = 151
Rater 1 = 3
Rater 2 = 4

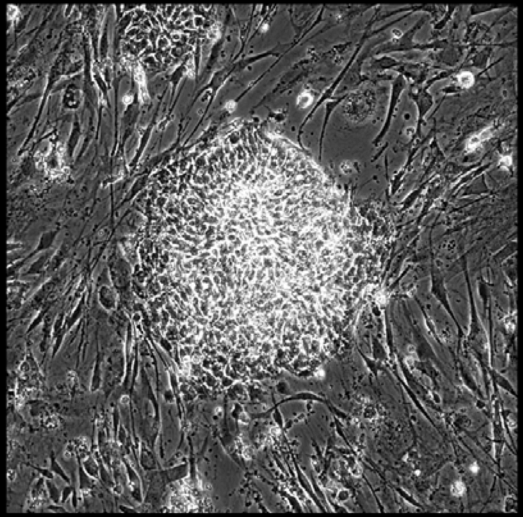

Well = 1, Colony = 152
Rater 1 = 3
Rater 2 = 3

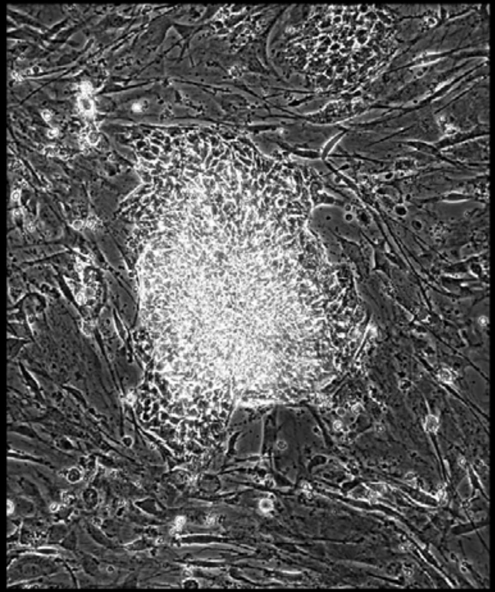

Well = 1, Colony = 153
Rater 1 = 4
Rater 2 = 4

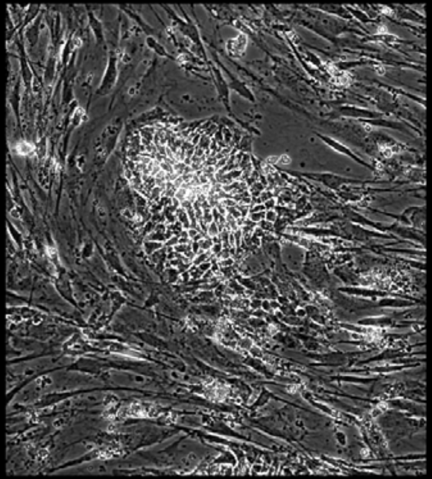

Well = 1, Colony = 154
Rater 1 = 4
Rater 2 = 4

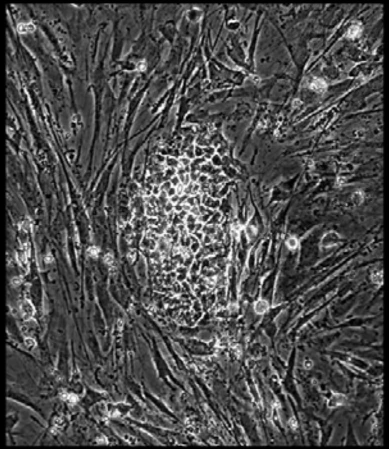

Well = 1, Colony = 155
Rater 1 = 4
Rater 2 = 4

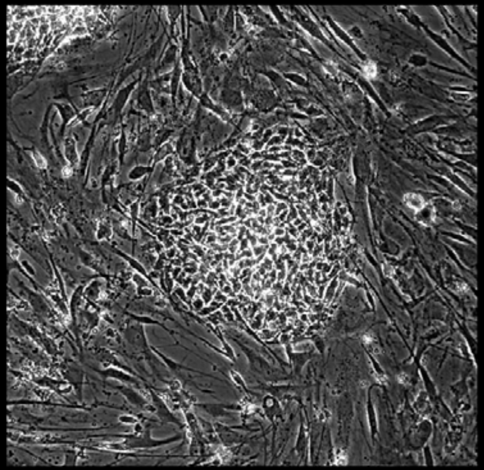

Well = 1, Colony = 156
Rater 1 = 4
Rater 2 = 3

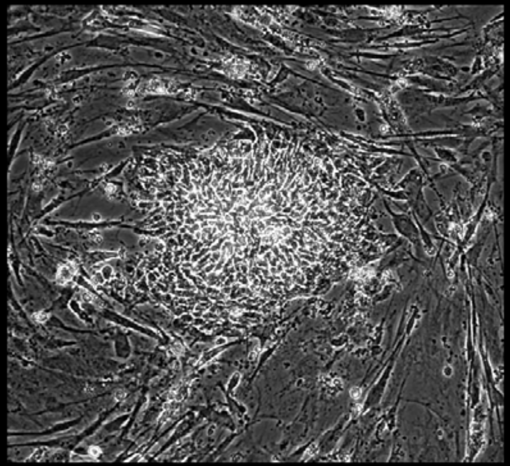

Well = 1, Colony = 157
Rater 1 = 4
Rater 2 = 4

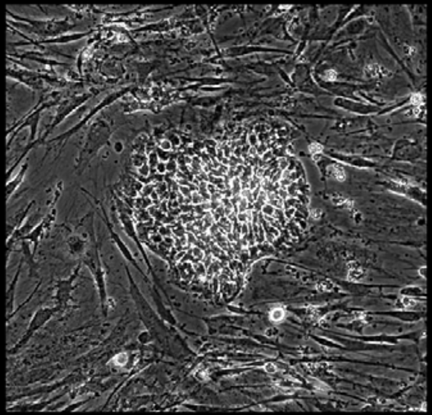

Well = 1, Colony = 158
Rater 1 = 4
Rater 2 = 4

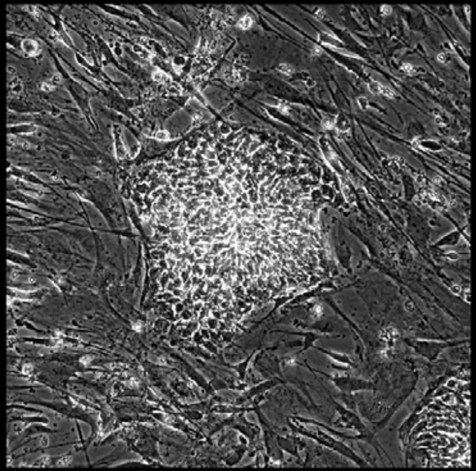

Well = 1, Colony = 159
Rater 1 = 4
Rater 2 = 3

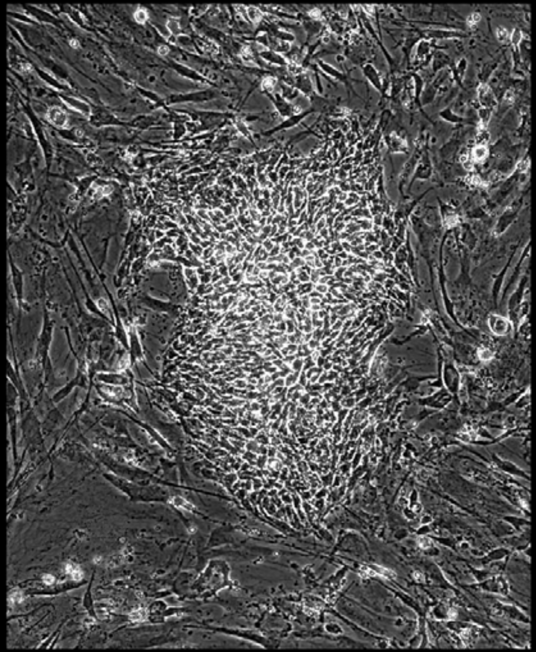

Well = 1, Colony = 160
Rater 1 = 4
Rater 2 = 3

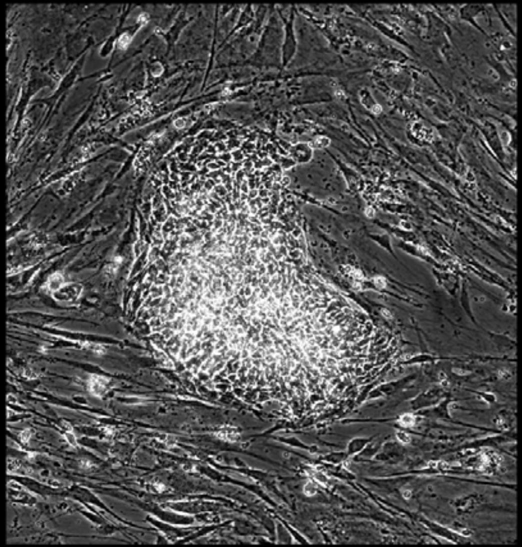

Well = 1, Colony = 161
Rater 1 = 4
Rater 2 = 4

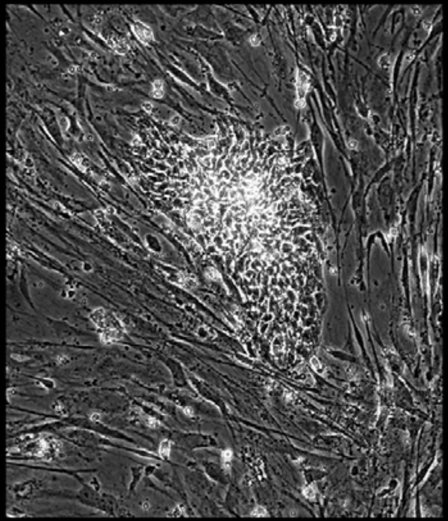

Well = 1, Colony = 162
Rater 1 = 3
Rater 2 = 3

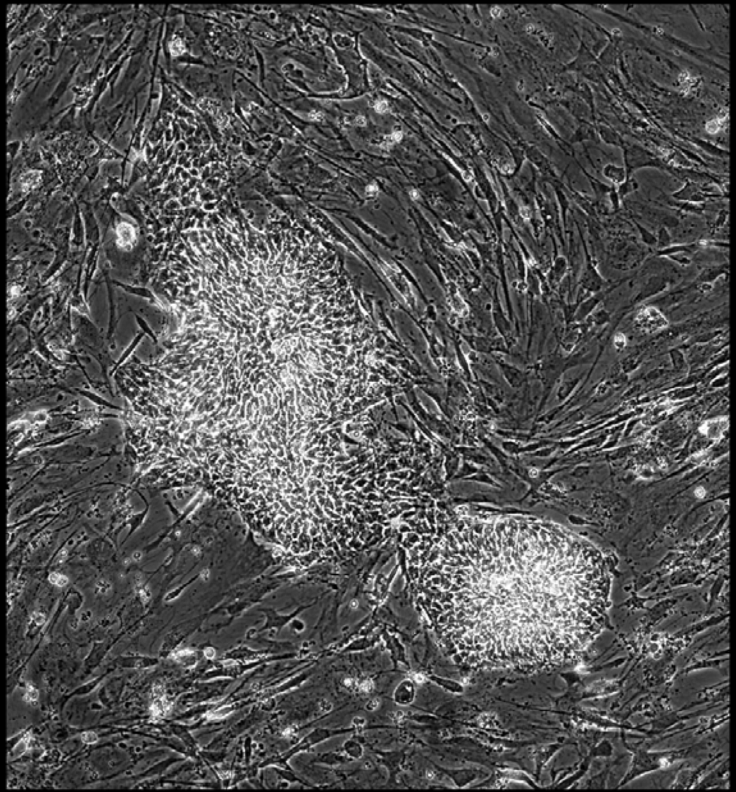

Well = 1, Colony = 163
Rater 1 = 4
Rater 2 = 5

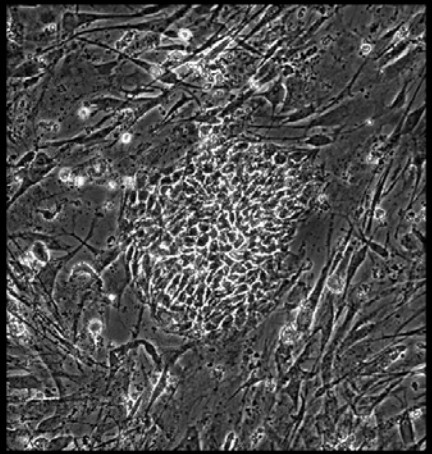

Well = 1, Colony = 164
Rater 1 = 5
Rater 2 = 5

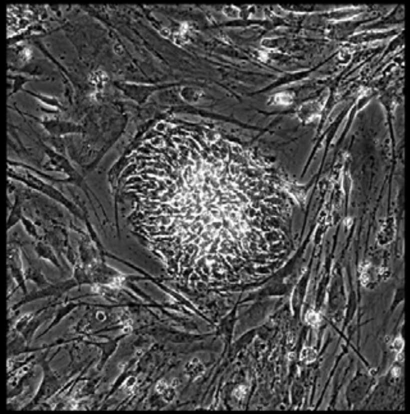

Well = 1, Colony = 165
Rater 1 = 3
Rater 2 = 3

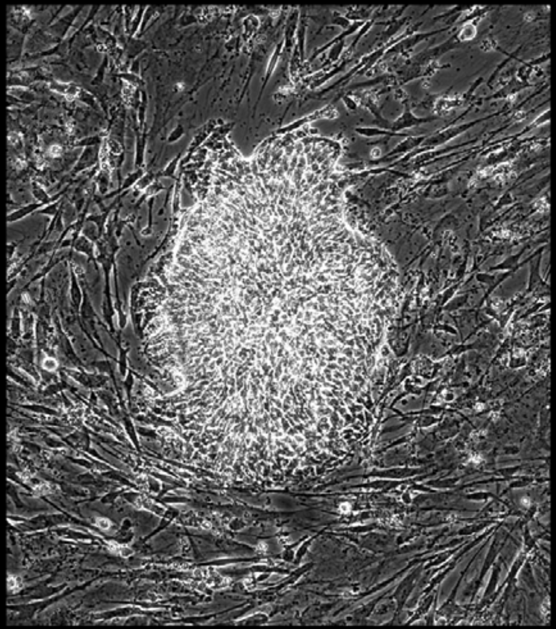

Well = 1, Colony = 166
Rater 1 = 4
Rater 2 = 4

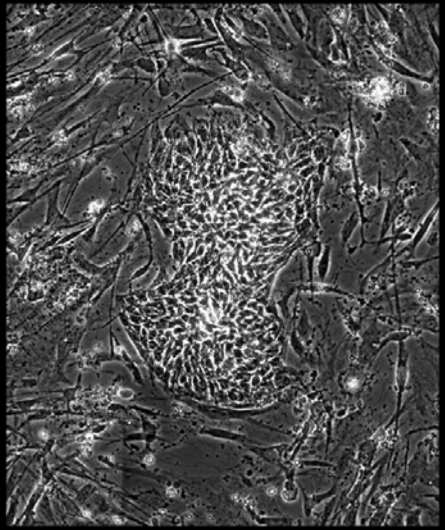

Well = 1, Colony = 167
Rater 1 = 4
Rater 2 = 3

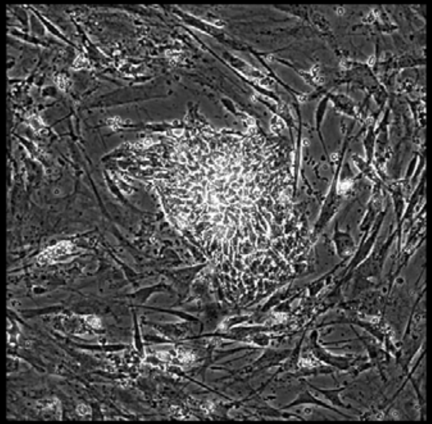

Well = 1, Colony = 168
Rater 1 = 2
Rater 2 = 2

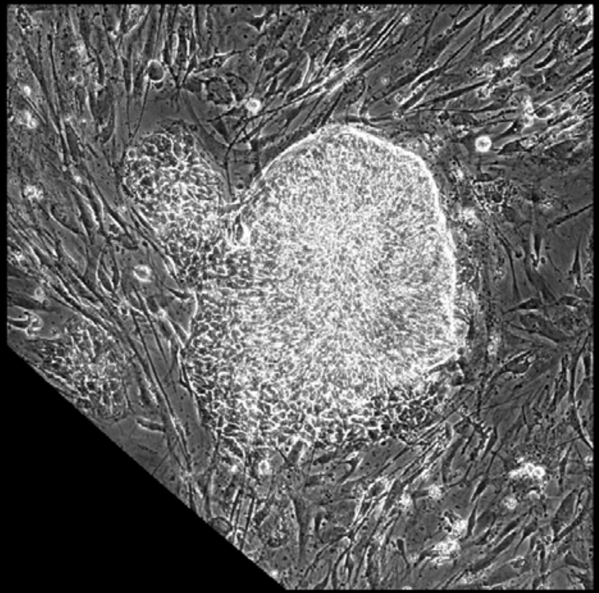

Well = 1, Colony = 169
Rater 1 = 2
Rater 2 = 1

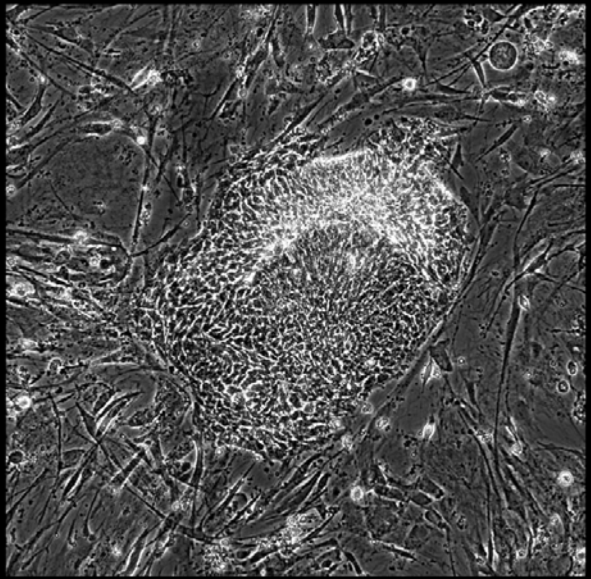

Well = 1, Colony = 170
Rater 1 = 4
Rater 2 = 4

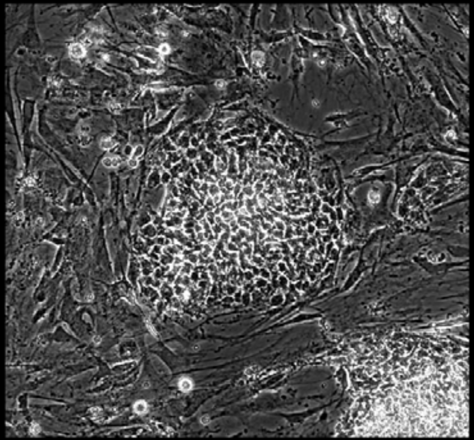

Well = 1, Colony = 171
Rater 1 = 3
Rater 2 = 3

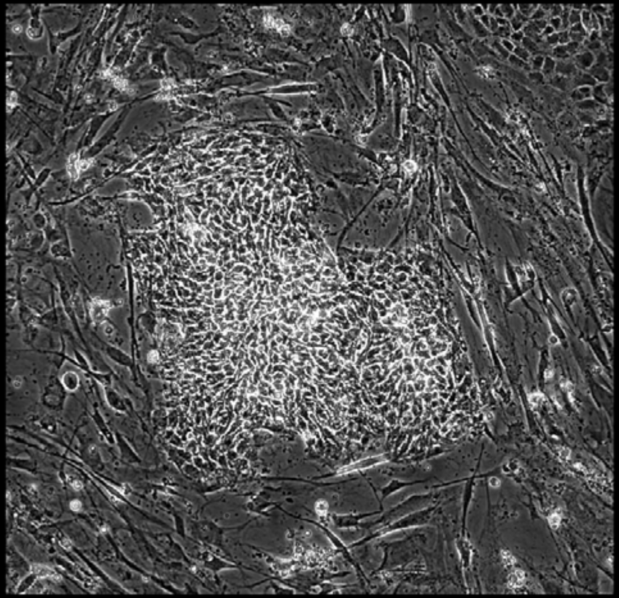

Well = 1, Colony = 172
Rater 1 = 2
Rater 2 = 2

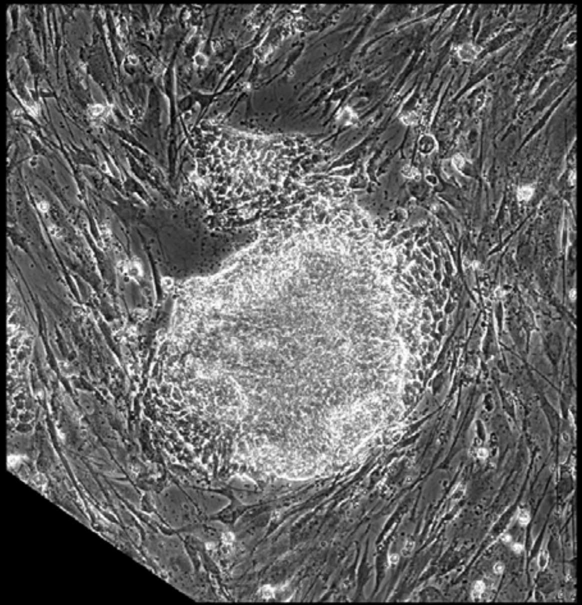

Well = 1, Colony = 173
Rater 1 = 2
Rater 2 = 2

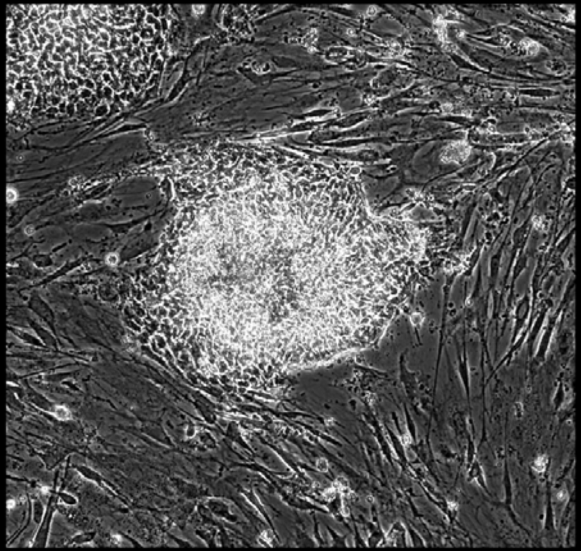

Well = 1, Colony = 174
Rater 1 = 4
Rater 2 = 4

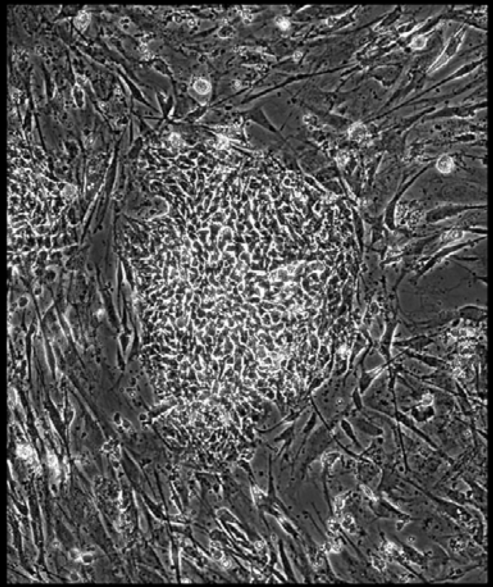

Well = 1, Colony = 175
Rater 1 = 1
Rater 2 = 1

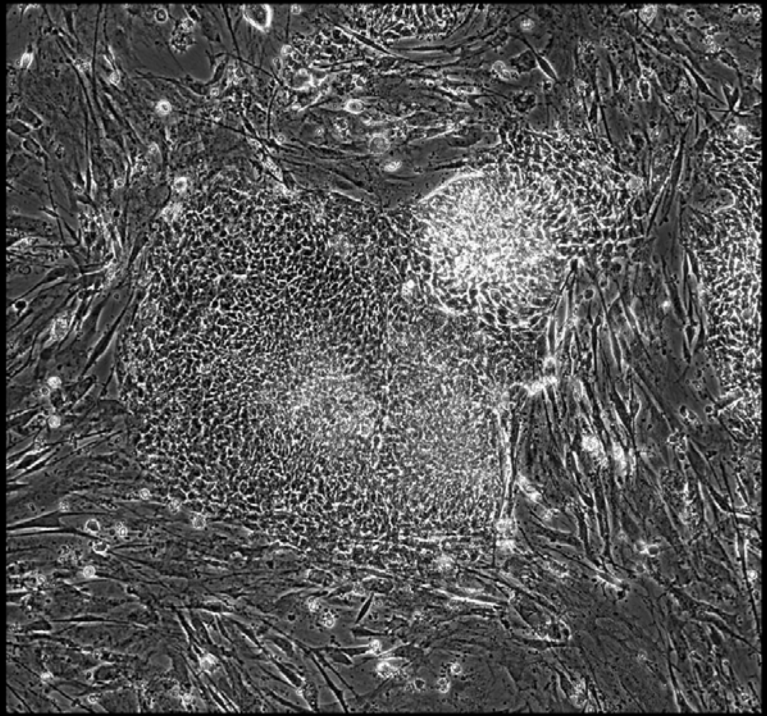

Well = 1, Colony = 176
Rater 1 = 4
Rater 2 = 3

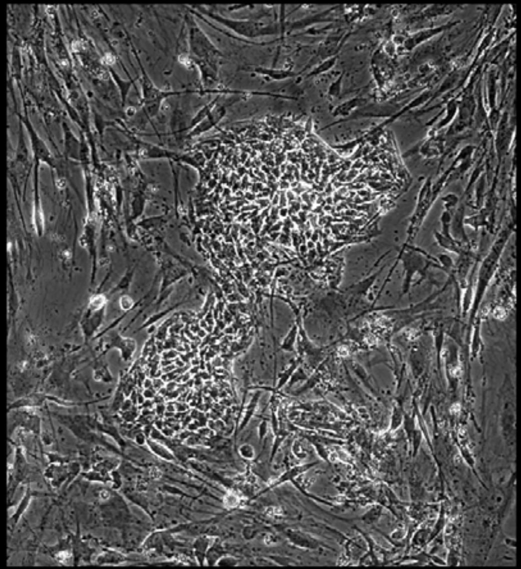

Well = 1, Colony = 177
Rater 1 = 4
Rater 2 = 4

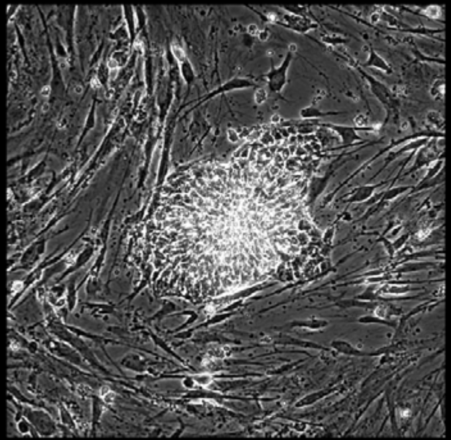

Well = 1, Colony = 178
Rater 1 = 4
Rater 2 = 3

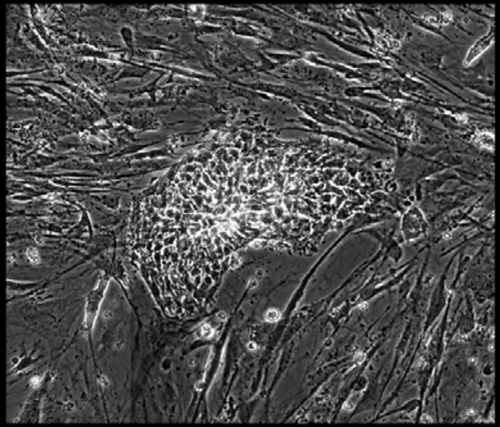

Well = 1, Colony = 179
Rater 1 = 5
Rater 2 = 5

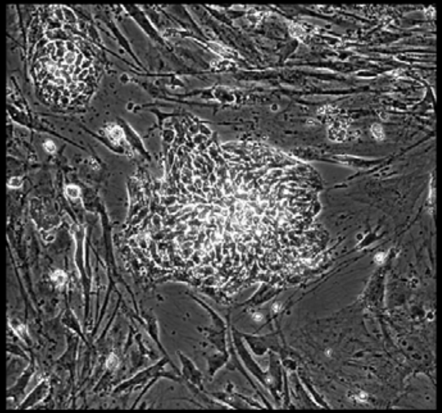

Well = 1, Colony = 180
Rater 1 = 5
Rater 2 = 4

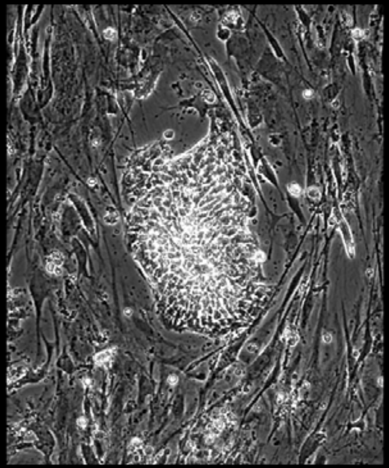

Well = 1, Colony = 181
Rater 1 = 2
Rater 2 = not scored

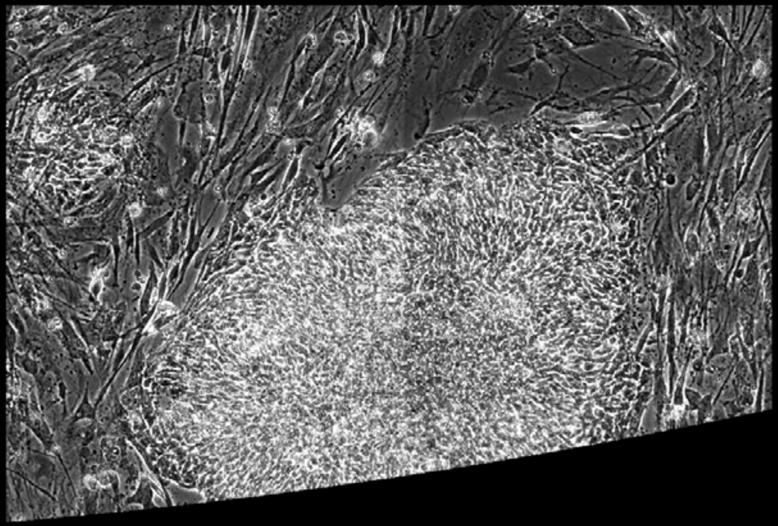

Well = 1, Colony = 182
Rater 1 = 4
Rater 2 = 5

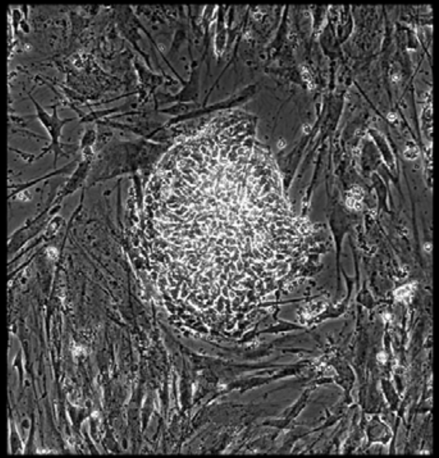

Well = 2, Colony = 1
Rater 1 = 1
Rater 2 = 2

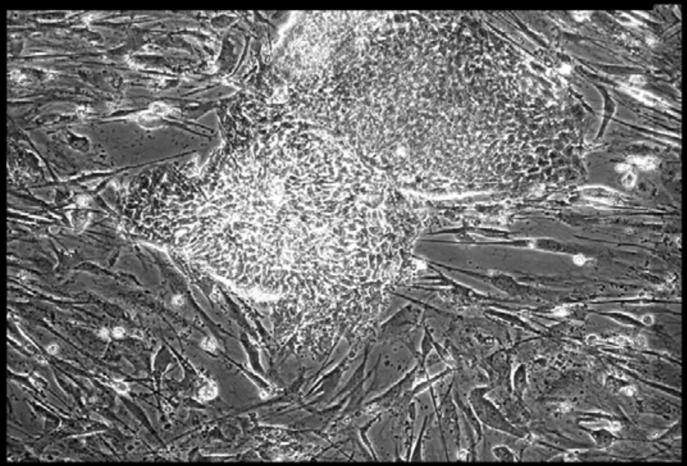

Well = 2, Colony = 2
Rater 1 = 1
Rater 2 = 2

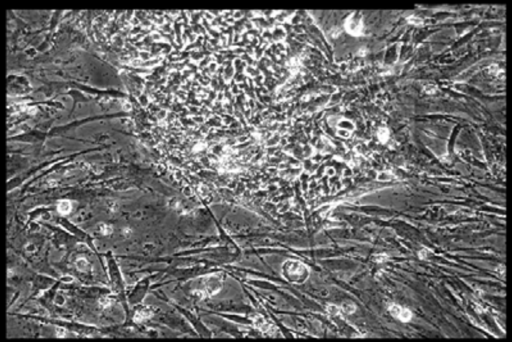

Well = 2, Colony = 3
Rater 1 = 3
Rater 2 = 2

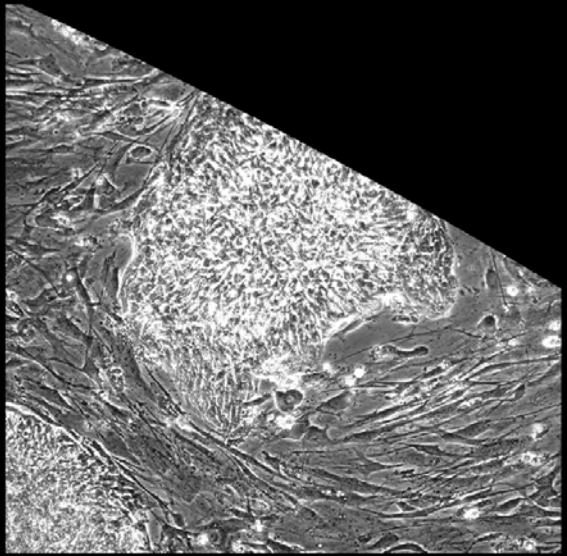

Well = 2, Colony = 4
Rater 1 = 2
Rater 2 = 2

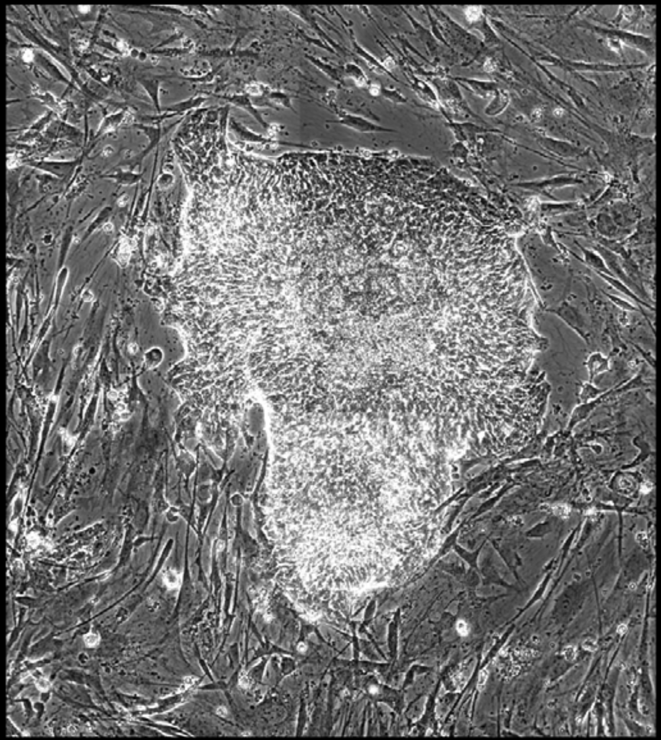

Well = 2, Colony = 5
Rater 1 = 2
Rater 2 = 2

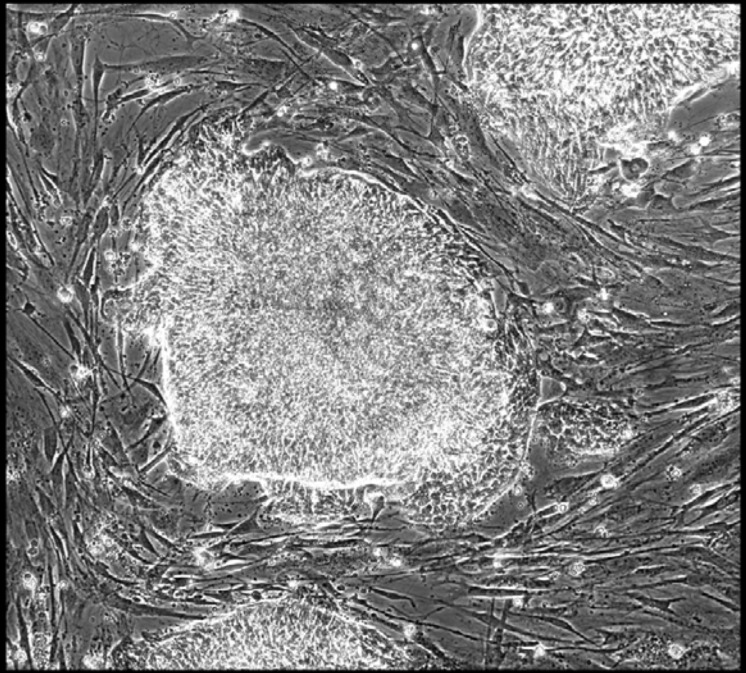

Well = 2, Colony = 6
Rater 1 = 3
Rater 2 = 2

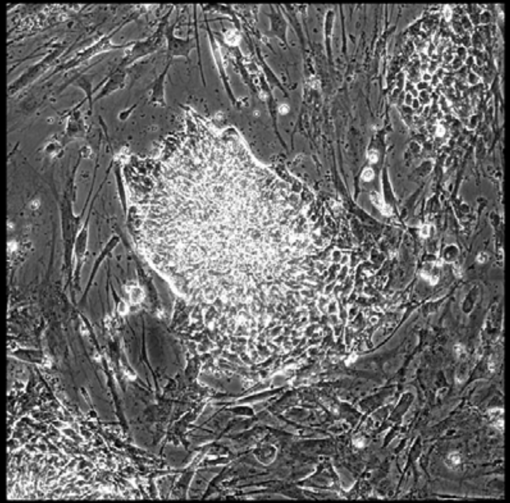

Well = 2, Colony = 7
Rater 1 = 3
Rater 2 = 4

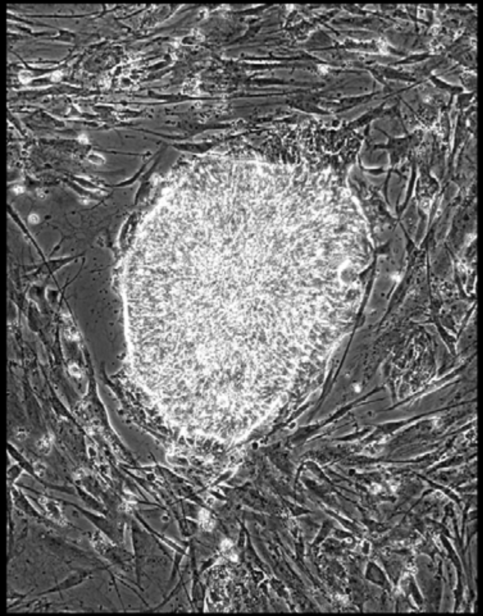

Well = 2, Colony = 8
Rater 1 = 4
Rater 2 = 5

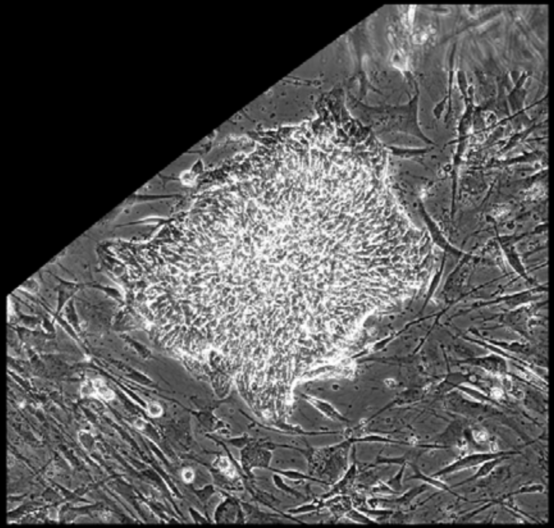

Well = 2, Colony = 9
Rater 1 = 4
Rater 2 = 4

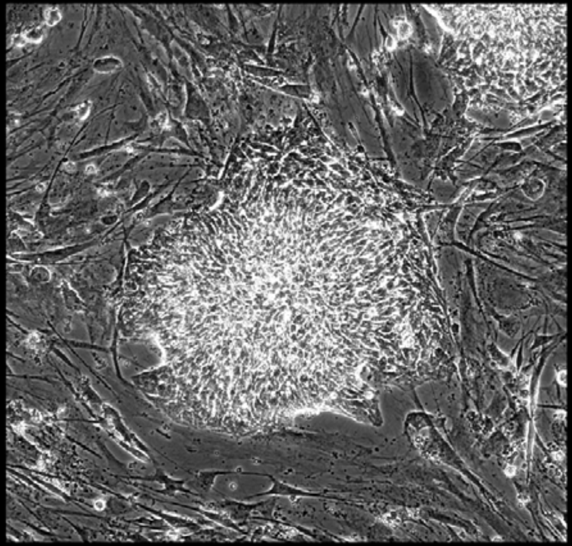

Well = 2, Colony = 10
Rater 1 = 4
Rater 2 = 3

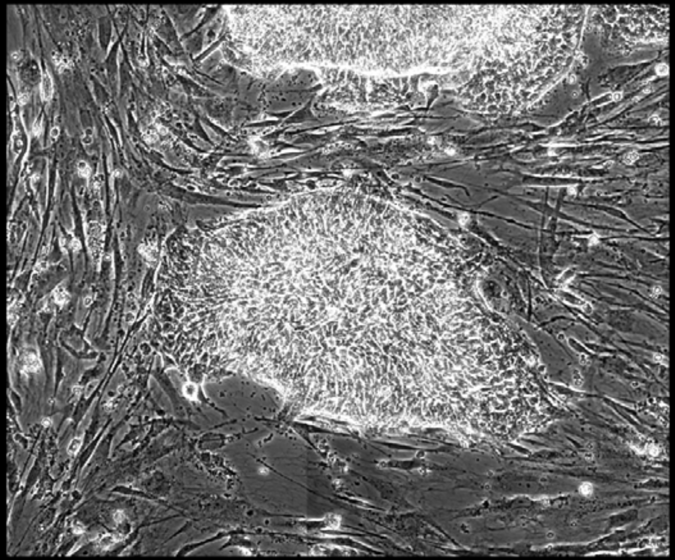

Well = 2, Colony = 11
Rater 1 = 4
Rater 2 = 4

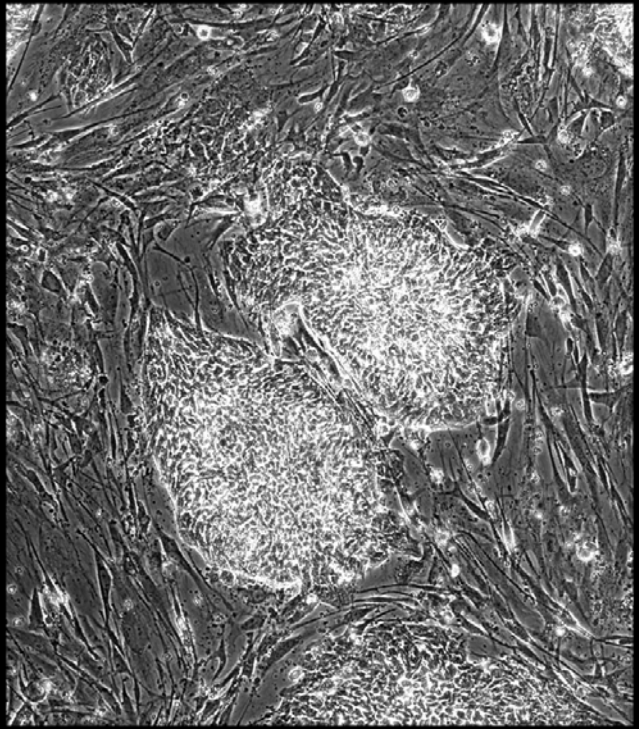

Well = 2, Colony = 12
Rater 1 = 3
Rater 2 = 4

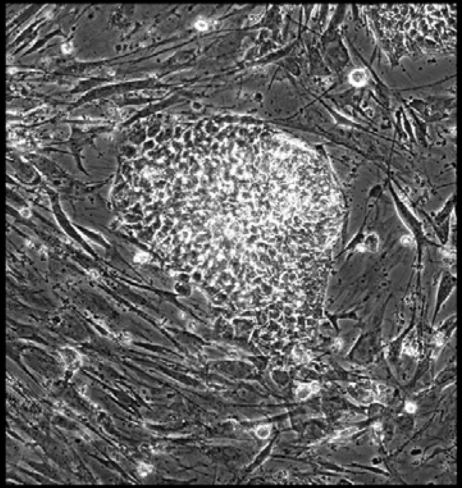

Well = 2, Colony = 13
Rater 1 = 3
Rater 2 = not scored

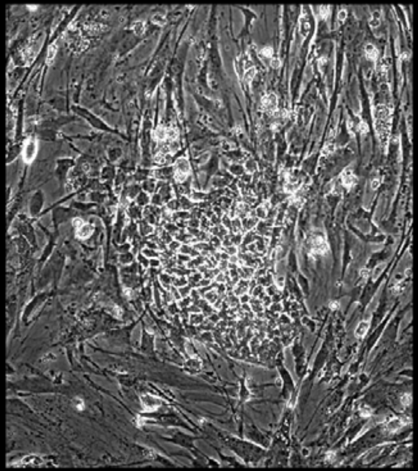

Well = 2, Colony = 14
Rater 1 = 3
Rater 2 = not scored

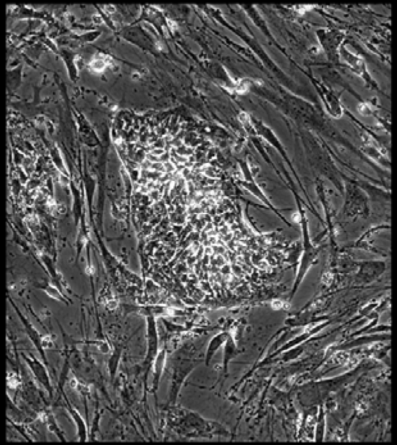

Well = 2, Colony = 15
Rater 1 = 4
Rater 2 = 3

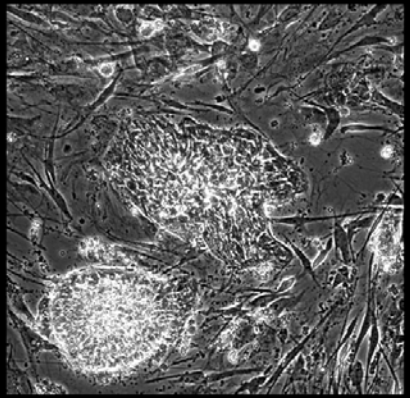

Well = 2, Colony = 16
Rater 1 = 4
Rater 2 = 3

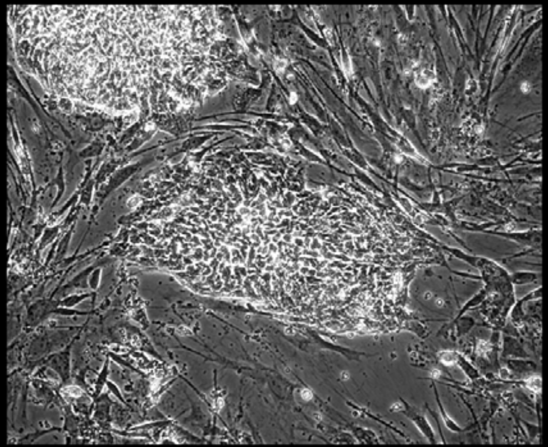

Well = 2, Colony = 17
Rater 1 = 4
Rater 2 = 4

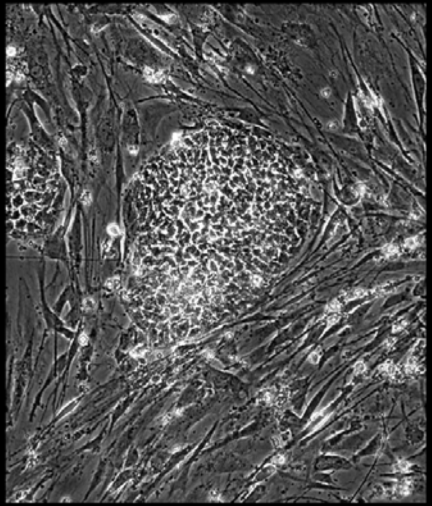

Well = 2, Colony = 18
Rater 1 = 4
Rater 2 = 3

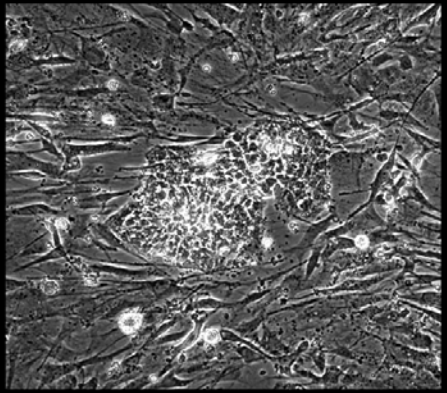

Well = 2, Colony = 19
Rater 1 = 3
Rater 2 = 2

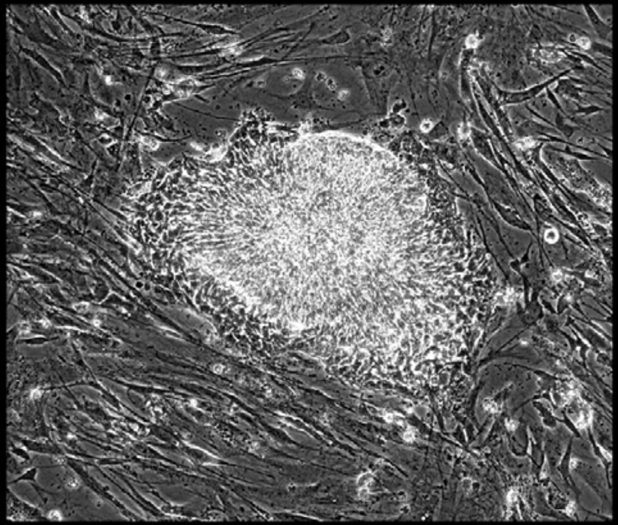

Well = 2, Colony = 20
Rater 1 = 3
Rater 2 = 3

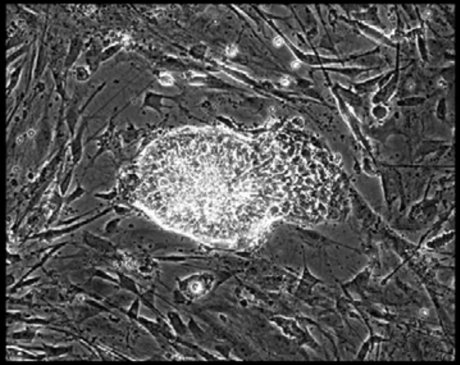

Well = 2, Colony = 21
Rater 1 = 4
Rater 2 = 4

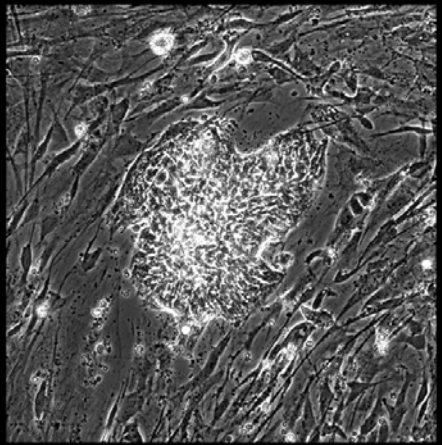

Well = 2, Colony = 22
Rater 1 = 4
Rater 2 = 4

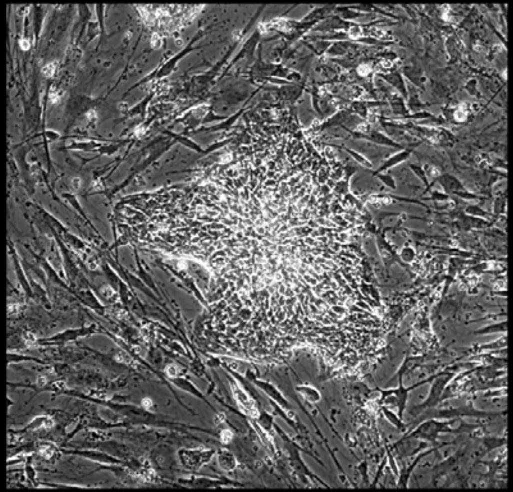

Well = 2, Colony = 23
Rater 1 = 3
Rater 2 = 3

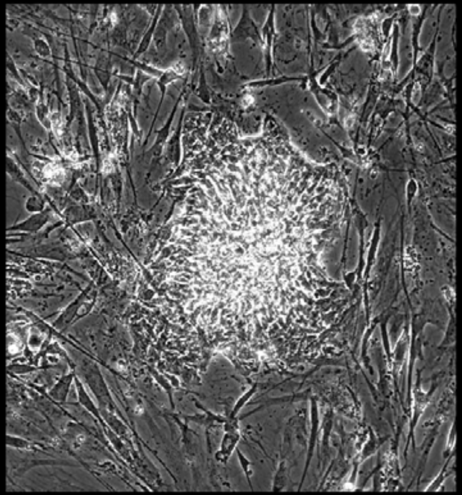

Well = 2, Colony = 24
Rater 1 = 3
Rater 2 = 3

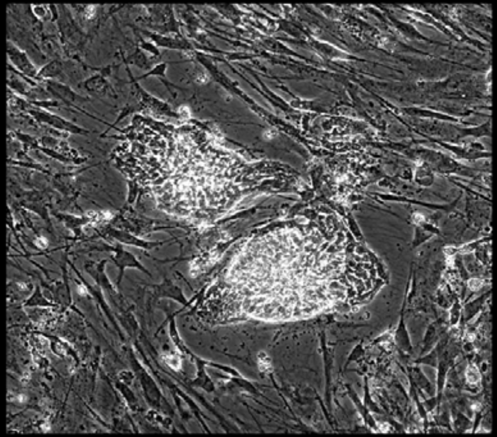

Well = 2, Colony = 25
Rater 1 = 3
Rater 2 = 4

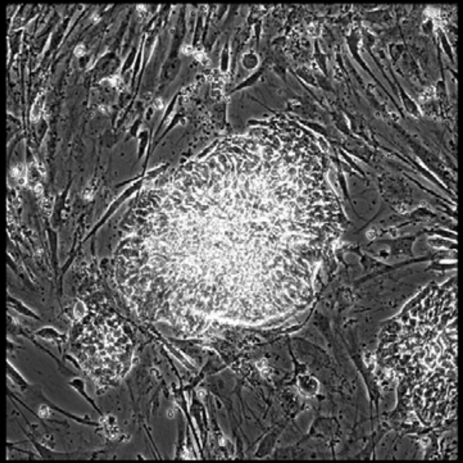

Well = 2, Colony = 26
Rater 1 = 3
Rater 2 = 4

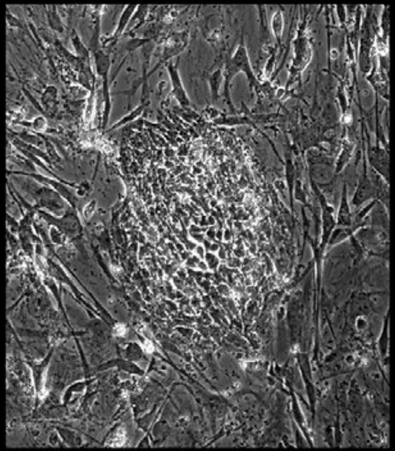

Well = 2, Colony = 27
Rater 1 = 3
Rater 2 = 4

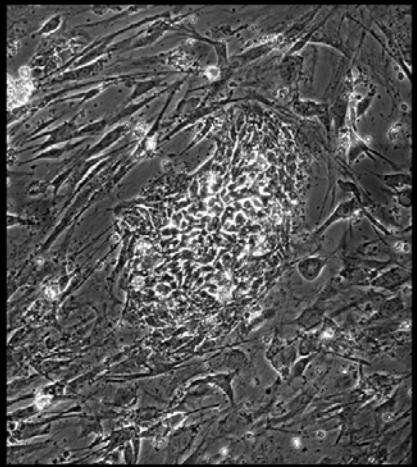

Well = 2, Colony = 28
Rater 1 = 4
Rater 2 = not scored

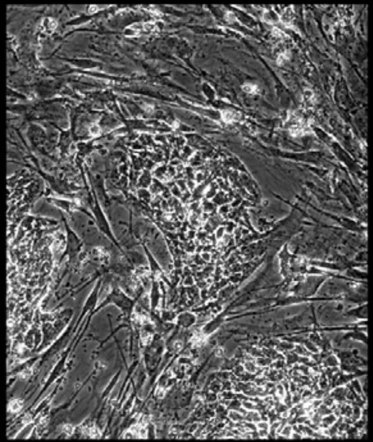

Well = 2, Colony = 29
Rater 1 = 4
Rater 2 = 4

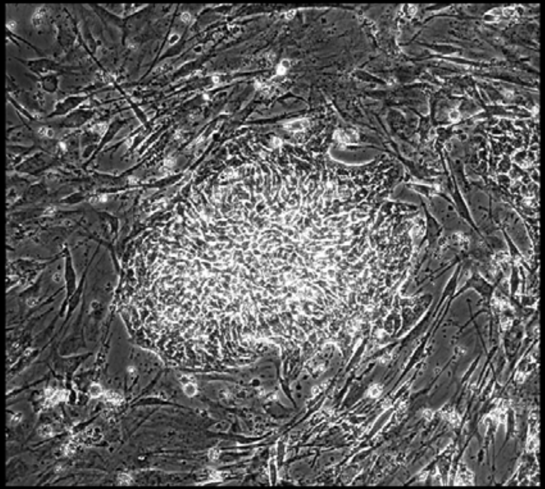

Well = 2, Colony = 30
Rater 1 = 2
Rater 2 = 2

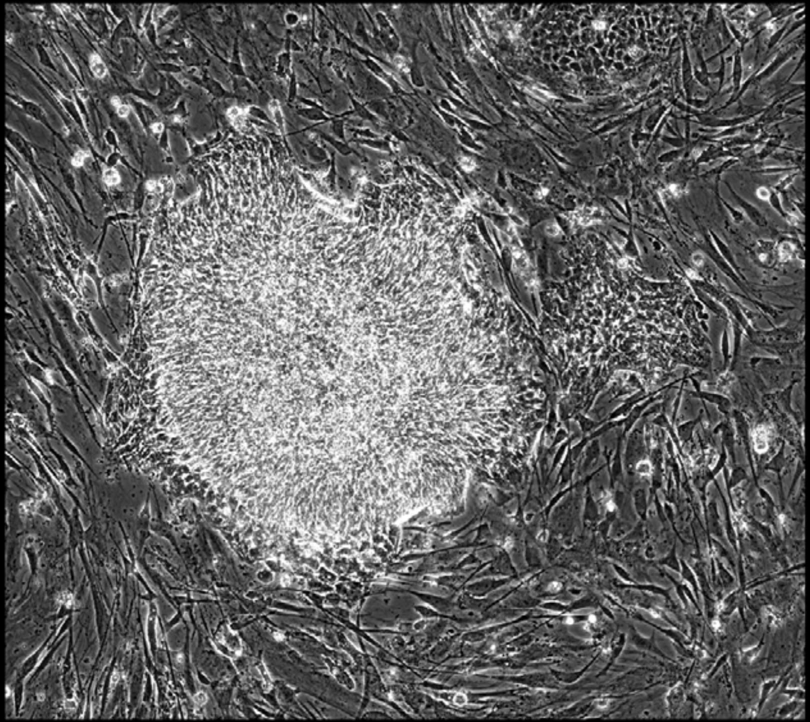

Well = 2, Colony = 31
Rater 1 = 4
Rater 2 = 4

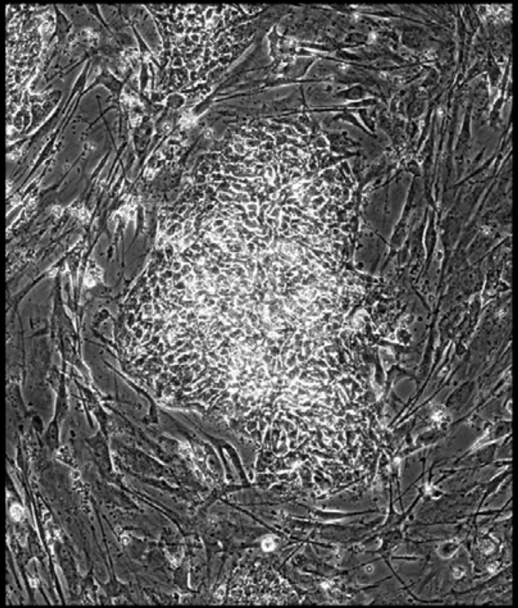

Well = 2, Colony = 32
Rater 1 = 2
Rater 2 = not scored

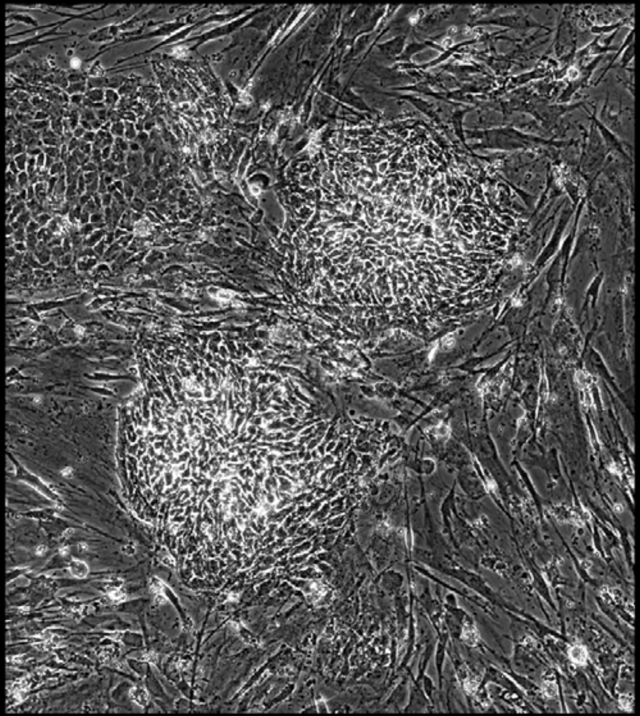

Well = 2, Colony = 33
Rater 1 = 1
Rater 2 = 2

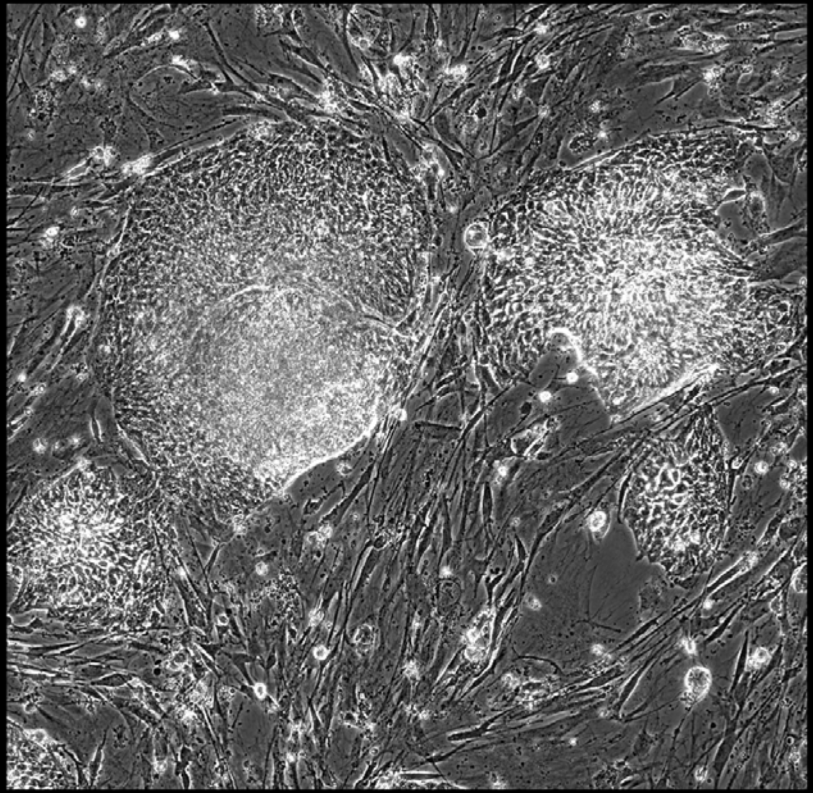

Well = 2, Colony = 34
Rater 1 = 4
Rater 2 = 5

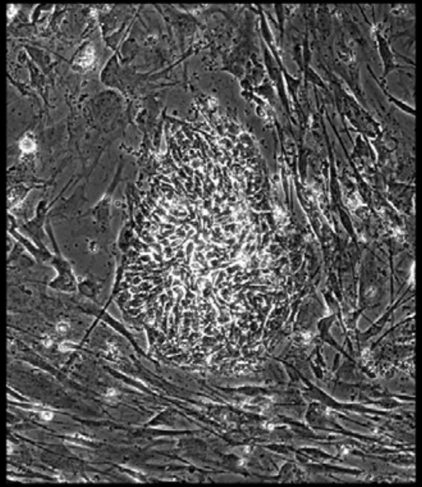

Well = 2, Colony = 35
Rater 1 = 3
Rater 2 = 3

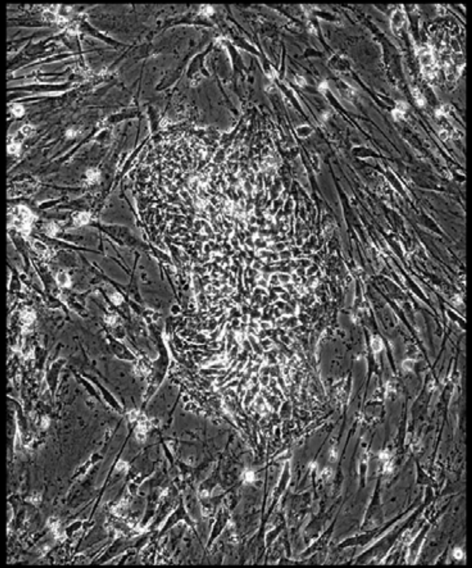

Well = 2, Colony = 36
Rater 1 = 3
Rater 2 = 4

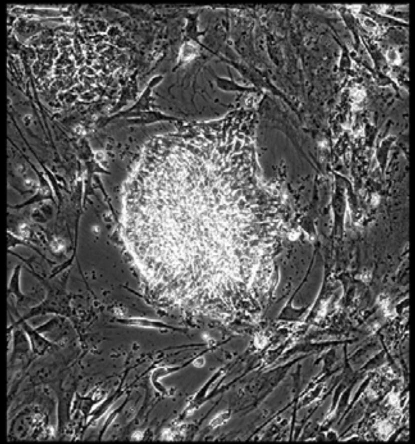

Well = 2, Colony = 37
Rater 1 = 4
Rater 2 = 4

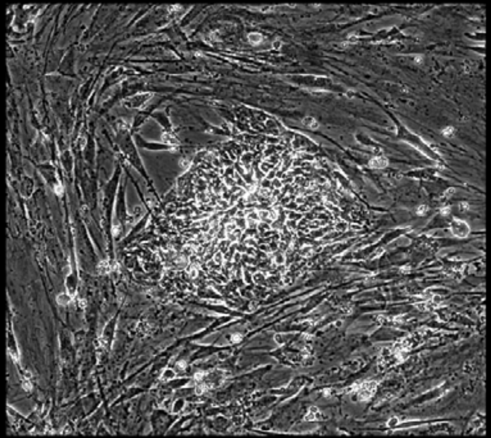

Well = 2, Colony = 38
Rater 1 = 2
Rater 2 = 2

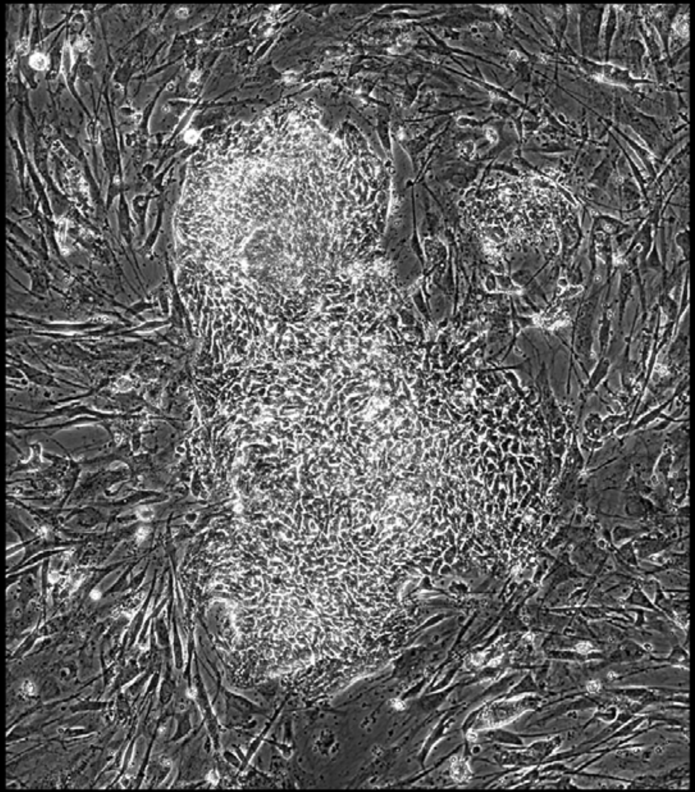

Well = 2, Colony = 39
Rater 1 = 3
Rater 2 = 4

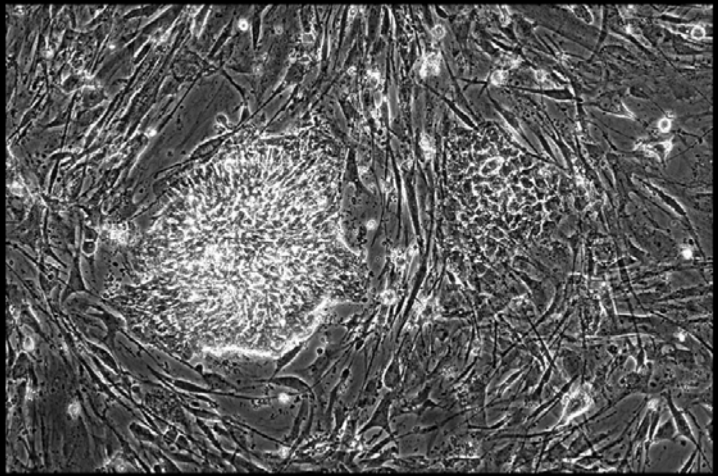

Well = 2, Colony = 40
Rater 1 = 4
Rater 2 = 4

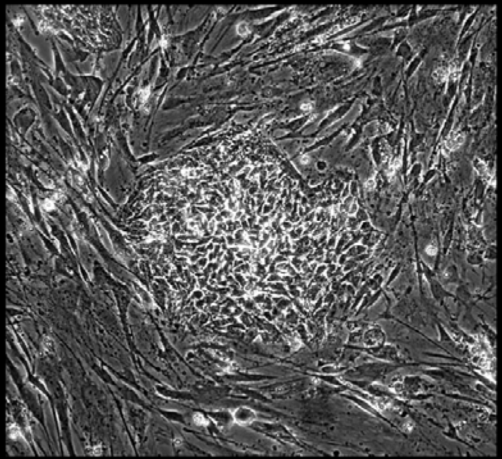

Well = 2, Colony = 41
Rater 1 = 3
Rater 2 = 4

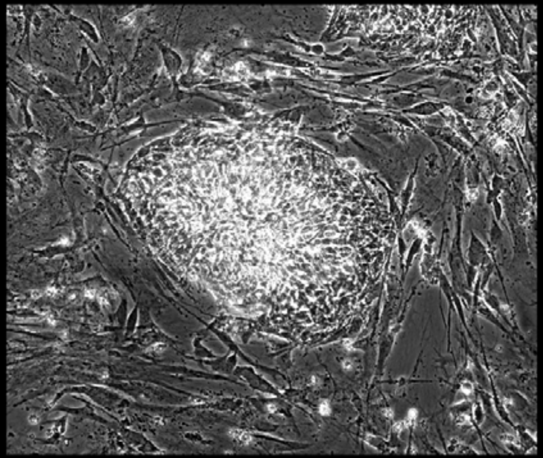

Well = 2, Colony = 42
Rater 1 = 4
Rater 2 = 3

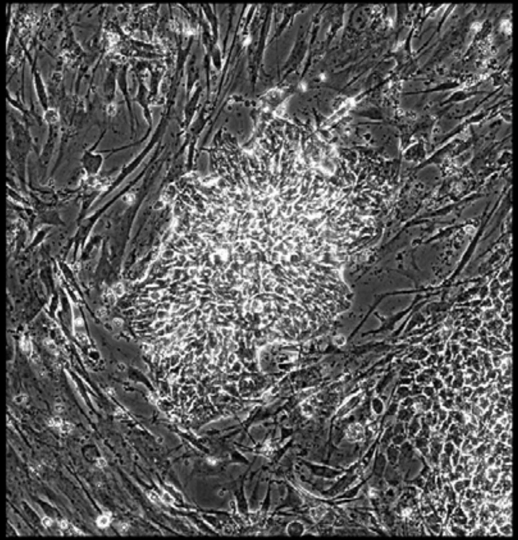

Well = 2, Colony = 43
Rater 1 = 3
Rater 2 = 2

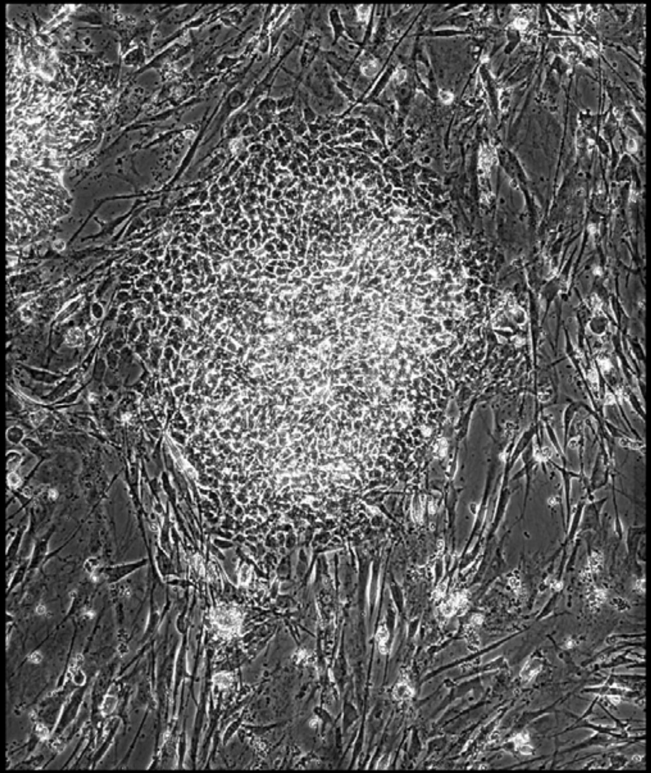

Well = 2, Colony = 44
Rater 1 = 3
Rater 2 = 4

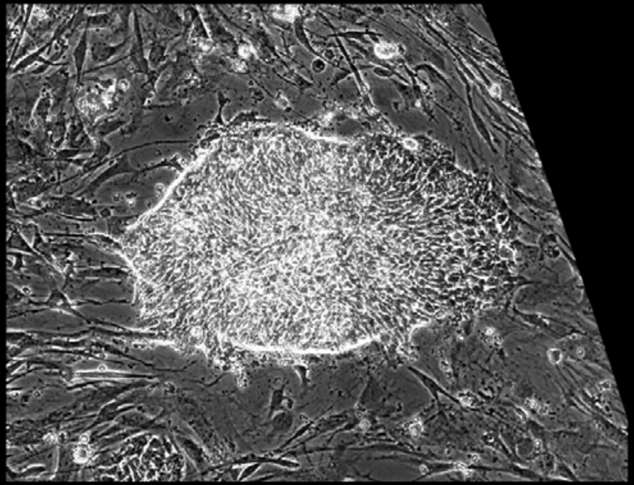

Well = 2, Colony = 45
Rater 1 = not scored
Rater 2 = not scored

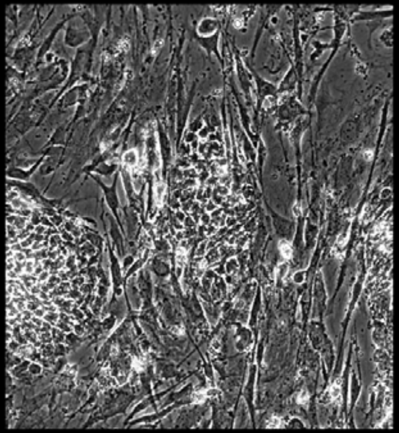

Well = 2, Colony = 46
Rater 1 = 3
Rater 2 = 3

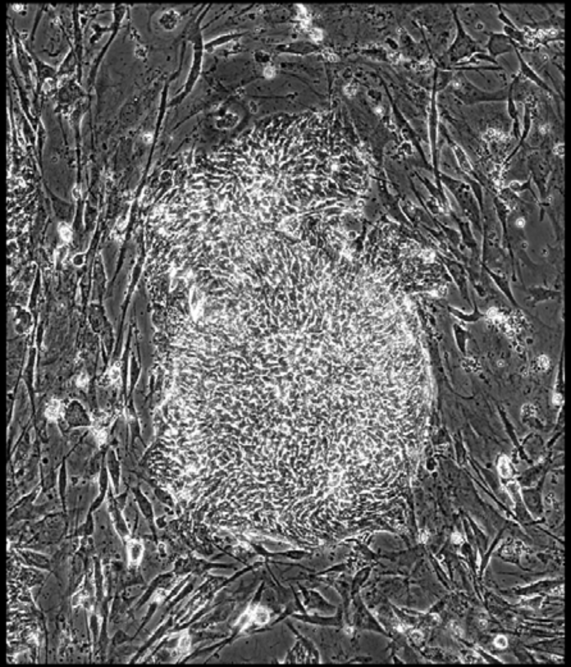

Well = 2, Colony = 47
Rater 1 = 4
Rater 2 = 5

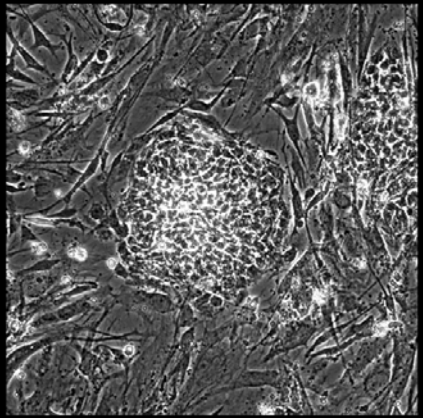

Well = 2, Colony = 48
Rater 1 = 4
Rater 2 = 5

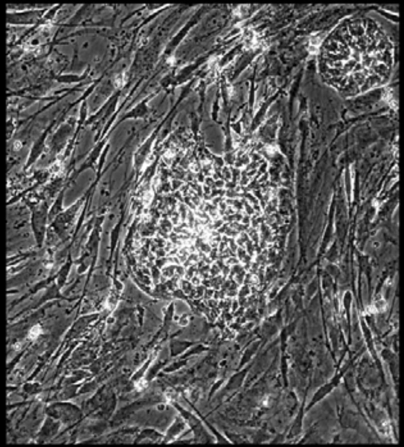

Well = 2, Colony = 49
Rater 1 = 3
Rater 2 = 3

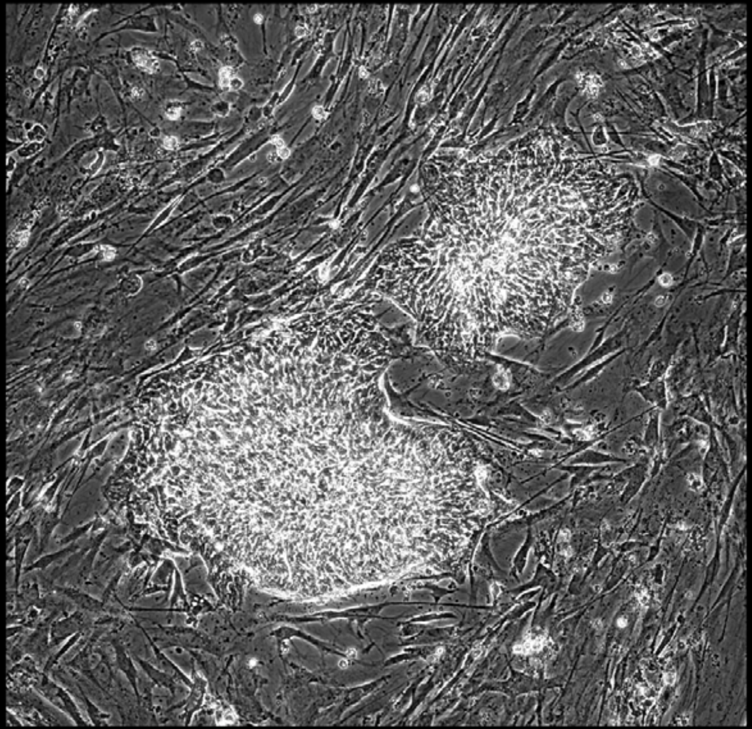

Well = 2, Colony = 50
Rater 1 = 4
Rater 2 = 4

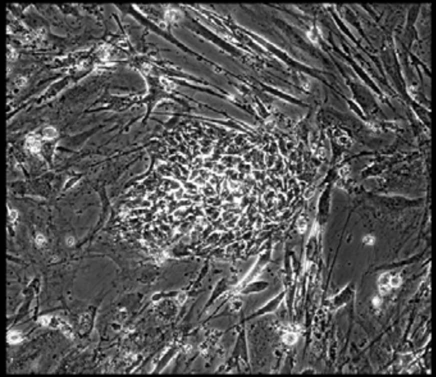

Well = 2, Colony = 51
Rater 1 = 4
Rater 2 = 4

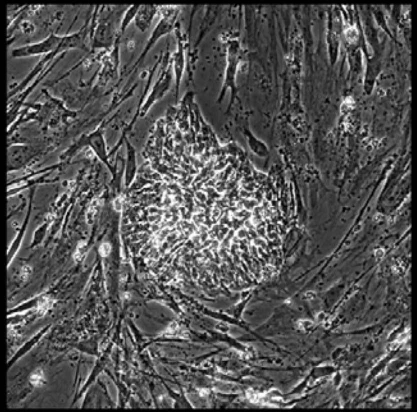

Well = 2, Colony = 52
Rater 1 = 4
Rater 2 = 4

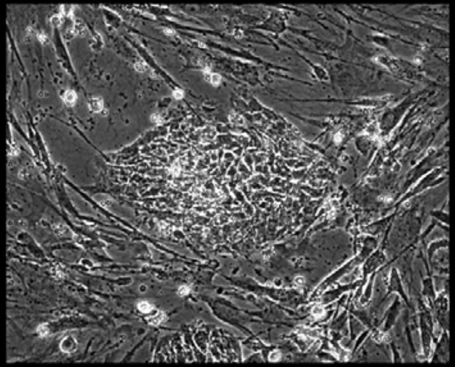

Well = 2, Colony = 53
Rater 1 = 4
Rater 2 = 3

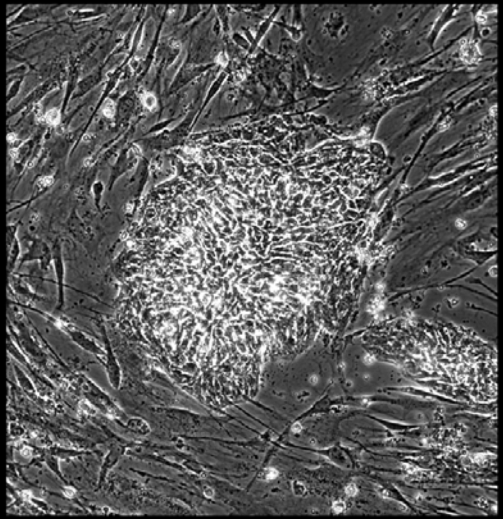

Well = 2, Colony = 54
Rater 1 = 3
Rater 2 = 3

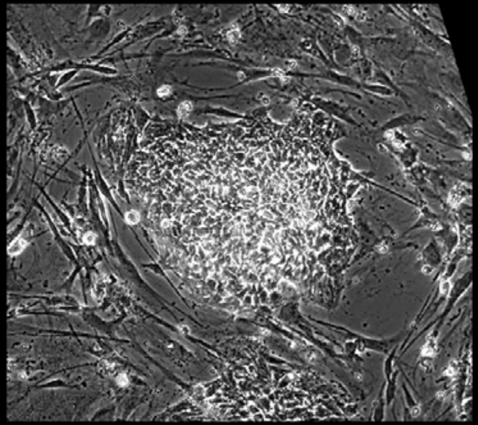

Well = 2, Colony = 55
Rater 1 = 4
Rater 2 = 4

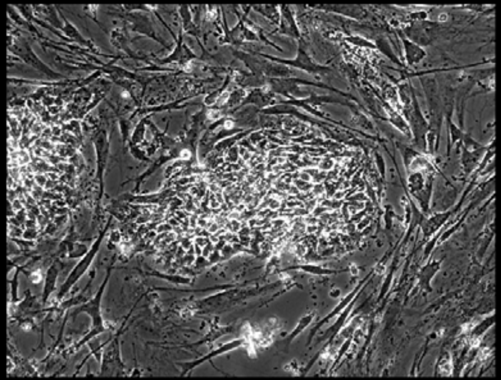

Well = 2, Colony = 56
Rater 1 = 4
Rater 2 = 3

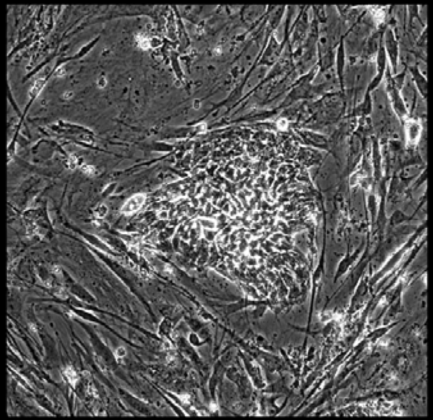

Well = 2, Colony = 57
Rater 1 = 3
Rater 2 = 4

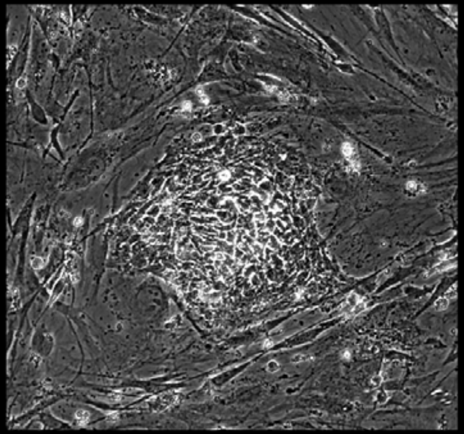

Well = 2, Colony = 58
Rater 1 = 3
Rater 2 = 3

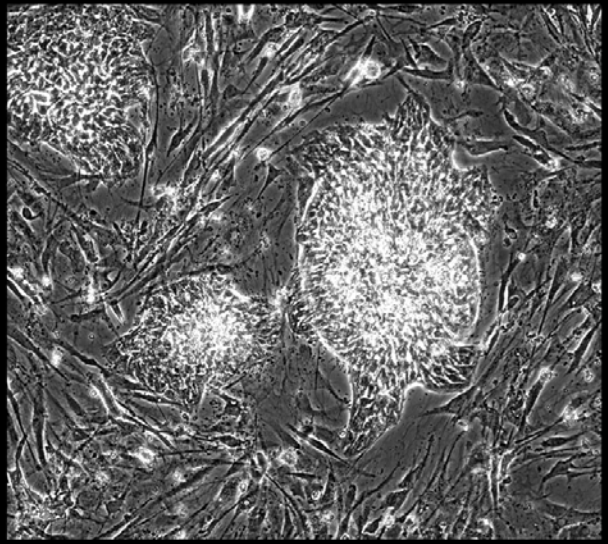

Well = 2, Colony = 59
Rater 1 = 4
Rater 2 = 5

Well = 2, Colony = 60
Rater 1 = 4
Rater 2 = 4

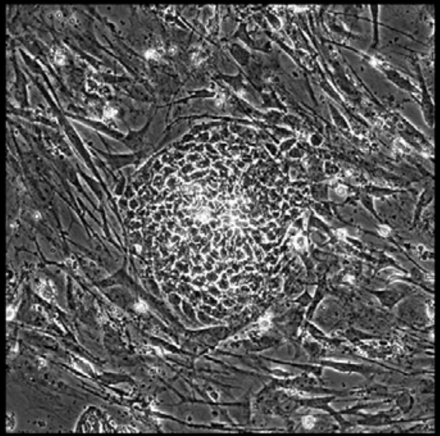

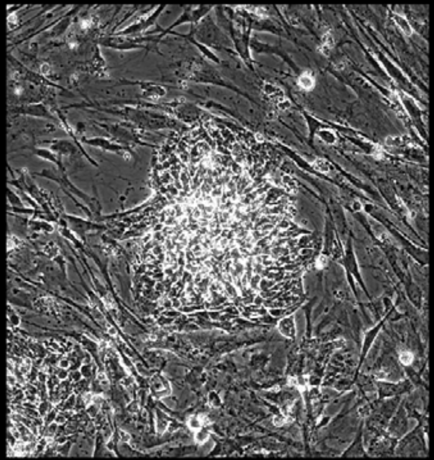

Well = 2, Colony = 61
Rater 1 = 2
Rater 2 = 2

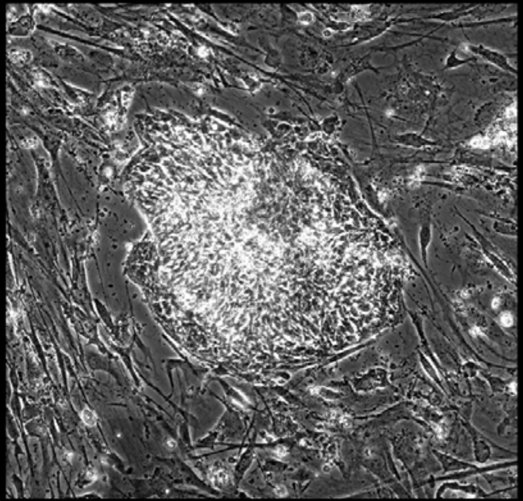

Well = 2, Colony = 62
Rater 1 = 2
Rater 2 = 2

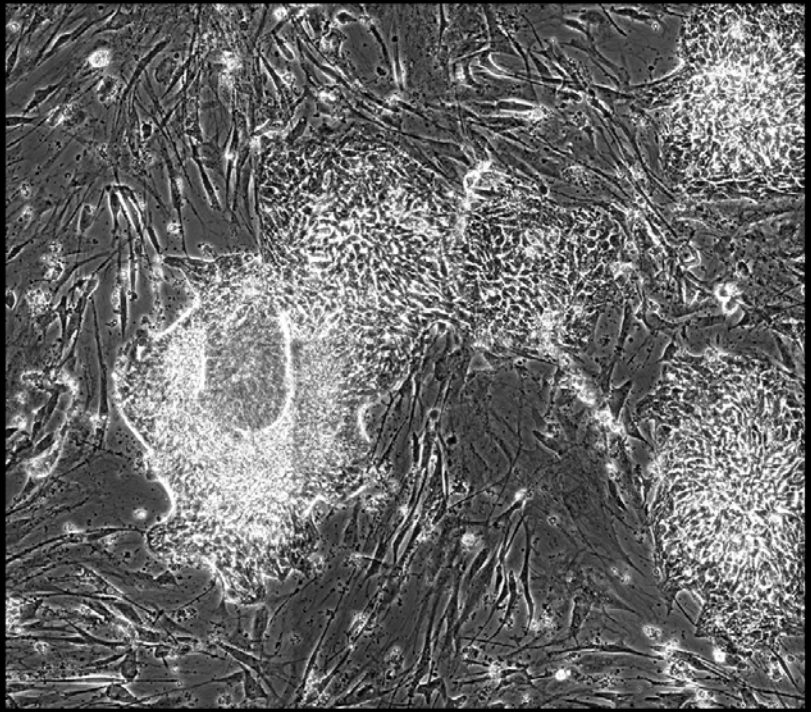

Well = 2, Colony = 63
Rater 1 = 4
Rater 2 = 4

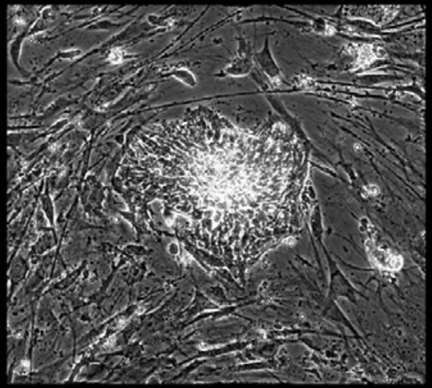

Well = 2, Colony = 64
Rater 1 = 3
Rater 2 = 2

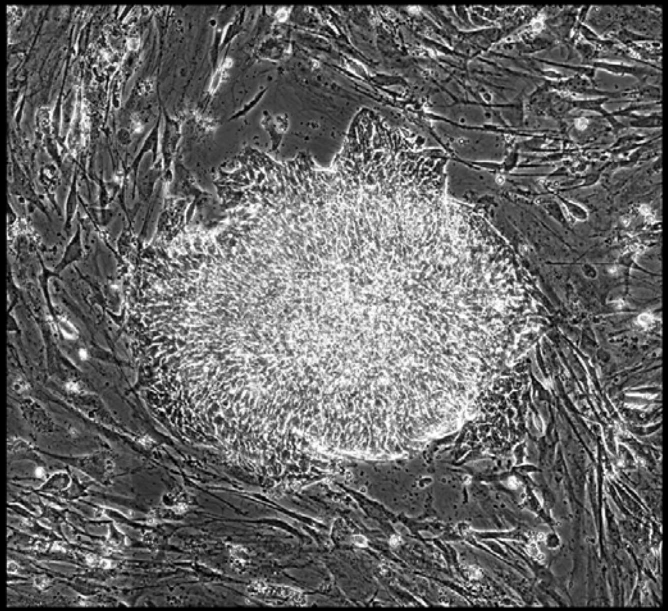

Well = 2, Colony = 65
Rater 1 = 2
Rater 2 = 3

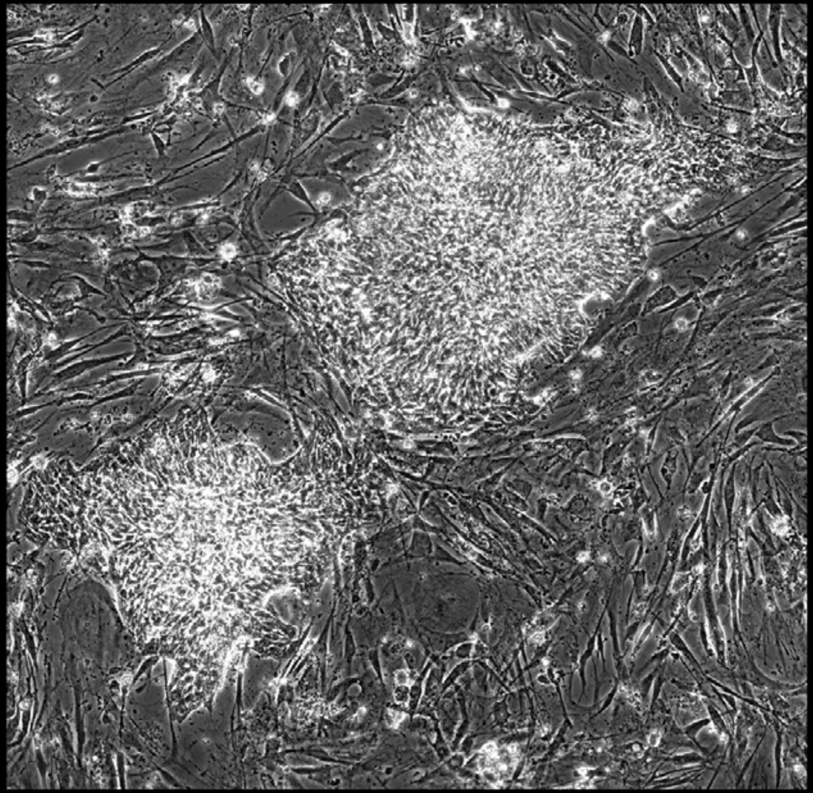

Well = 2, Colony = 66
Rater 1 = 2
Rater 2 = 2

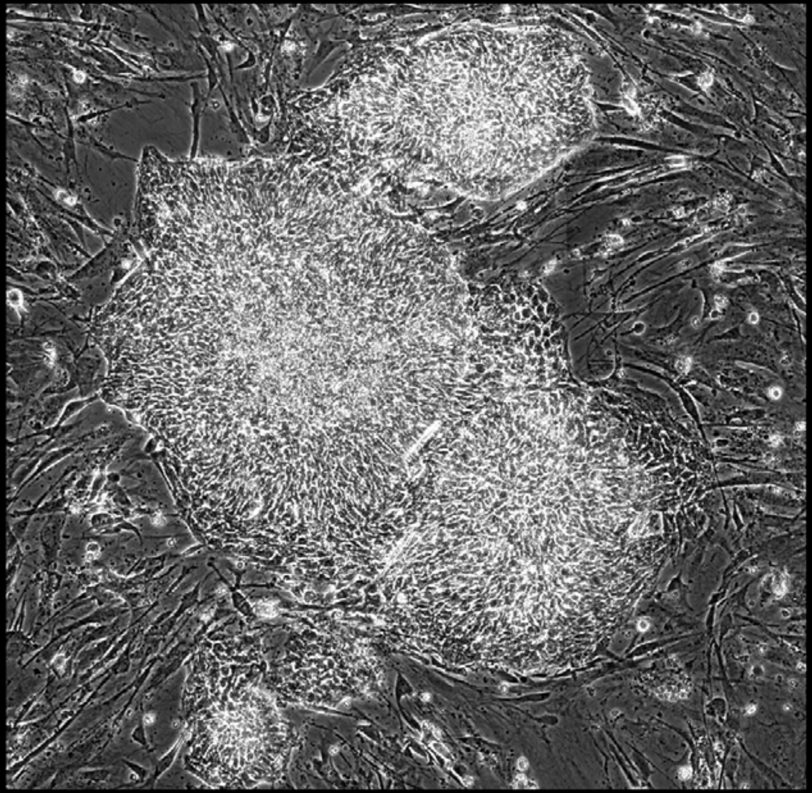

Well = 2, Colony = 67
Rater 1 = 4
Rater 2 = 4

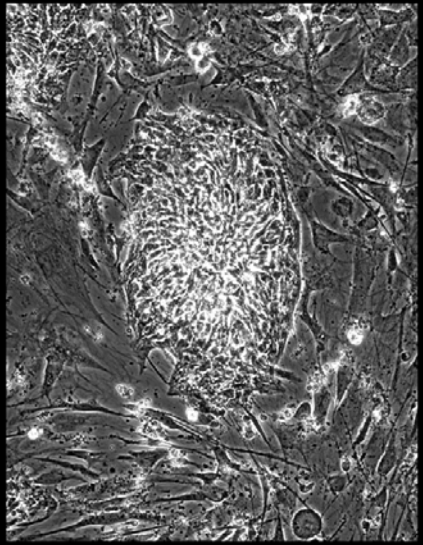

Well = 2, Colony = 68
Rater 1 = 4
Rater 2 = 3

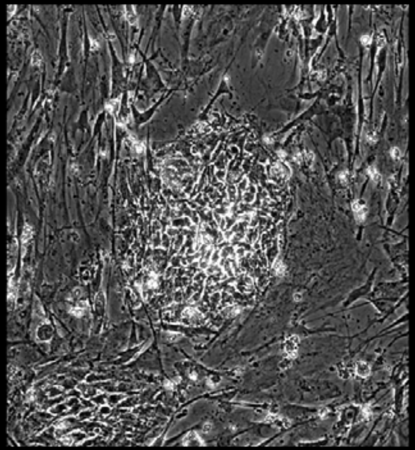

Well = 2, Colony = 69
Rater 1 = 5
Rater 2 = 4

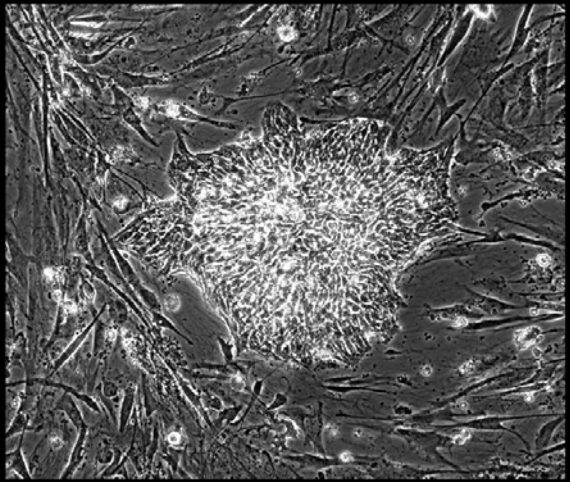

Well = 2, Colony = 70
Rater 1 = 4
Rater 2 = 4

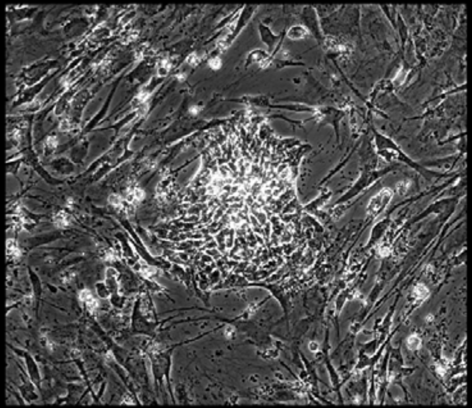

Well = 2, Colony = 71
Rater 1 = 4
Rater 2 = 3

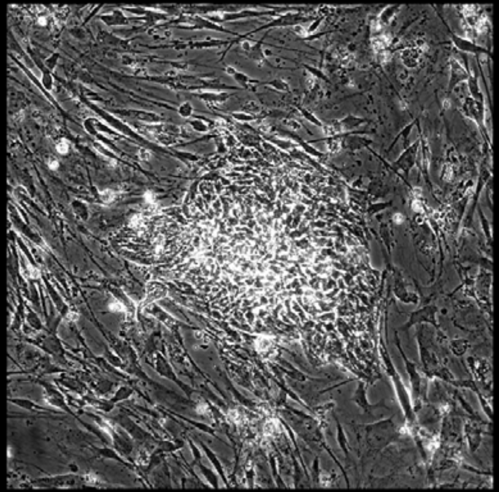

Well = 2, Colony = 72
Rater 1 = 5
Rater 2 = 4

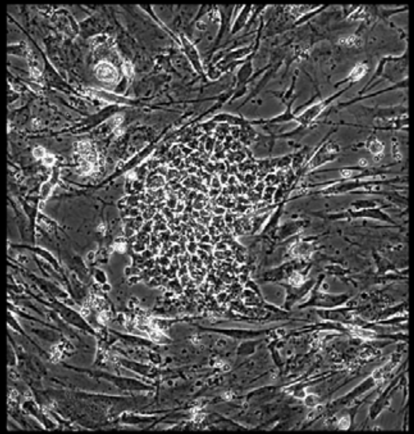

Well = 2, Colony = 73
Rater 1 = 4
Rater 2 = 4

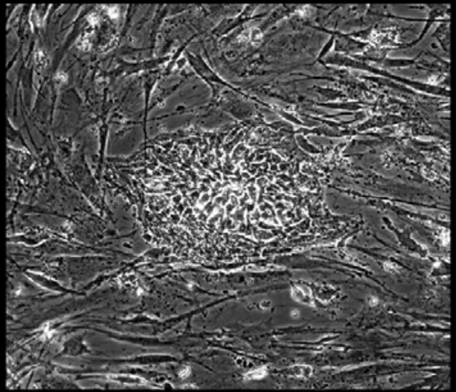

Well = 2, Colony = 74
Rater 1 = 3
Rater 2 = 2

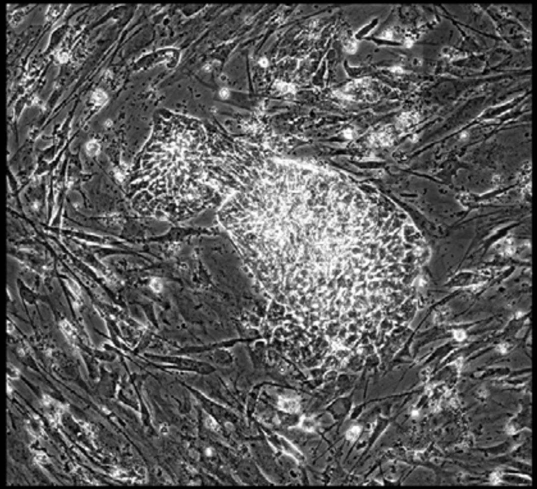

Well = 2, Colony = 75
Rater 1 = 4
Rater 2 = 3

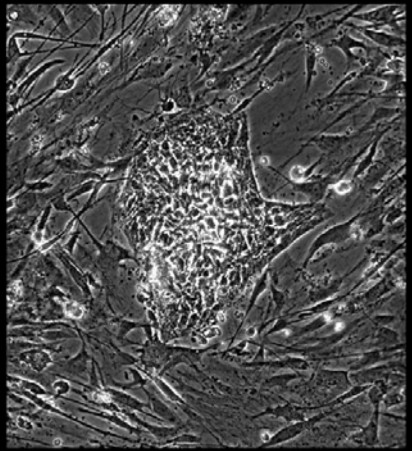

Well = 2, Colony = 76
Rater 1 = 4
Rater 2 = 4

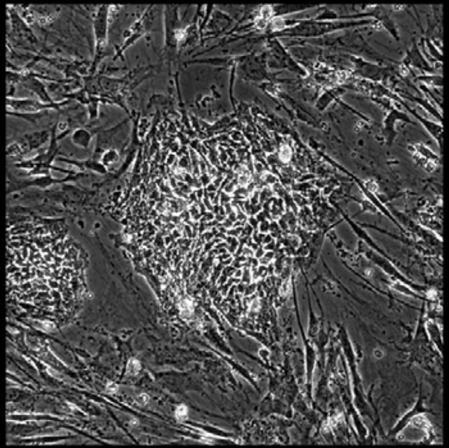

Well = 2, Colony = 77
Rater 1 = 4
Rater 2 = 3

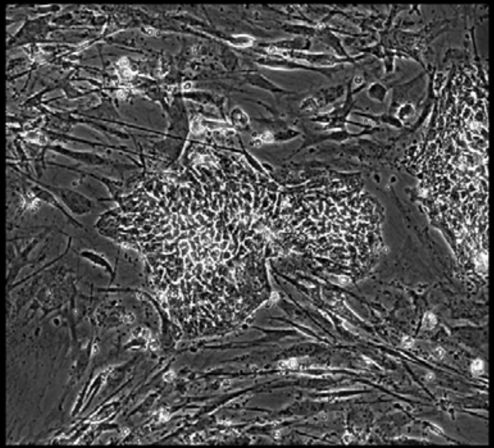

Well = 2, Colony = 78
Rater 1 = 4
Rater 2 = 4

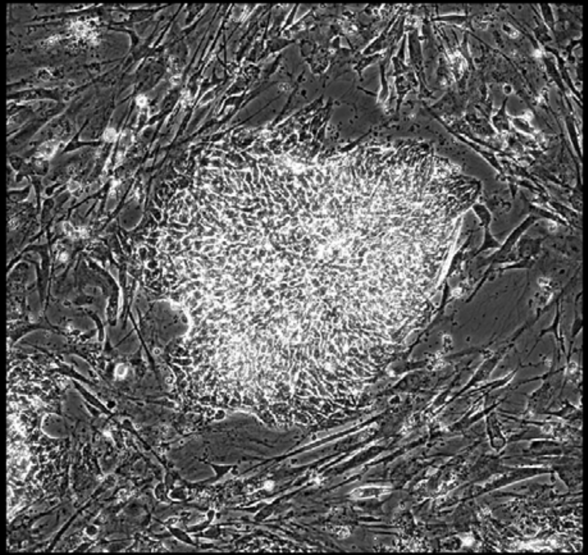

Well = 2, Colony = 79
Rater 1 = 4
Rater 2 = 3

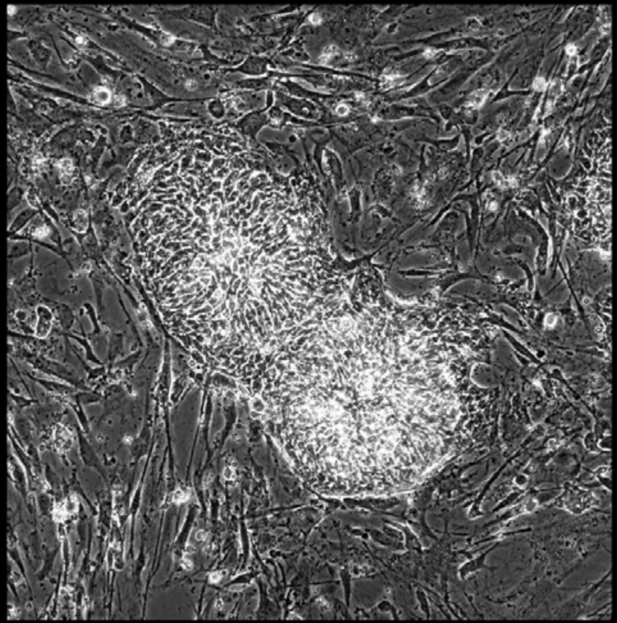

Well = 2, Colony = 80
Rater 1 = 3
Rater 2 = 4

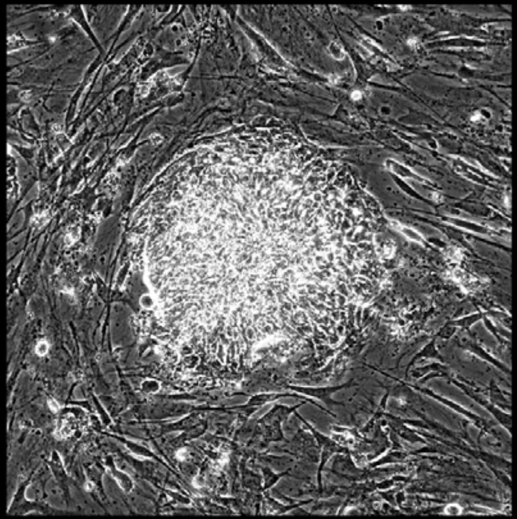

Well = 2, Colony = 81
Rater 1 = 3
Rater 2 = 5

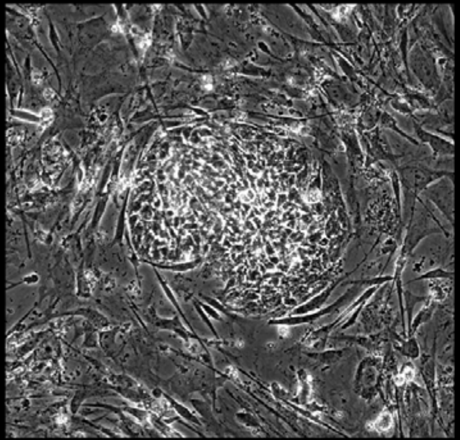

Well = 2, Colony = 82
Rater 1 = 2
Rater 2 = 3

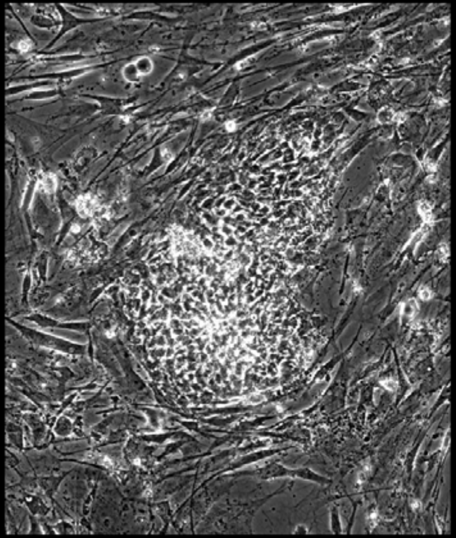

Well = 2, Colony = 83
Rater 1 = 4
Rater 2 = 4

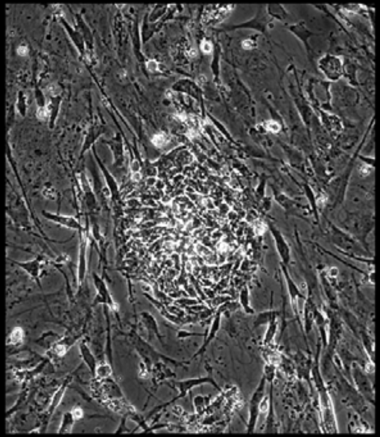

Well = 2, Colony = 84
Rater 1 = 4
Rater 2 = 4

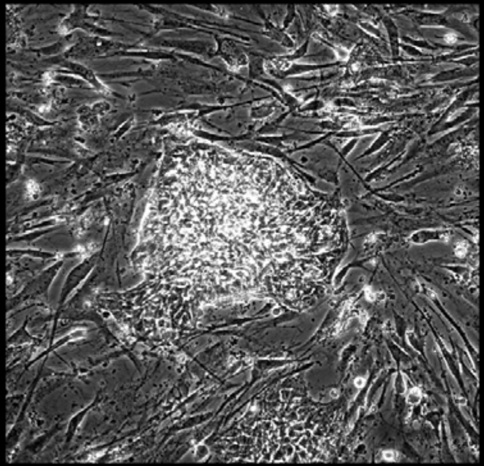

Well = 2, Colony = 85
Rater 1 = 5
Rater 2 = 4

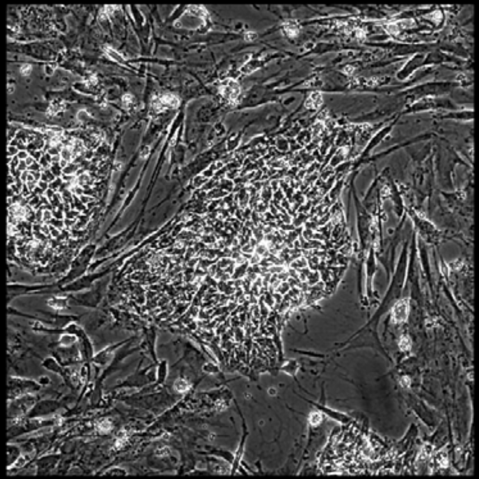

Well = 2, Colony = 86
Rater 1 = 5
Rater 2 = 4

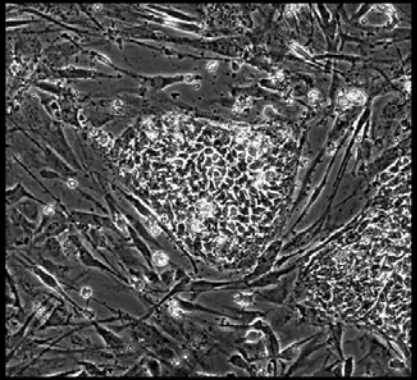

Well = 2, Colony = 87
Rater 1 = 5
Rater 2 = 5

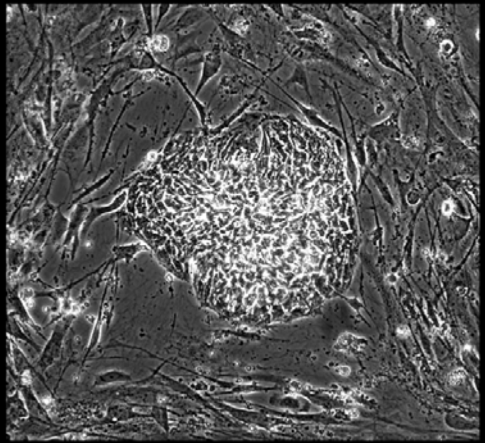

Well = 2, Colony = 88
Rater 1 = 4
Rater 2 = 3

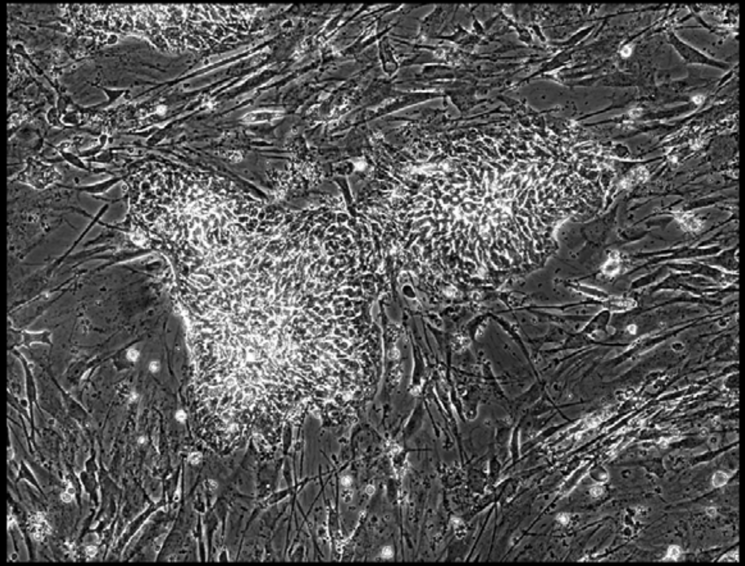

Well = 2, Colony = 89
Rater 1 = 4
Rater 2 = 2

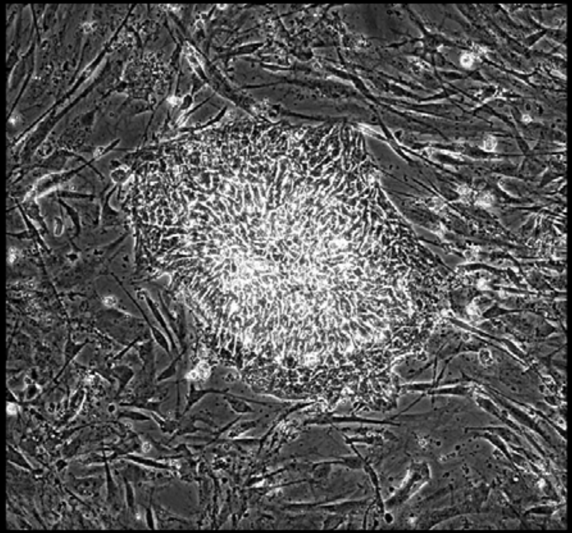

Well = 2, Colony = 90
Rater 1 = 1
Rater 2 = 1

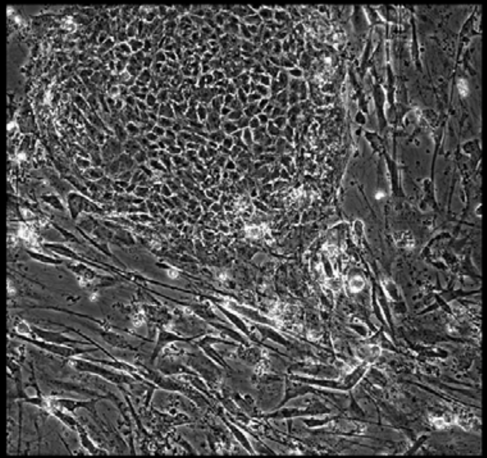

Well = 2, Colony = 91
Rater 1 = 4
Rater 2 = 4

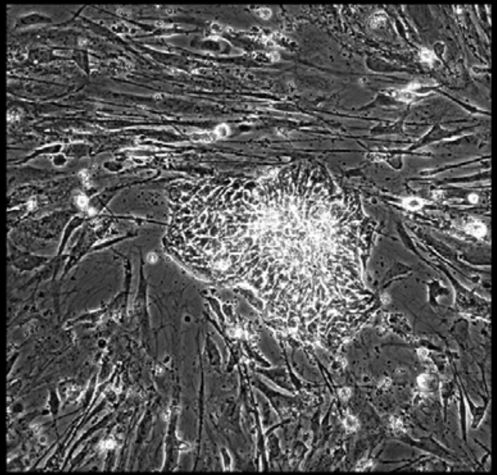

Well = 2, Colony = 92
Rater 1 = 4
Rater 2 = 4

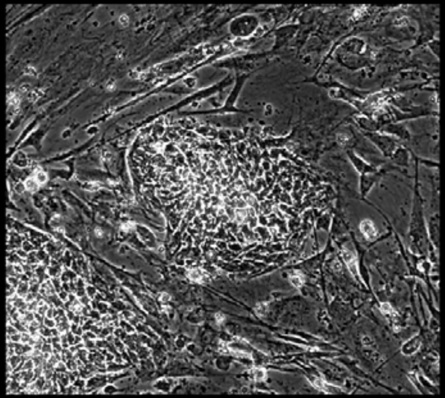

Well = 2, Colony = 93
Rater 1 = 4
Rater 2 = 4

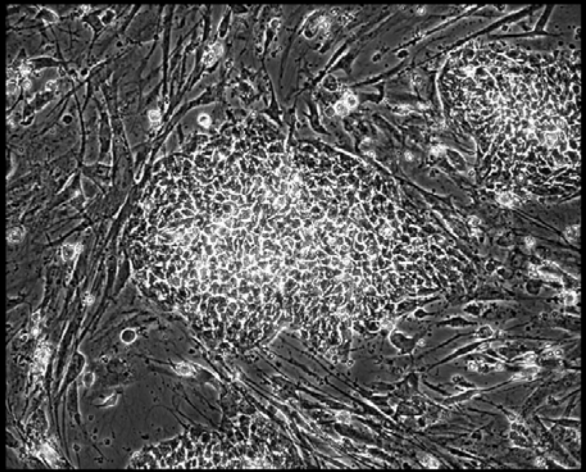

Well = 2, Colony = 94
Rater 1 = 4
Rater 2 = 4

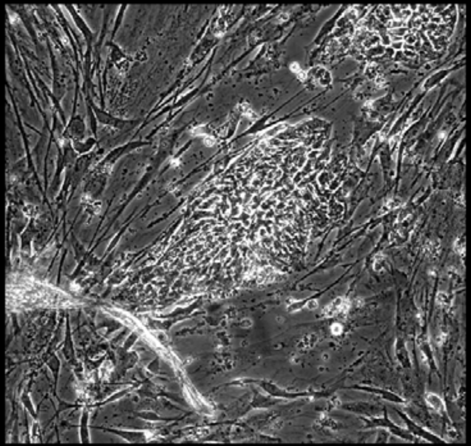

Well = 2, Colony = 95
Rater 1 = 3
Rater 2 = 4

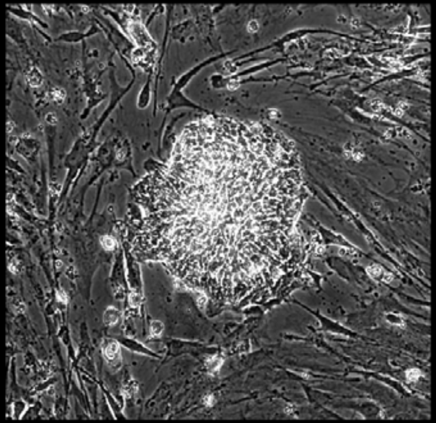

Well = 2, Colony = 96
Rater 1 = 4
Rater 2 = 5

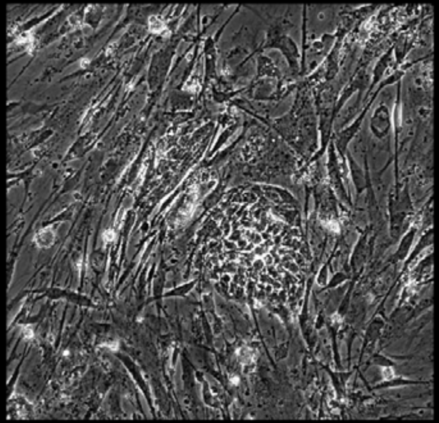

Well = 2, Colony = 97
Rater 1 = 3
Rater 2 = 3

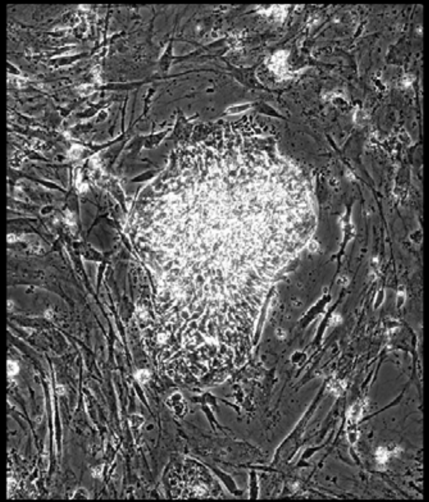

Well = 2, Colony = 98
Rater 1 = 4
Rater 2 = 5

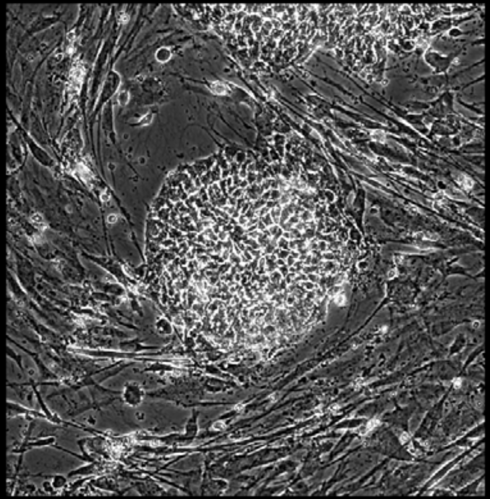

Well = 2, Colony = 99
Rater 1 = 2
Rater 2 = 4

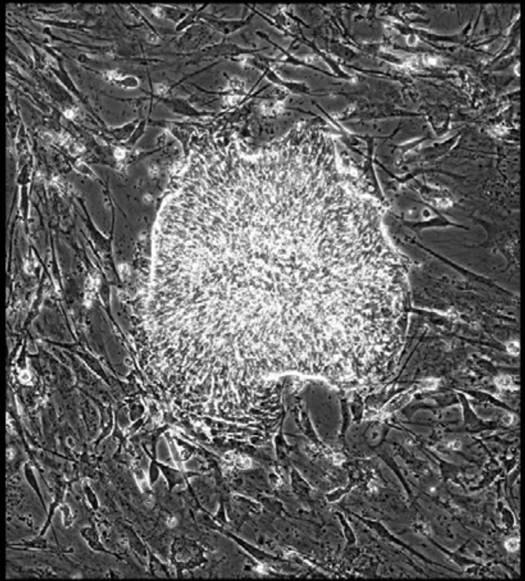

Well = 2, Colony = 100
Rater 1 = 4
Rater 2 = 4

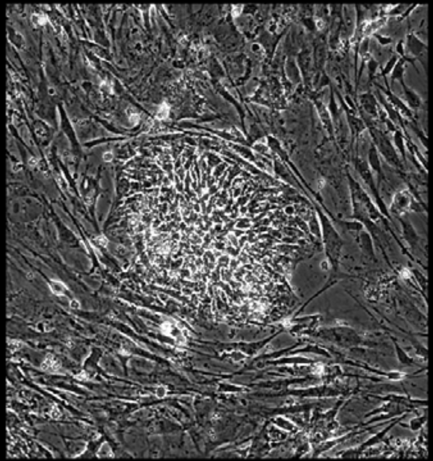

Well = 2, Colony = 101
Rater 1 = 4
Rater 2 = 4

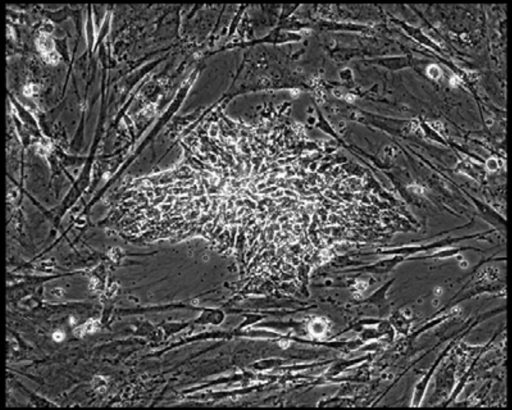

Well = 2, Colony = 102
Rater 1 = 4
Rater 2 = 4

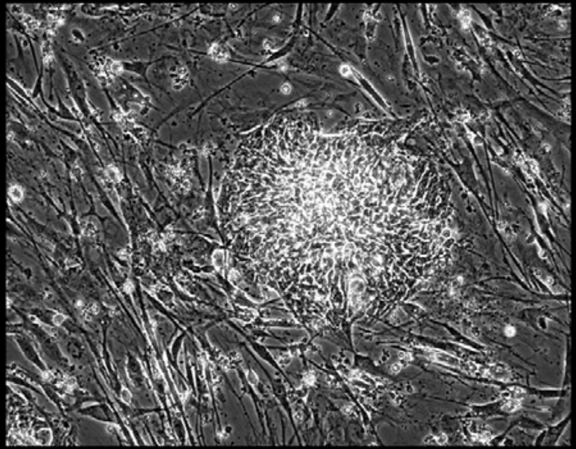

Well = 2, Colony = 103
Rater 1 = 3
Rater 2 = 3

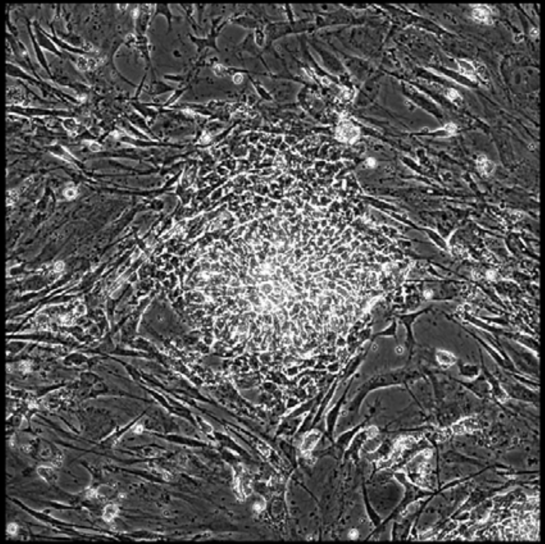

Well = 2, Colony = 104
Rater 1 = 2
Rater 2 = 3

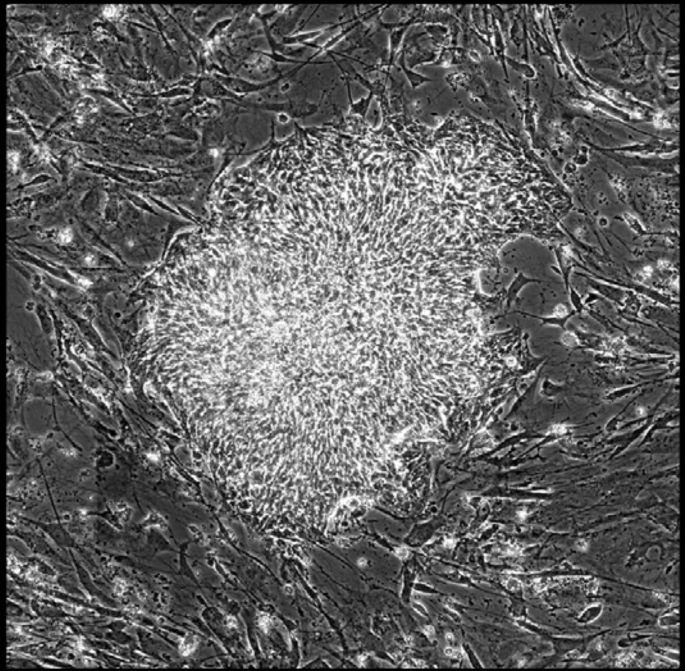

Well = 2, Colony = 105
Rater 1 = 3
Rater 2 = 3

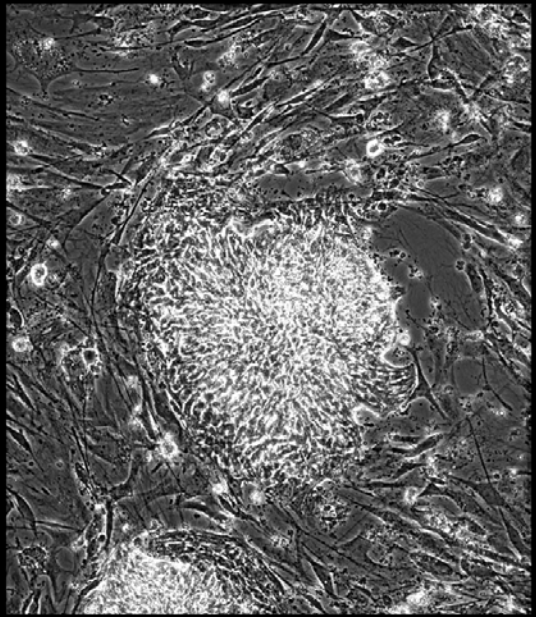

Well = 2, Colony = 106
Rater 1 = 3
Rater 2 = 3

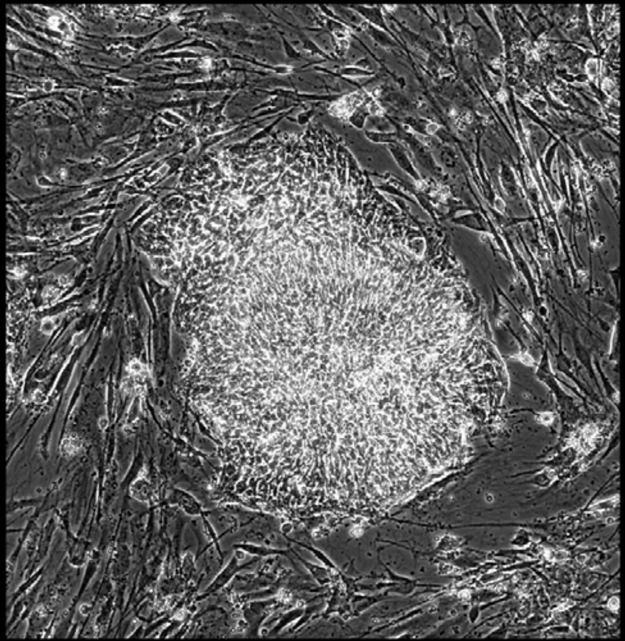

Well = 2, Colony = 107
Rater 1 = 3
Rater 2 = 4

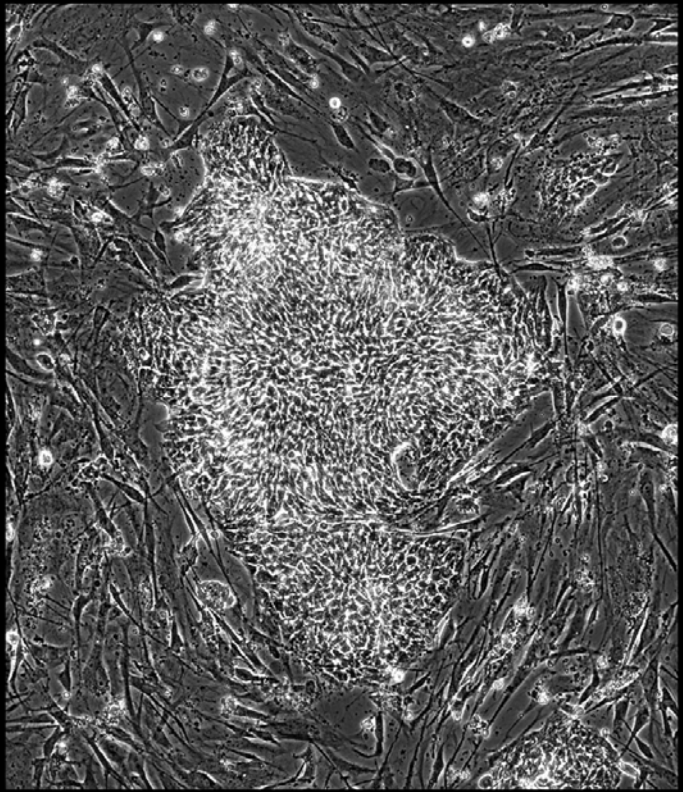

Well = 2, Colony = 108
Rater 1 = 4
Rater 2 = 4

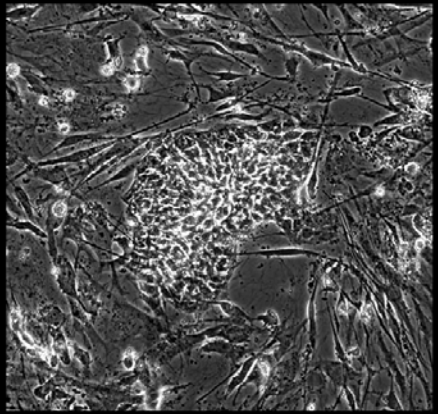

Well = 2, Colony = 109
Rater 1 = 4
Rater 2 = 5

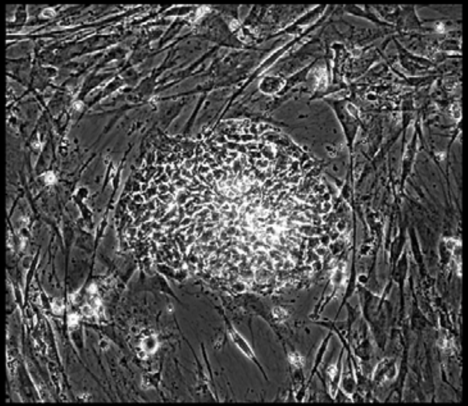

Well = 2, Colony = 110
Rater 1 = 4
Rater 2 = 3

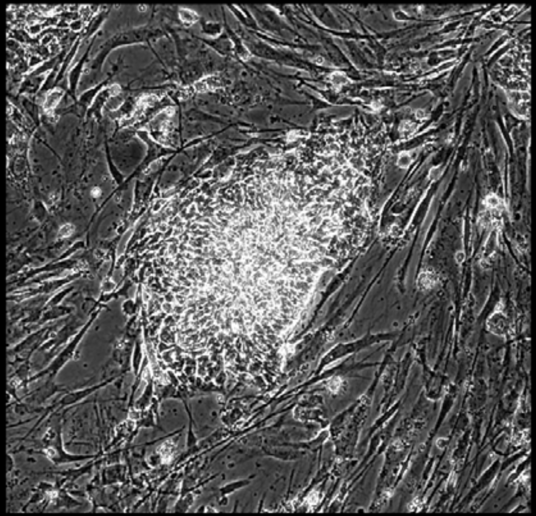

Well = 2, Colony = 111
Rater 1 = 4
Rater 2 = 3

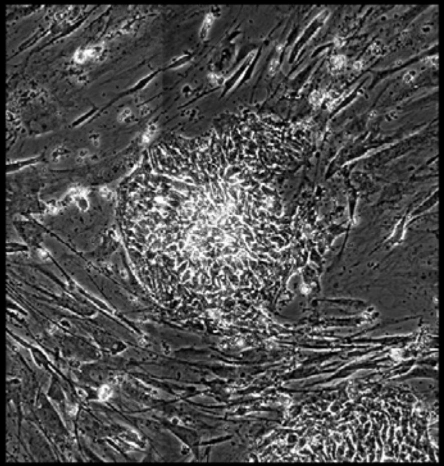

Well = 2, Colony = 112
Rater 1 = 3
Rater 2 = 2

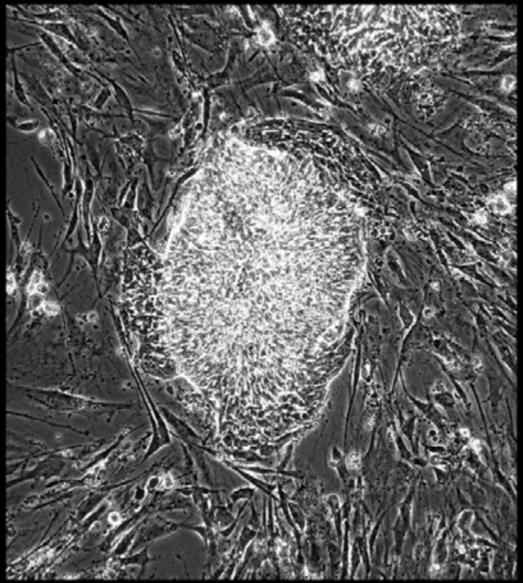

Well = 2, Colony = 113
Rater 1 = 4
Rater 2 = not scored

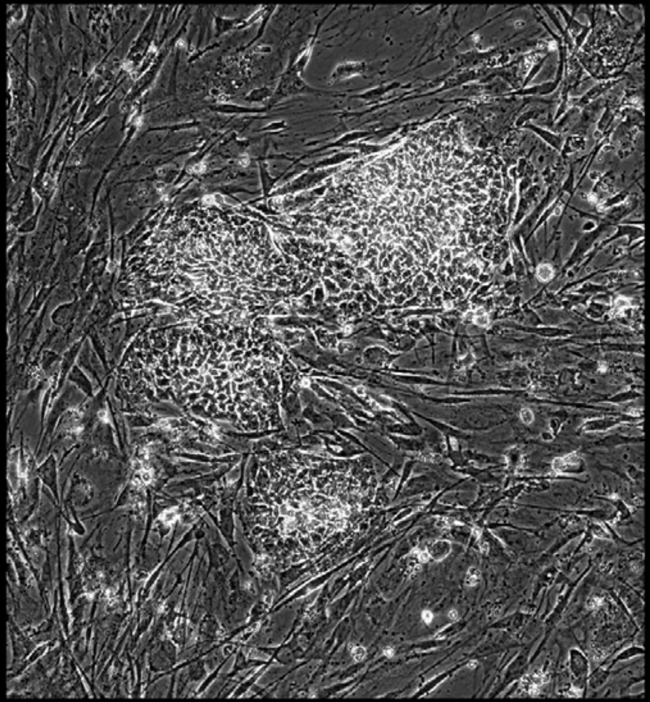

Well = 2, Colony = 114
Rater 1 = 4
Rater 2 = 5

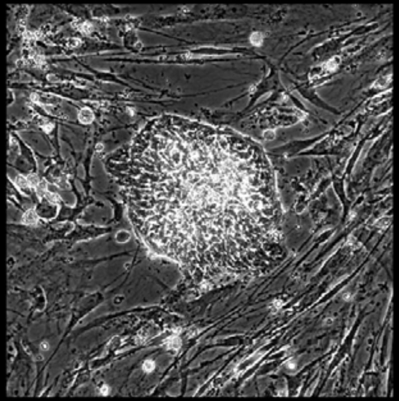

Well = 2, Colony = 115
Rater 1 = 4
Rater 2 = 5

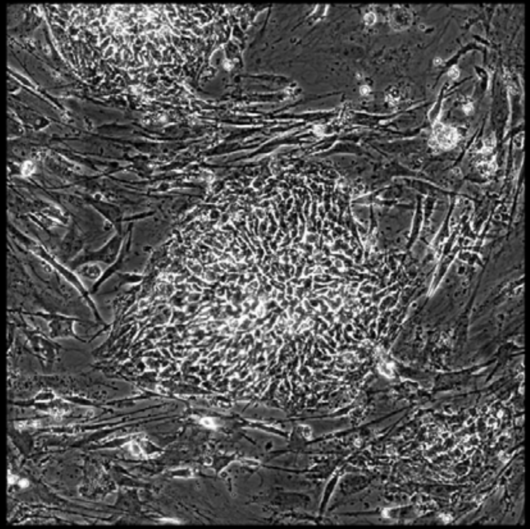

Well = 2, Colony = 116
Rater 1 = 4
Rater 2 = 5

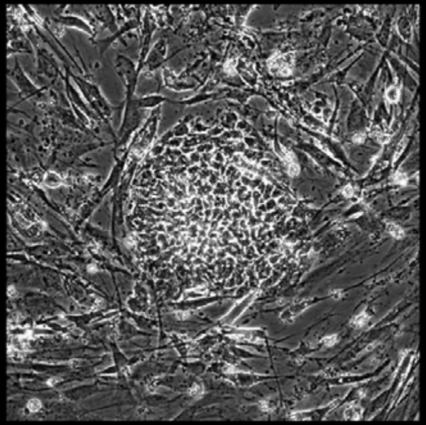

Well = 2, Colony = 117
Rater 1 = 4
Rater 2 = 4

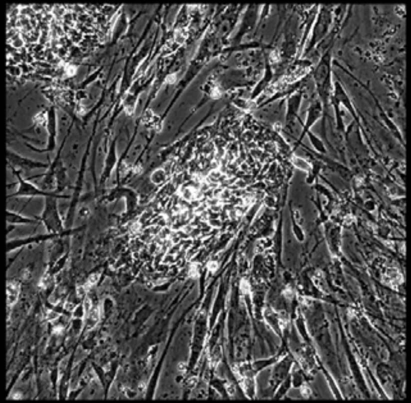

Well = 2, Colony = 118
Rater 1 = 3
Rater 2 = 4

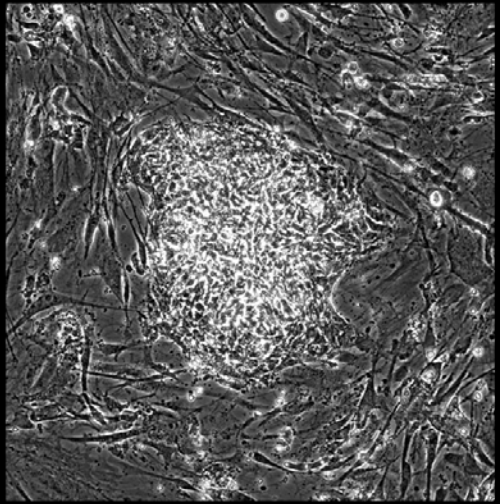

Well = 2, Colony = 119
Rater 1 = 4
Rater 2 = 4

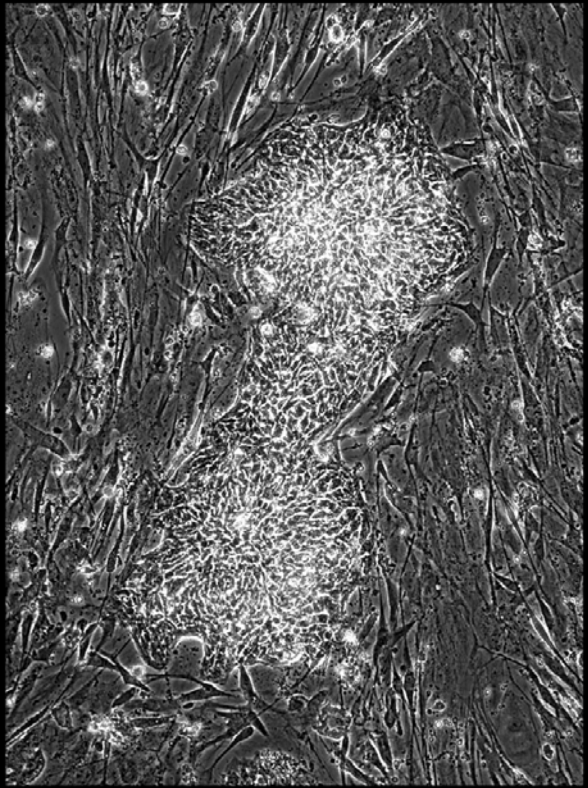

Well = 2, Colony = 120
Rater 1 = 2
Rater 2 = 3

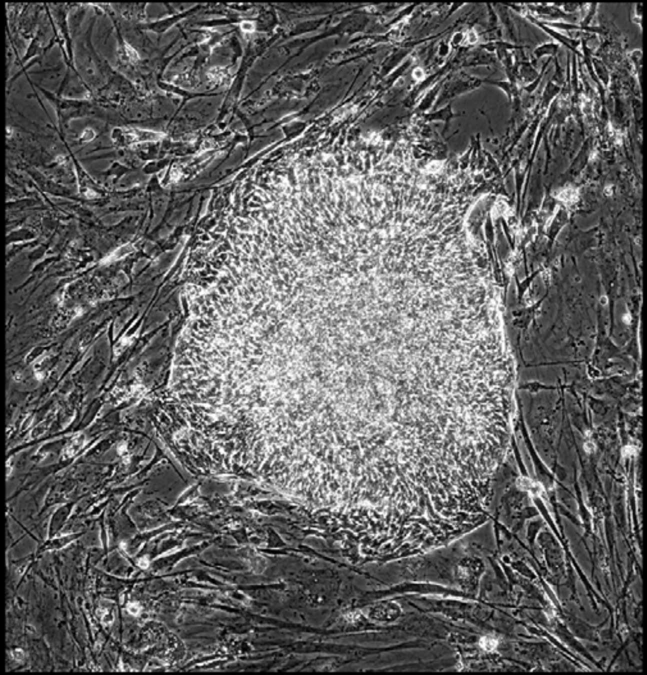

Well = 2, Colony = 121
Rater 1 = 4
Rater 2 = not scored

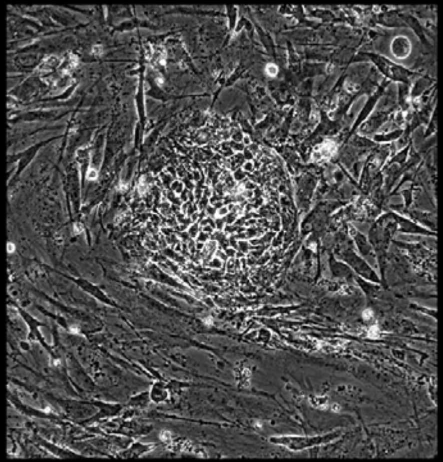

Well = 2, Colony = 122
Rater 1 = 4
Rater 2 = 4

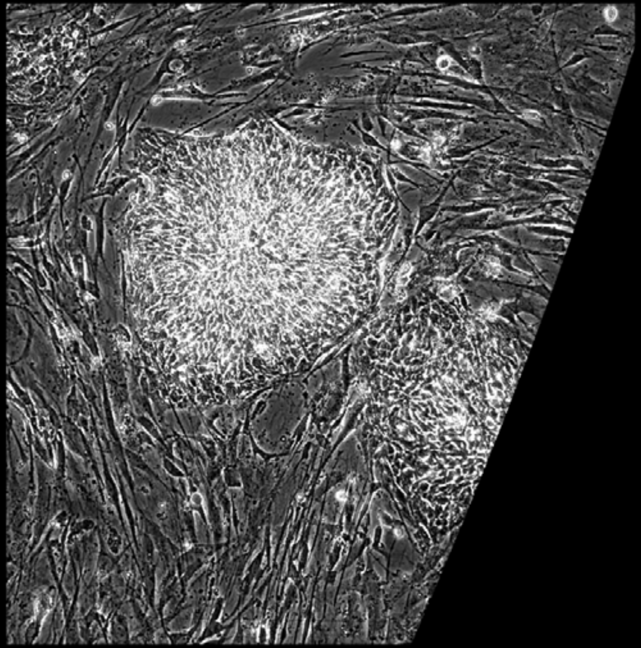

Well = 2, Colony = 123
Rater 1 = 2
Rater 2 = not scored

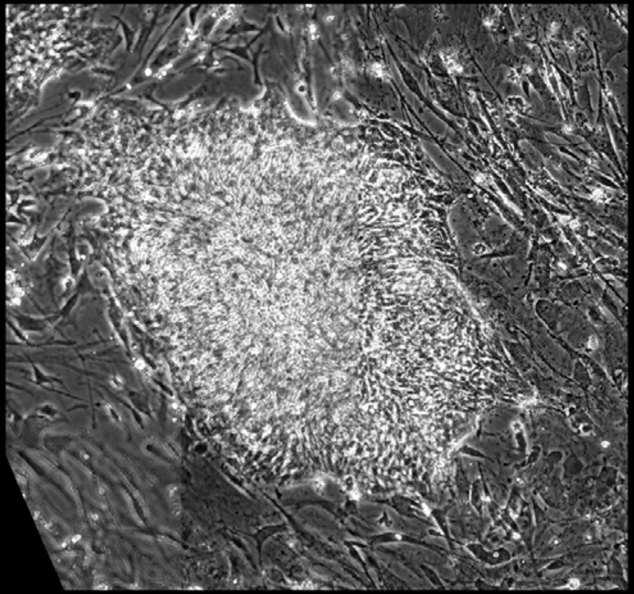

Well = 2, Colony = 124
Rater 1 = 4
Rater 2 = 4

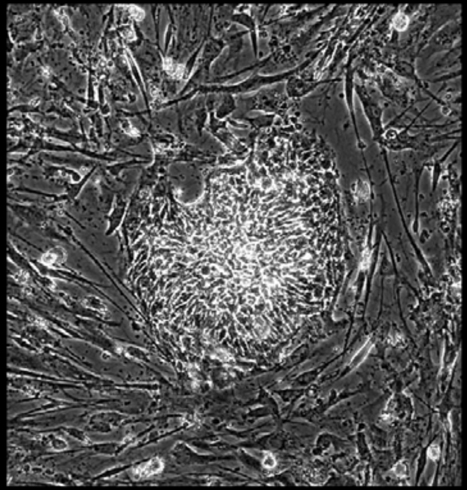

Well = 2, Colony = 125
Rater 1 = 4
Rater 2 = 4

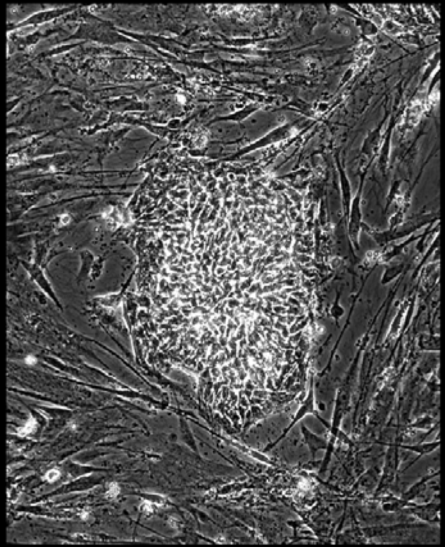

Well = 2, Colony = 126
Rater 1 = 4
Rater 2 = 3

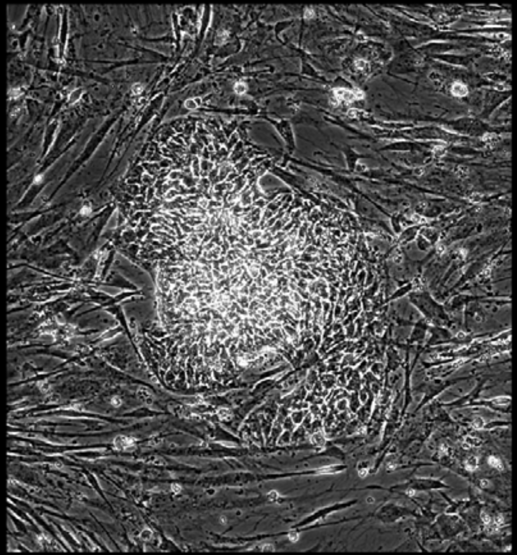

Well = 2, Colony = 127
Rater 1 = 2
Rater 2 = 2

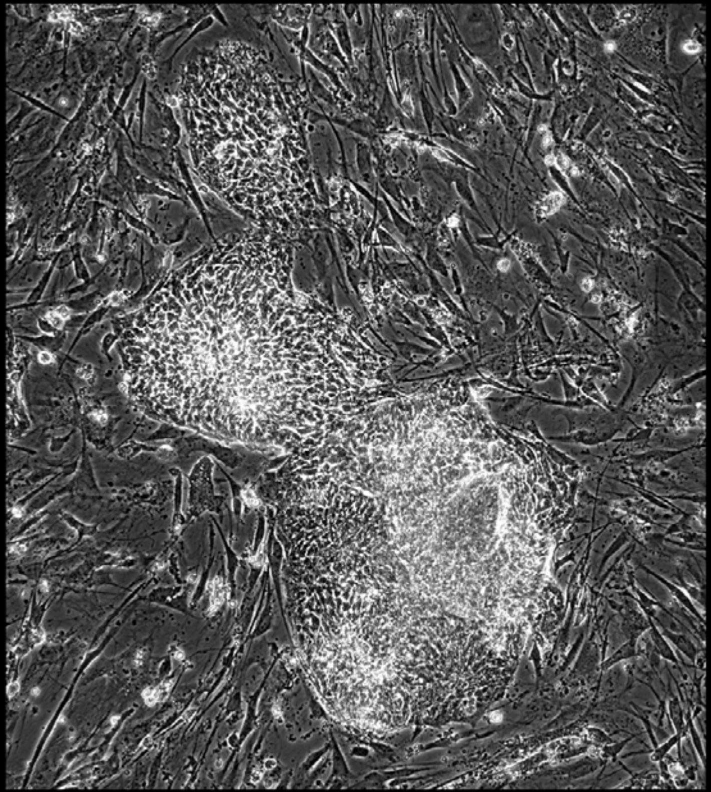

Well = 2, Colony = 128
Rater 1 = 3
Rater 2 = 2

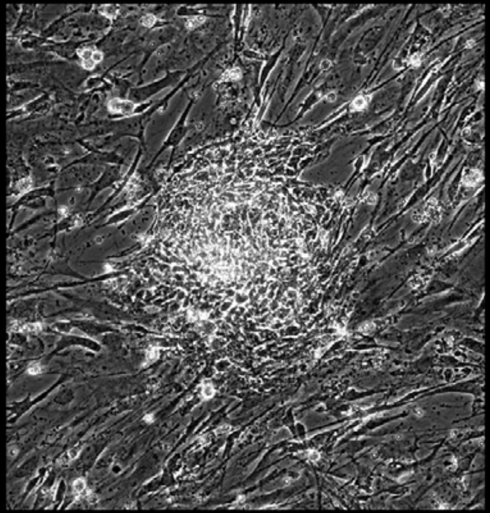

Well = 2, Colony = 129
Rater 1 = 3
Rater 2 = 4

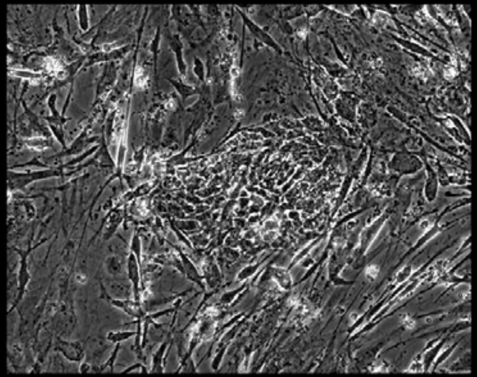

Well = 2, Colony = 130
Rater 1 = 3
Rater 2 = 4

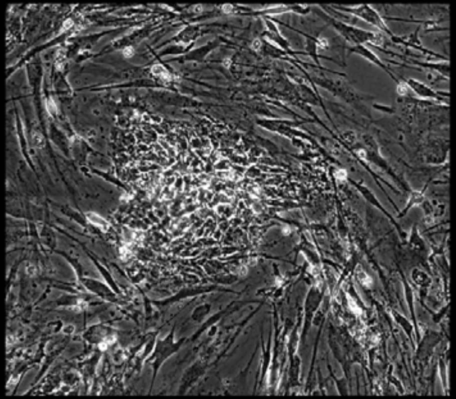

Well = 2, Colony = 131
Rater 1 = 3
Rater 2 = not scored

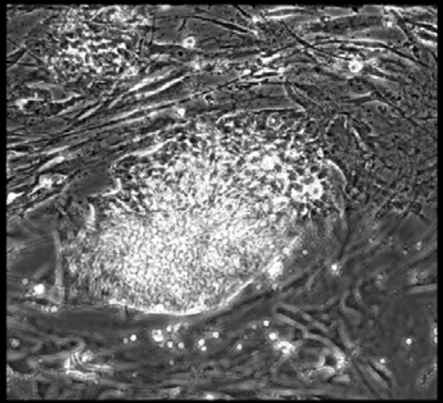

Well = 2, Colony = 132
Rater 1 = 3
Rater 2 = 2

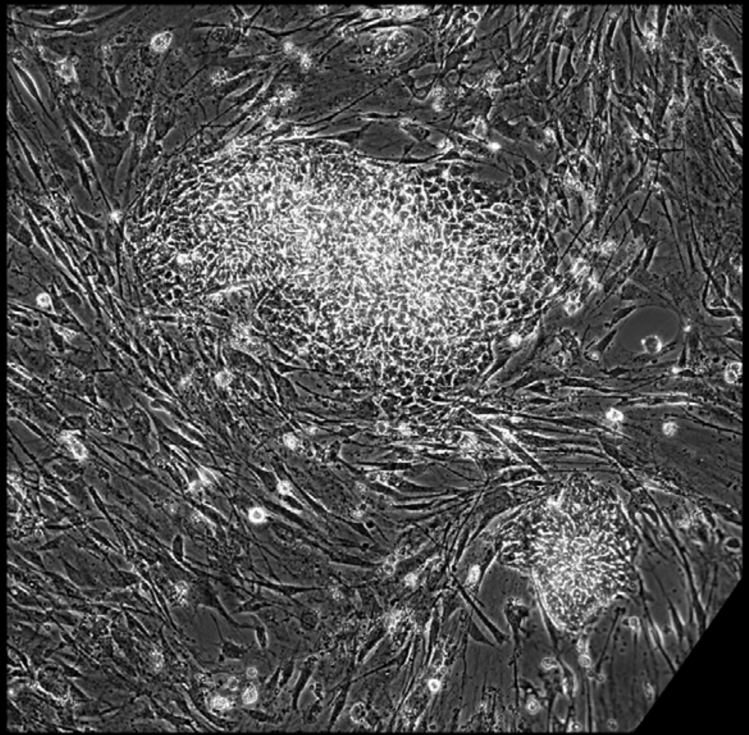

Well = 2, Colony = 133
Rater 1 = 4
Rater 2 = 3

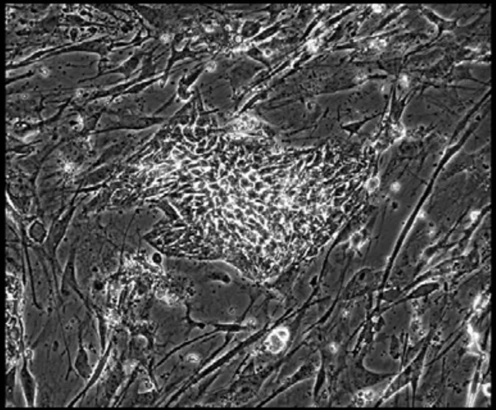

Well = 2, Colony = 134
Rater 1 = 4
Rater 2 = 5

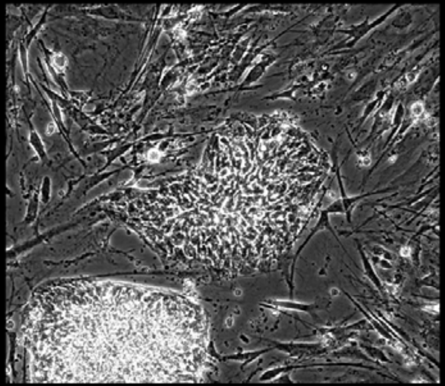

Well = 2, Colony = 135
Rater 1 = 4
Rater 2 = 4

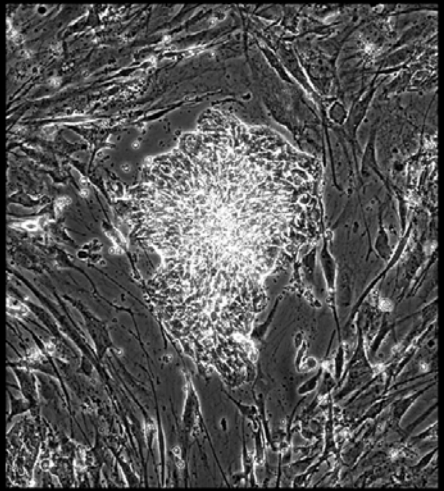

Well = 2, Colony = 136
Rater 1 = 2
Rater 2 = 2

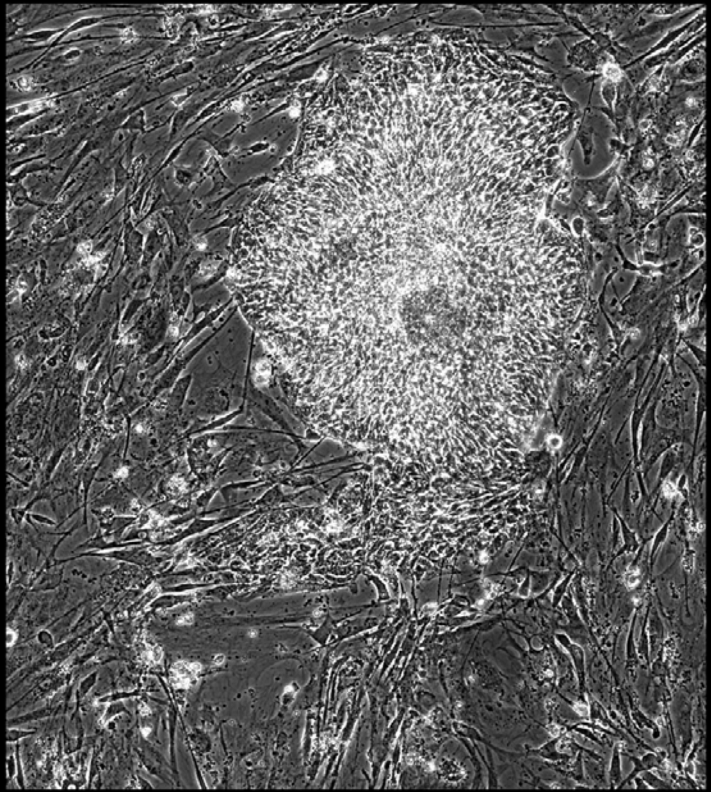

Well = 2, Colony = 137
Rater 1 = 4
Rater 2 = 3

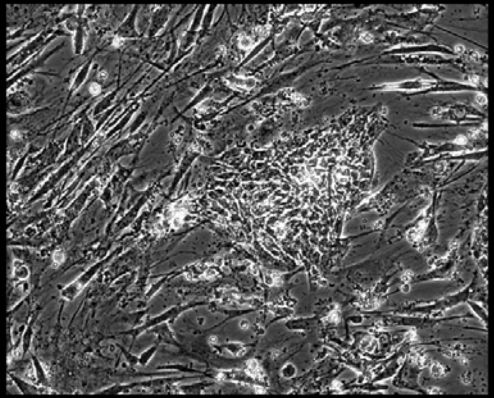

Well = 2, Colony = 138
Rater 1 = 3
Rater 2 = 4

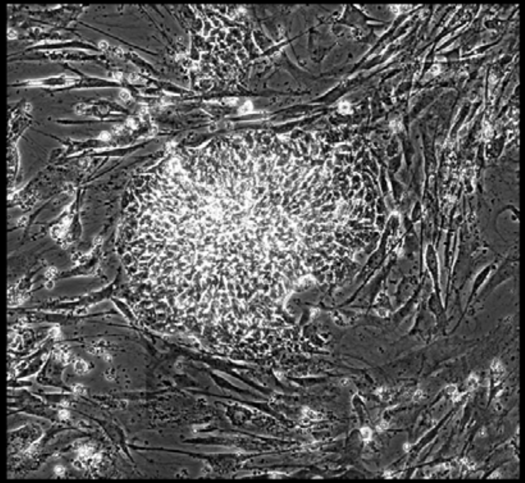

Well = 2, Colony = 139
Rater 1 = 4
Rater 2 = 5

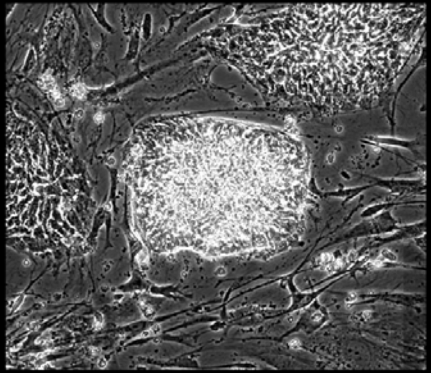

Well = 2, Colony = 140
Rater 1 = 5
Rater 2 = 5

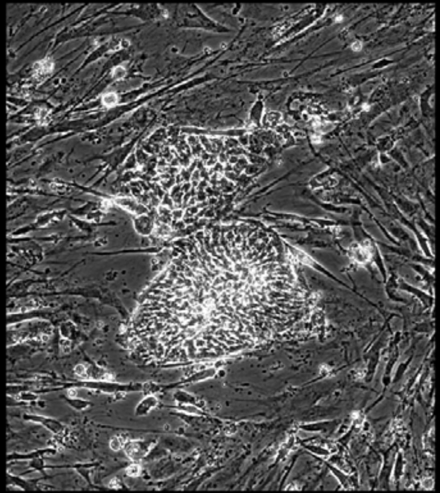

Well = 2, Colony = 141
Rater 1 = 2
Rater 2 = 2

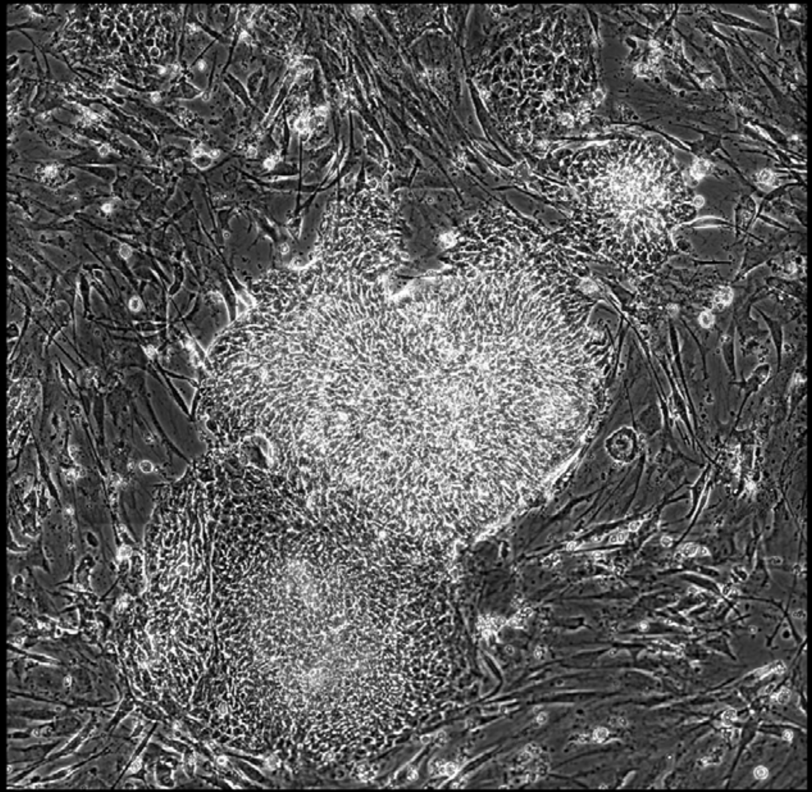

Well = 2, Colony = 142
Rater 1 = 3
Rater 2 = 2

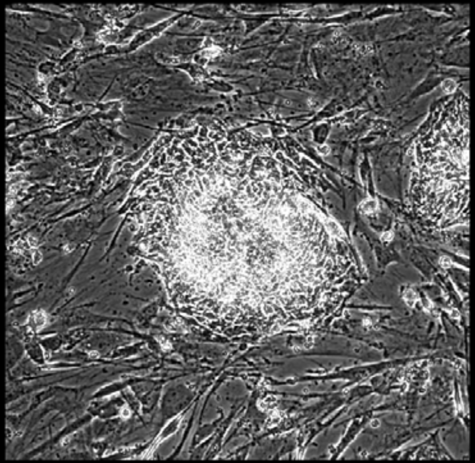

Well = 2, Colony = 144
Rater 1 = 3
Rater 2 = 2

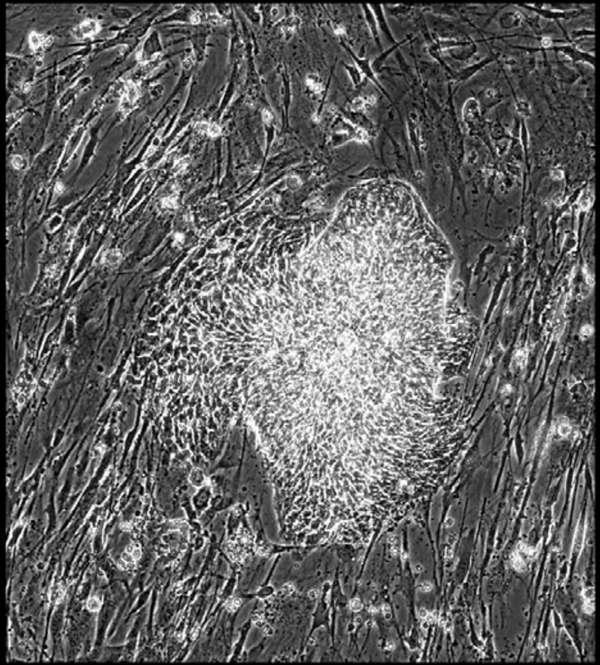

Well = 2, Colony = 145
Rater 1 = 4
Rater 2 = 4

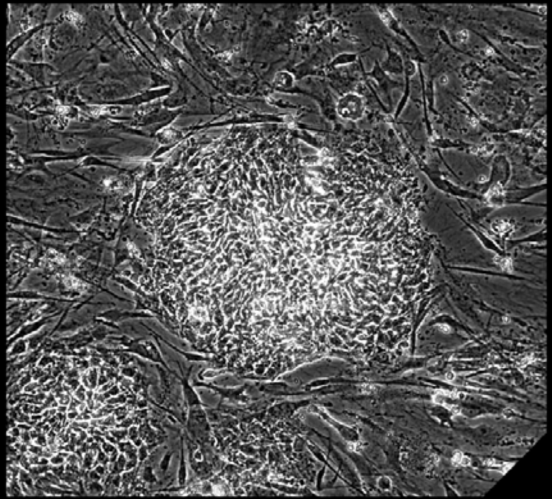

Well = 2, Colony = 146
Rater 1 = 4
Rater 2 = 4

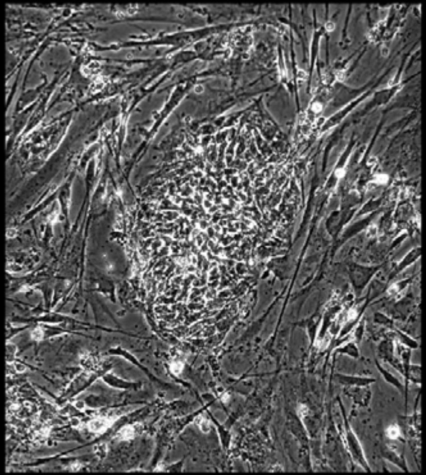

Well = 2, Colony = 147
Rater 1 = 4
Rater 2 = 4

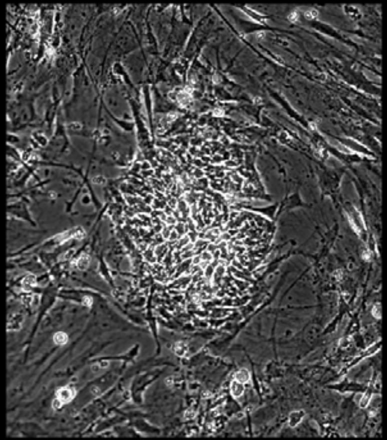

Well = 2, Colony = 148
Rater 1 = 3
Rater 2 = 3

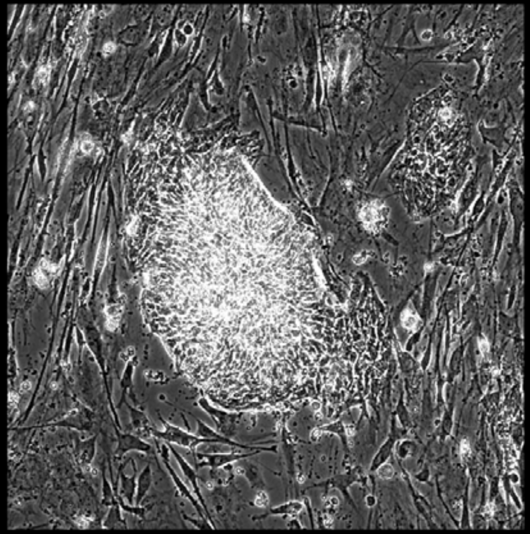

Well = 2, Colony = 149
Rater 1 = 3
Rater 2 = 4

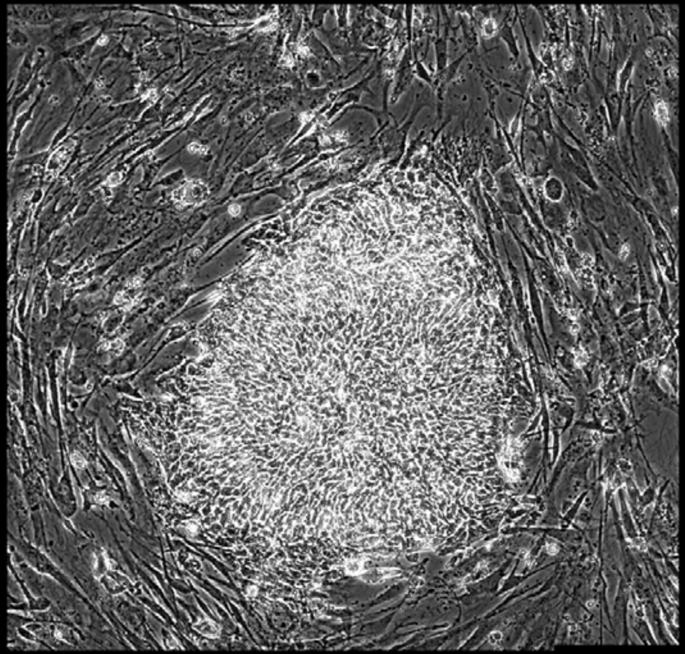

Well = 2, Colony = 150
Rater 1 = 4
Rater 2 = 4

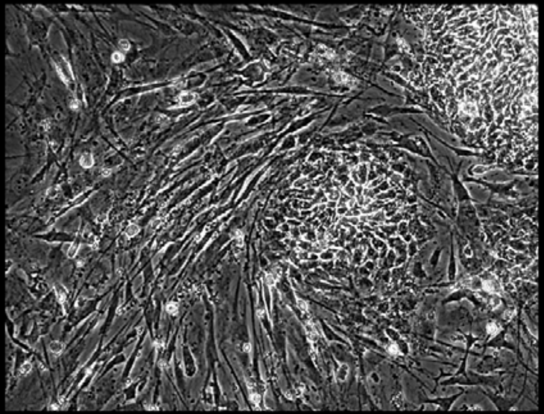

Well = 2, Colony = 151
Rater 1 = 3
Rater 2 = 2

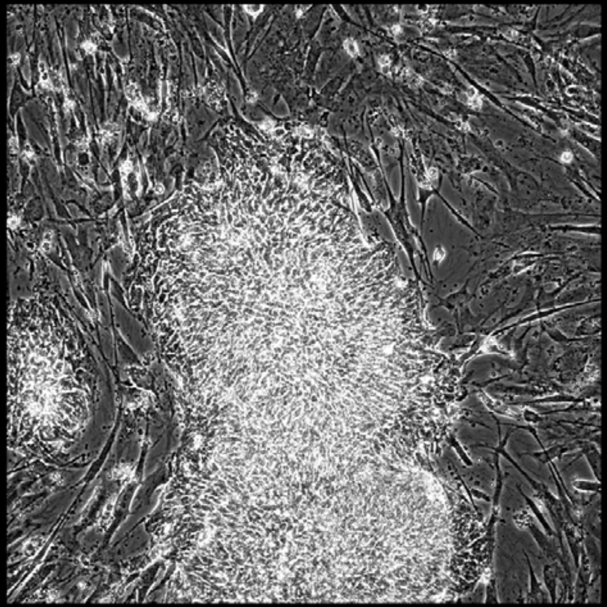

Well = 2, Colony = 152
Rater 1 = 3
Rater 2 = 4

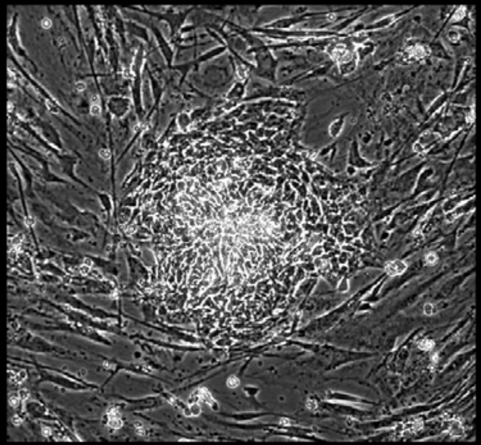

Well = 2, Colony = 153
Rater 1 = 4
Rater 2 = not scored

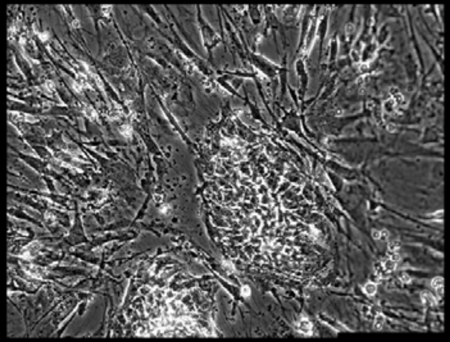

Well = 3, Colony = 1
Rater 1 = 4
Rater 2 = 4

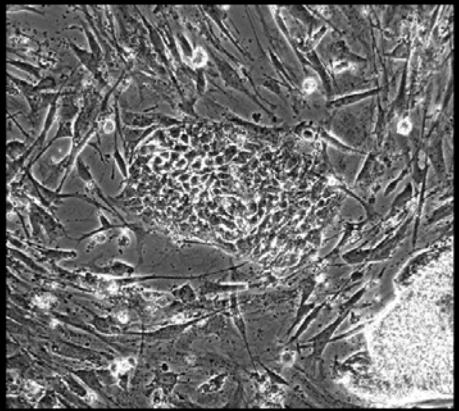

Well = 3, Colony = 2
Rater 1 = 2
Rater 2 = 2

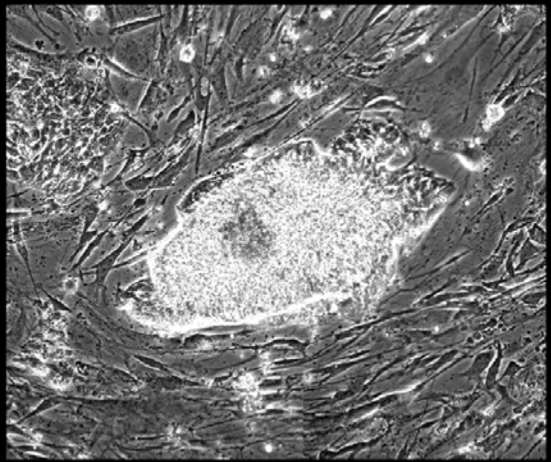

Well = 3, Colony = 3
Rater 1 = 2
Rater 2 = 2

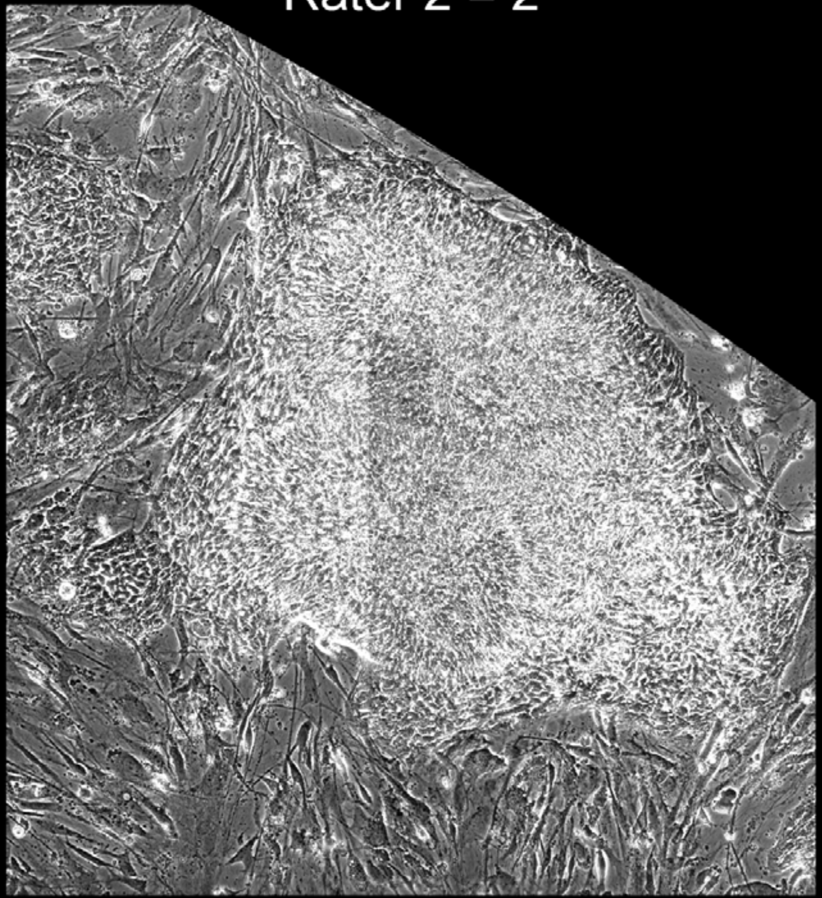

Well = 3, Colony = 4
Rater 1 = 4
Rater 2 = 4

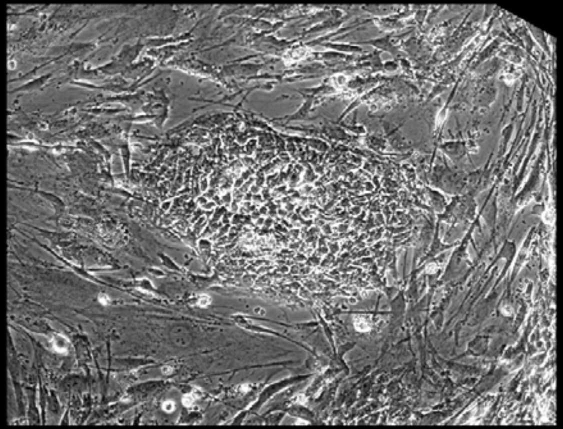

Well = 3, Colony = 5
Rater 1 = 4
Rater 2 = 4

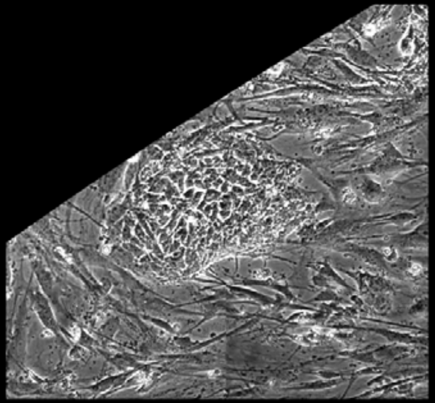

Well = 3, Colony = 6
Rater 1 = 2
Rater 2 = 3

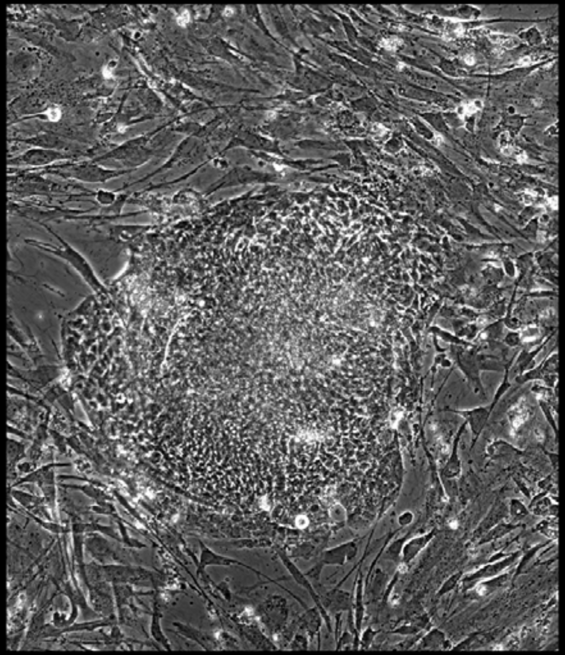

Well = 3, Colony = 7
Rater 1 = 4
Rater 2 = 5

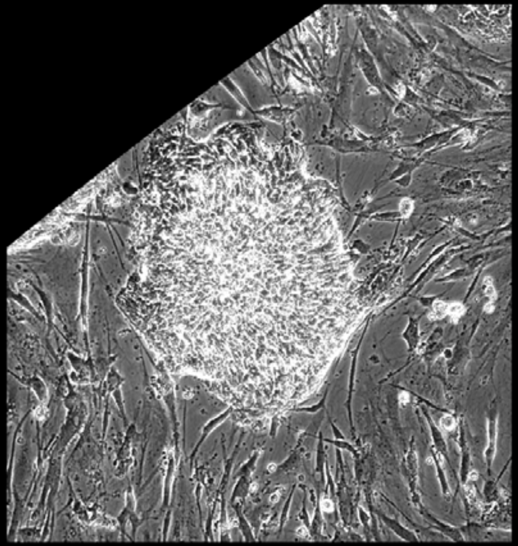

Well = 3, Colony = 8
Rater 1 = 3
Rater 2 = 3

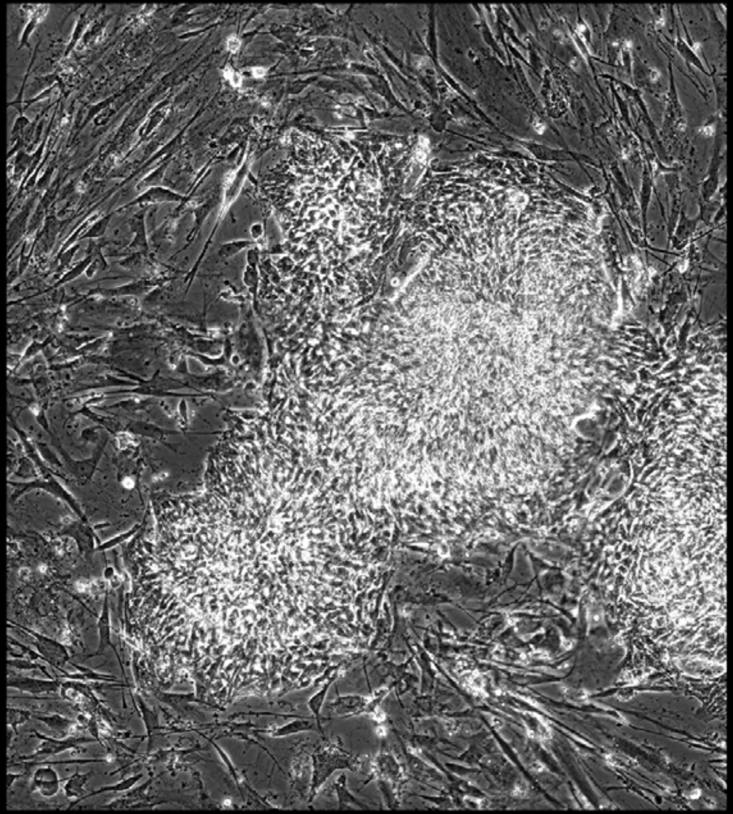

Well = 3, Colony = 9
Rater 1 = 3
Rater 2 = 4

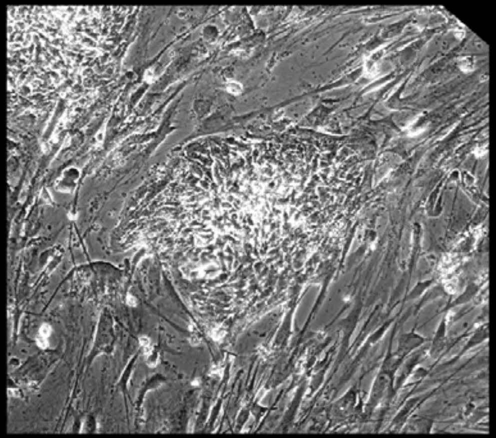

Well = 3, Colony = 10
Rater 1 = 3
Rater 2 = 3

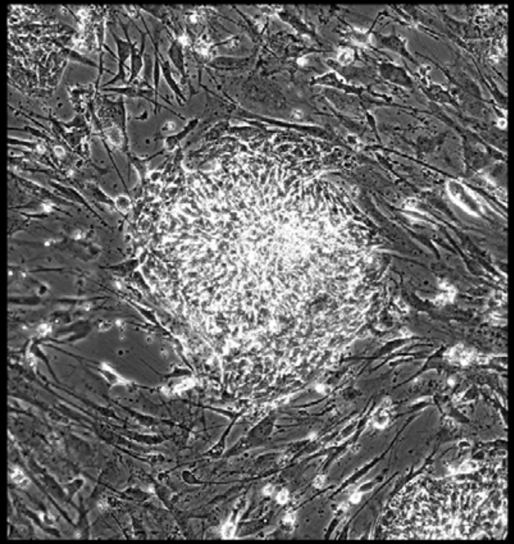

Well = 3, Colony = 11
Rater 1 = 3
Rater 2 = 2

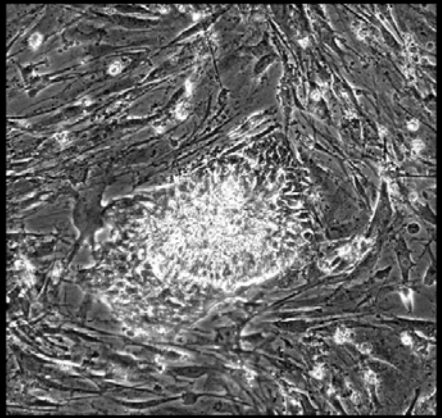

Well = 3, Colony = 12
Rater 1 = 3
Rater 2 = 3

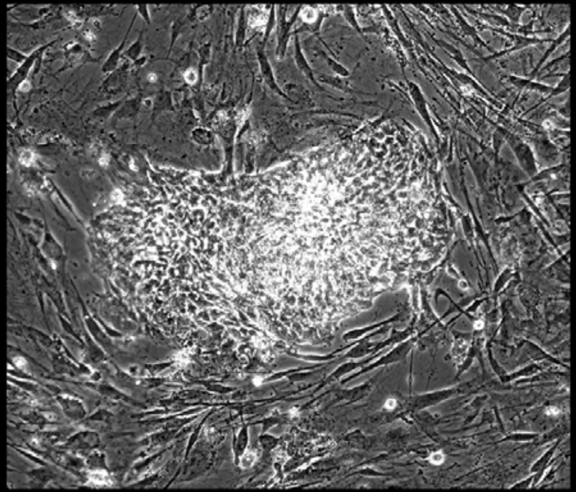

Well = 3, Colony = 13
Rater 1 = 2
Rater 2 = 2

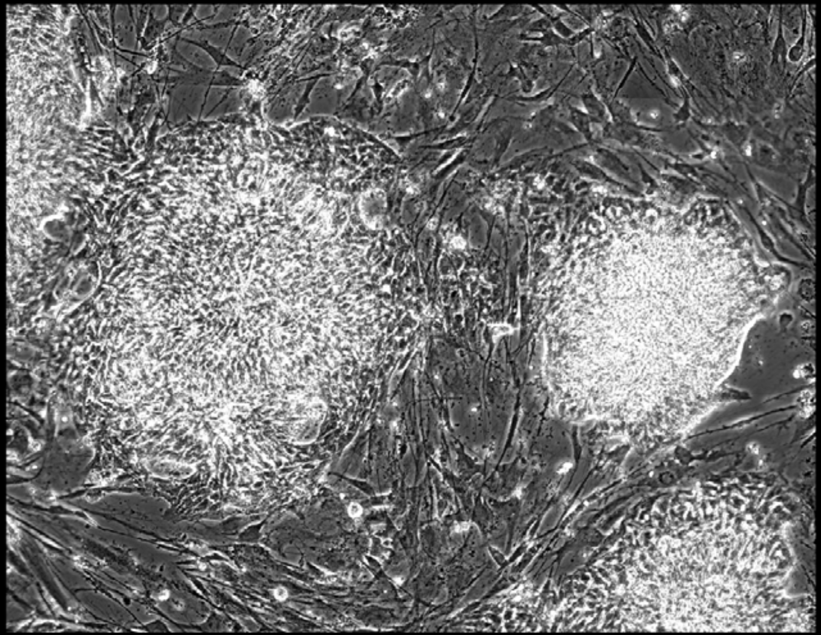

Well = 3, Colony = 14
Rater 1 = 3
Rater 2 = 4

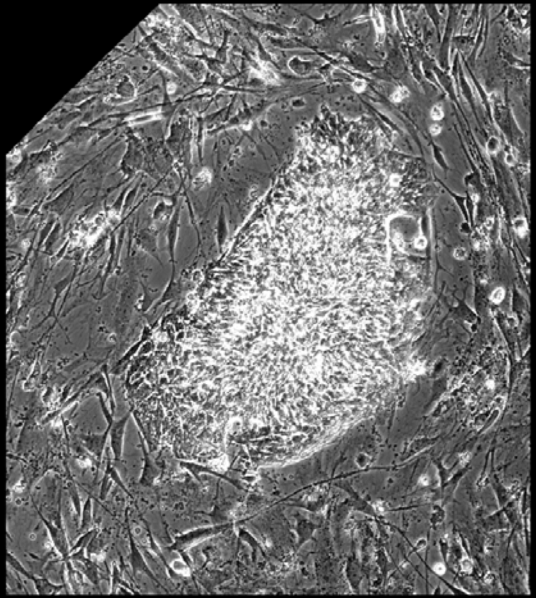

Well = 3, Colony = 15
Rater 1 = 4
Rater 2 = 4

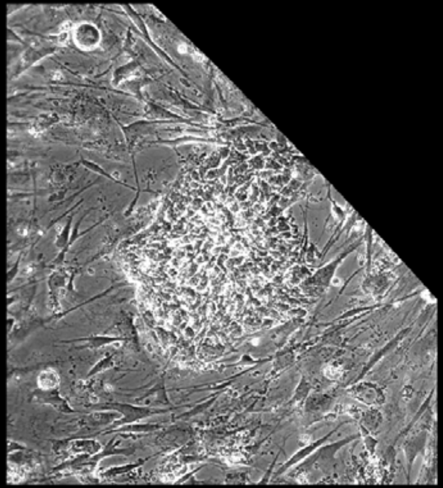

Well = 3, Colony = 16
Rater 1 = 3
Rater 2 = 5

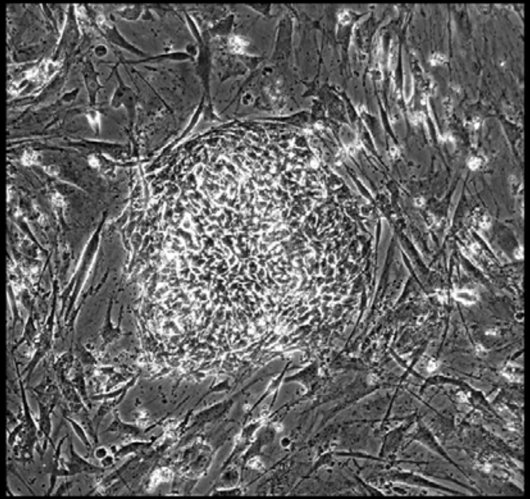

Well = 3, Colony = 17
Rater 1 = 3
Rater 2 = not scored

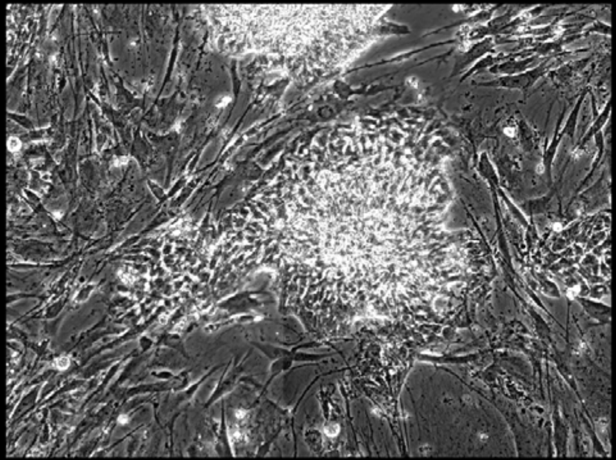

Well = 3, Colony = 18
Rater 1 = 4
Rater 2 = 3

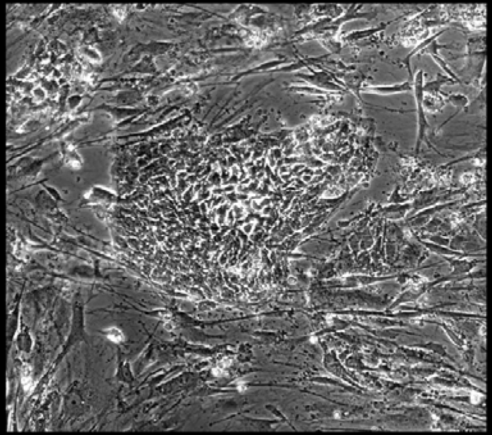

Well = 3, Colony = 19
Rater 1 = 2
Rater 2 = 2

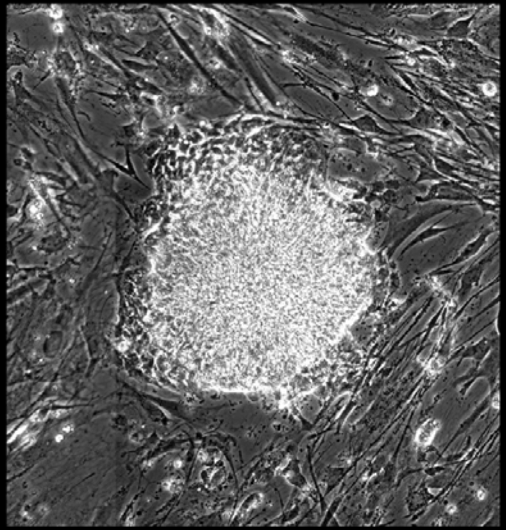

Well = 3, Colony = 20
Rater 1 = 3
Rater 2 = 5

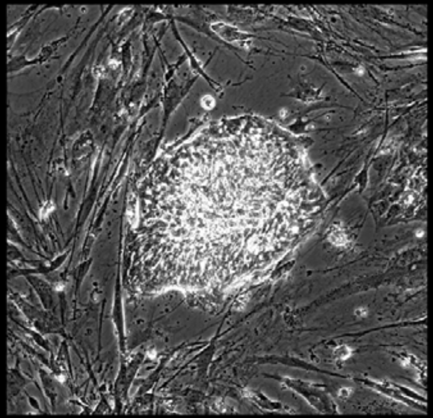

Well = 3, Colony = 21
Rater 1 = 4
Rater 2 = 4

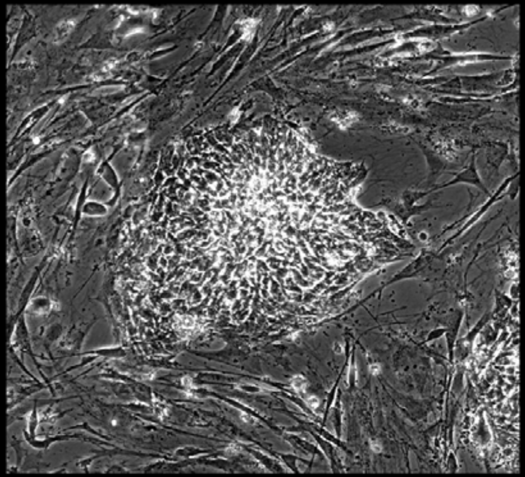

Well = 3, Colony = 22
Rater 1 = 4
Rater 2 = 4

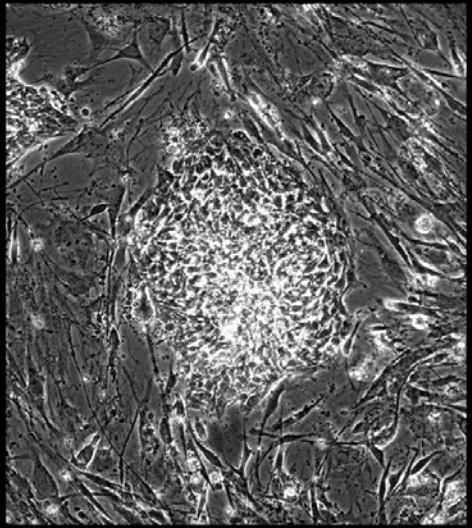

Well = 3, Colony = 23
Rater 1 = 4
Rater 2 = 4

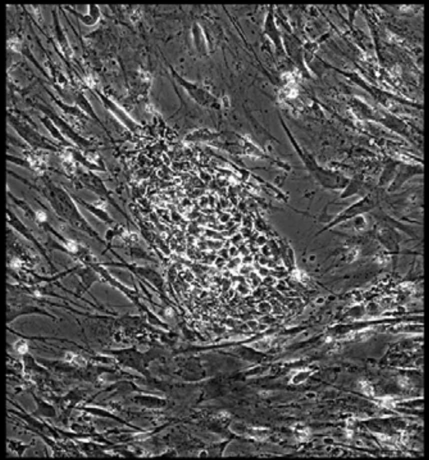

Well = 3, Colony = 24
Rater 1 = 5
Rater 2 = 5

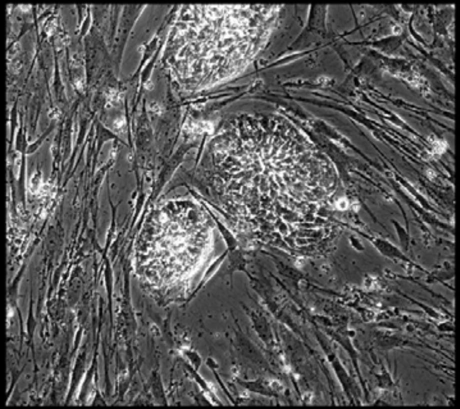

Well = 3, Colony = 25
Rater 1 = 4
Rater 2 = 4

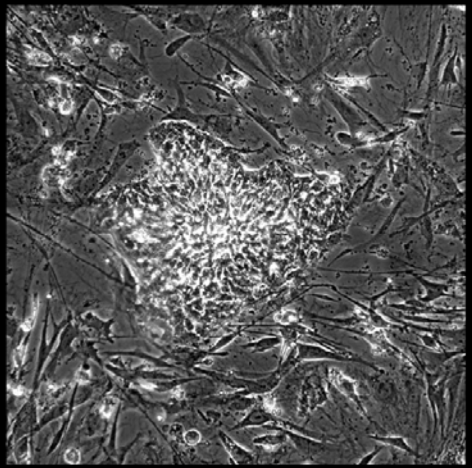

Well = 3, Colony = 26
Rater 1 = 2
Rater 2 = 4

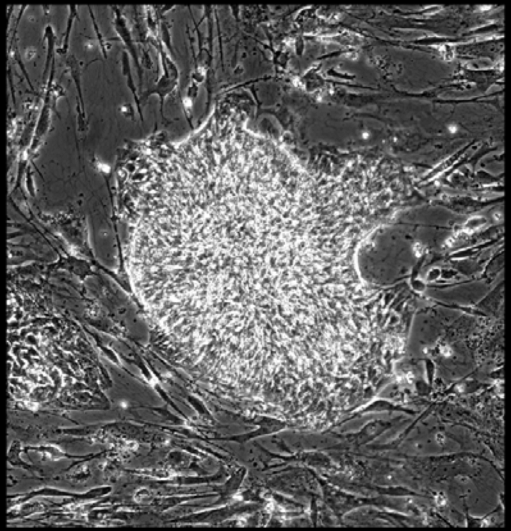

Well = 3, Colony = 27
Rater 1 = 3
Rater 2 = 3

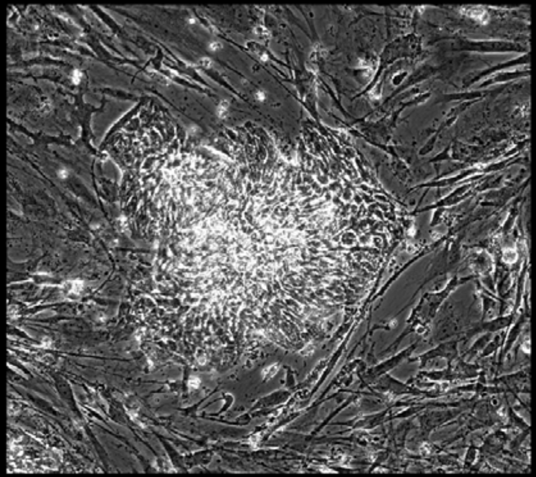

Well = 3, Colony = 28
Rater 1 = 5
Rater 2 = 4

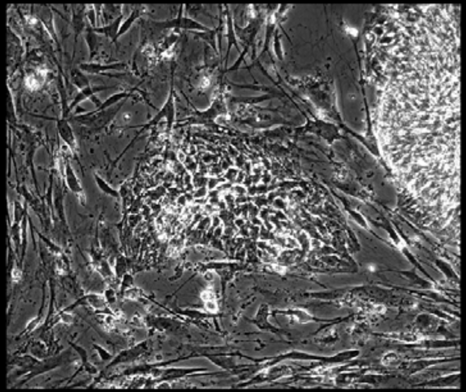

Well = 3, Colony = 29
Rater 1 = 3
Rater 2 = 3

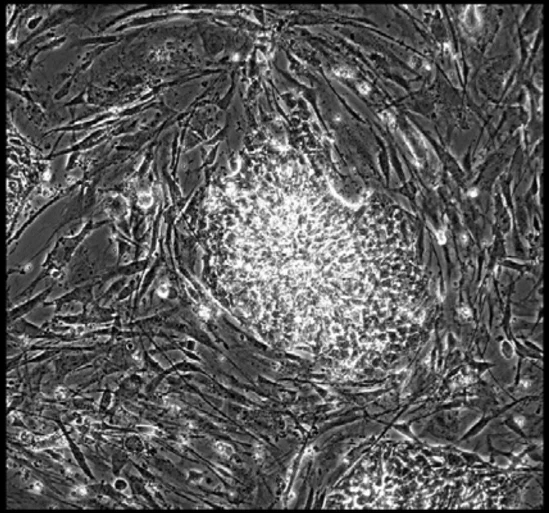

Well = 3, Colony = 30
Rater 1 = 5
Rater 2 = 5

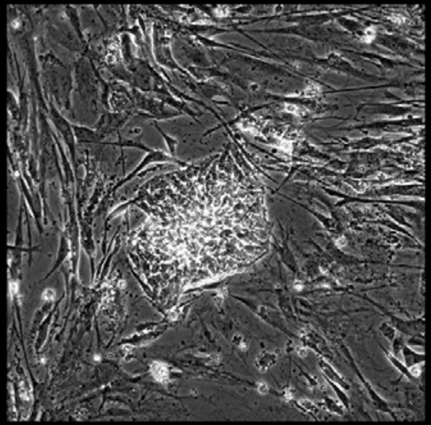

Well = 3, Colony = 31
Rater 1 = 3
Rater 2 = 4

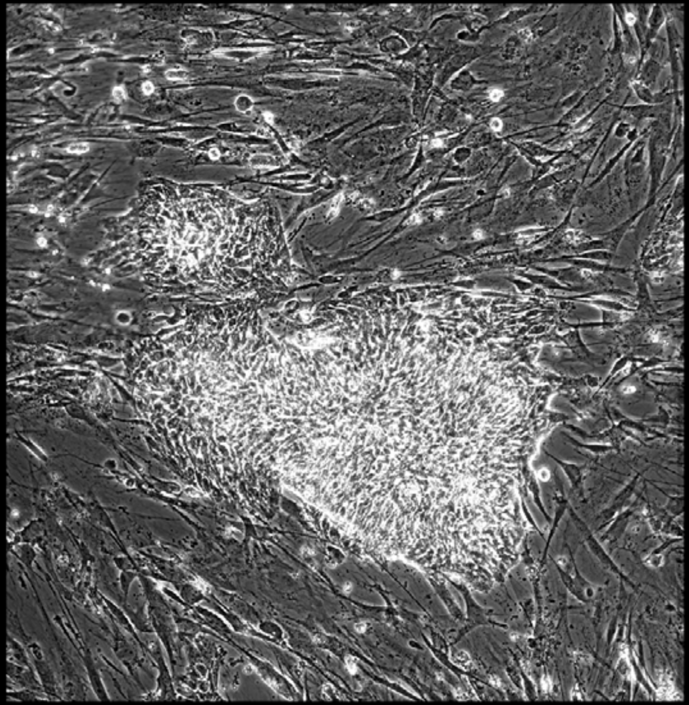

Well = 3, Colony = 32
Rater 1 = 4
Rater 2 = 4

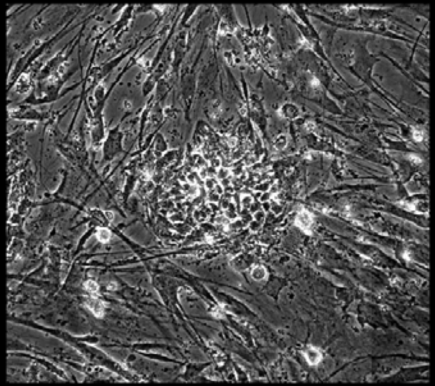

Well = 3, Colony = 33
Rater 1 = 4
Rater 2 = 5

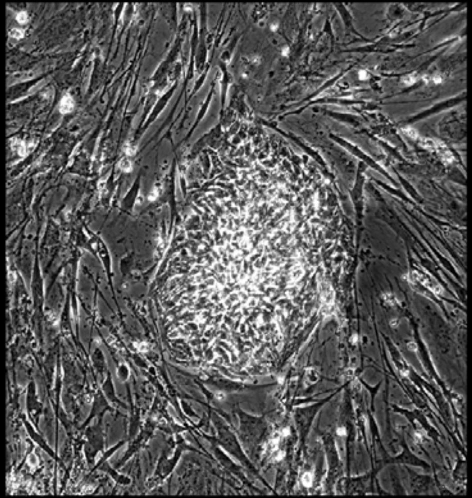

Well = 3, Colony = 34
Rater 1 = 3
Rater 2 = 4

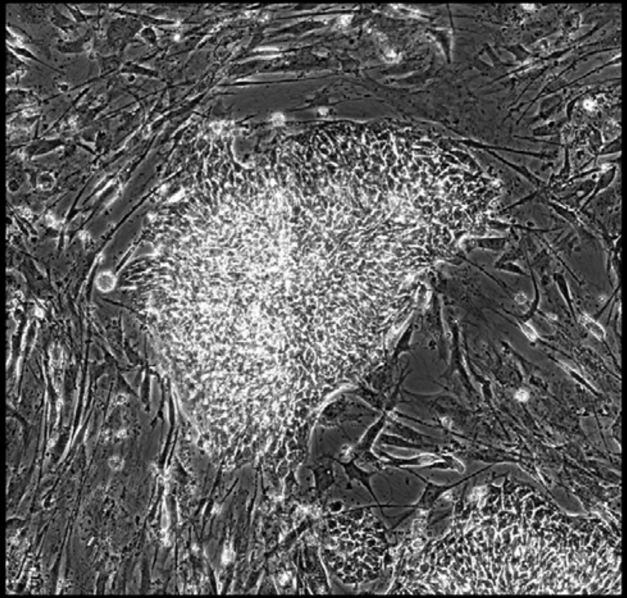

Well = 3, Colony = 35
Rater 1 = 3
Rater 2 = 5

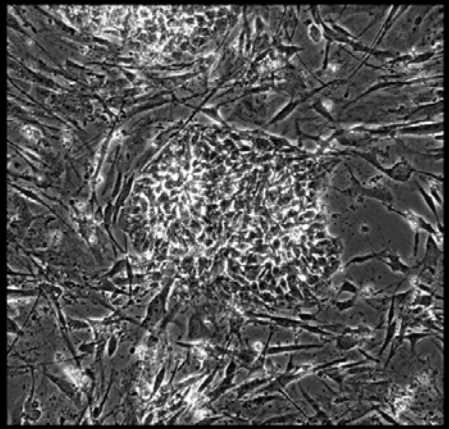

Well = 3, Colony = 36
Rater 1 = 2
Rater 2 = 5

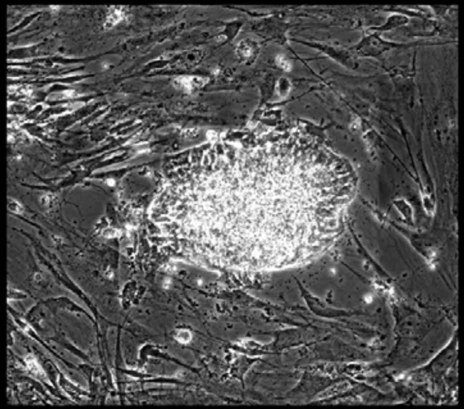

Well = 3, Colony = 37
Rater 1 = 3
Rater 2 = 3

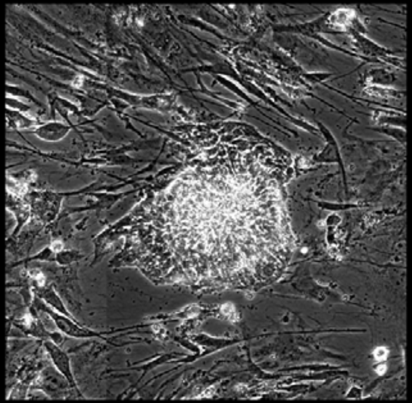

Well = 3, Colony = 38
Rater 1 = 2
Rater 2 = 5

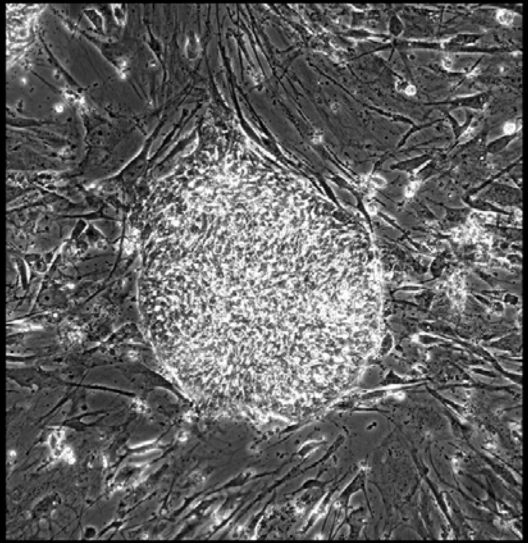

Well = 3, Colony = 39
Rater 1 = 3
Rater 2 = 4

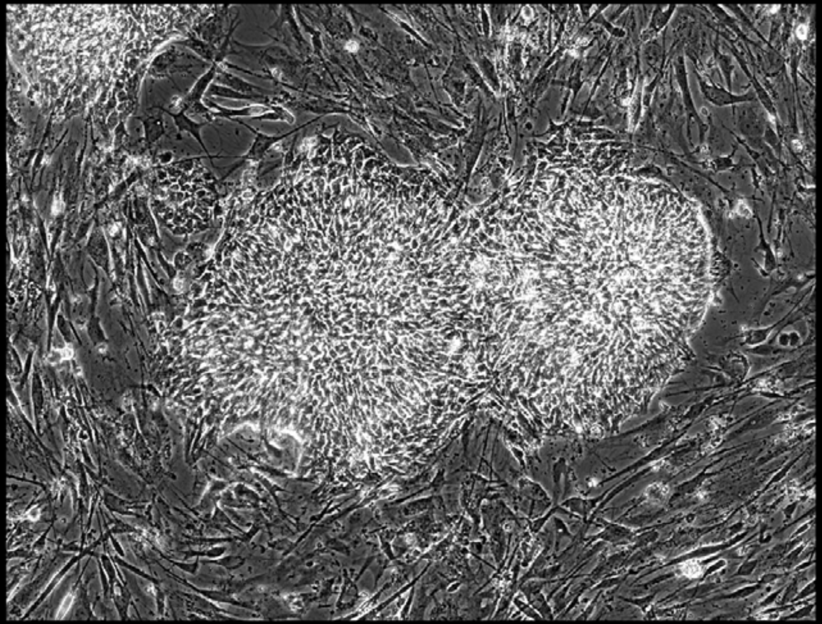

Well = 3, Colony = 40
Rater 1 = 5
Rater 2 = 5

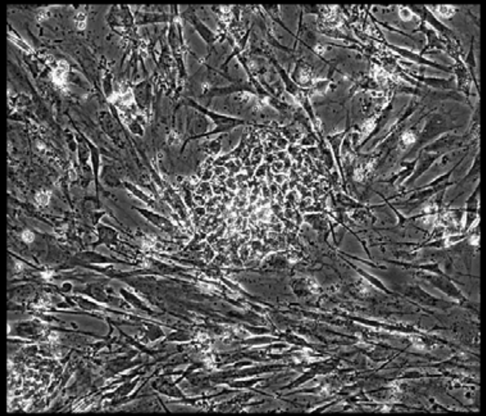

Well = 3, Colony = 41
Rater 1 = 3
Rater 2 = 4

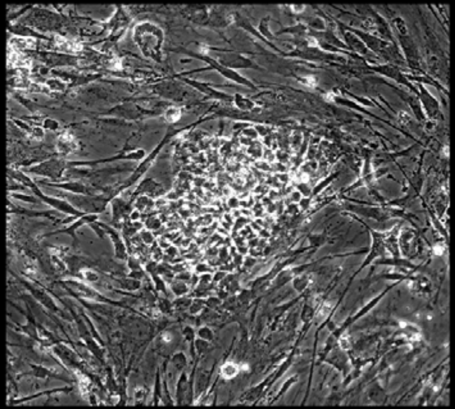

Well = 3, Colony = 42
Rater 1 = 4
Rater 2 = 5

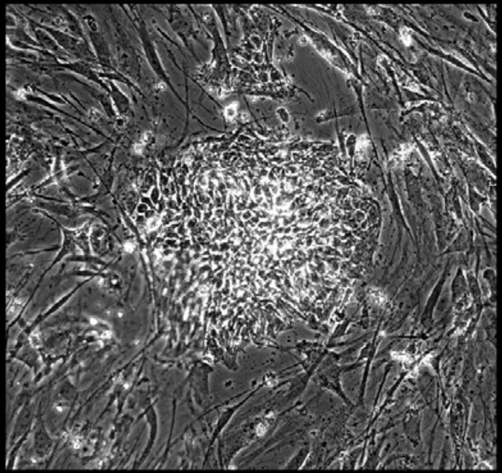

Well = 3, Colony = 43
Rater 1 = 4
Rater 2 = 5

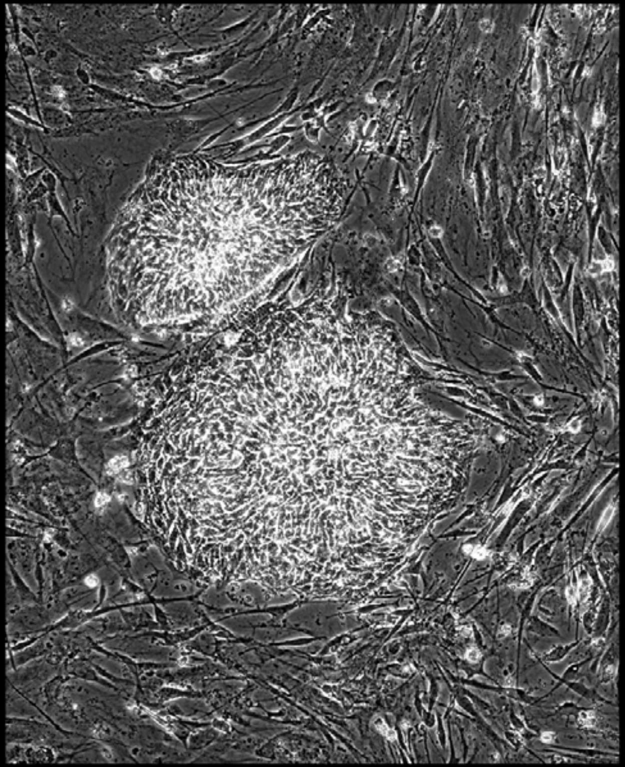

Well = 3, Colony = 44
Rater 1 = not scored
Rater 2 = 4

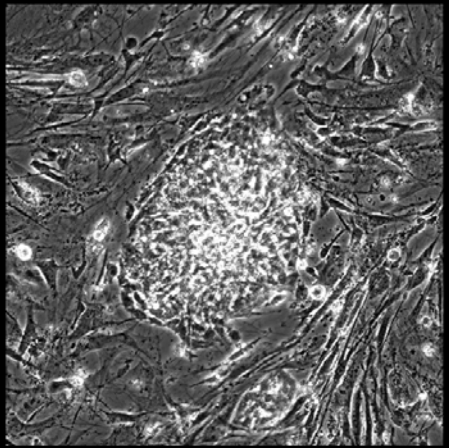

Well = 3, Colony = 45
Rater 1 = 3
Rater 2 = 3

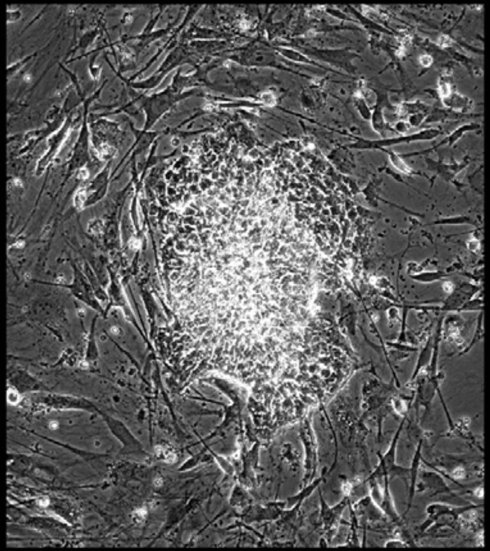

Well = 3, Colony = 46
Rater 1 = 2
Rater 2 = 3

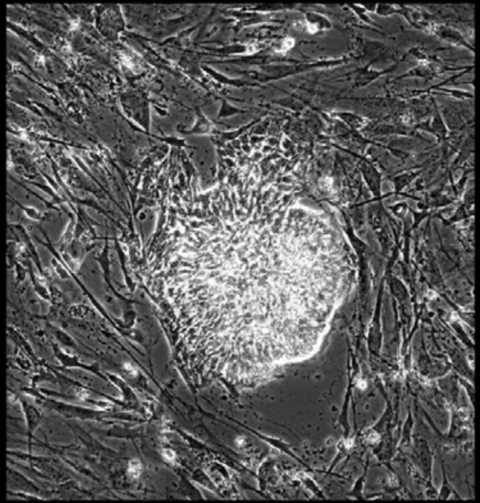

Well = 3, Colony = 47
Rater 1 = 3
Rater 2 = 3

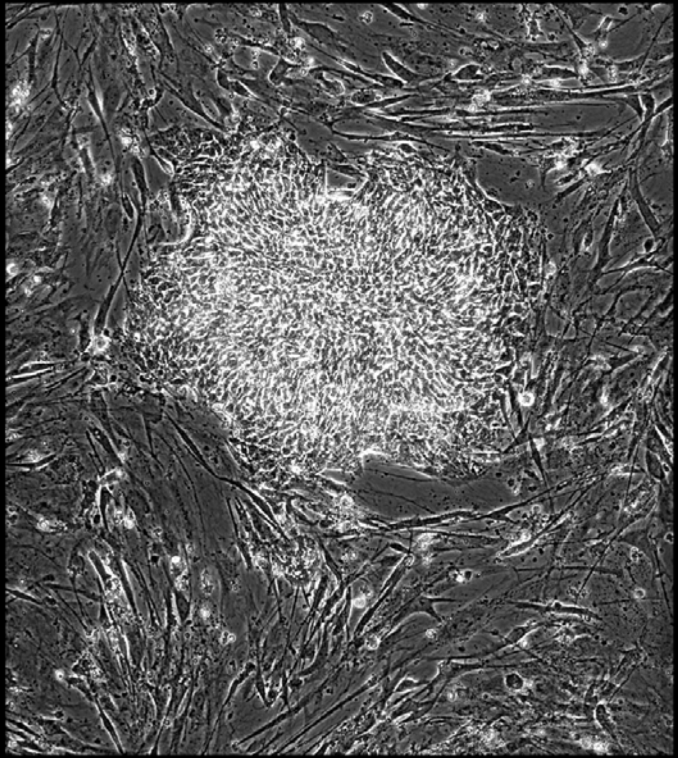

Well = 3, Colony = 48
Rater 1 = 5
Rater 2 = 4

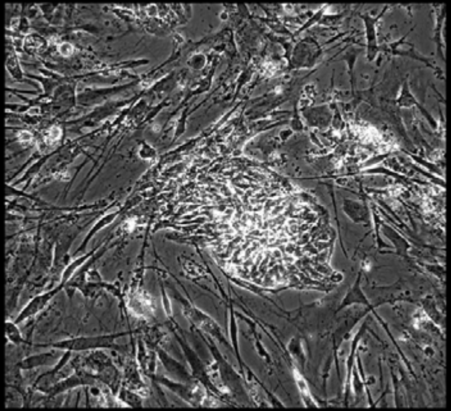

Well = 3, Colony = 49
Rater 1 = 1
Rater 2 = 1

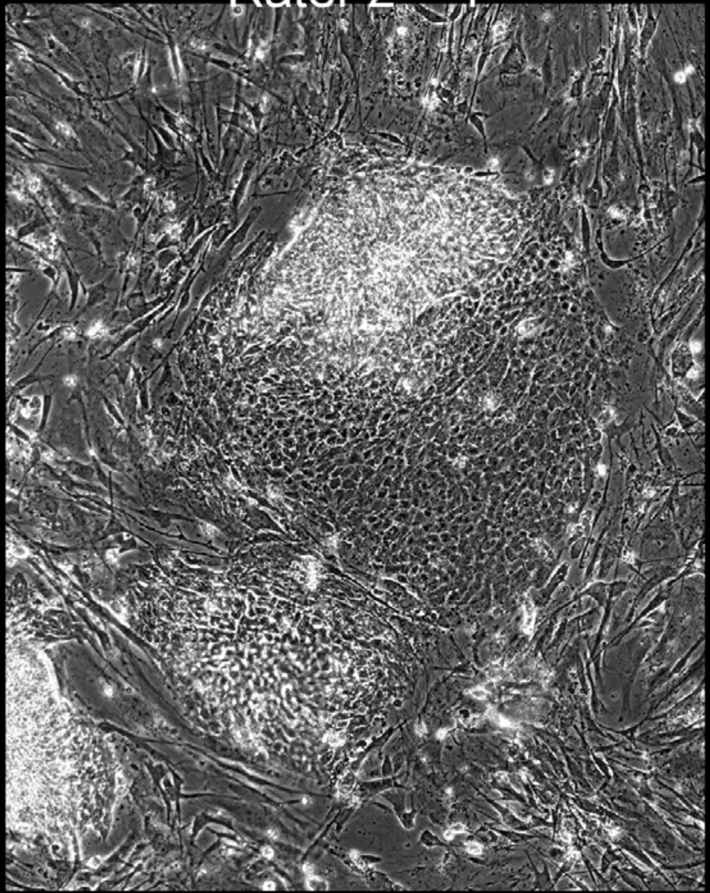

Well = 3, Colony = 50
Rater 1 = 3
Rater 2 = 3

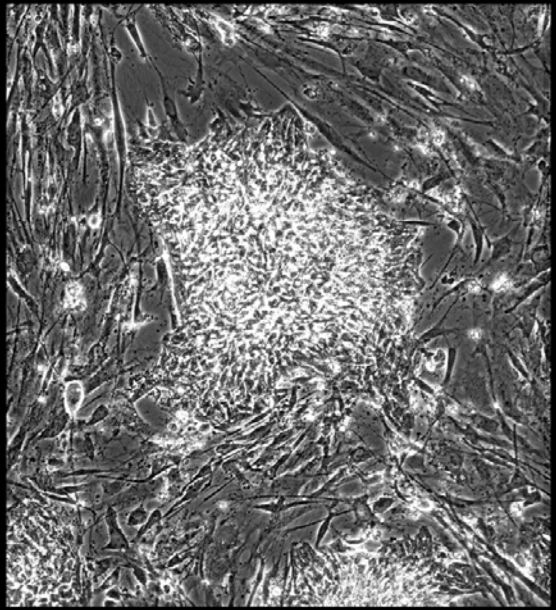

Well = 3, Colony = 51
Rater 1 = 5
Rater 2 = 4

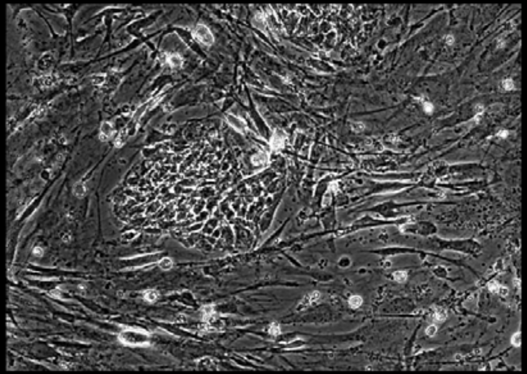

Well = 3, Colony = 52
Rater 1 = 5
Rater 2 = 4

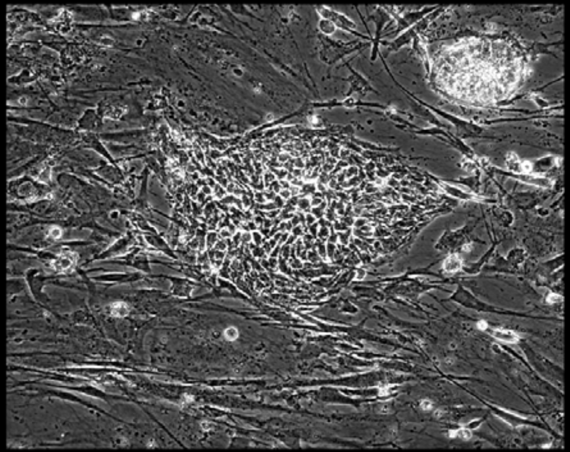

Well = 3, Colony = 53
Rater 1 = 2
Rater 2 = 3

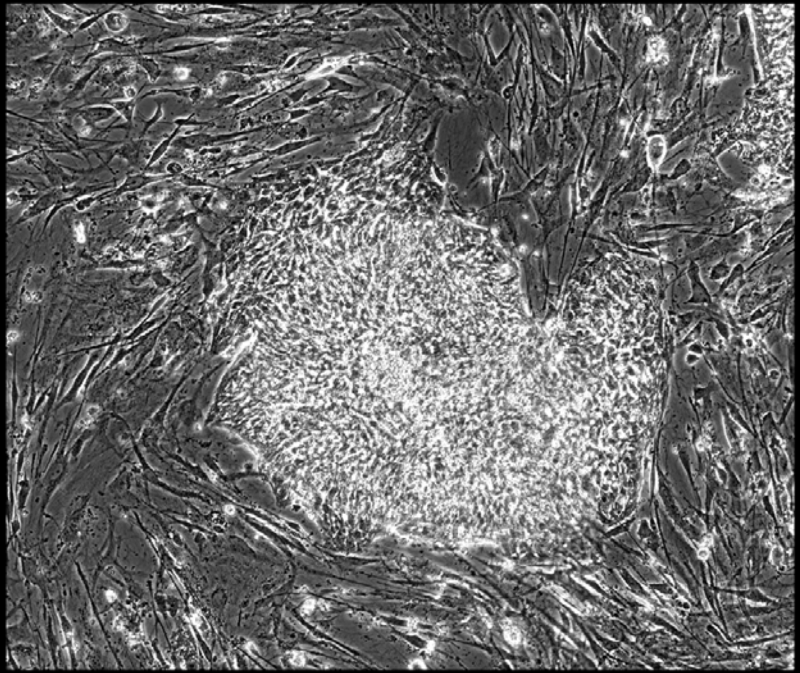

Well = 3, Colony = 54
Rater 1 = 3
Rater 2 = 3

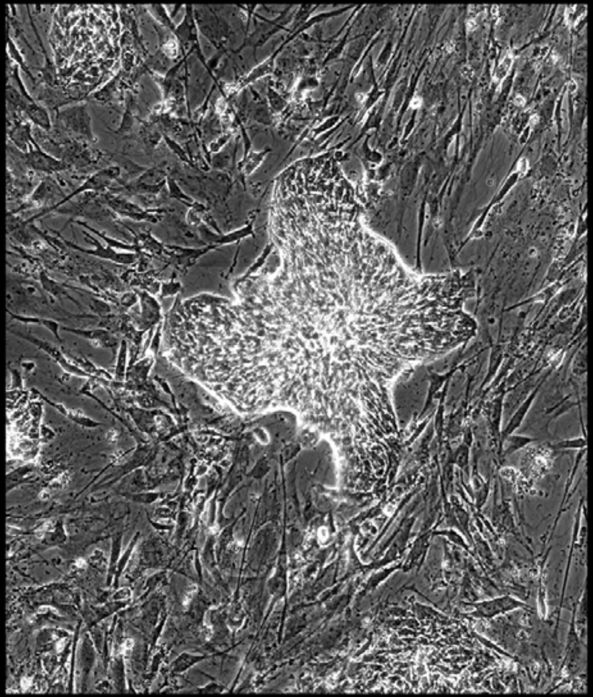

Well = 3, Colony = 55
Rater 1 = 5
Rater 2 = 4

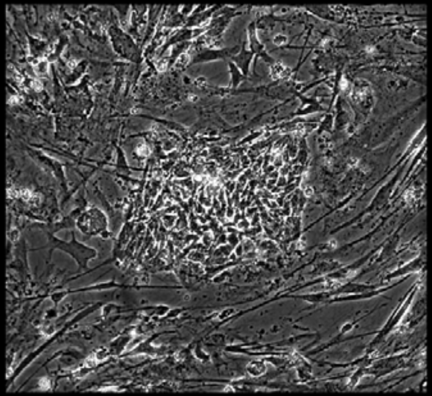

Well = 3, Colony = 56
Rater 1 = 2
Rater 2 = 3

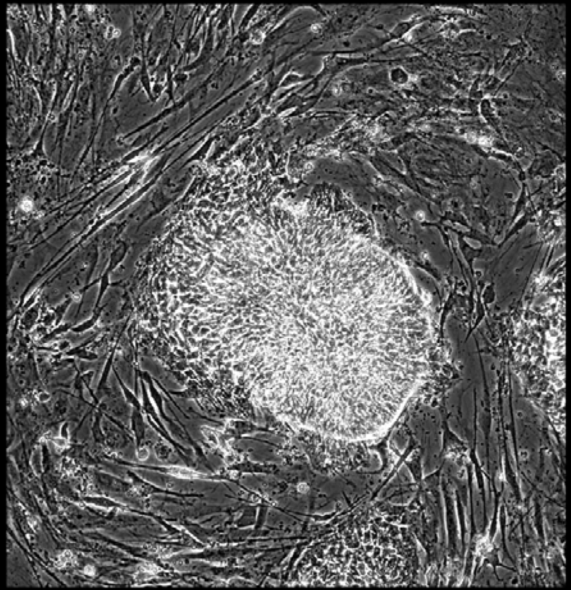

Well = 3, Colony = 57
Rater 1 = 4
Rater 2 = not scored

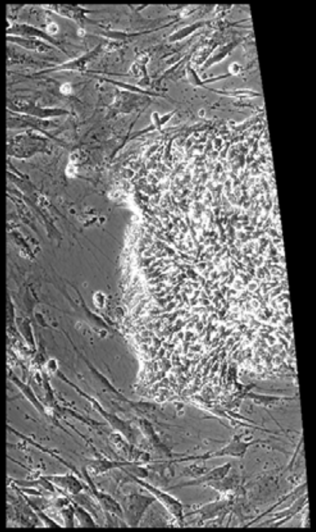

Well = 3, Colony = 58
Rater 1 = 5
Rater 2 = 5

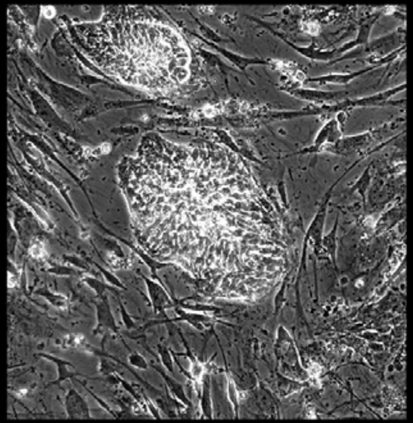

Well = 3, Colony = 59
Rater 1 = 2
Rater 2 = 2

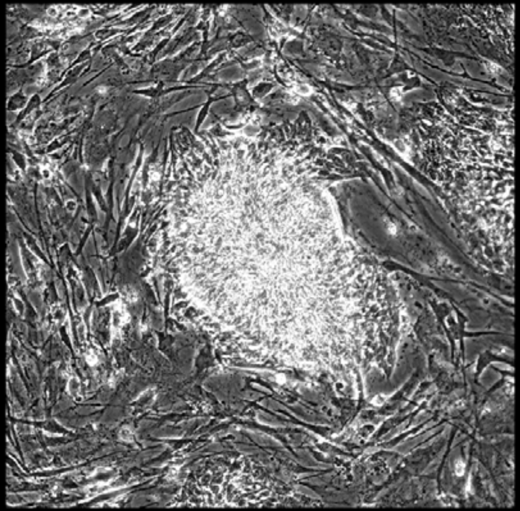

Well = 3, Colony = 60
Rater 1 = 2
Rater 2 = 3

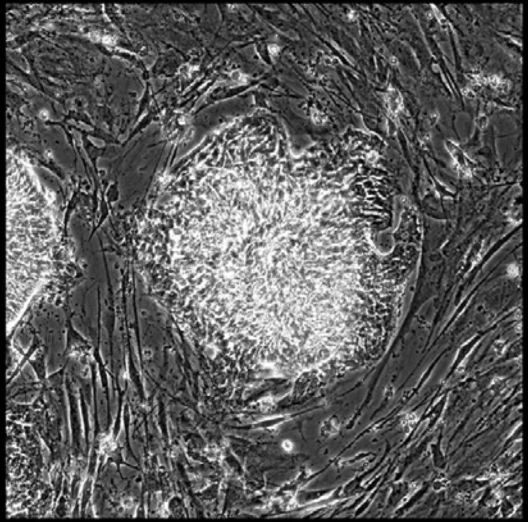

Well = 3, Colony = 61
Rater 1 = 4
Rater 2 = 4

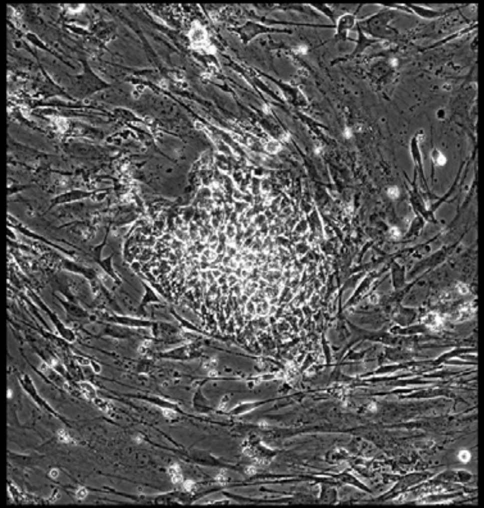

Well = 3, Colony = 62
Rater 1 = 4
Rater 2 = 4

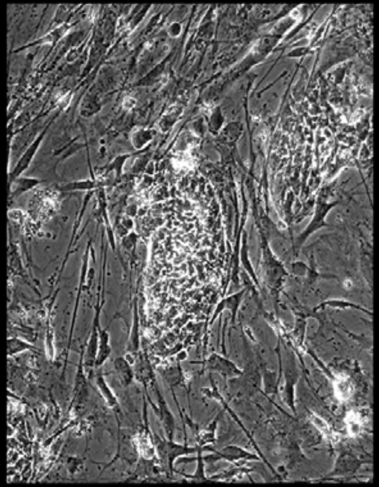

Well = 3, Colony = 63
Rater 1 = 3
Rater 2 = 5

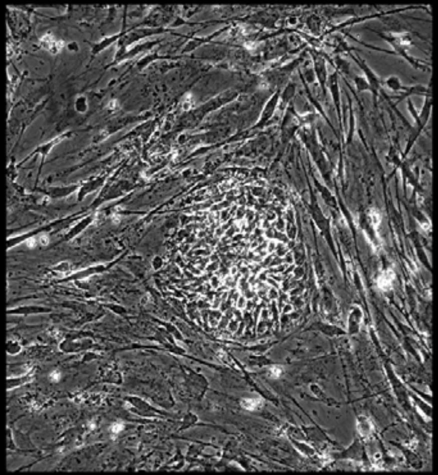

Well = 3, Colony = 64
Rater 1 = 4
Rater 2 = 5

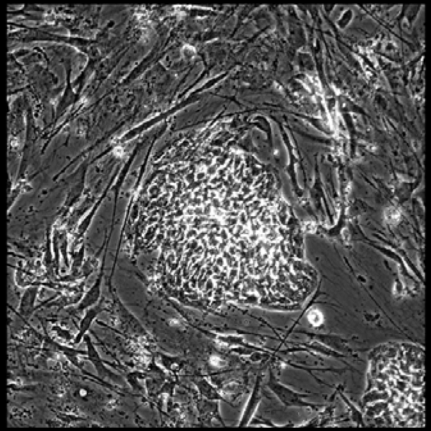

Well = 3, Colony = 65
Rater 1 = 4
Rater 2 = 3

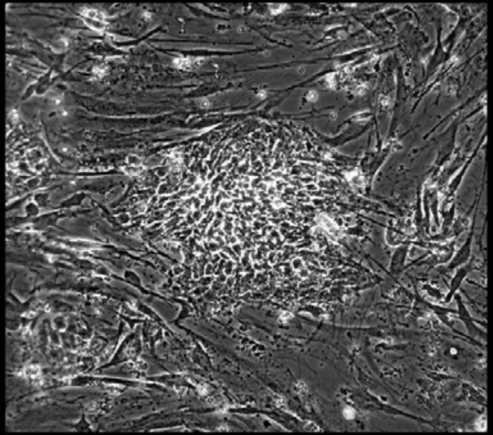

Well = 3, Colony = 66
Rater 1 = 3
Rater 2 = 3

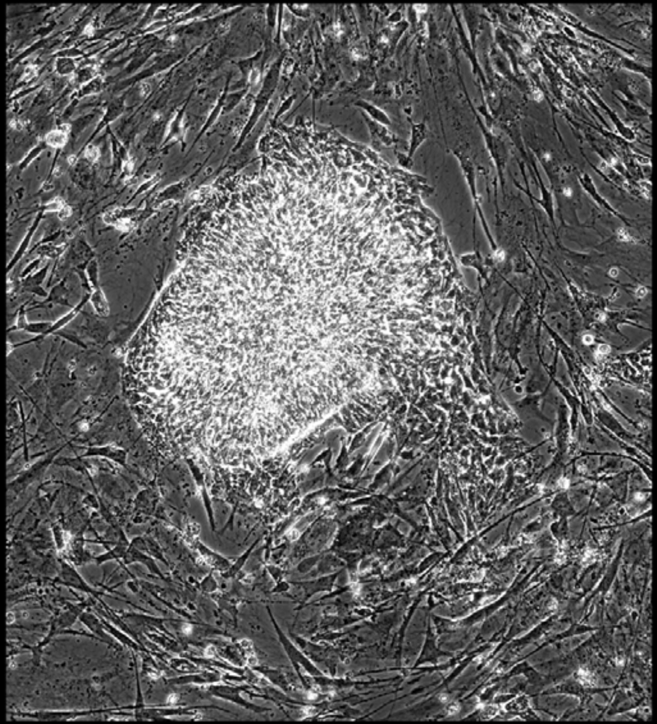

Well = 3, Colony = 67
Rater 1 = 4
Rater 2 = 4

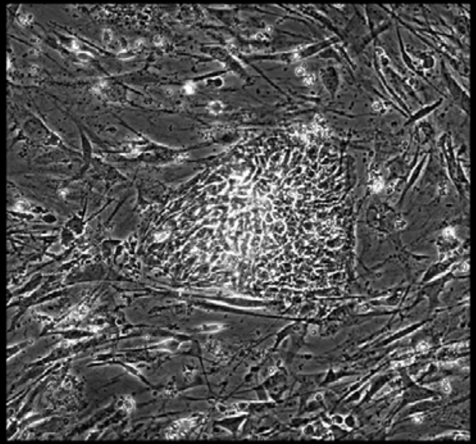

Well = 3, Colony = 68
Rater 1 = 5
Rater 2 = 4

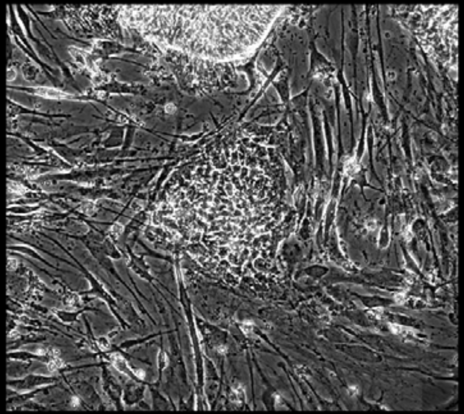

Well = 3, Colony = 69
Rater 1 = 2
Rater 2 = 3

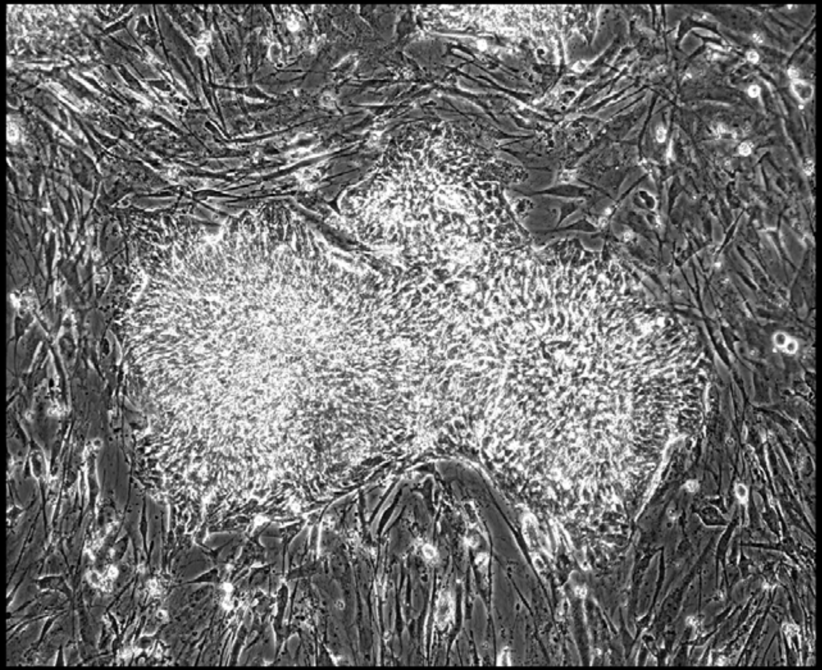

Well = 3, Colony = 70
Rater 1 = 4
Rater 2 = 5

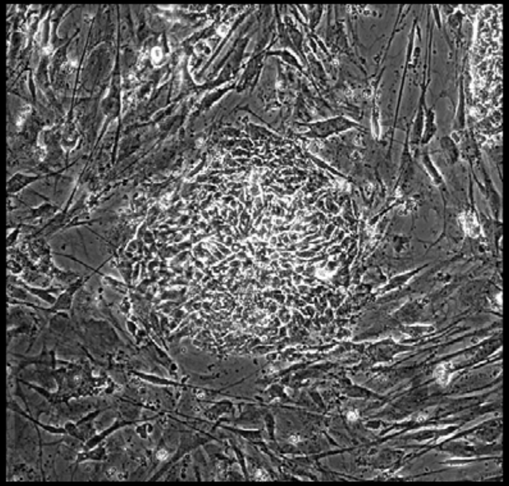

Well = 3, Colony = 71
Rater 1 = 4
Rater 2 = 4

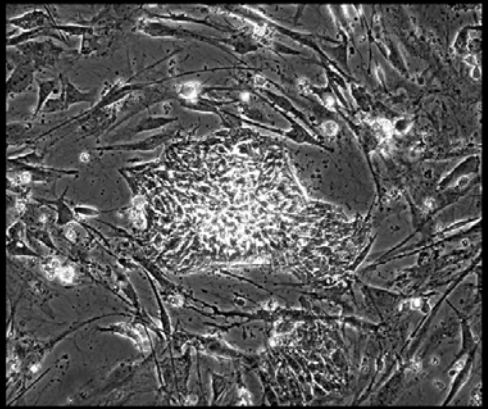

Well = 3, Colony = 72
Rater 1 = 3
Rater 2 = 4

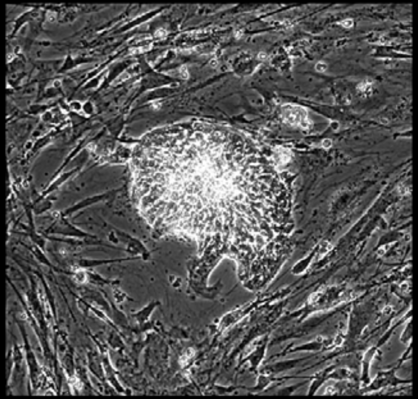

Well = 3, Colony = 73
Rater 1 = 2
Rater 2 = 3

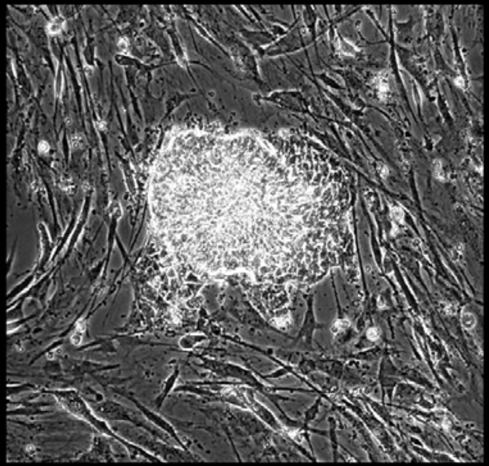

Well = 3, Colony = 74
Rater 1 = 4
Rater 2 = 4

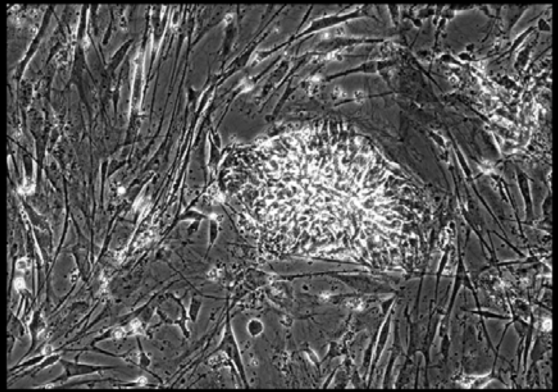

Well = 3, Colony = 75
Rater 1 = 3
Rater 2 = 3

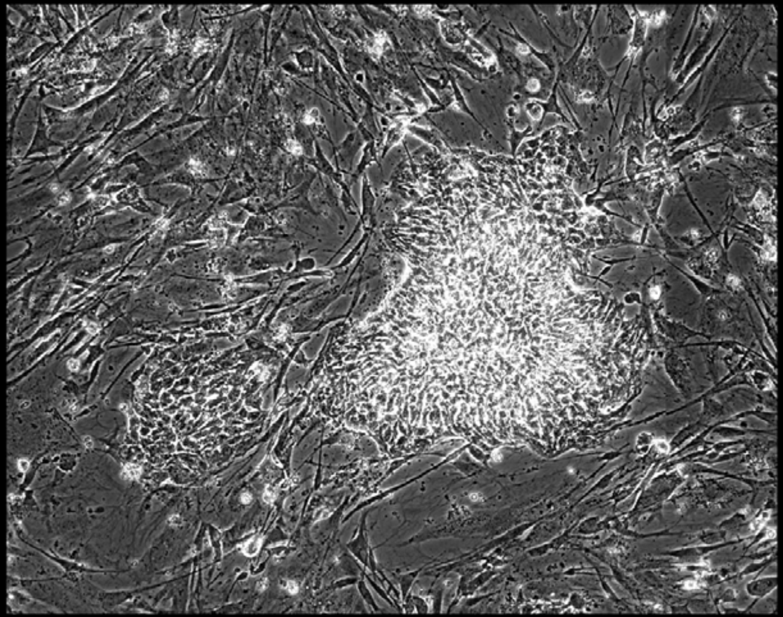

Well = 3, Colony = 76
Rater 1 = 3
Rater 2 = not scored

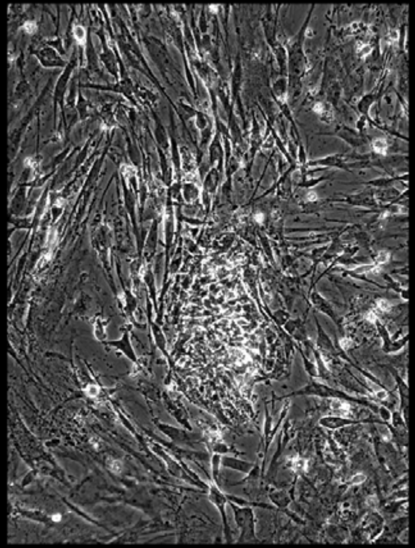

Well = 3, Colony = 77
Rater 1 = 4
Rater 2 = 5

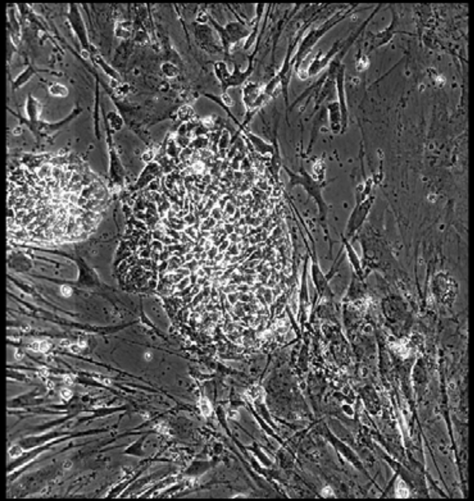

Well = 3, Colony = 78
Rater 1 = 4
Rater 2 = 4

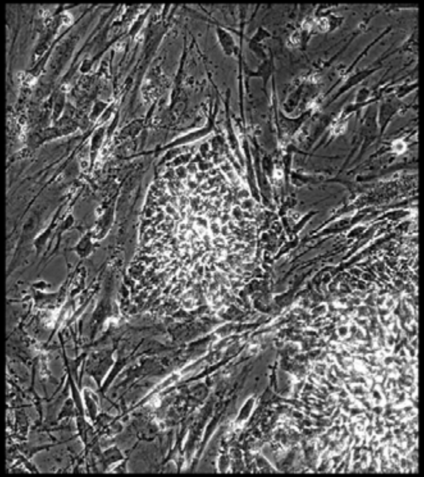

Well = 3, Colony = 79
Rater 1 = 3
Rater 2 = 2

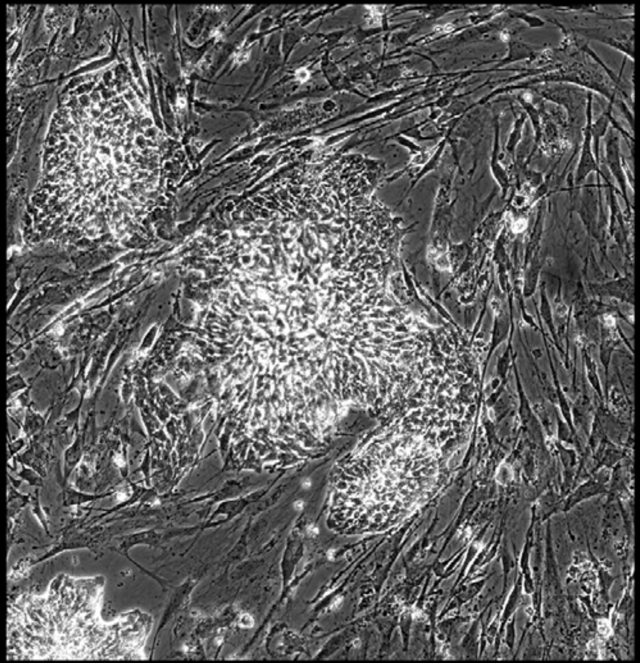

Well = 3, Colony = 80
Rater 1 = 3
Rater 2 = 3

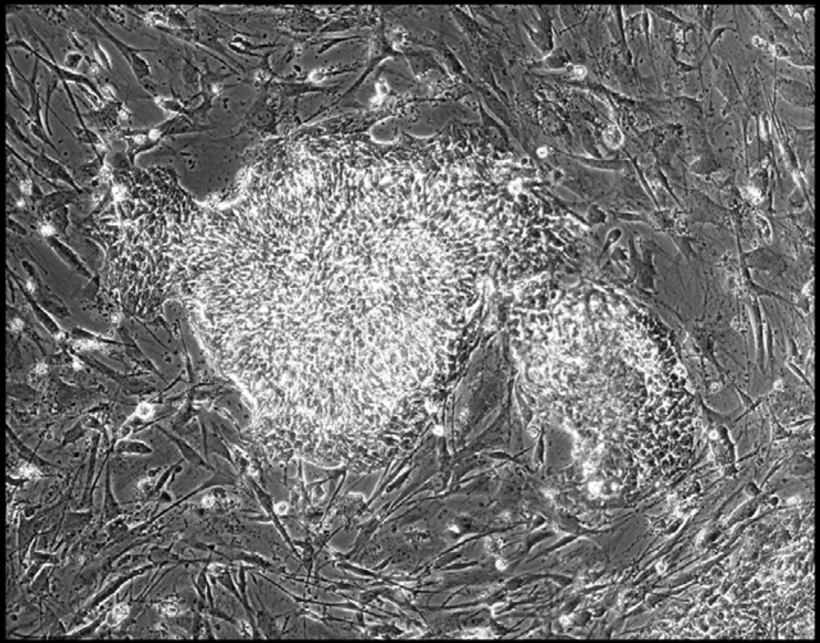

Well = 3, Colony = 81
Rater 1 = 1
Rater 2 = 2

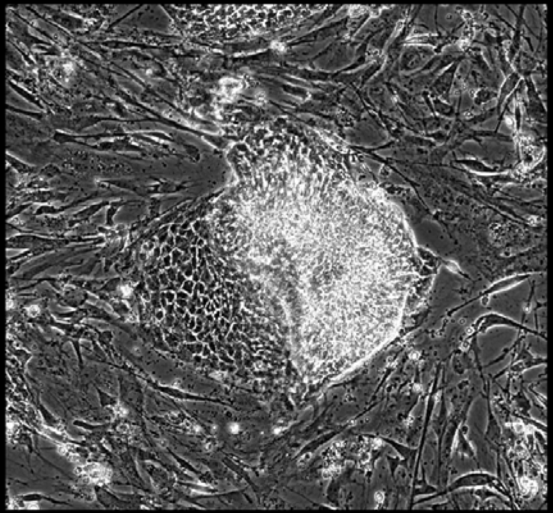

Well = 3, Colony = 82
Rater 1 = 3
Rater 2 = 5

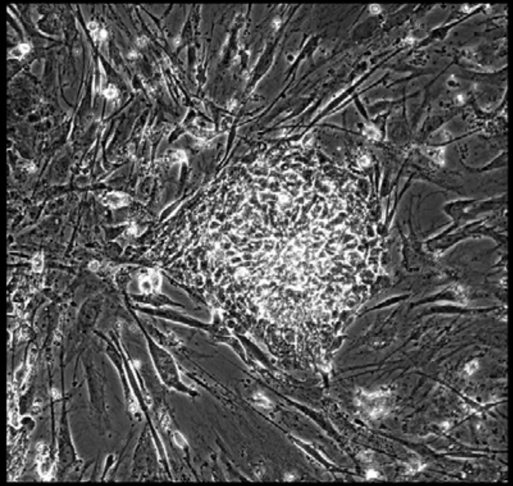

Well = 3, Colony = 83
Rater 1 = 4
Rater 2 = 5

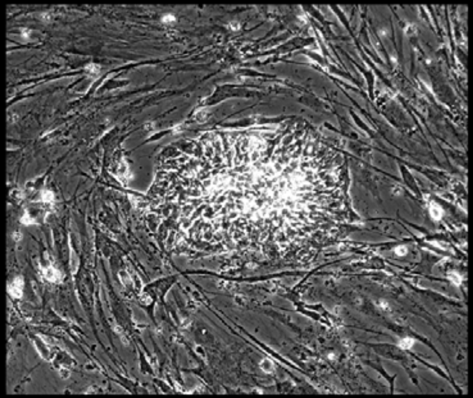

Well = 3, Colony = 84
Rater 1 = 2
Rater 2 = 3

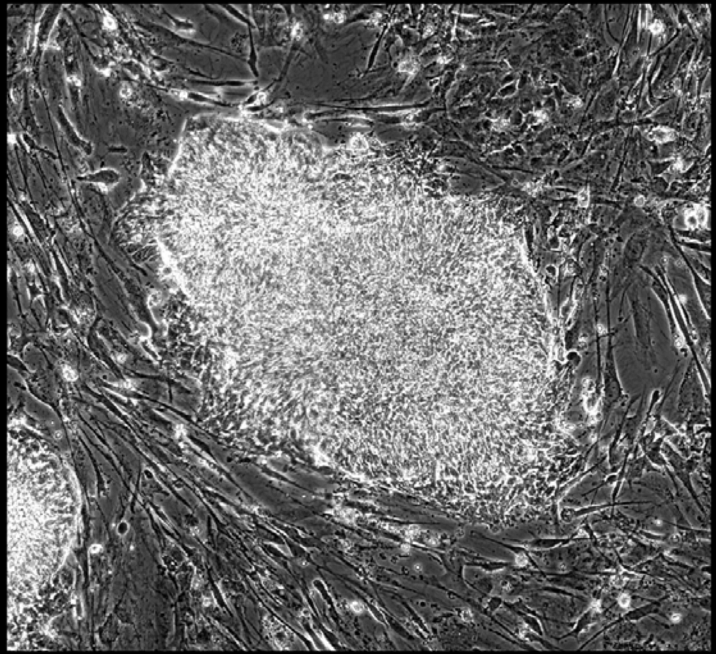

Well = 3, Colony = 85
Rater 1 = 4
Rater 2 = 4

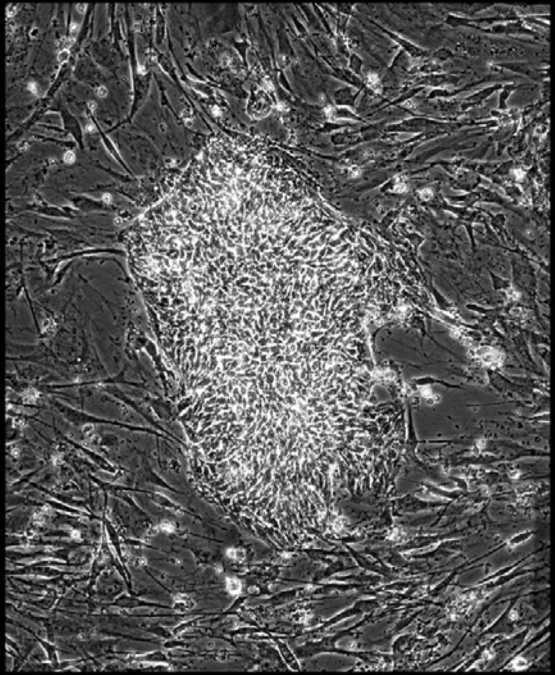

Well = 3, Colony = 86
Rater 1 = 3
Rater 2 = 4

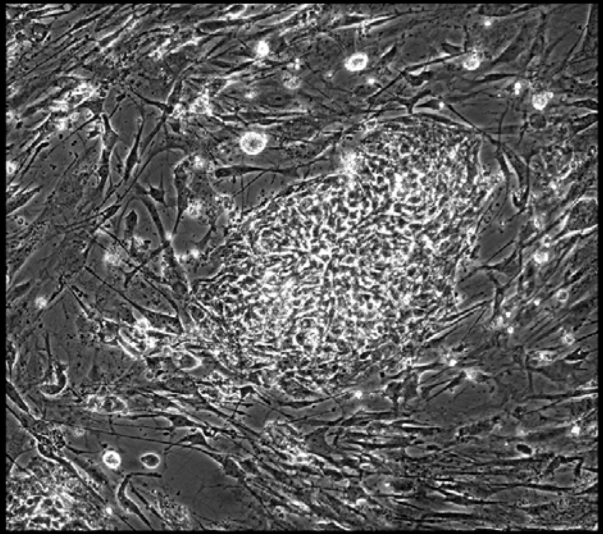

Well = 3, Colony = 87
Rater 1 = 2
Rater 2 = 3

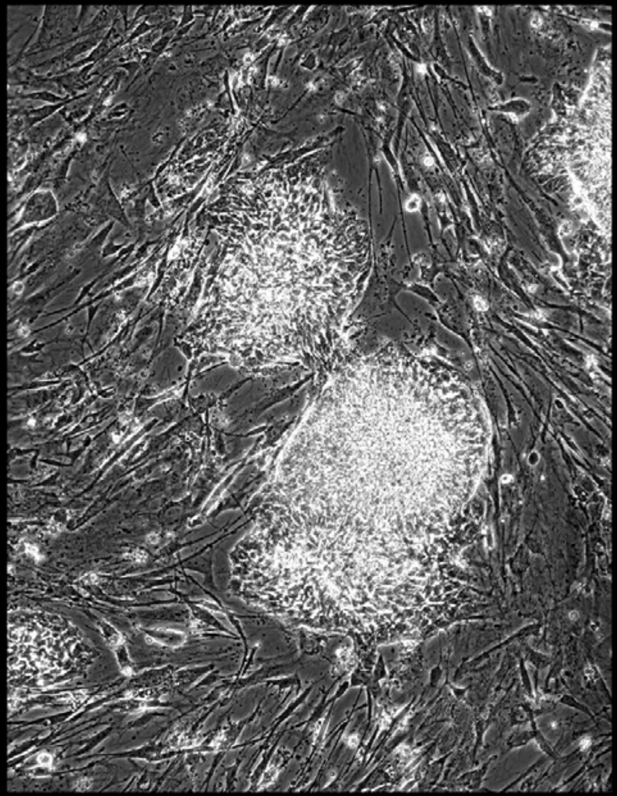

Well = 3, Colony = 88
Rater 1 = 3
Rater 2 = 3

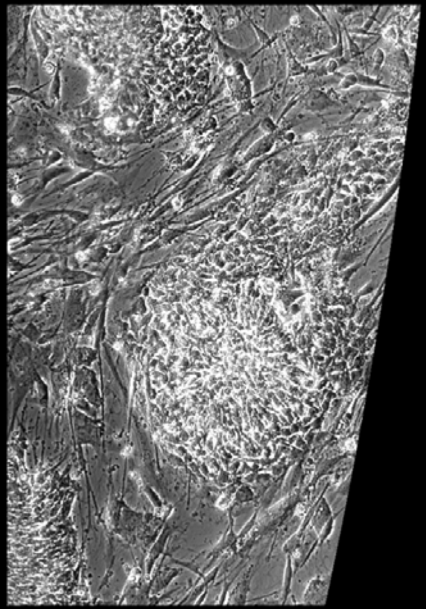

Well = 3, Colony = 89
Rater 1 = 3
Rater 2 = 4

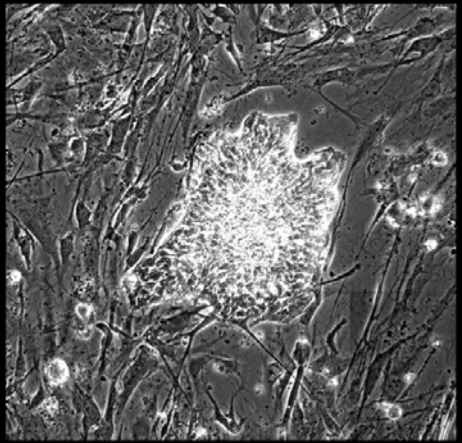

Well = 3, Colony = 90
Rater 1 = 4
Rater 2 = 4

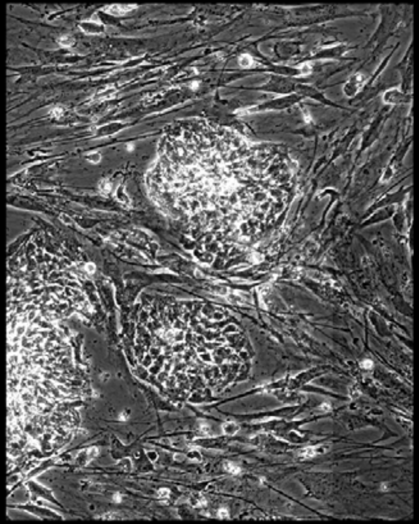

Well = 3, Colony = 91
Rater 1 = 4
Rater 2 = 4

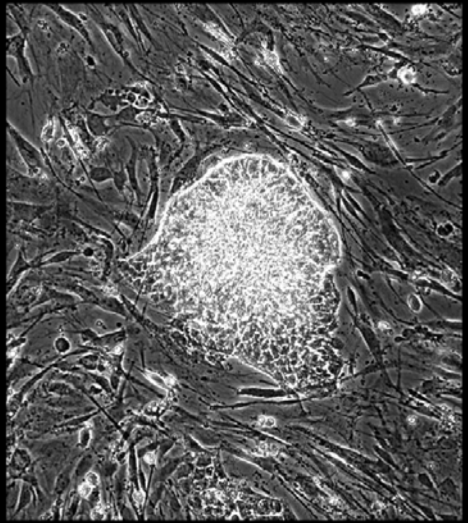

Well = 3, Colony = 92
Rater 1 = 2
Rater 2 = 3

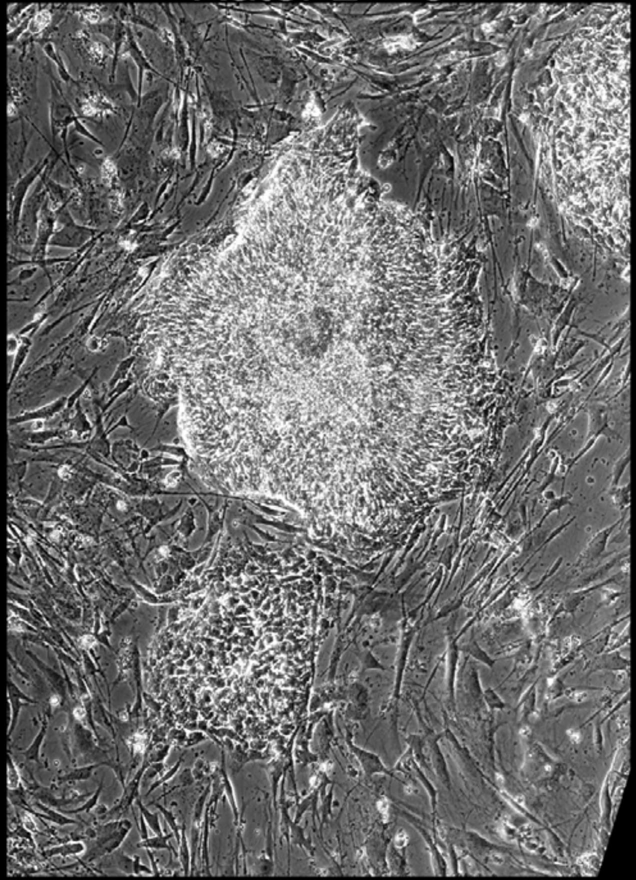

Well = 3, Colony = 93
Rater 1 = 5
Rater 2 = 5

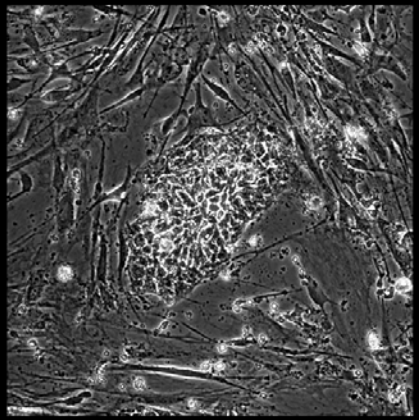

Well = 3, Colony = 94
Rater 1 = 3
Rater 2 = 5

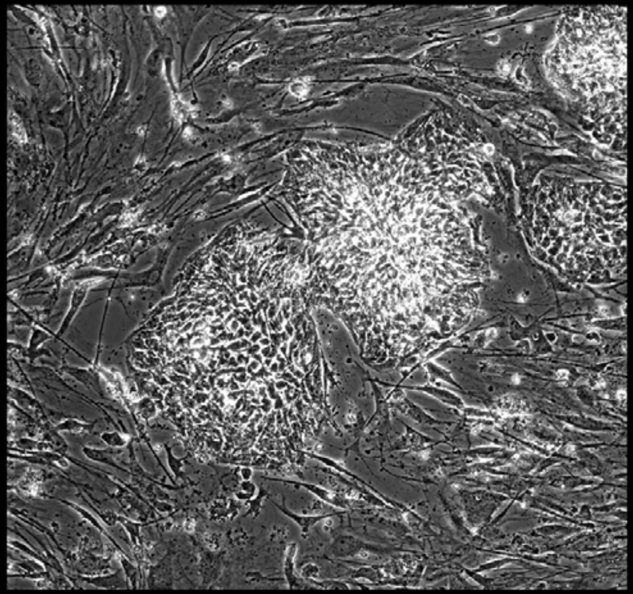

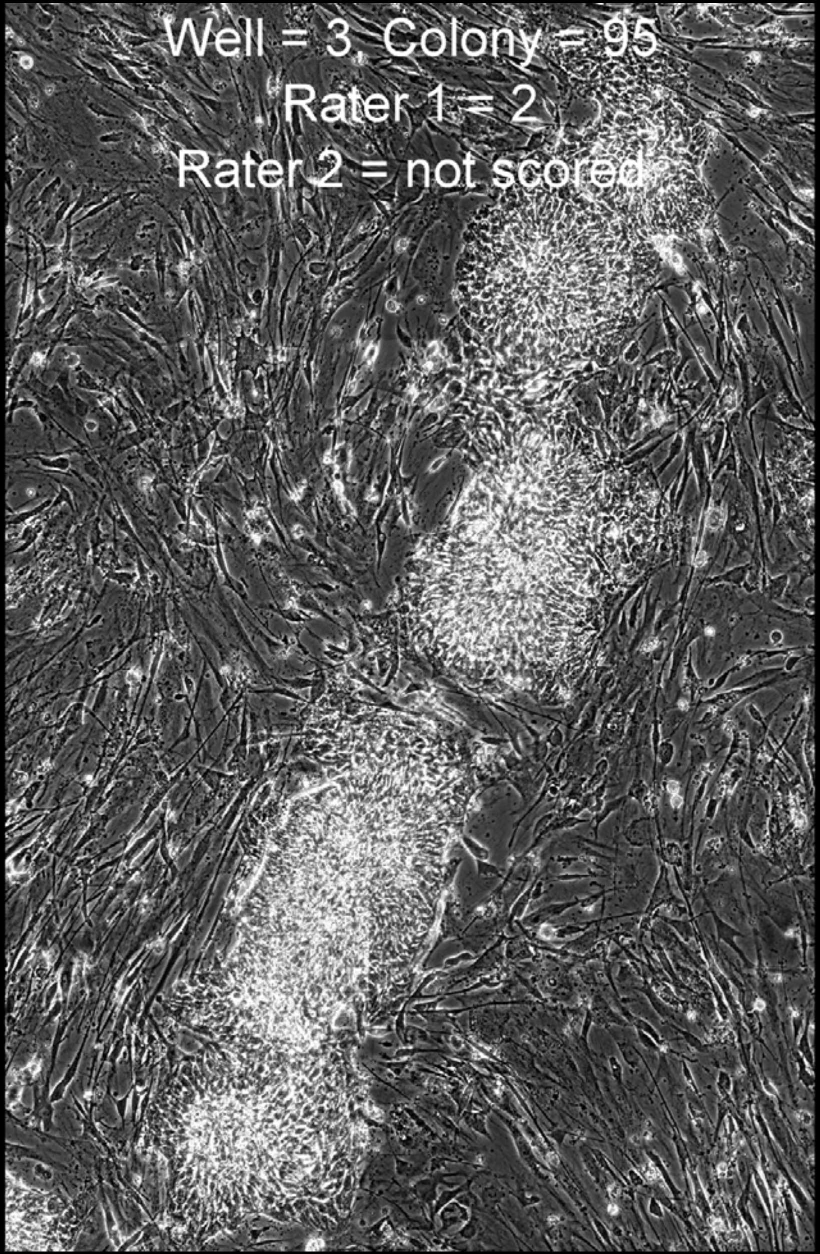


Well = 3, Colony = 96
Rater 1 = 4
Rater 2 = 5

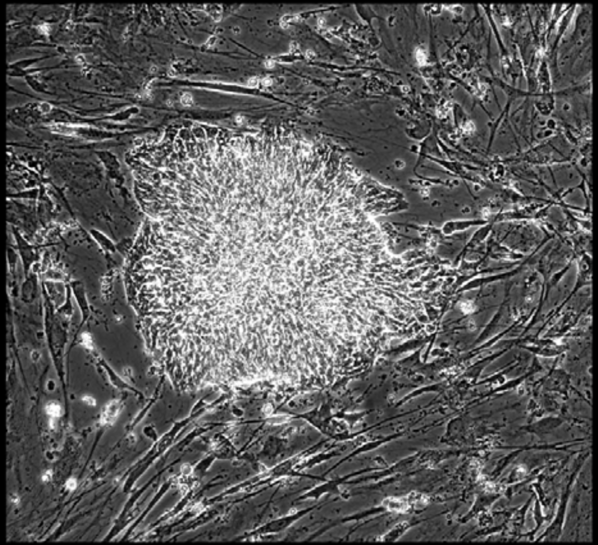

Well = 3, Colony = 97
Rater 1 = 3
Rater 2 = not scored

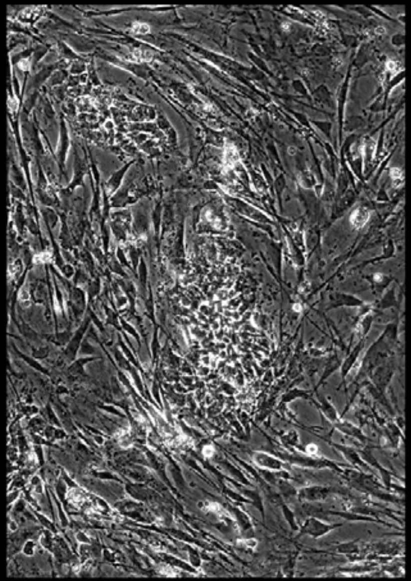

Well = 3, Colony = 98
Rater 1 = 2
Rater 2 = 4

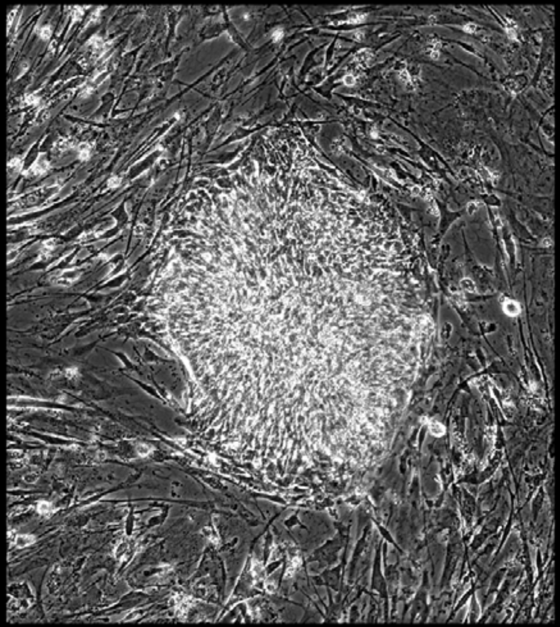

Well = 3, Colony = 99
Rater 1 = 4
Rater 2 = 5

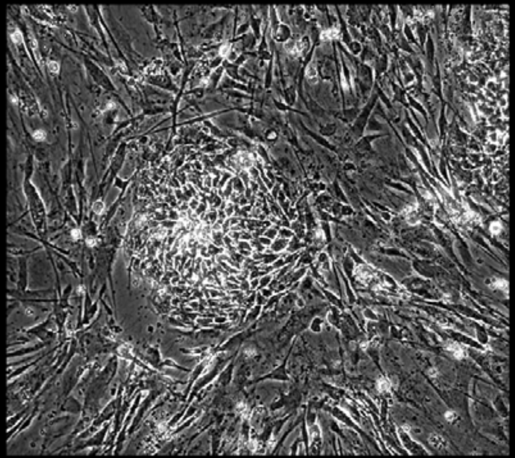

Well = 3, Colony = 100
Rater 1 = 2
Rater 2 = not scored

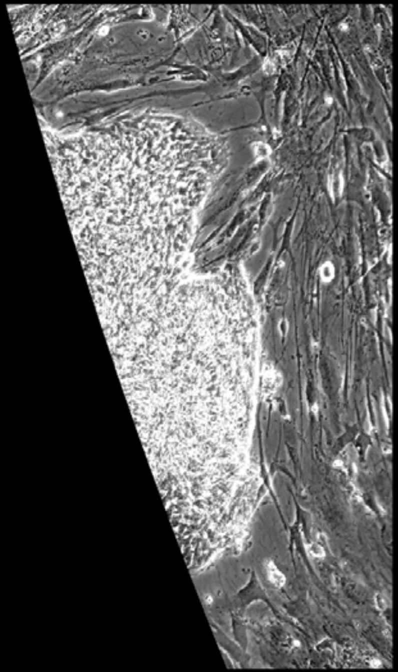

Well = 3, Colony = 101
Rater 1 = 1
Rater 2 = 1

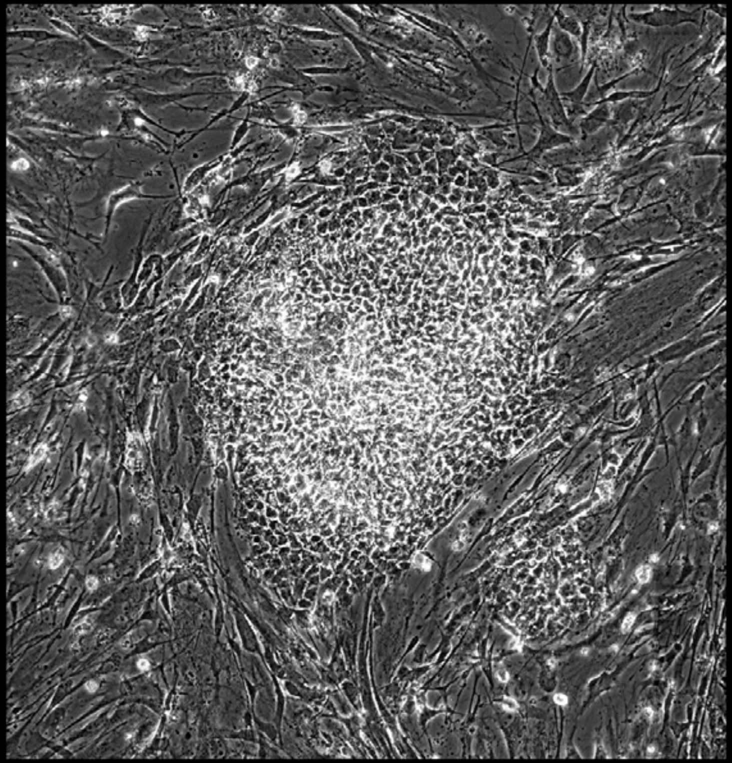

Well = 3, Colony = 102
Rater 1 = 4
Rater 2 = 5

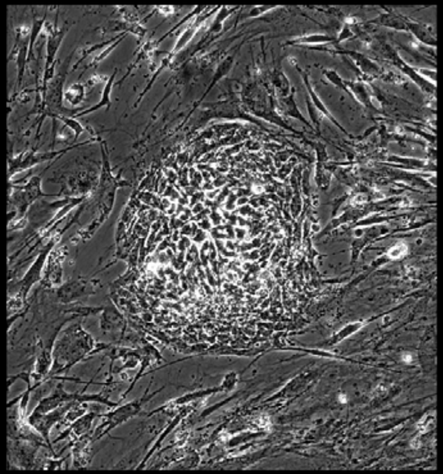

Well = 3, Colony = 103
Rater 1 = 3
Rater 2 = 3

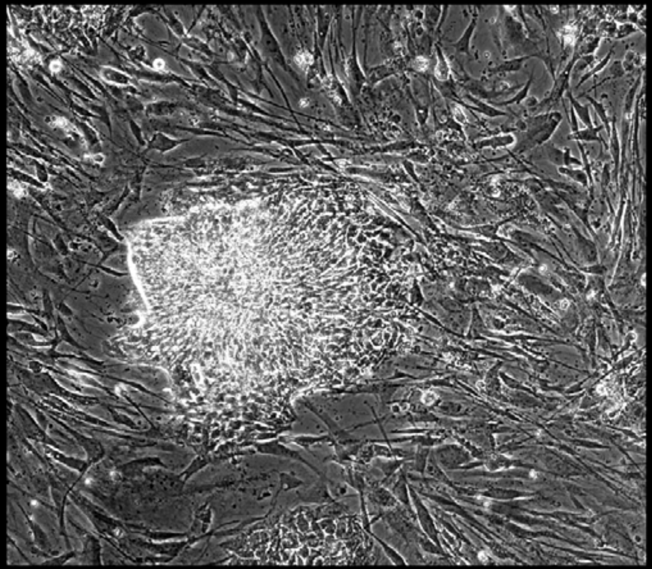

Well = 3, Colony = 104
Rater 1 = 3
Rater 2 = 5

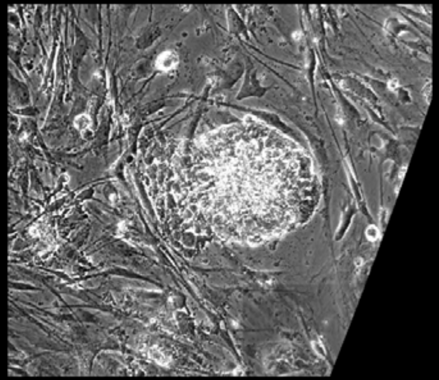

Well = 3, Colony = 105
Rater 1 = 4
Rater 2 = 4

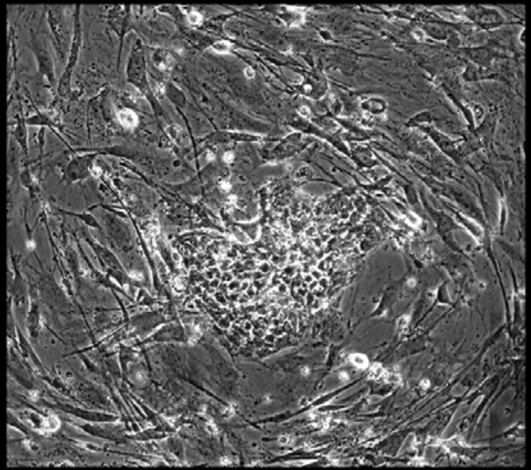

Well = 3, Colony = 106
Rater 1 = 3
Rater 2 = 3

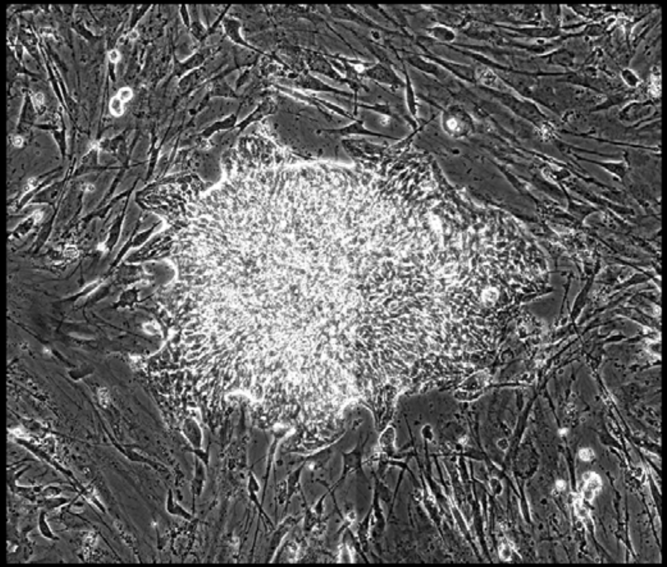

Well = 3, Colony = 107
Rater 1 = 2
Rater 2 = 4

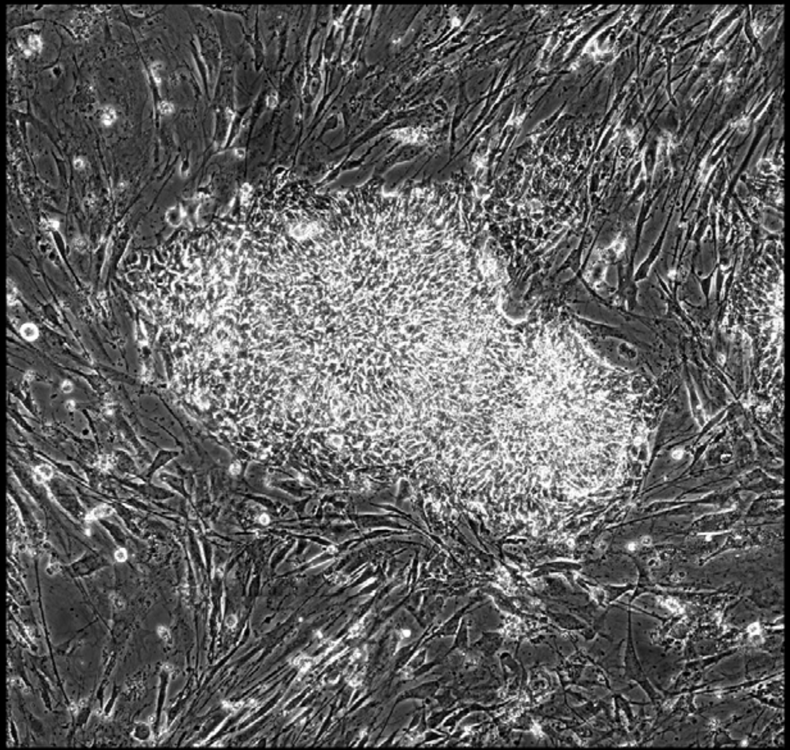

Well = 3, Colony = 108
Rater 1 = 3
Rater 2 = 5

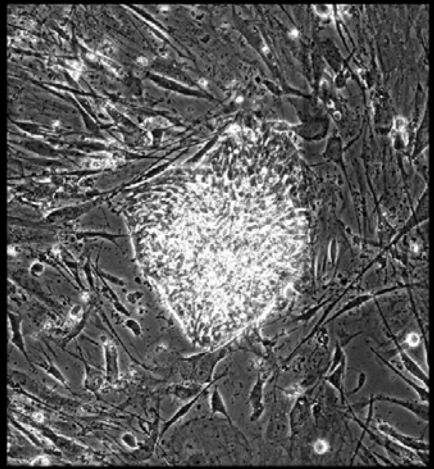

Well = 3, Colony = 109
Rater 1 = 3
Rater 2 = 5

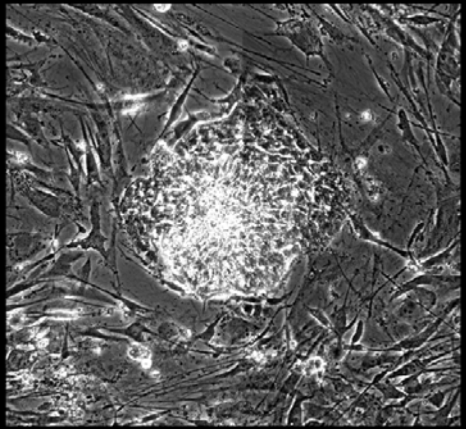

Well = 3, Colony = 110
Rater 1 = 2
Rater 2 = 2

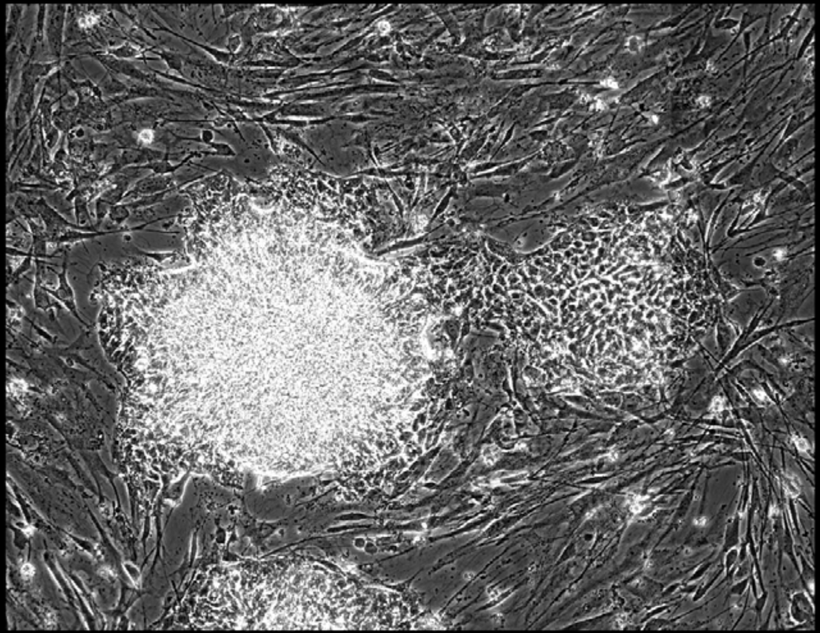

Well = 3, Colony = 111
Rater 1 = 2
Rater 2 = not scored

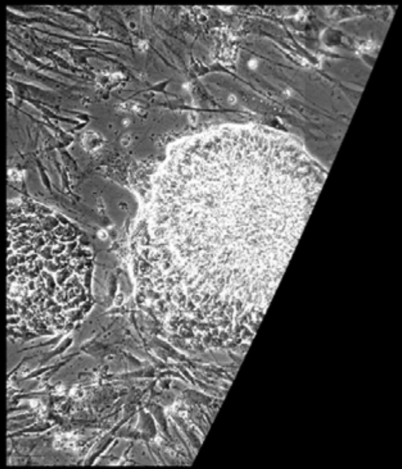

Well = 3, Colony = 112
Rater 1 = 5
Rater 2 = 5

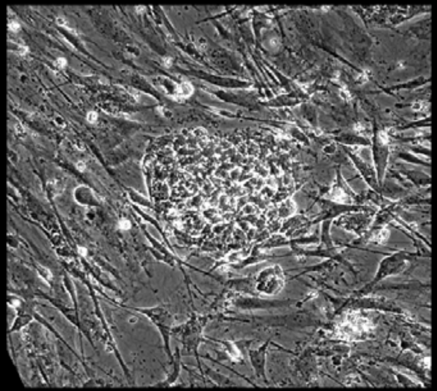

Well = 3, Colony = 113
Rater 1 = 4
Rater 2 = 5

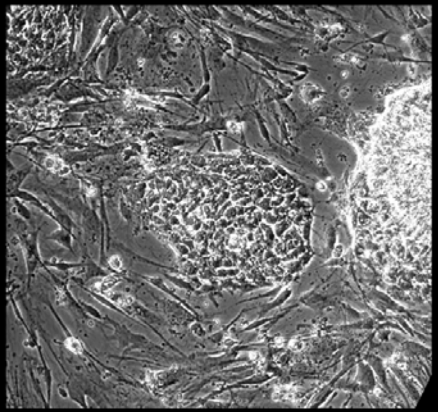

Well = 3, Colony = 114
Rater 1 = 4
Rater 2 = 4

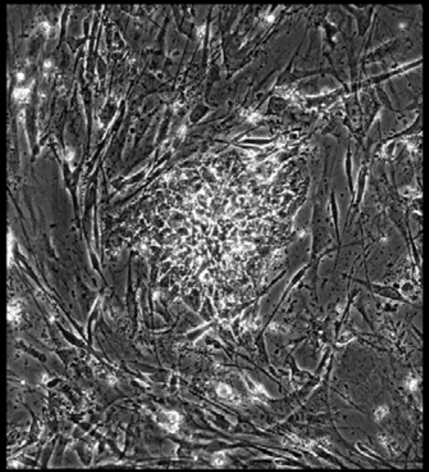

Well = 3, Colony = 115
Rater 1 = 2
Rater 2 = 4

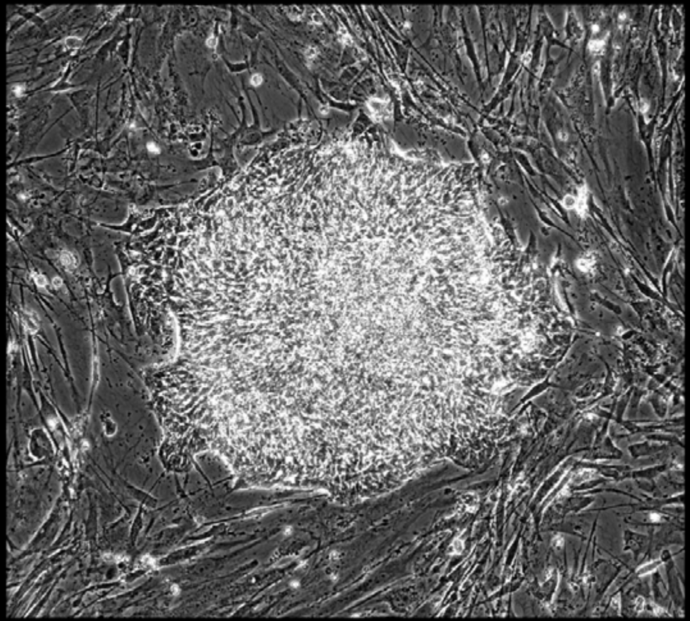

Well = 3, Colony = 116
Rater 1 = 4
Rater 2 = 4

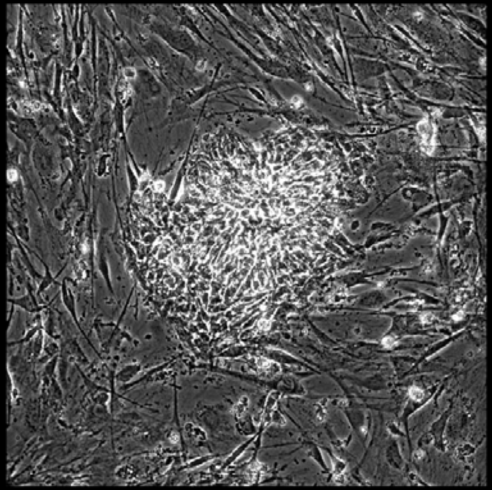

Well = 3, Colony = 117
Rater 1 = 2
Rater 2 = 4

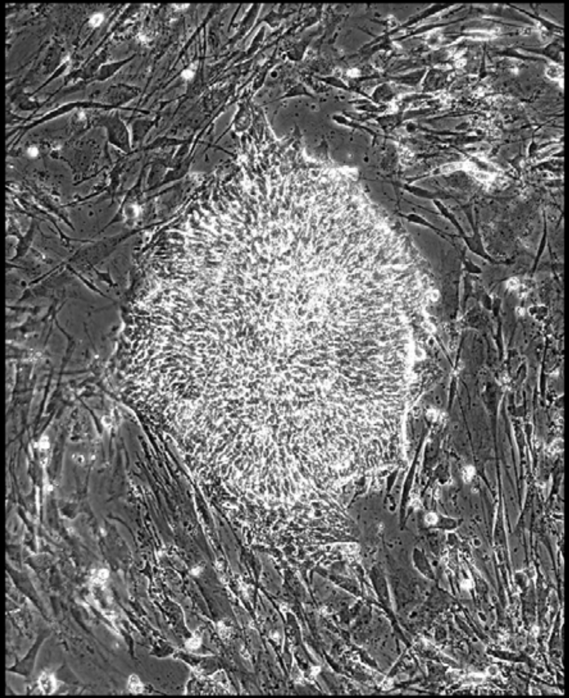

Well = 3, Colony = 118
Rater 1 = 4
Rater 2 = 5

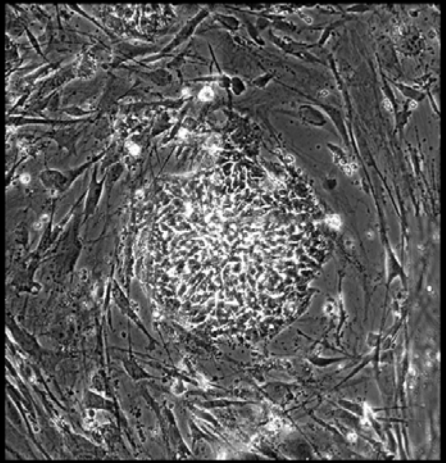

Well = 3, Colony = 119
Rater 1 = 2
Rater 2 = 3

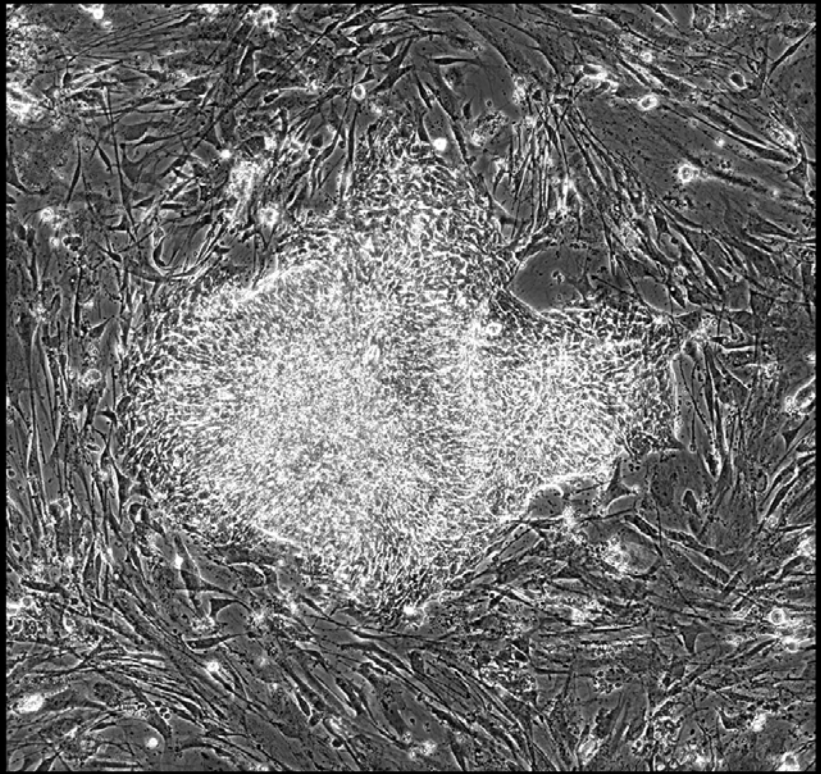

Well = 3, Colony = 120
Rater 1 = 3
Rater 2 = 2

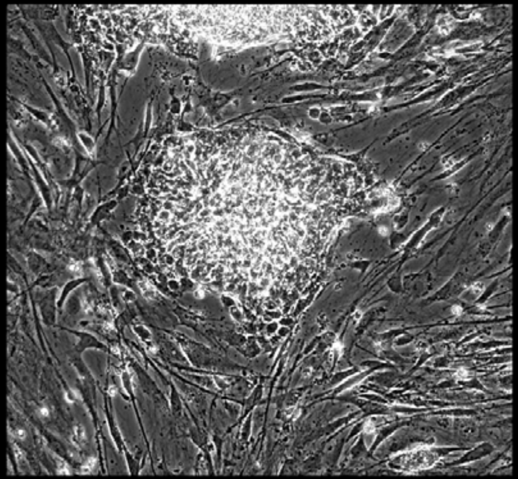

Well = 3, Colony = 121
Rater 1 = 4
Rater 2 = 4

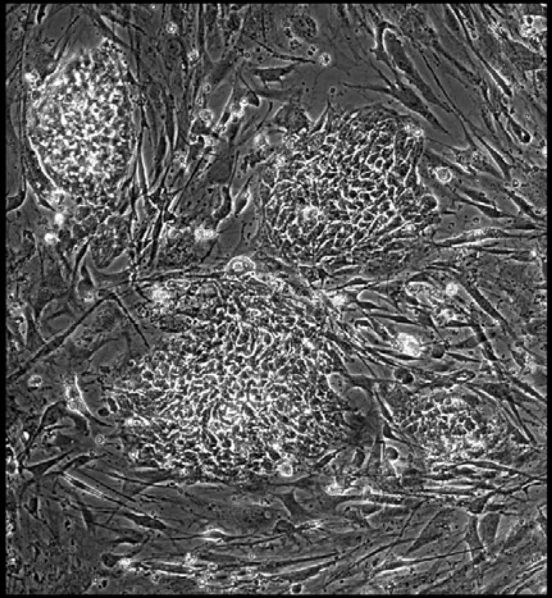

Well = 3, Colony = 124
Rater 1 = 4
Rater 2 = 5

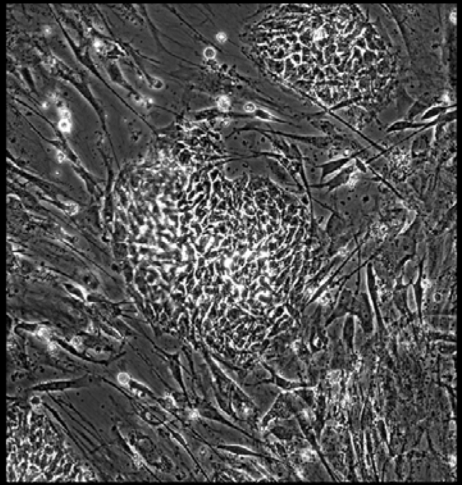

Well = 3, Colony = 125
Rater 1 = 3
Rater 2 = 4

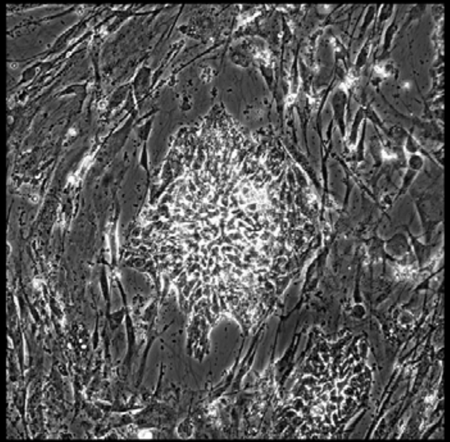

Well = 3, Colony = 126
Rater 1 = 3
Rater 2 = 3

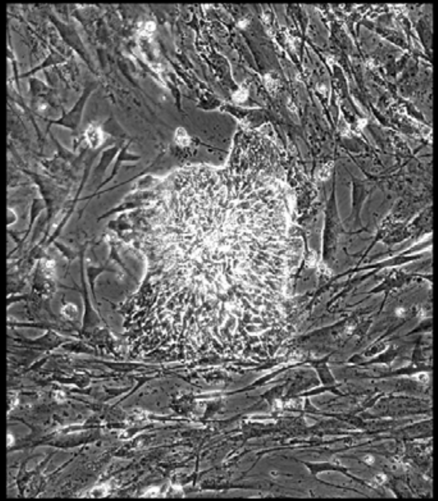

Well = 3, Colony = 127
Rater 1 = 5
Rater 2 = 4

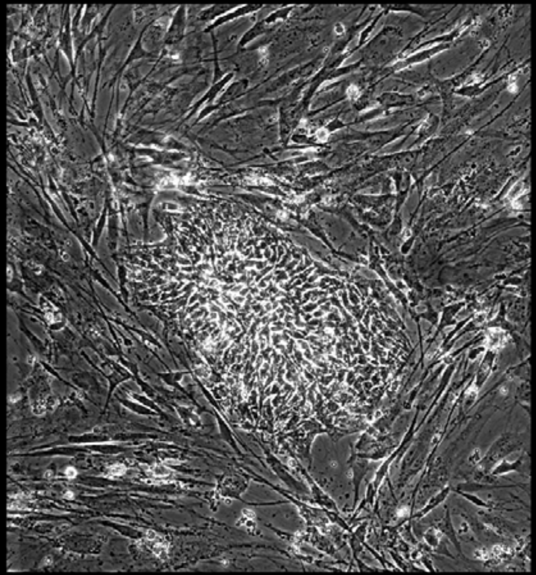

Well = 3, Colony = 128
Rater 1 = 3
Rater 2 = 4

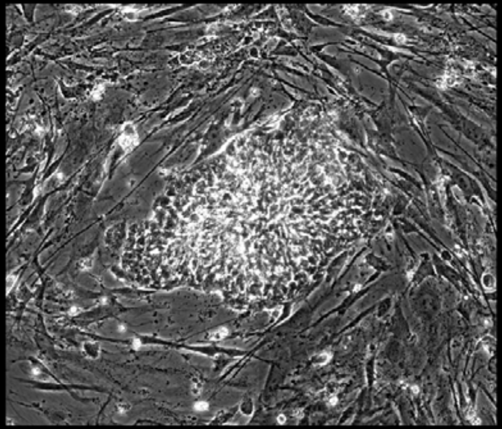

Well = 3, Colony = 129
Rater 1 = 3
Rater 2 = 3

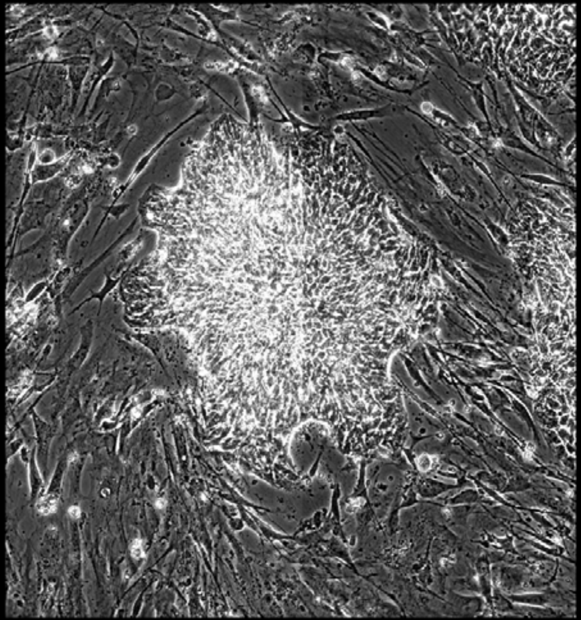

Well = 3, Colony = 130
Rater 1 = 3
Rater 2 = not scored

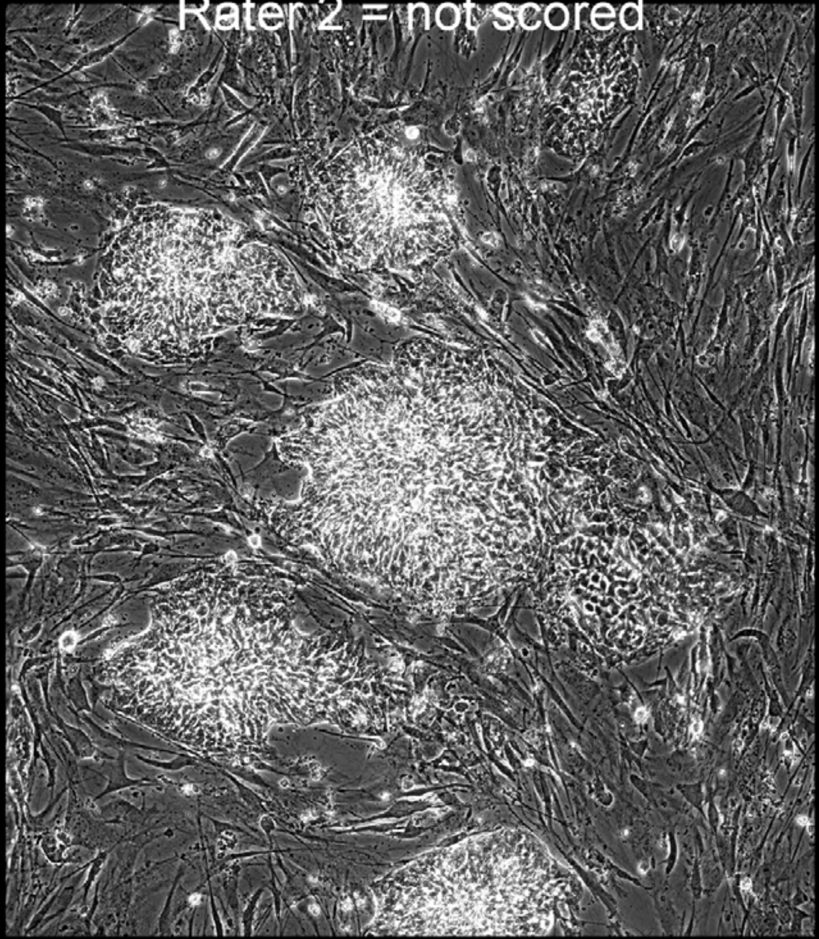

Well = 3, Colony = 131
Rater 1 = 4
Rater 2 = 4

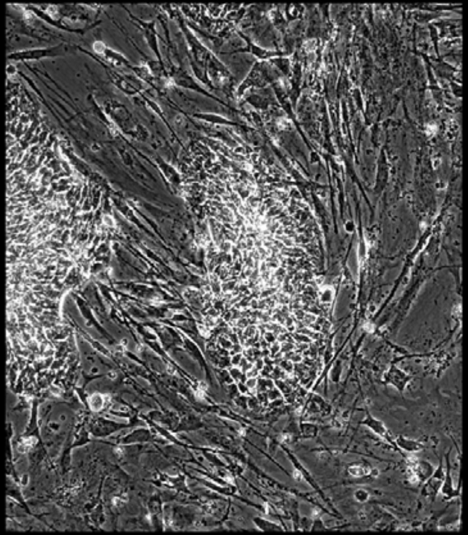

Well = 3, Colony = 132
Rater 1 = 4
Rater 2 = 5

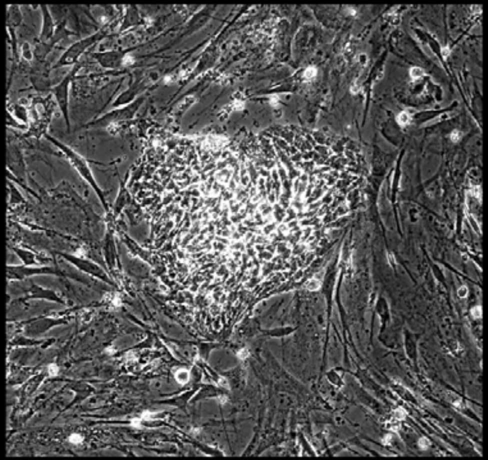

Well = 3, Colony = 133
Rater 1 = 3
Rater 2 = 4

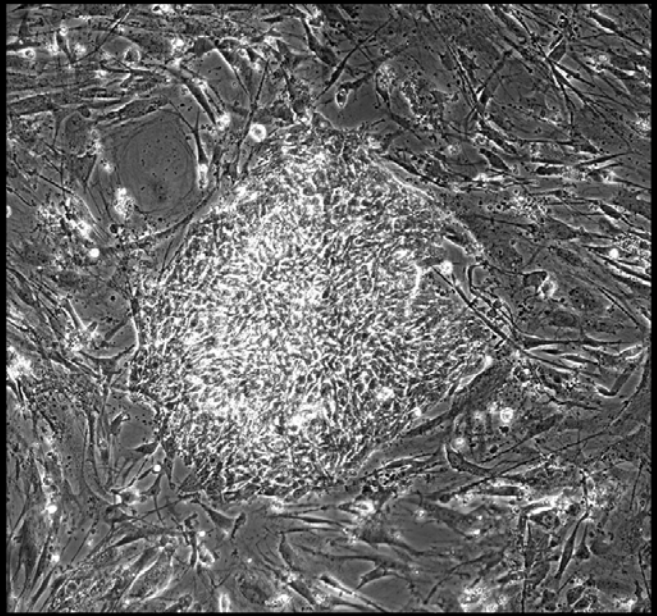

Well = 3, Colony = 134
Rater 1 = 5
Rater 2 = 5

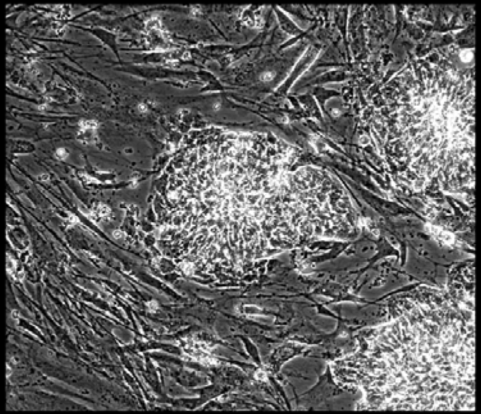

Well = 3, Colony = 135
Rater 1 = 5
Rater 2 = 5

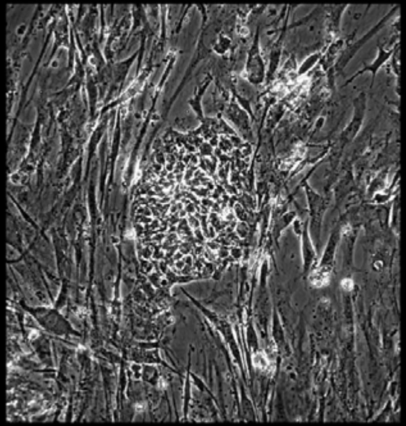

Well = 3, Colony = 136
Rater 1 = 4
Rater 2 = 5

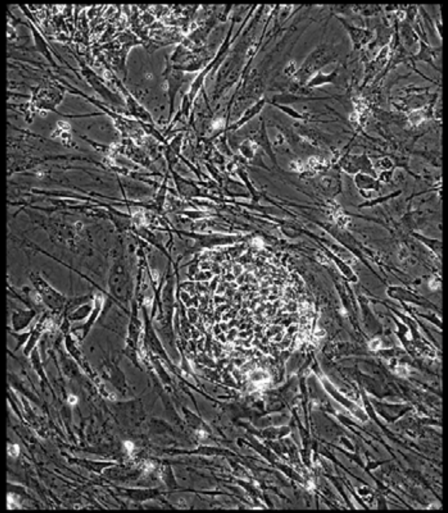

Well = 3, Colony = 137
Rater 1 = 2
Rater 2 = 5

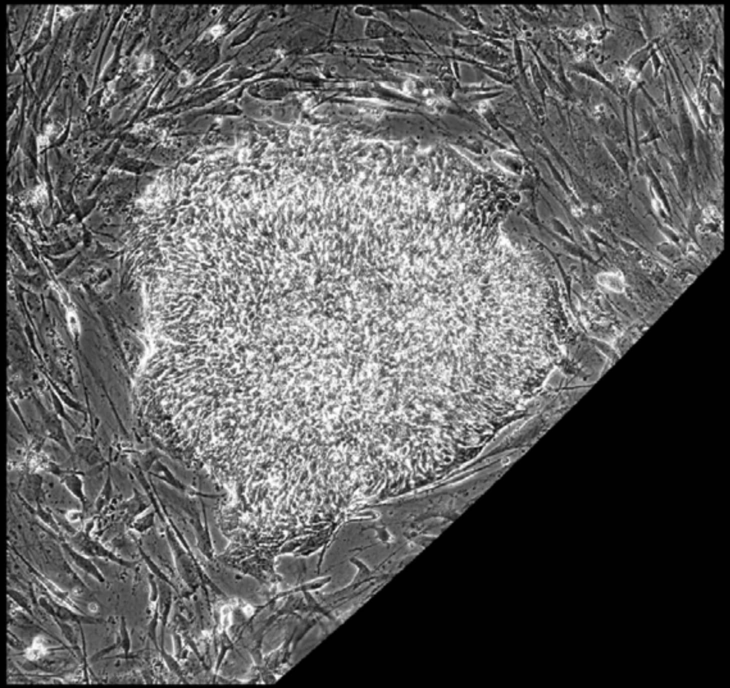

Well = 3, Colony = 138
Rater 1 = 4
Rater 2 = not scored

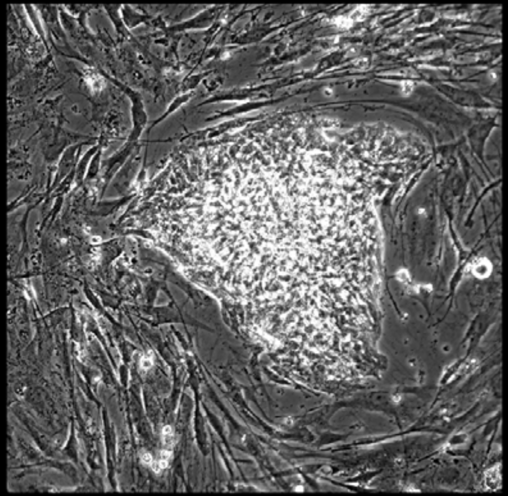

Well = 3, Colony = 139
Rater 1 = 2
Rater 2 = 4

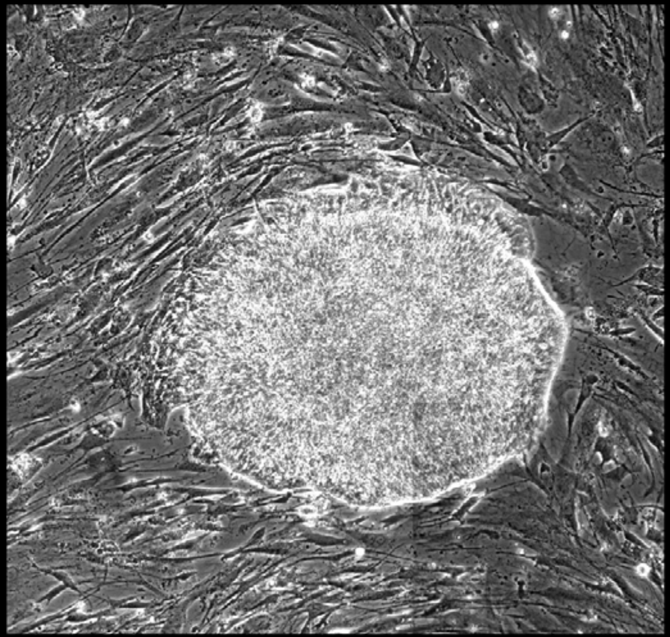

Well = 3, Colony = 140
Rater 1 = 5
Rater 2 = 5

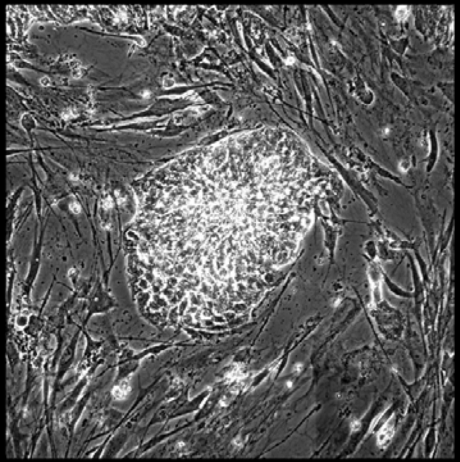

Well = 3, Colony = 141
Rater 1 = 3
Rater 2 = 5

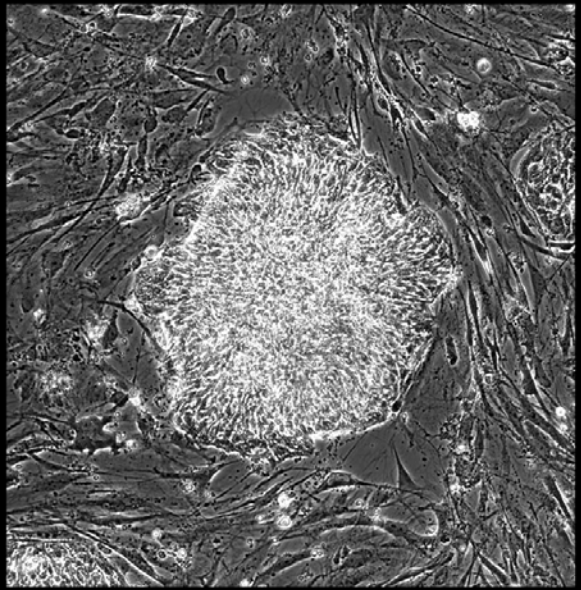

Well = 3, Colony = 142
Rater 1 = 3
Rater 2 = 4

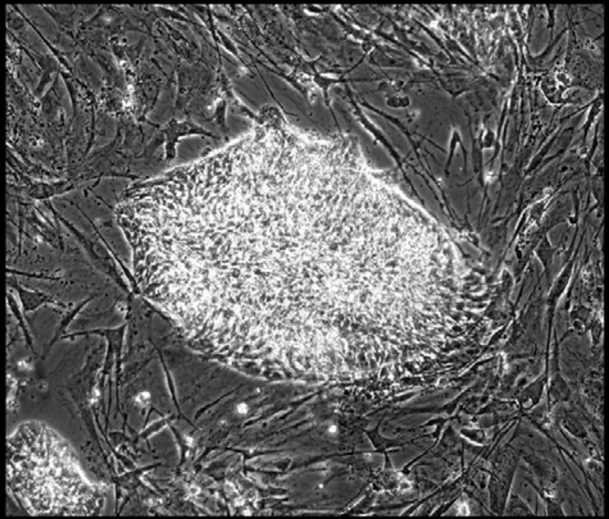

Well = 3, Colony = 143
Rater 1 = 4
Rater 2 = 5

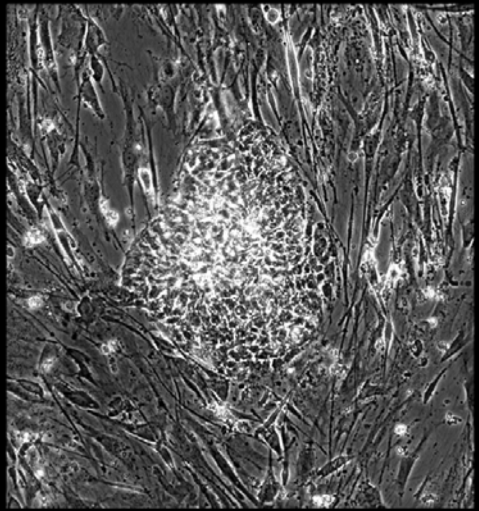

Well = 3, Colony = 144
Rater 1 = 5
Rater 2 = 5

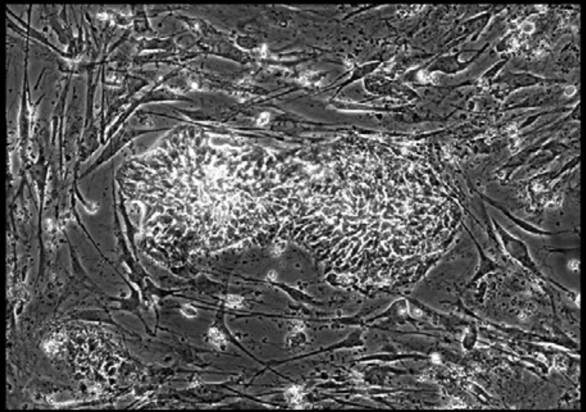

Well = 3, Colony = 145
Rater 1 = 3
Rater 2 = 4

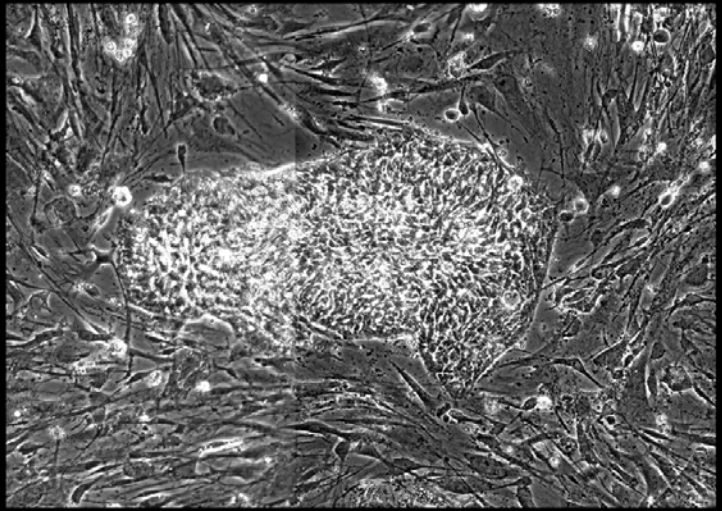

Well = 3, Colony = 146
Rater 1 = 3
Rater 2 = not scored

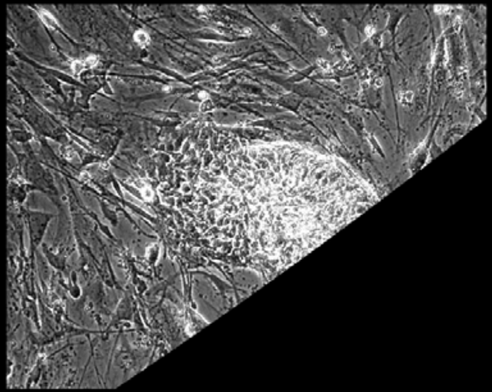

Well = 3, Colony = 147
Rater 1 = 3
Rater 2 = 4

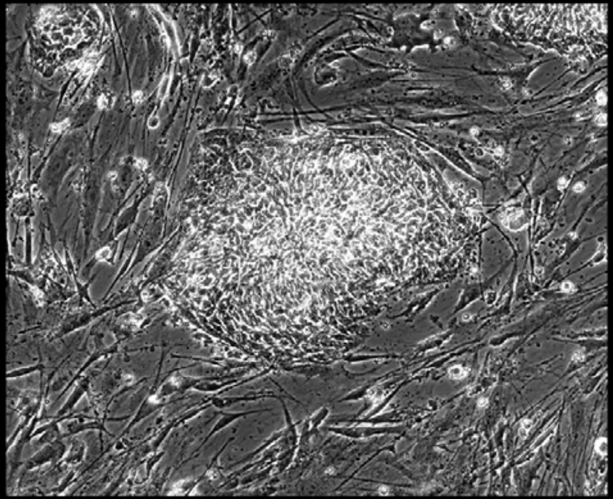

Well = 3, Colony = 148
Rater 1 = 4
Rater 2 = 3

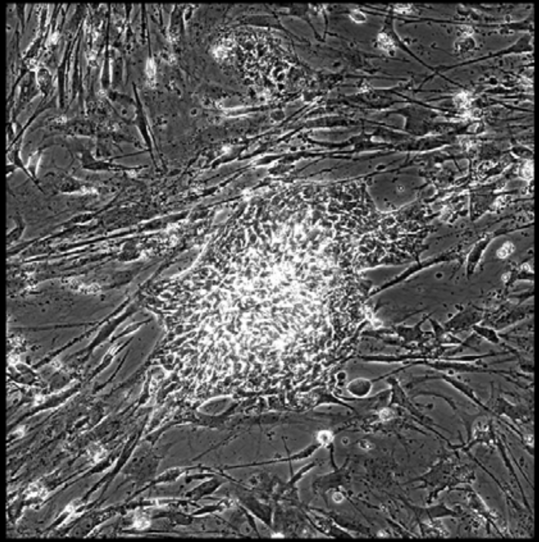
